# In-cell penetration selection—mass spectrometry produces noncanonical peptides for antisense delivery

**DOI:** 10.1101/2022.04.13.488231

**Authors:** Carly K. Schissel, Charlotte E. Farquhar, Andrei Loas, Annika B. Malmberg, Bradley L. Pentelute

**Author notes:** These authors contributed equally to this work (Carly K. Schissel, Charlotte E. Farquhar).

## Abstract

Peptide-mediated delivery of macromolecules in cells has significant potential therapeutic benefits, but no therapy employing cell-penetrating peptides (CPPs) has reached the market after 30 years of investigation due to challenges in the discovery of new, more efficient sequences. We developed a method for in-cell penetration selection-mass spectrometry (in-cell PS-MS) to discover peptides from a synthetic library capable of delivering macromolecule cargo to the cytosol. This method was inspired by recent in vivo selection approaches for cell-surface screening, with an added spatial dimension resulting from subcellular fractionation. A representative peptide discovered in the cytosolic extract, Pep1a, is nearly 100-fold more active toward antisense phosphorodiamidate morpholino oligomer (PMO) delivery compared to a sequence identified from a whole cell extract, which includes endosomes. Pep1a is composed of D-amino acids and two non-α-amino acids. Pulse-chase and microscopy experiments revealed that while the PMO-Pep1a conjugate is likely taken up by endosomes, it can escape to localize to the nucleus. In-cell PS-MS introduces a means to empirically discover unnatural synthetic peptides for subcellular delivery of therapeutically relevant cargo.

## Introduction

After 30 years of investigation, therapies involving cell-penetrating peptides (CPPs) are beginning to advance to late-stage clinical trials.^1,2^ These sequences, composed typically of fewer than 20 amino acids and endowed with diverse physicochemical properties, are able to penetrate the cellular membrane and at times deliver otherwise non-penetrant cargo.^3^ Because of these properties, CPPs have potential applications for the treatment of disease, including cancer, genetic disorders, inflammation, and diabetes. An example is SRP-5051, in which a CPP is covalently linked to an antisense phosphorodiamidate morpholino oligomer (PMO), and is currently under investigation in a Phase 2 clinical trial in patients with Duchenne muscular dystrophy.^4^ Despite these recent advances, to our knowledge no CPP-based therapy has reached the commercial market yet.^5^

While there are several limitations that have slowed the clinical advancement of CPPs, one that we are particularly interested in addressing is the empirical design of novel, more efficient sequences. Historically, CPPs, also known as protein transduction domains, were derived from transmembrane portions of viral and transcriptional proteins. For example, the polyarginine peptide TAT was derived from the HIV-trans-activator of transcription protein and was found to penetrate into the nucleus and target gene expression.^6,7^ From this and similar sequences, synthetic peptides could be designed, including some tailored for delivery of PMO cargo such as Bpep, which relies on a polyarginine sequence to improve uptake and the unnatural residues β-alanine and 6-amino-hexanoic acid to reduce endosomal trapping.^8,9^ Beyond empirical design using derivatives of polyarginine sequences, the rational design of new sequences remains challenging. Methods involving some rational design include synthetic molecular evolution^10,11^ and in silico methods.^12–17^ The latter include our own recent work that leverages machine learning to design new sequences using a model trained with a library tested for the desired activity: nuclear localization.^18–20^ Finally, another common strategy involves screening platforms employing libraries from phage or mRNA display.^21–24^ For example, a screening platform identified several “phylomer” CPPs from bacterial and viral genomes that were then shown to deliver antisense cargo *in vivo*.^25^ Still, a persistent limitation with these approaches is the difficulty of incorporating D-chiral or unnatural amino acids, which would provide access to an augmented chemical space. Unnatural amino acids are more easily incorporated into synthetic one-bead one-compound (OBOC) libraries, although discovery of CPPs by these methods often relies on synthetic vesicles.^26^ Improved screening platforms for discovery of enhanced CPPs could be developed by using unnatural peptides to access greater chemical diversity and proteolytic stability, and by incorporating biologically relevant screening conditions into the protocol, such as in-cell selection and inclusion of the specific cargo to be delivered.

Classic affinity selection involves screening peptide ligands from synthetic libraries (OBOC), phage or mRNA display against immobilized protein targets, and decoding hits.^27,28^ These methods advanced to biologically relevant conditions in on-cell selection platforms for the discovery of new ligands with affinity for the external surface of cells and tissues.^29–31^ Again, biological display techniques are restricted to the use of mostly natural amino acids, limiting the resulting library diversity and proteolytic stability^32,33^, and even those mirror image techniques that allow D-peptide discovery still have difficulty incorporating non-canonical residues.^34,35^ Screening of a synthetic one-bead one-compound (OBOC) library eases the incorporation of non-canonical and D-residues. Our group has recently demonstrated that in vivo affinity selection-mass spectrometry (AS-MS) could identify an erythrocyte-targeting D-peptide.^36^ Such label-free techniques applied to the cell surface can be used to discover novel, non-canonical, D-peptide binders without the addition of display scaffolds or encoding tags.

While most works have focused on affinity screening at the cell surface, there has been some success pushing these techniques to discover peptides that cross the cell membrane. As mentioned, phage display and encoded peptide libraries have been used to discover novel cell penetrating peptides, but these methods have limited advancements for discovery of peptides that deliver cargo to subcellular compartments.^22,25^ Recently, the first example of a DNA-encoded small molecule library screen inside living cells resulted in several chemical motifs that bind the over-expressed protein targets inside oocytes.^37^ This strategy of in-cell selection (rather than on-cell) would be beneficial for discovery of peptides that can deliver macromolecules to the cytosol.

Here we have combined CPP library design and AS-MS selection approaches into a new method: in-cell penetration selection-mass spectrometry (in-cell PS-MS, Fig. 1). Bringing together our expertise in both MS-based selection methodologies and in-cell localization, this technique enables direct recovery of “hit” peptides that deliver a specific type of antisense cargo into the cytosol of cells. Our PS-MS methodology allows the detection of non-canonical peptide-cargo conjugates in-cell, with only the addition of a small biotin handle for extraction. In addition, the PS-MS platform allows additional spatial resolution, separating peptides extracted from whole cell lysates from those extracted from the cytosolic fraction. Resulting peptides are tested in a validation assay selects for peptides that can effectively deliver the cargo to the nucleus, giving an additional layer of spatial resolution. The PMO-CPP library used in the screening demonstrated antisense delivery activity, not necessarily due to a few highly active sequences but due to the activity of the library as a whole. This method led to the discovery of a potent antisense delivery peptide, Pep1a, isolated from the cytosol of cells. This peptide was more active and more efficiently localized to the nucleus compared to peptides that were isolated from the whole cell extracts, which include endosomes.

**Figure 1.**
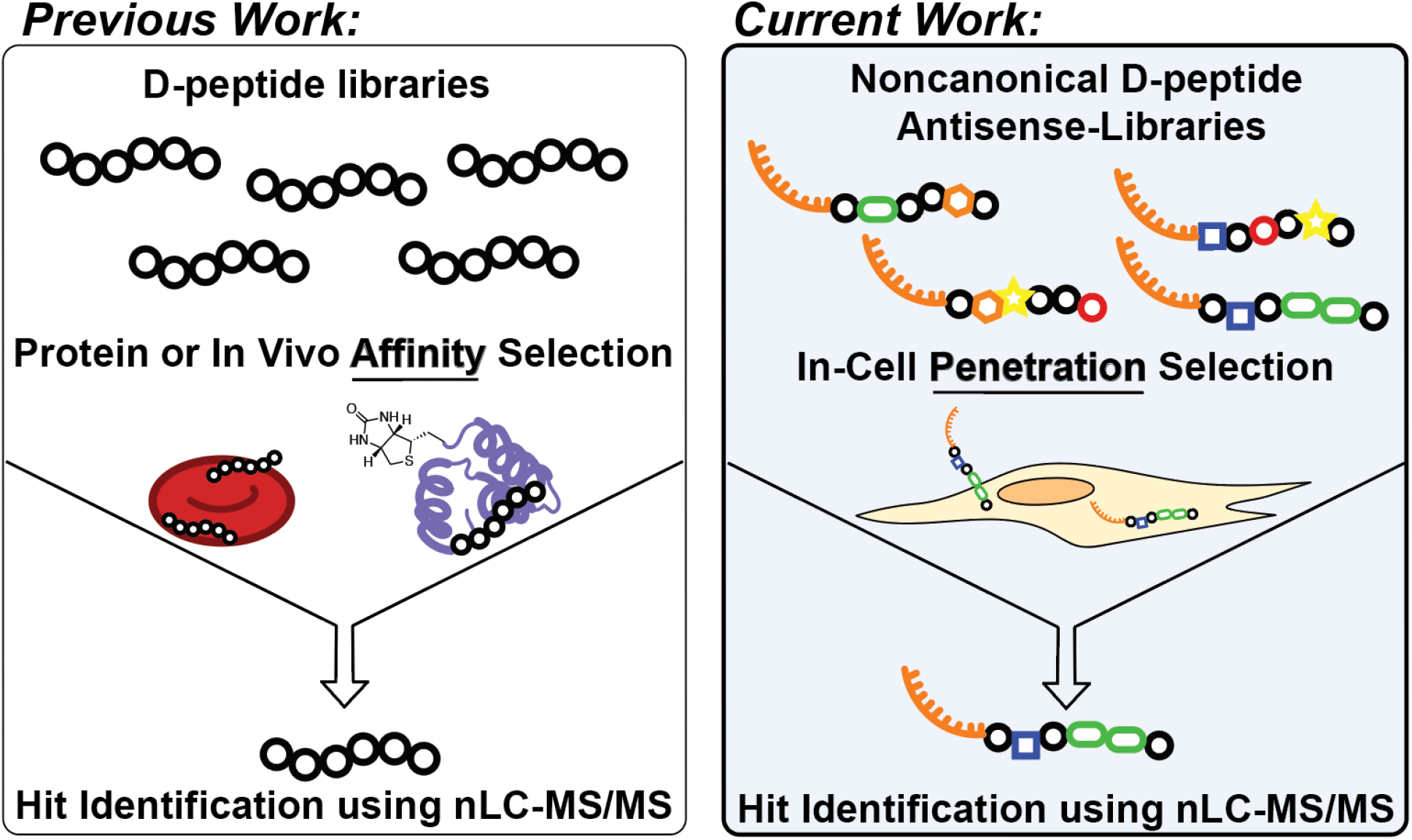
In-cell penetration selection—mass spectrometry identifies noncanonical peptides that access the cytosol. Building on prior methods for identifying binders to proteins and cells (left), in-cell PS-MS identifies noncanonical peptides that carry macromolecular cargo into the cytosol of cells (right).

## Results and Discussion

### Library preparation

The library was prepared with a “CPP-like” C-terminal sequence and six variable positions containing D- and unnatural amino acids (Fig. 2A). Split-and-pool synthesis afforded 0.016 µg of peptide per bead for a low-redundancy, 95,000-member library with a theoretical diversity greater than 10^8^. A D-kwkk motif, derived from the established cell-penetrating peptide penetratin^38^, was installed at the C-terminus to give the library a boost in activity. We have previously shown that these fixed constraints and C-terminal charge also increase peptide recovery in AS-MS.^27,39^ Unnatural amino acids were chosen to expand the chemical diversity and potentially enhance cell penetration of the library peptides. The library includes unnatural residues with non-α backbones to promote endosomal escape (γ-aminobutyric acid and β-alanine),^8^ residues with hydrophobic and aromatic functionality to increase membrane penetration (homoleucine, norleucine, naphthylalanine, and diphenylalanine),^40,41^ and additional charged residues and arginine analogues to enhance membrane penetration (diaminobutyric acid, aminopiperidine-carboxylic acid, aminomethylphenylalanine, and 2-amino-4-guanidinobutanoic acid)^42,43^ (Fig. 2B). The oxidative cleavable linker isoseramox was installed by reductive amination immediately following the variable region as previously reported,^44^ followed by a trypsin cleavage site to prevent the recovery of non-internalized peptides. Finally, azidolysine and biotin capped the N-terminus of the sequences to allow for PMO conjugation and affinity capture, respectively. Following cleavage from the resin, a portion of the library was conjugated by azide-alkyne cycloaddition to a model PMO derivatized with dibenzocyclooctyne (DBCO), monitored by LC-MS. Quality control analysis of the library by Orbitrap nano-liquid chromatography-tandem mass spectrometry (nLC-MS/MS) confirmed successful synthesis and exhibited a range of incorporated residues (Fig. 2C, Supporting Information Appendix III). The canonical and noncanonical residues are evenly incorporated and distributed throughout the variable region, with the exception of guanidyl residues, which showed reduced incorporation consistent with previous observations in solid-phase peptide synthesis.^45^ This library design ensured the isolation of 10-mer peptides with a native N-terminus, suitable for sequencing via tandem mass spectrometry.^44^

**Figure 2.**
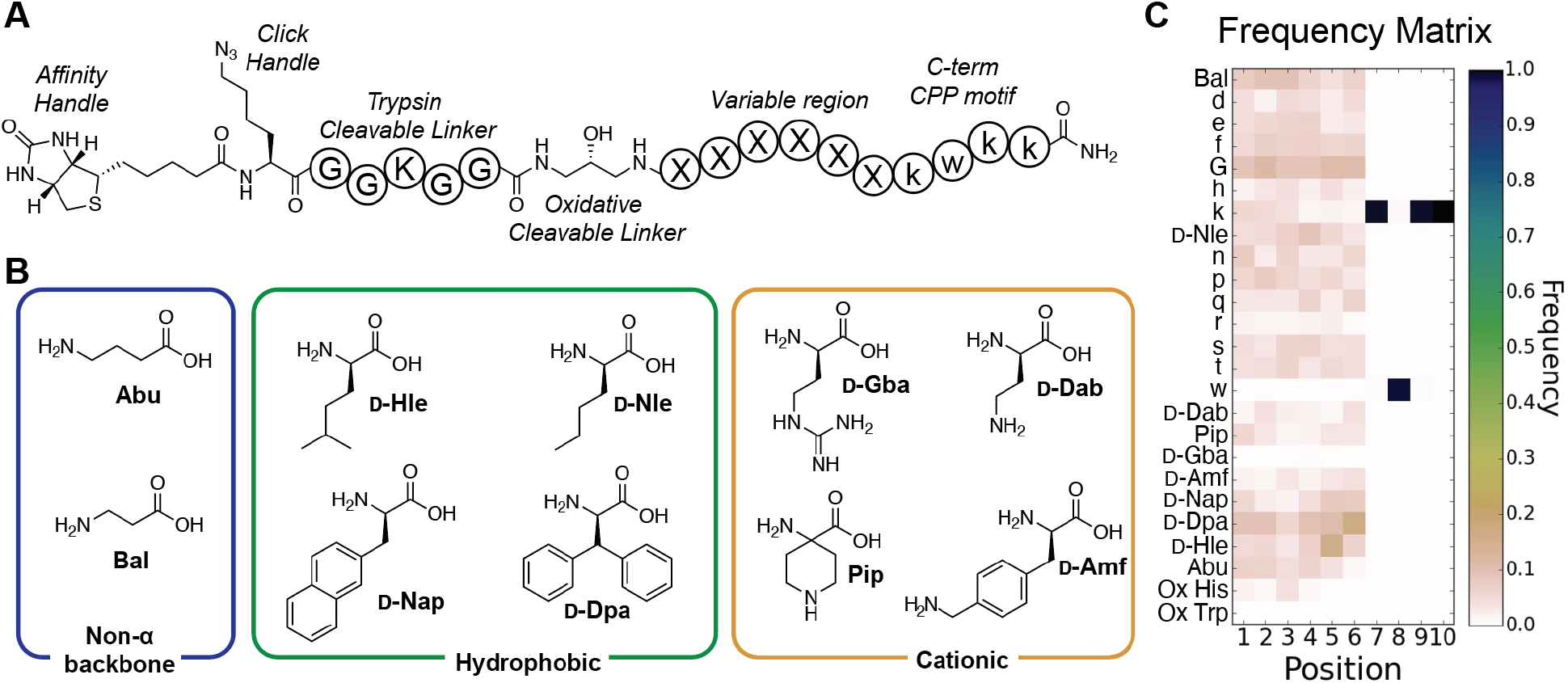
A combinatorial library was prepared with unnatural and D-amino acids. (A) Design of the library. An N-terminal biotin and azidolysine provide an affinity handle and conjugation handle, respectively. A trypsin-cleavable linker prevents isolation of extracellular conjugates, and isoseramox cleavable linker permits oxidative cleavage of the conjugates from streptavidin beads. Finally, there are six variable positions within the library peptides, with a “CPP-like” motif capping the C-terminus. (B) Structures of the unnatural monomers used. All natural-backbone monomers were in D-form. (C) Heat map of the quality control showing relative abundance of the various amino acids of the sequence up to the isoseramox linker. Positions 7-10 show the D-KWKK motif, with positions 1-6 showing the varied composition of the variable region. Abu (γ-aminobutyric acid), Bal (β-alanine), D-Hle (homoleucine), D-Nle (norleucine), D-Nap (naphthylalanine), D-Dpa (diphenylalanine), D-Dab (diaminobutyric acid), Pip (aminopiperidine-carboxylic acid), D-Amf (aminomethylphenylalanine), and D-Gba (2-amino-4-guanidinobutanoic acid).

We then performed a series of *in vitro* experiments to confirm that the peptides within the library had nuclear-localizing activity. The phenotypic assay used correlates with the amount of active PMO delivered to the nucleus by resulting in corrective splicing to produce enhanced green fluorescent protein (EGFP), quantified by flow cytometry. First, PMO-library aliquots demonstrated a concentration-dependent increase in activity (Fig. 3A). At the same time, the library at these concentrations did not exhibit any membrane disruption or toxicity as determined by a lactate dehydrogenase (LDH) release assay (Fig. 3B). We also found that testing the same concentrations of library aliquots containing different amounts of sequences, thereby increasing diversity, showed no difference in activity as well as no demonstrated membrane toxicity as determined by the LDH release assay (Fig. 3C, Fig. S3).

**Figure 3.**
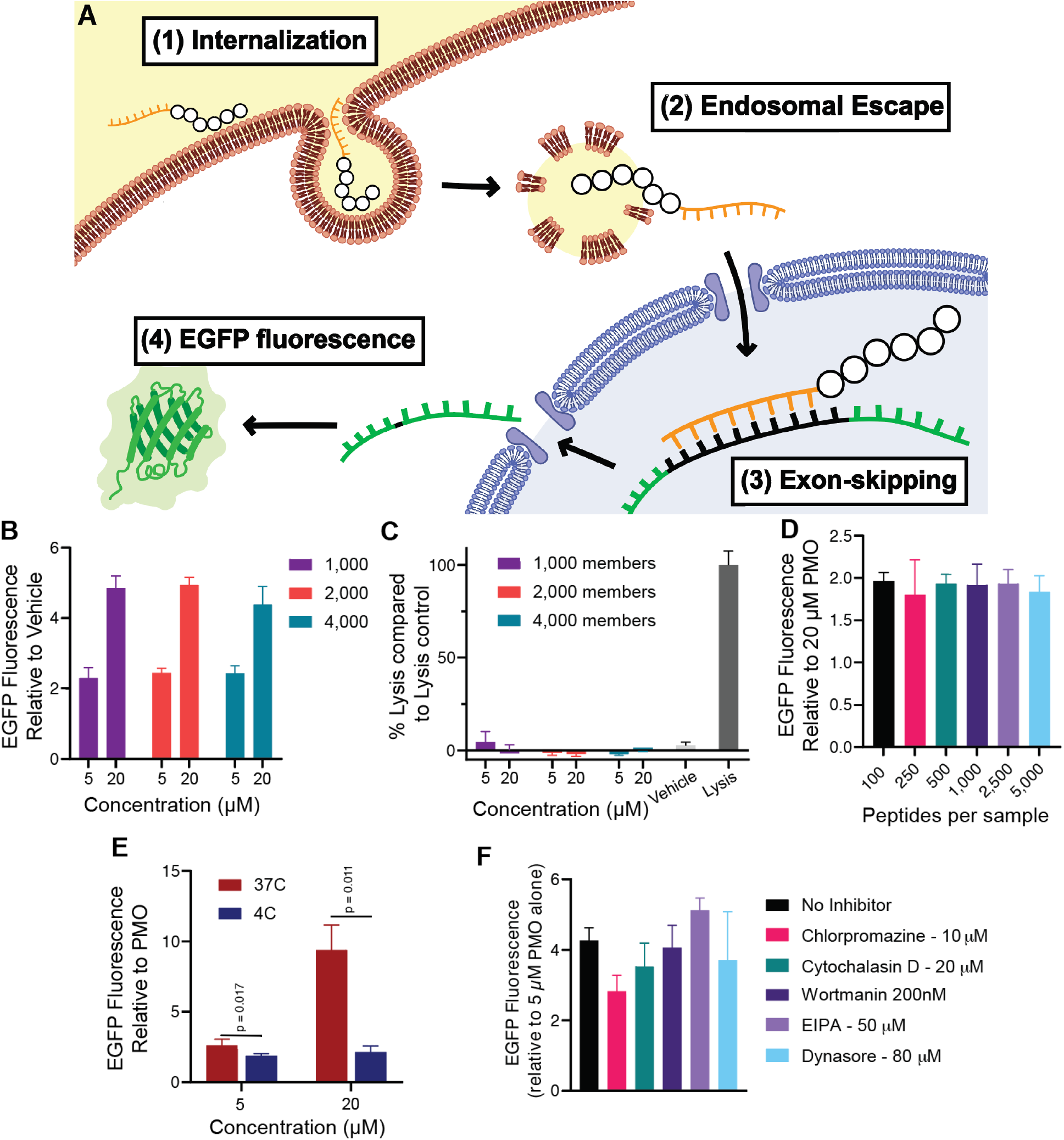
The CPP library can deliver PMO regardless of member size and enters via active transport. (A) HeLa 654 cells were treated with 5 or 20 µM of PMO-Library containing ∼1000, ∼2000, or ∼4000 members for 22 h prior to flow cytometry. Results are given as the mean EGFP fluorescence of cells treated with PMO-peptide relative to the fluorescence of cells alone. Bars represent mean ± SD, N = 3. PMO-Library samples show concentration-dependent PMO delivery at all library sizes tested. (B) HeLa 654 cells were treated with 5 or 20 µM PMO-Library for 22 h, then tested for LDH released into the cell media. Results are given as LDH release above vehicle relative to fully lysed cells. Bars represent mean ± SD, N = 3. No compounds showed LDH release significantly above vehicle-treated cells. (C) HeLa 654 cells were treated with 20 µM PMO-Library of varying member sizes or 20 µM PMO alone for 22 h prior to flow cytometry. Results are given relative to the fluorescence of PMO-treated cells. Bars represent mean ± SD, N = 3. There is no significant difference in EGFP fluorescence between the libraries of different sizes. Experiment was repeated twice with similar results, shown in Figure S1. (D) HeLa 654 cells were pre-incubated at 4 °C or 37 °C for 30 min prior to treatment with 20 µM PMO-Library or 20 µM PMO alone for 2 h at the indicated temperature. After treatment, cells were washed with 0.1 mg/mL heparin and incubated in media for 22 h prior to flow cytometry. Results are given relative to the fluorescence of PMO-treated cells. Bars represent mean ± SD, N = 3. There was a significant difference between the 4 °C and 37 °C treatment conditions at 5 µM (p = 0.017) and 20 µM (p = 0.011) of PMO-library. (E) Plot of EGFP mean fluorescence intensity relative to PMO for cells treated with different endocytosis inhibitors. The cells were pre-incubated for 30 min with the indicated compound and then 10 µM PMO-library (1,000 members) was added. After treatment with the construct for 3 h, the cells were washed with 0.1 mg/mL heparin and the media was exchanged for fresh, untreated media for 22 h prior to flow cytometry. At 10 µM chlorpromazine, EGFP fluorescence significantly decreased (p = 0.004). P-values determined by two-sided unpaired student’s t-test, N = 3. Endocytosis experiments were repeated with similar results, shown in Figure S2.

The PMO delivery activity of the library is likely energy-dependent, similar to PMO-CPP conjugates previously investigated.^18,19,46^ A 1,000-member portion of the PMO-library was incubated with cells at 4 °C, conditions that arrest energy-dependent uptake. After incubation with the PMO-CPP conjugates, each well was washed extensively with PBS and heparin in order to disrupt and remove membrane-bound conjugates.^47^ The cells were warmed back up to 37 °C and the assay continued in standard format and analyzed by flow cytometry. Temperature treatments did not result in membrane toxicity as determined by LDH release assay (Fig. S4). The significant decrease in library PMO delivery relative to PMO alone at 4 °C for both 5 µM and 20 µM library incubation conditions suggests energy-dependent uptake for the PMO-CPPs (Fig. 3D). Previous PMO-CPPs discovered in our laboratory are hypothesized to enter cells via clathrin-mediated endocytosis, as demonstrated through incubation with a panel of chemical endocytosis inhibitors, suggesting this could be a likely mechanism of energy-dependent uptake for a PMO-CPP library. ^18,19,46^ To confirm this, a 1,000 member library was tested with a series of chemical endocytosis inhibitors in a pulse-chase format EGFP assay, in which HeLa 654 cells were pre-incubated with inhibitors to arrest various endocytosis pathways before PMO-CPPs were added. Following 3 h co-incubation, cells were washed extensively with heparin to dissociate membrane-bound constructs.^18,19,46^ Activity of the library sample was reduced by 10 µM chlorpromazine (Fig. 3E), a known inhibitor of clathrin-mediated endocytosis. Again, no membrane toxicity was observed under these conditions (Fig. S5).

### In-cell penetration selection-mass spectrometry

We subjected the library to the in-cell penetration selection-mass spectrometry platform (in-cell PS-MS) to discover sequences that are present in the whole cell lysate and in the cytosol (Fig. 4). Here we are profiling for sequences that access the cytosol, making them more likely to access the nucleus and the RNA target of the PMO cargo. The protocol for extracting biotinylated sequences was adapted from our recent method of profiling mixtures of PMO-D-CPPs from the cytosol and whole cell using MALDI-ToF.^48,49^ Confluent HeLa cells in a 12-well plate were treated with 20 µM of biotin-library or PMO-biotin-library (∼10^3^ members, 3.5 nmol individual peptide per bead) for 1 h, before being washed with PBS and heparin to dissociate membrane-bound conjugates. Cells were then lifted and extracellular conjugates digested with Trypsin, pelleted, and washed with PBS. Cells were gently lysed using either RIPA buffer (for whole cell extraction) or digitonin buffer (for cytosolic extraction).^48,50^ Digitonin extraction is selective to cholesterol-rich membranes, leaving endosomes and some organelles intact while permeabilizing the cell membrane and semi-permeabilizing the nucleus.^51^ Exclusion of endosomes in the cytosolic fraction was confirmed by Western blot, in which the endosomal marker, Rab5, is absent (Fig. S6). In addition to the experimental samples, half of the no-treatment lysates were spiked with library as positive controls (Supporting Information Appendix IV).

**Figure 4.**
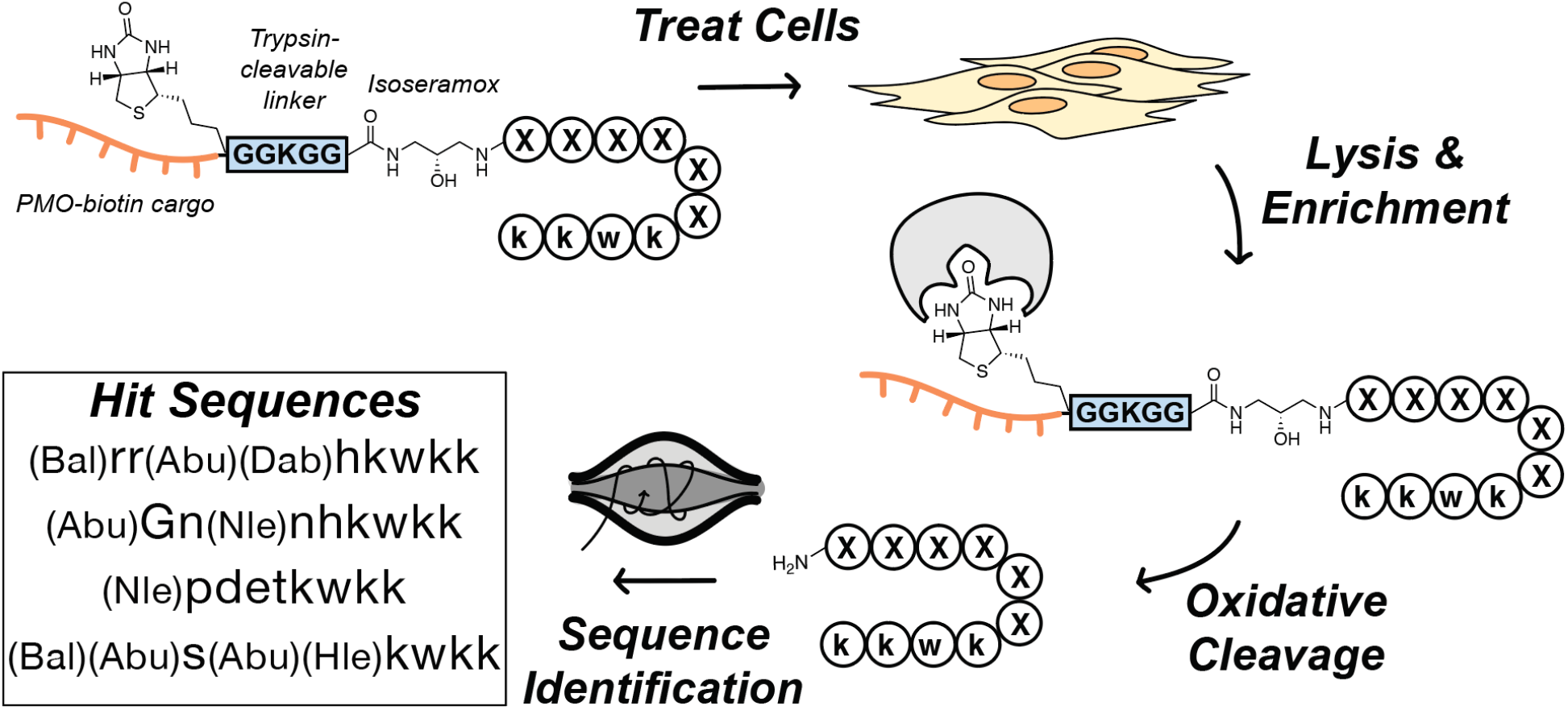
Workflow of in-cell penetration selection-mass spectrometry. HeLa cells were treated with 20 µM PMO-biotin-library or biotin-library (1,000 members) for 1 h at 37 °C. Cells were then extensively washed with PBS and 0.1 mg/mL heparin before lysis with RIPA (whole cell extract) or digitonin (cytosolic extract). Lysates were incubated with magnetic streptavidin beads, and the C-terminal native peptides were cleaved from the beads under oxidative conditions. The peptides were desalted through solid-phase extraction and sequenced by LC-MS/MS. Hit PMO-delivering sequences were then identified as those peptides found only in the PMO-library fractions that do not overlap with peptides found in the cell only control or the samples treated with the biotin-library.

Biotinylated species in the lysates were affinity captured with magnetic streptavidin beads, and ultimately released by oxidative cleavage using brief incubation with sodium periodate. We had previously used this cleavable linker to recover a single PMO-CPP conjugate from inside HeLa cells, and it was found to reliably cleave library peptides from streptavidin beads to isolate native peptides for sequencing by mass spectrometry.^44^ The isolated peptides were desalted by solid-phase extraction and analyzed via Orbitrap tandem mass spectrometry using a mixed fragmentation method optimized for cationic peptides, consisting of electron-transfer dissociation (ETD), higher-energy ETD, and higher-energy collisional dissociation (HCD). Sequences matching the library design were then identified using a Python script.^27,39^

### Hit peptides identified from PS-MS show nuclear PMO delivery, but are not solely responsible for the PMO delivery activity of the library

Several hit peptides were selected for experimental validation and showed differential activities depending on the fraction in which they were found. We selected two sequences found exclusively in the cytosolic extract (Pep1a, Pep1b) and two from only the whole cell extract (Pep1c, Pep1d), with sequences shown in Fig. 5A-B. These peptides were synthesized via semi-automated solid-phase fast-flow peptide synthesis^52^ with identical sequences to the library design with the exception of a D-Ser residue to replace the isoseramox linker. These sequences were tested first in a concentration-response EGFP assay. The sequences extracted from the cytosol showed significantly increased activity compared to the sequences from the whole cell lysate, with Pep1a showing an EC50 of 43 µM compared to Pep1c with EC50 of 380 µM (Fig. 5C). It was also confirmed that these sequences did not exhibit membrane toxicity at the concentrations tested (Fig. 5D). The peptides showed a positive correlation between charge and activity, with the highest performing peptide (Pep1a) having a charge of +7, compared to Pep1c with a charge of +2. This trend of positive charge correlating with PMO activity has been observed consistently in our lab.^18–20,46^ Although the hit peptides do not show higher activity than the parent peptide penetratin in its native L-form, they do show less toxicity and membrane disruption than penetratin at 25 µM in an LDH release assay (Fig. 5D), demonstrating their potential utility as PMO delivery vehicles. We also compared Pep1a and Pep1c to the known endosomal escape peptide Bpep^48^, composed of eight Arg residues interspaced with non-α-backbone residues β-alanine and 6-aminohexanoic acid, in its D-form which shows an EC50 closer to 3 µM (Fig. 5E). Interestingly, Pep1a shares some similar motifs to Bpep, namely two Arg residues flanked by two non-α-backbone residues. If these motifs are responsible for endosomal escape, it is not surprising that Pep1a was found in the cytosol and confirmed to have significant PMO delivery activity.

**Figure 5.**
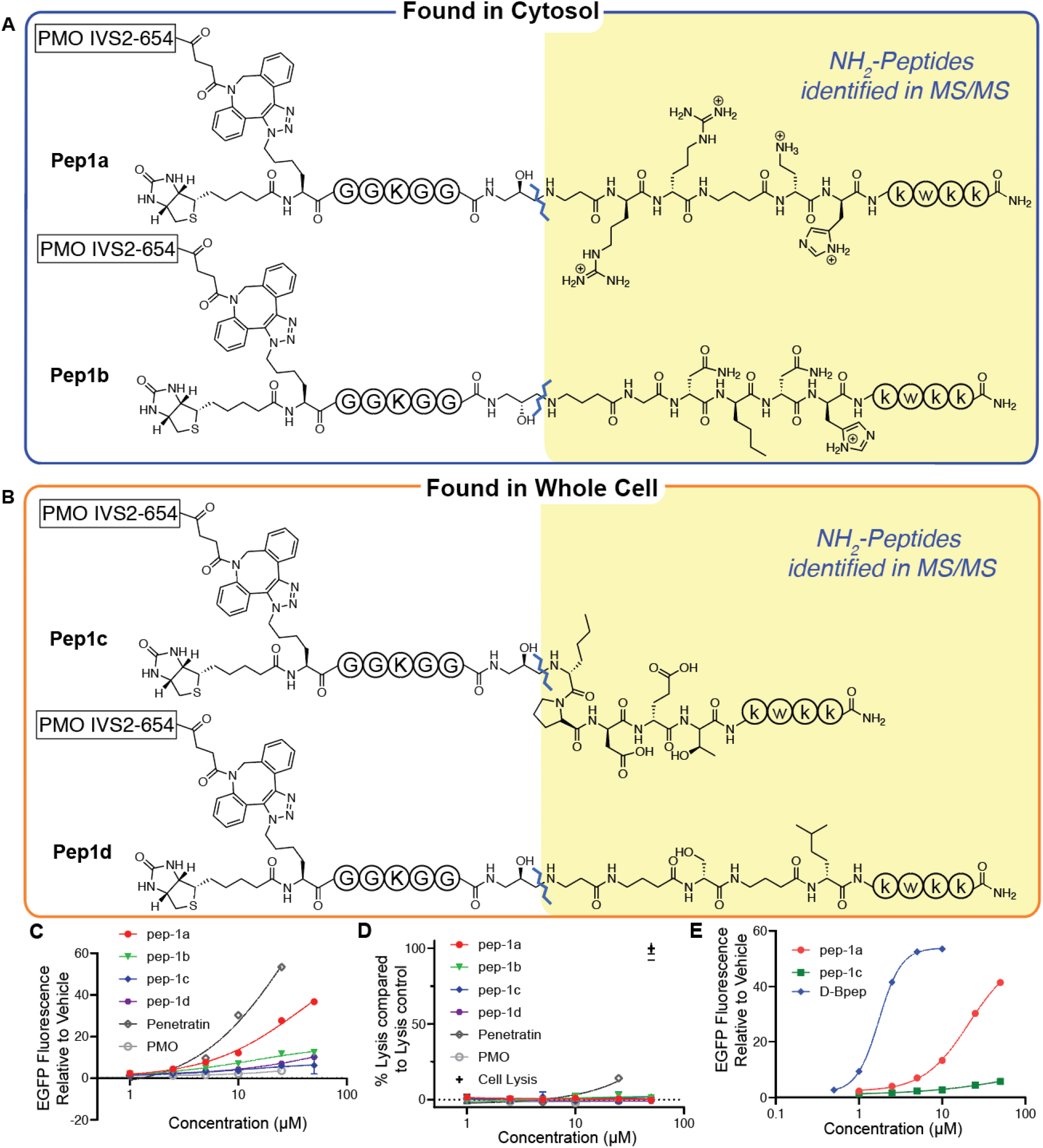
Candidate peptides from PS-MS demonstrate PMO delivery. Shown are the sequences of the four candidate peptides, grouped by their extraction from either (A) cytosol or (B) whole-cell lysate. On the N-terminus are biotin and PMO IVS2-654 linked through a dibenzocyclooctyne-azidolysine. The fixed residues of the sequence are shown within circles while the regions unique to each sequence are fully drawn structures. The isoseramox linker is cleaved during library screening, and is substituted for D-serine in the PMO-peptides synthesized for hit validation. (C) HeLa 654 cells were treated with 1, 2.5, 5, 10, 25, or 50 µM PMO-CPP for 22 h prior to flow cytometry. Results are given as the mean EGFP fluorescence of cells treated with PMO-peptide relative to the fluorescence of cells treated with vehicle only. Bars represent mean ± SD, N = 3. All 4 peptides show similar activity across biological replicates (Figure S7). (D) Cell supernatant from (C) was tested for LDH release. Results are given as percent LDH release above vehicle relative to fully lysed cells. Only L-penetratin showed significant (p = 0.008) LDH release above vehicle-treated cells. (E) HeLa 654 cells were treated with 0.5, 1, 2.5, 5, 10, 25, or 50 µM PMO-CPP for 22 h. Based on the average of three biological replicates, Pep1a has a significantly lower EC50 (24.6 µM) compared to pep1c with EC50 of 360 µM (p = 0.0013). Biological replicate demonstrated similar activity (Figure S8), and LDH assay demonstrated no membrane toxicity (Figure S9).

The bioactive hit peptides are not solely responsible for the PMO delivery exhibited by the entire 1,000-member library. Within a 20 µM treatment dose of a 1,000 member library *in vitro*, each individual peptide should hypothetically be present below 100 nM, a concentration at which no single peptide is known to deliver PMO cargo. To investigate whether overall library PMO delivery efficacy could be due to a few highly active peptides, we treated HeLa 654 cells with a 250-member library at 20 µM and compared the activity to HeLa cells treated with the same library with a penetrant peptide (either Pep1a or the positive control D-Bpep) spiked in at roughly the concentration of the individual library members (Fig. S10A, S11). There was no significant difference in PMO delivery between the 250-member library alone and the library with potent peptides spiked in, showing that the overall, combined library penetration is unlikely to be affected by the activity of a few members. To confirm library activity is not affected by small differences in concentration between individual library members, we repeated the experiment with a larger library of 2,500 members and spiked in penetrant peptides at 10-fold higher concentrations than the individual library members, which also showed no significant change in library PMO delivery.

Instead of a few highly active members, library penetration is more likely the result of many cationic peptides acting in tandem. There has been previous evidence that at high concentrations (> 20 µM), cell entry of highly cationic peptides can be caused by non-specific flooding via non-endocytic pathways, via a positive feed-back loop that involves alteration of the plasma membrane composition.^53^ We demonstrated that the library enters cells in an energy-dependent manner, ruling out this and other mechanisms of energy-independent non-specific library entry. However, the concept of multiple cationic peptides acting in tandem at the plasma membrane suggests that such an effect could be responsible for the overall penetration of the library, from an “ensemble” of cationic peptides at the cell membrane. To test this hypothesis, we modeled the library on a much smaller scale, using only five peptides. Since we demonstrated that libraries ranging in size from 100 to 5,000 members show similar activity, we expect that even a small model may represent the activity of library peptides. Thus, we tested the four hit library candidates, as well as a “library peptide” found in the quality control sequencing of the library, but not extracted from cells, both individually and in combination for PMO delivery (Fig. S10B, S12). The 5 µM “combined peptides” sample contains each individual peptide at 1 µM, yet this five-member library shows significantly more PMO delivery than any of the individual peptides at 1 µM. In fact, the PMO delivery of the peptides in combination at 5 µM total peptide more closely matches the averaged values of all five peptides individually at 5 µM, further suggesting that the activity observed from the library is due to an ensemble effect from the activity of many cationic individual peptides and not due to a few highly active sequences. These experimental conditions did not result in membrane toxicity (Fig. S13, S14)

### Hit peptide demonstrates high endosomal escape activity and PMO delivery

These PMO-CPPs likely enter the cell via clathrin-mediated endocytosis as we have found for previous constructs and the parent library. Pep1a was tested with a series of chemical endocytosis inhibitors in the same pulse-chase format EGFP assay used to assess the endocytosis mechanism of the 1,000 member library sample. Activity of Pep1a was impacted by 10 µM chlorpromazine (Fig. 6A), while none of the endocytosis inhibitors demonstrated cell-associated toxicity or membrane lysis (Fig. S18). The increased activity seen with EIPA is likely a result of autofluorescence from the inhibitor itself. Chlorpromazine is a known inhibitor of clathrin-mediated endocytosis, indicating that this conjugate may participate in this pathway. Moreover, the 4 °C condition also significantly impacted the activities of each conjugate, indicating that, like the entire library sample (Fig. 3D), uptake of the four hit peptides is energy-dependent, and the peptides are most likely entering the cells through endocytosis (Fig. 6B).

**Figure 6.**
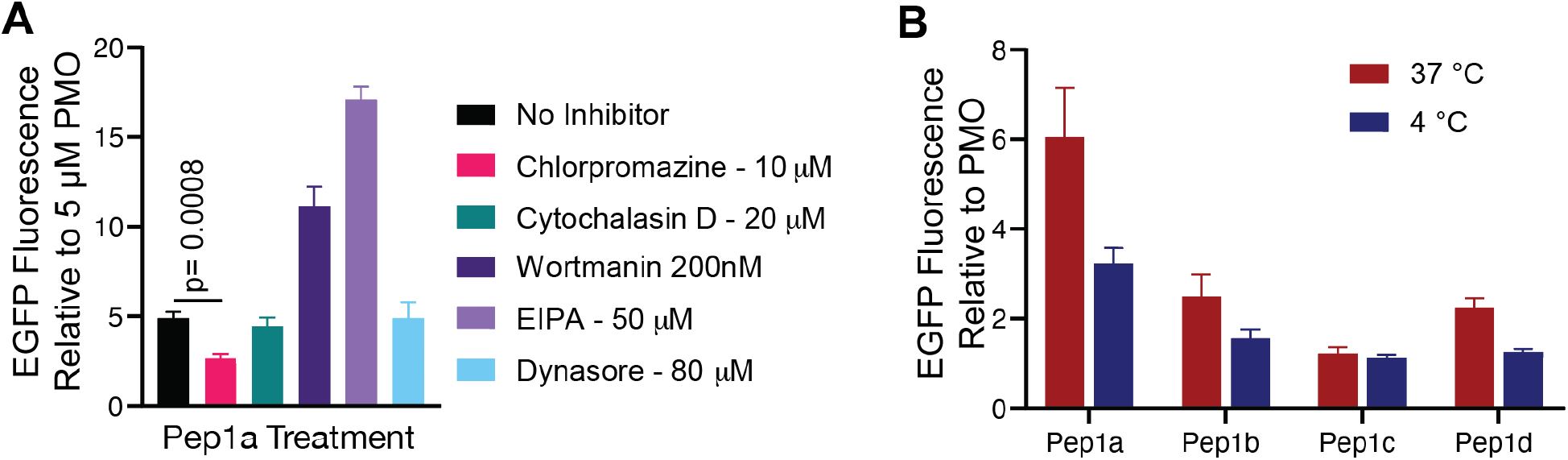
Hit peptides likely deliver PMO via clathrin-mediated endocytosis. (A) Plot of EGFP mean fluorescence intensity relative to PMO for cells treated with different endocytosis inhibitors. The cells were pre-incubated for 30 min with the indicated compound and then 5 µM PMO-Pep1a was added. After treatment with the construct for 3 h, the cells were washed with 0.1 mg/mL heparin and the media was exchanged for fresh, untreated media for 22 h prior to flow cytometry. Results are given as the mean EGFP fluorescence of cells treated with PMO-peptide relative to the fluorescence of cells treated with 5µM PMO. Bars represent mean ± SD, N = 3. At 10 µM chlorpromazine, EGFP fluorescence significantly decreased (p = 0.0008). (B) Plot of EGFP mean fluorescence intensity relative to PMO for cells incubated with PMO-CPPs at 4 °C or 37 °C. The cells were pre-incubated for 30 min at 4 °C or 37 °C, followed by the addition of PMO-peptide conjugate to each well at a concentration of 5 µM. After incubation at 4°C or 37 °C for 2 h, the cells were washed with 0.1 mg/mL heparin and the media was exchanged for fresh, untreated media for 22 h prior to flow cytometry. Results are given as the mean EGFP fluorescence of cells treated with PMO-peptide relative to the fluorescence of cells treated with vehicle only. Bars represent mean ± SD, N = 3. The experiments were repeated with similar results (Figure S15, S16). These treatment conditions did not result in membrane toxicity as measured by LDH release assay (Figure S17, S18)

We further investigated the differences in activity between the sequences found in the cytosol versus the whole cell lysate and compared them to a benchmark compound, PMO-D-Bpep, using flow cytometry.^48^ For this purpose, several SulfoCy5-labeled PMO-CPPs were generated and tested to ensure the fluorophore did not impact PMO delivery activity (Fig. 7A). Comparing the results of the EGFP assay of conjugates with and without the fluorophore, no significant differences were found between the constructs’ EC50 values (Fig. 7B-D).

**Figure 7.**
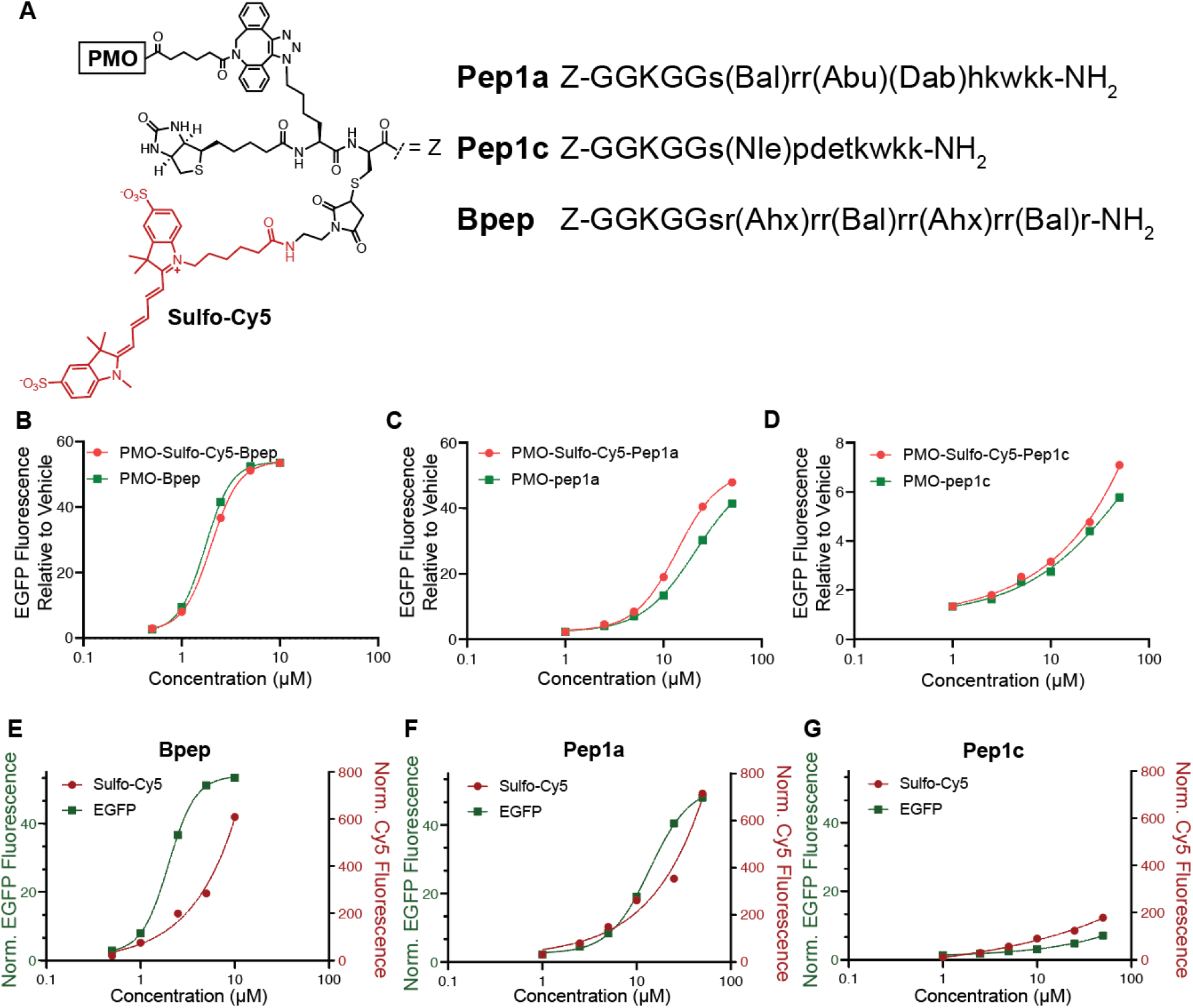
SulfoCy5 label does not impact PMO delivery. (A) Sequences of PMO-SulfoCy5-CPP constructs, with the N-terminal cargo fully drawn out (Z). Lowercase letters denote D-amino acids. (B-D) HeLa 654 cells were treated with 1, 2.5, 5, 10, 25, or 50 µM PMO-CPP or PMO-SulfoCy5-CPP for 22 h prior to flow-cytometry. Results are given as the mean EGFP fluorescence of cells treated with PMO-peptide relative to the fluorescence of cells treated with vehicle only. Bars represent mean ± SD, N = 3. (B) Treatment with D-Bpep constructs. (C) Treatment with Pep1a constructs. (D) Treatment with Pep1c constructs. Pep1a showed no significant difference with the addition of SulfoCy5 at 5 µM and below (p<0.05), while Pep1c and D-Bpep showed no significant difference between the SulfoCy5 and standard constructs at 25 µM and below (p<0.05). No peptides had significant differences in EC50 between SulfoCy5 and unlabeled constructs. Bal (beta-Alanine), Abu (γ-aminobutyric acid), Dab (D-diaminobutyric acid), Nle (D-Norleucine), Ahx (6-aminohexanoic acid). Experiments shown in B-D were repeated with similar results (Figure S19) (E-G) HeLa 654 cells were treated with 1, 2.5, 5, 10, 25, or 50 µM PMO-SulfoCy5-CPP for 22 h prior to flow cytometry. Results are given as the mean fluorescence of cells treated with PMO-SulfoCy5-peptide relative to the fluorescence of cells treated with vehicle only for each channel. Bars represent mean ± SD, N = 3. (E) Treatment with D-Bpep constructs. (F) Treatment with Pep1a constructs. (G) Treatment with Pep1c constructs. Experiments shown in E-G were repeated with similar results (Figure S20). Cells from experiments B-G did not show membrane toxicity as determined by LDH release assay (Figure S21).

After confirming the Cy5 label did not interfere with efficacy, we then looked at the uptake and nuclear delivery of the PMO-SulfoCy5-CPPs. This study was performed by monitoring both EGFP fluorescence elicited by nuclear-localized compound and Cy5 fluorescence to measure total cellular uptake of each analog. Each conjugate demonstrated similar concentration-dependent increases in both EGFP and Cy5 fluorescence (Fig. 7E-G). Pep1c showed low fluorescence signal in both the EGFP and Cy5 channels. On the other hand, Pep1a showed higher fluorescence signals in each channel, indicating greater uptake and nuclear localization compared to Pep1c. This readout underscores the utility of the PS-MS methodology in identifying peptides that can efficiently escape endosomes. Interestingly, D-Bpep showed greater EGFP fluorescence but slightly diminished Cy5 fluorescence than Pep1a, indicating that D-Bpep may access the nucleus more efficiently once taken up in endosomes, but with a lower total uptake compared to Pep1a. This pattern has been observed previously, in that the endosomal escape motifs present in Bpep enhance endosomal escape while diminishing cellular uptake.^8^

The uptake of the fluorescent conjugates into HeLa cells was also evaluated via confocal microscopy. HeLa cells were treated with 5 µM or 25 µM of SulfoCy5-labeled conjugates for 30 min, followed by a wash in complete media. Hoechst and LysoTracker Green in complete media were added to the cells immediately prior to imaging. D-Bpep was again used as a control here as it is a known highly active sequence able to escape the endosome and localize to the nucleus.^48^ Indeed, at both concentrations diffuse fluorescence was observed in the cytosol and nucleus of PMO-Bpep-treated cells, in addition to punctate fluorescence that co-localized with LysoTracker, suggesting accumulation in endosomes (Fig. 8). Pep1a also demonstrated perinuclear localization, especially at the higher concentration (Fig. 8B). In contrast, Pep1c showed significantly reduced overall fluorescence inside the cell, and exclusively as punctate fluorescence within endosomes.

**Figure 8.**
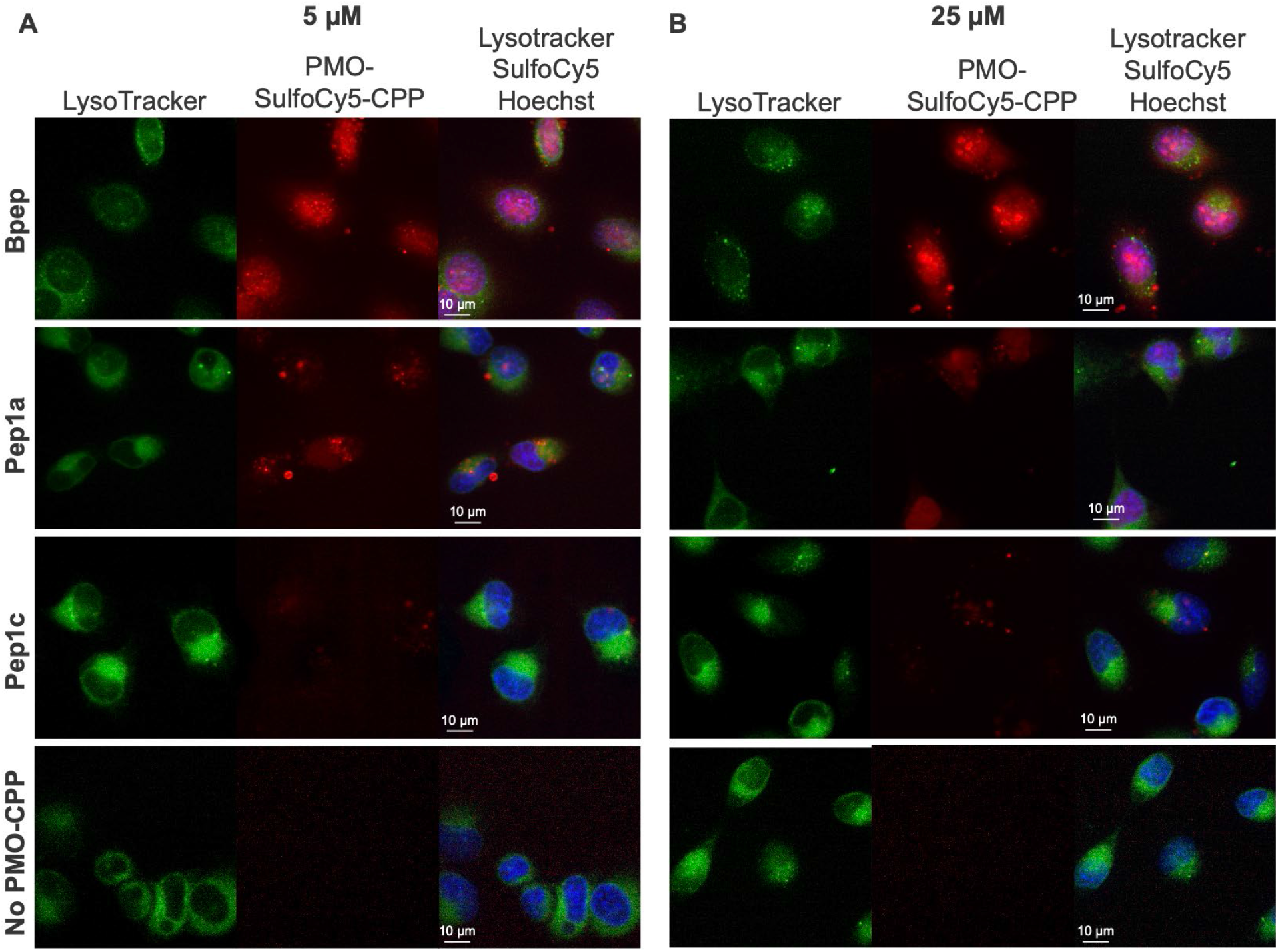
Pep1a localizes to cytosol and nucleus. Confocal micrographs of HeLa cells treated with (A) 5 µM or (B) 25 µM PMO-SulfoCy5-Bpep, PMO-SulfoCy5-Pep1a, or PMO-SulfoCy5-Pep1c. Hoechst labels the nuclei and Lyostracker Green labels the endosomes. SulfoCy5-labeled PMO-CPPs can be observed in the endosomes, cytosol, and nuclei of the cells.

To confirm the hit peptides do not simply permeabilize endosomes, we traced the integrity of endosomes in cells treated with Cy5-labeled hit peptides. HeLa cells were first preincubated with DEAC-k5, an endosomal-localizing peptide composed of D-lysine residues.^54^ The DEAC-k5 is visible as blue puncta in the no-CPP treatment control, indicating the expected endosomal localization (Fig. S22). After treatment with PMO-SulfoCy5-CPP constructs, the DEAC-k5 continues to occupy endosomes, indicating that Pep1a and Pep1c do not non-specifically permeabilize the endosome to release other endosomal cargo.

### Conclusions

Affinity selection-mass spectrometry (AS-MS) techniques have traditionally been used to probe protein-protein interactions in vitro.^28^ Our group has recently shown that chemical libraries may reach the diversity of other display techniques for identification of peptide binders to proteins^27^, and that this strategy can be applied for cell-surface selection in vivo.^36^ In this work, we have added an additional spatial element to this strategy, by extracting the cytosol for in-cell selection of fully synthetic peptide libraries conjugated to a model antisense cargo. By comparing these sequences to those found in a whole cell extract, we can exclude sequences that accumulate in the endosomes.

In-cell PS-MS combined with subcellular fractionation resulted in the identification of a novel, abiotic peptide capable of accessing the cytosol and delivering PMO to the nucleus. Pep1a, like the positive control peptide Bpep, was able to deliver PMO to the nucleus by escaping endosomes. Furthermore, Pep1a does not appear to permeabilize the endosome to allow the escape of other endosomal cargo, nor does it demonstrate cell membrane toxicity at the measured concentrations in an LDH release assay. All peptides discovered through this novel platform demonstrated lower toxicity than the CPP penetratin at 25 µM, which contributed the fixed “CPP-like” C-terminal region in the library. Endowed with lower toxicity and superior chemical diversity provided by the noncanonical residues, the peptides discovered with the in-cell PS-MS platform show advantages over the library’s parent peptide. The few active PMO-CPPs individually sequenced and validated are not solely responsible for the overall cell penetration of the library, however. In fact, it is more likely that the PMO delivery arises from the combined activity of a number of peptides at low concentration, including the hits discovered with our platform, despite the varying activity of the individual CPPs within the library.

Current investigations in our laboratory aim to further combine this method with orthogonal approaches in order to better focus the selection on successful peptides. As such, we envisage using chromatographic fractionation of the peptide libraries to enrich for penetrant peptides within the library before conducting the PS-MS uptake assay. This chromatographic pre-enrichment would allow for screening of an overall less active, but more diverse library, perhaps obviating the need for a pre-installed fixed C-terminal penetrating motif.

## Experimental Section

### Reagents and Solvents

H-Rink Amide-ChemMatrix resin was obtained from PCAS BioMatrix Inc. (St-Jean-sur-Richelieu, Quebec, Canada) and TentaGel was obtained from Rapp Polymere (Tuebingen, Germany). 1-[Bis(dimethylamino)methylene]-1*H*-1,2,3-triazolo[4,5-b]pyridinium-3-oxid-hexafluorophosphate (HATU), Fmoc-L-Lys(N_3_), Fmoc-β-Ala-OH, Fmoc-norleucine, *N*α-Fmoc-*N*γ-Boc-D-2,4-diaminobutyric acid, Fmoc-D-homoleucine, Fmoc-3,3-diphenyl-D-alanine, Fmoc-3-(1-naphthyl)-D-alanine, Fmoc-4-(Boc-aminomethyl)-D-phenylalanine, 1-Boc-piperidine-4-Fmoc-amino-4-carboxylic acid, Fmoc-γ-aminobutyric acid, and 2-(Fmoc-amino)-4-(bis-Boc-guanidino)-D-butyric acid were purchased from Chem-Impex International (Wood Dale, IL). Fmoc-protected D-amino acids (Fmoc-Arg(Pbf)-OH; Fmoc-Asn(Trt)-OH; Fmoc-Asp-(O*t*-Bu)-OH; Fmoc-Gln(Trt)-OH; Fmoc-Glu(O*t*-Bu)-OH; Fmoc-Gly-OH; Fmoc-His(Trt)-OH; Fmoc-Lys(Boc)-OH; Fmoc-Phe-OH; Fmoc-Pro-OH; Fmoc-Ser(But)-OH; Fmoc-Thr(*t*-Bu)-OH; Fmoc-Trp(Boc)-OH), were purchased from the Novabiochem-line from MilliporeSigma. Sulfo-Cyanine5 maleimide was purchased from Lumiprobe (Cockeysville, MD), and 7-diethylaminocoumarin-3-carboxylic acid was purchased from AAT Bioquest (Sunnyvale, CA). Dibenzocyclooctyne acid was purchased from Click Chemistry Tools (Scottsdale, AZ). Cytochalasin D was obtained from Santa Cruz Biotech. Peptide synthesis-grade *N,N*-dimethylformamide (DMF), CH_2_Cl_2_ (DCM), diethyl ether, and HPLC-grade acetonitrile were obtained from VWR International (Radnor, PA). All other reagents were purchased from Sigma-Aldrich (St. Louis, MO). Milli-Q water was used exclusively. Hoechst 33342 and LysoTracker™ Green were purchased from ThermoFisher Scientific (Walthan, MA). The LDH Assay kit was purchased from Promega (Madison, WI).

### Liquid chromatography-mass spectrometry

LC-MS analyses were performed on an Agilent 6550 iFunnel Q-TOF LC-MS system (abbreviated as 6550) coupled to an Agilent 1290 Infinity HPLC system. Mobile phases were: 0.1% formic acid in water (solvent A) and 0.1% formic acid in acetonitrile (solvent B). The following LC-MS method was used for characterization:

### Method A: 1-61% B over 6 min, Zorbax C3 column (6550)

#### LC

Agilent EclipsePlus C18 RRHD column: 2.1 × 50 mm, 1.8 μm, column temperature: 40 °C, gradient: 0-1 min 1% B, 1-6 min, 1-61% B, 6-7 min, 91% B, 7-8 min, 1% B; flow rate: 0.5 mL/min.

#### MS

Positive electrospray ionization (ESI) extended dynamic range mode in mass range 300–3000 m/z. MS is on from 1 to 6 min.

All data were processed using Agilent MassHunter software package. Y-axis in all chromatograms shown represents total ion current (TIC) unless noted.

### General peptide preparation

#### Fast-flow Peptide Synthesis

Peptides were synthesized on a 0.1 mmol scale using an automated fast-flow peptide synthesizer for L-peptides and a semi-automated fast-flow peptide synthesizer for D-peptides.^52^ Automated synthesis conditions were used as previously reported.^55^ Briefly, a 100 mg portion of ChemMatrix Rink Amide HYR resin was loaded into a reactor maintained at 90 °C. All reagents were flowed at 40 mL/min with HPLC pumps through a stainless-steel loop maintained at 90 °C before introduction into the reactor. For each coupling, 10 mL of a solution containing 0.4 M amino acid and 0.38 M HATU in DMF were mixed with 600 μL of diisopropylethylamine and delivered to the reactor. Fmoc removal was accomplished using 10.4 mL of 20% (v/v) piperidine. Between each step, DMF (15 mL) was used to wash out the reactor. To couple unnatural amino acids or to cap the peptide (e.g., with 4-pentynoic acid), the resin was incubated for 30 min at room temperature with amino acid (1 mmol) dissolved in 2.5 mL of 0.4 M HATU in DMF with 500 μL of diisopropylethylamine (DIEA). After completion of the synthesis, the resin was washed 3 times with dichloromethane and dried under vacuum.

Semi-automated synthesis was carried out as previously described.^52^ 1 mmol of amino acid was combined with 2.5 mL of 0.4 M HATU and 500 µL of DIEA and mixed before being delivered to the reactor containing resin via syringe pump at 6 mL/min. The reactor was submerged in a water bath heated to 70 °C. An HPLC pump delivered either DMF (20 mL) for washing or 20% piperidine/DMF (6.7 mL) for Fmoc deprotection, at 20 mL/min.

#### Peptide Cleavage and Deprotection

Each peptide was subjected to simultaneous global side-chain deprotection and cleavage from resin by treatment with 5 mL of 94% trifluoroacetic acid (TFA), 2.5% thioanisole, 2.5% water, and 1% triisopropylsilane (TIPS) (v/v) at room temperature for 2 to 4 h. The cleavage cocktail was first concentrated by bubbling N_2_ through the mixture, and cleaved peptide was precipitated and triturated with 40 mL of cold ether (chilled in dry ice). The crude product was pelleted by centrifugation for three minutes at 4,000 rpm and the ether was decanted. This wash step was repeated two more times. After the third wash, the pellet was dissolved in 50% water and 50% acetonitrile containing 0.1% TFA, filtered through a fritted syringe to remove the resin and lyophilized.

#### Peptide Purification

The peptides were dissolved in water and acetonitrile containing 0.1% TFA, filtered through a 0.22 μm nylon filter and purified by mass-directed semi-preparative reversed-phase HPLC. Solvent A was water with 0.1% TFA additive and Solvent B was acetonitrile with 0.1% TFA additive. A linear gradient that changed at a rate of 0.5% B/min was used. Most of the peptides were purified on an Agilent Zorbax SB C18 column: 9.4 × 250 mm, 5 μm. Based on target ion mass data recorded for each fraction, only pure fractions were pooled and lyophilized. The purity of the fraction pool was confirmed by LC-MS.

#### Preparation of PMO-Peptides

PMO IVS2-654 (50 mg, 8 µmol) obtained from Sarepta Therapeutics was dissolved in 150 µL DMSO. To the solution was added a solution containing 2 equivalents of dibenzocyclooctyne acid (5.3 mg, 16 µmol) activated with HBTU (37.5 µL of 0.4 M HBTU in DMF, 15 µmol) and DIEA (2.8 µL, 16 µmol) in 40 µL DMF (Final reaction volume = 0.23 mL). The reaction proceeded for 25 min before being quenched with 1 mL of water and 2 mL of ammonium hydroxide. The ammonium hydroxide hydrolyzed any ester formed during the course of the reaction. After 1 hour, the solution was diluted to 40 mL in water/acetonitrile and purified using reverse-phase HPLC (Agilent Zorbax SB C3 column: 21.2 × 100 mm, 5 µm) and a linear gradient from 2 to 60% B (solvent A: water; solvent B: acetonitrile) over 58 min (1% B / min). Using mass data about each fraction from the instrument, only pure fractions were pooled and lyophilized. The purity of the fraction pool was confirmed by LC-MS.

#### Conjugation to peptides

PMO-DBCO (1 eq, 5 mM, water) was conjugated to azido-peptides (1 eq, 5 mM, water) at room temperature for 2 h. Reaction progress was monitored by LC-MS and additional stock of 5 mM azido-peptide was added until all PMO-DBCO was consumed. The purity of the final construct was confirmed by LC-MS to be >95%.

### Preparation of peptide libraries

Split-and-pool synthesis was carried out on 300 μm TentaGel resin (0.23 mmol/g) for a 95,000 member library. Splits were performed by suspending the resin in DCM and dividing it evenly (via pipetting) among 22 plastic fritted syringes on a vacuum manifold. Couplings were carried out as follows: solutions of Fmoc-protected amino acids (10 equivalents relative to the resin loading), PyAOP (0.38 M in DMF; 0.9 eq. relative to amino acid), and DIEA (1.1 eq. for histidine; 3 eq. for all other amino acids) were each added to individual portions of resin. Couplings were allowed to proceed for 60 min. Resin portions were recombined and washed with DCM and DMF. Fmoc removal was carried out by treatment of the resin with 20% piperidine in DMF (1x flow wash; 2x 5 min batch treatments). Resin was washed again with DMF and DCM before the next split.

### EGFP Assay

HeLa 654 cells obtained from the University of North Carolina Tissue Culture Core facility were maintained in MEM supplemented with 10% (v/v) fetal bovine serum (FBS) and 1% (v/v) penicillin-streptomycin at 37 °C and 5% CO_2_. 18 h prior to treatment, the cells were plated at a density of 5,000 cells per well in a 96-well plate in MEM supplemented with 10% FBS and 1% penicillin-streptomycin.

For individual peptide testing, PMO-peptides were dissolved in PBS without Ca^2+^ or Mg^2+^ at a concentration of 1 mM (determined by UV) before being diluted in MEM. Cells were incubated at the designated concentrations for 22 h at 37 °C and 5% CO_2_. Next, the treatment media was removed, and the cells were washed once before being incubated with 0.25 % Trypsin-EDTA for 15 min at 37 °C and 5% CO_2_. Lifted cells were transferred to a V-bottom 96-well plate and washed once with PBS, before being resuspended in PBS containing 2% FBS and 2 µg/mL propidium iodide (PI). Flow cytometry analysis was carried out on a BD LSRII flow cytometer. Gates were applied to the data to ensure that cells that were positive for propidium iodide or had forward/side scatter readings that were sufficiently different from the main cell population were excluded. Each sample was capped at 5,000 gated events.

Analysis was conducted using Graphpad Prism 7 and FlowJo. For each sample, the mean fluorescence intensity (MFI) and the number of gated cells was measured. To report activity, triplicate MFI values were averaged and normalized to the PMO alone condition. For the final set of PMO-peptides evaluated, three biological replicates were performed.

### Endocytosis Inhibition Assay

Chemical endocytosis inhibitors were used to probe the mechanism of delivery of PMO by these peptides in a pulse-chase format. We have conducted such analysis on similar PMO-peptide constructs previously with comparable outcomes.^46^ For the PMO constructs, HeLa 654 cells were preincubated with various chemical inhibitors for 30 minutes before treatment with PMO-CPP constructs for three hours. The panel of endocytosis inhibitors included: 10 µM chlorpromazine (CPZ), which is demonstrated to interfere with clathrin-mediated endocytosis; 20 µM cytochalasin D (CyD), which inhibits phagocytosis and micropinocytosis; 200 nM wortmannin (Wrt), which alters various endocytosis pathways by inhibiting phosphatidylinositol kinases; 50 µM EIPA (5-(*N*-ethyl-*N*-isopropyl)amiloride), which inhibits micropinocytosis; and 80 µM Dynasore (Dyn), which also inhibits clathrin-mediated endocytosis.^56,57^ Treatment media was then replaced with fresh media and the cells were incubated for 22 h at 37 °C and 5% CO_2_. Cells were then lifted as previously described and EGFP synthesis was measured by flow cytometry.

### LDH Assay

Cytotoxicity assays were performed in HeLa 654 cells. Cell supernatant following treatment for flow cytometry was transferred to a new 96-well plate for analysis of LDH release. To each well of the 96-well plate containing supernatant was added CytoTox 96 Reagent (Promega). The plate was shielded from light and incubated at room temperature for 30 min. Equal volume of Stop Solution was added to each well, mixed, and the absorbance of each well was measured at 490 nm. The measurement of vehicle-treated cells was subtracted from each measurement, and % LDH release was calculated as % cytotoxicity = 100 × Experimental LDH Release (OD490) / Maximum LDH Release (OD490).

### Microscopy

HeLa cells were plated at a density of 8,000 cells/well in a 96-well cover glass-bottomed plate the day before the experiment. For standard localization imaging, cells were treated with PMO-Sulfo-Cy5-CPP conjugates at 5 µM or 25 µM for 30 min at 37 °C and 5% CO_2_. Each well was washed with media and incubated in fresh media for 1 h before Hoechst (nuclear) and Lysotracker Green (endosomal) fluorescent tracking dyes were added, and imaged immediately. The endosomal release experiment was adapted from a previously reported protocol.^54^ Cells were treated with 50 µM 7-Diethylaminocoumarin-3-carboxylic acid (DEAC)-k5 for 1 h at 37 °C and 5% CO_2_ before being washed with media. Then, PMO-Sulfo-Cy5-CPP conjugates were added as before. Sytox Green was added immediately before imaging in order to exclude observation of nonviable cells. Imaging was performed at the Whitehead Institute’s Keck Imaging Facility on an RPI Spinning Disk Confocal Microscope at 40x objective.

### Uptake Assay

#### Cell treatment

Cells were plated either in 6-well or 12-well plates at a density such that they reached 80% confluency the following day. CPP or PMO-CPP stock solutions were made fresh to 1 mM in cation-free PBS, as determined by UV-Vis. Treatment solution was then prepared by adding the stock solution to cell media at the concentrations described. Two wells were left untreated as controls. The plates were then incubated at 37 °C and 5% CO_2_ for the designated time. For the experiment to arrest energy-dependent uptake, the plate was incubated at 4 °C. Following incubation, the cells were washed three times with media, followed by 0.1 mg/mL heparin in PBS for 5 min. Supernatant was aspirated and cells were lifted by incubating in trypsin-EDTA for 10 min at 37 °C. Trypsin was quenched by adding cell media, and cells were transferred to Eppendorf tubes and pelleted at 500 rcf for 3 min. Pellets were washed by mixing with PBS, repeated twice.

#### Lysis

To acquire whole cell lysate, 50 µL RIPA buffer, protease inhibitor cocktail, water) was added to the cell pellet, mixed gently, and placed on ice for 1 h. To extract the cytosol, 50 µL digitonin buffer (0.05 mg/mL digitonin, 250 mM sucrose, PBS) was added to a cell pellet, mixed very gently, and placed on ice for 10 min. Samples were then pelleted by centrifugation at 16,000 rcf for 5 min. Supernatants were transferred to new Eppendorf tubes and kept on ice. Extracted protein from the cell-only control samples was quantified using Pierce Rapid Gold BCA Protein Assay Kit (Thermo Fisher). 10 µg protein from each sample was then analyzed by sodium dodecyl sulfate– polyacrylamide gel electrophoresis (SDS-PAGE) gel for 35 min at 165 V and then transferred to a nitrocellulose membrane soaked in 48 mM Tris, 39 mM glycine, 0.0375% SDS, 20% methanol using a TransBlot Turbo Semi-Dry Transfer Unit (BioRad) for 7 min. The membrane was blocked at 4 °C overnight in LI-Cor Odyssey blocking buffer in PBS. The membrane was then immunostained for 1 h with anti-Erk1/2 and anti-Rab5 (Cell Signaling) in PBS-Tween at room temperature. After incubation, the membrane was washed three times with TBST and incubated with the appropriate secondary antibody in TBST for 1 h at room temperature, then washed with TBST. The membrane was imaged on a ChemiDoc MP Imaging System (Bio-Rad).

#### Penetration Selection

10 µL Dynabeads™ MyOne™ Streptavidin T1 (Thermo Fisher) were transferred to tubes in a magnet stand and washed with PBS. Cell extracts were added to the corresponding bead-containing tube and rotated at 4 °C for 2 h. To two of the cell only sample lysates was added each 0.5 µL of PMO-biotin-library and 0.5 µL of biotin-library, and was combined with 50 µL of Streptavidin beads. Following pulldown, the beads were washed with 6 M guanidinium chloride (GuHCl, pH 6.8, 2 × 200 µL) and suspended in 100 µL PBS. Then, NaIO4 (1 mM in H_2_O, 2 µL) was added and the beads incubated for 5 min in absence of light, followed by quench solutions: Na_2_SO_3_ (100 mM, 5 µL) and NH_2_OH (100 mM, 5 µL). The supernatants were transferred to new tubes and the beads were washed with 6 M GuHCl (2 × 100 µL). Pooled supernatant fractions were then desalted by solid-phase extraction (SPE) using C18 ZipTips, lyophilized, and rehydrated in 10 µL 1 M GuHCl in water containing 0.1% formic acid.

#### Orbitrap LC-MS/MS

Analysis was performed on an EASY-nLC 1200 (Thermo Fisher Scientific) nano-liquid chromatography handling system connected to an Orbitrap Fusion Lumos Tribrid Mass Spectrometer (Thermo Fisher Scientific). Samples were run on a PepMap RSLC C18 column (2 μm particle size, 15 cm × 50 μm ID; Thermo Fisher Scientific, P/N ES901). A nanoViper Trap Column (C18, 3 μm particle size, 100 Å pore size, 20 mm × 75 μm ID; Thermo Fisher Scientific, P/N 164946) was used for desalting. The standard nano-LC method was run at 40 °C and a flow rate of 300 nL/min with the following gradient: 1% solvent B in solvent A ramping linearly to 41% B in A over 55 min, where solvent A = water (0.1% FA), and solvent B = 80% acetonitrile, 20% water (0.1% FA). Positive ion spray voltage was set to 2200 V. Orbitrap detection was used for primary MS, with the following parameters: resolution = 120,000; quadrupole isolation; scan range = 150–1200 m/z; RF lens = 30%; AGC target = 250%; maximum injection time = 100 ms; 1 microscan. Acquisition of secondary MS spectra was done in a data-dependent manner: dynamic exclusion was employed such that a precursor was excluded for 30 s if it was detected four or more times within 30 s (mass tolerance: 10.00 ppm); monoisotopic precursor selection used to select for peptides; intensity threshold was set to 2 × 10^4^; charge states 2–10 were selected; and precursor selection range was set to 200–1400 m/z. The top 15 most intense precursors that met the preceding criteria were subjected to subsequent fragmentation. Two fragmentation modes— higher-energy collisional dissociation (HCD), and electron-transfer/higher-energy collisional dissociation (EThcD)—were used for acquisition of secondary MS spectra. Detection was performed in the Orbitrap (resolution = 30,000; quadrupole isolation; isolation window = 1.3 m/z; AGC target = 2 × 10^4^; maximum injection time = 100 ms; 1 microscan). For HCD, a stepped collision energy of 3, 5, or 7% was used. For EThcD, a supplemental activation collision energy of 25% was used.

#### De novo peptide sequencing and filtering

De novo peptide sequencing of the acquired data was performed in PEAKS 8 (BioInformatics Solutions Inc.). Using PEAKS, spectra were prefiltered to remove noise, and sequenced. All non-canonical amino acids were sequenced as post-translational modifications based on the canonical amino acid most closely matching their molecular mass. Norleucine and β-Alanine were sequenced as leucine and alanine, respectively. In addition, His-oxide and Met-oxide were allowed as variable post-translational modifications. Twenty candidate sequence assignments were created for each secondary scan.

An automated Python-based routine was used for postprocessing data analysis to eliminate noise, synthetic impurities, duplicates, resolve certain sequencing ambiguities, and to select the best candidate sequence assignment for each MS/MS scan. The script eliminates all sequence candidates of length other than 9 or 10 and all candidates not bearing the C-terminal KWKK motif. Next, for each remaining spectrum, a single candidate is kept, discarding all other peptides with lower sequencing scores from PEAKS (average local confidence (ALC) scores, from 0 to 99), and duplicate sequences are labeled as non-unique. Finally, the resulting unique sequence assignments are refined further by excluding prominent synthetic impurities that were not eliminated in the previous steps. If two unique sequences have an identifiable main product/side product relationship, the side product is eliminated. In this way, peptides containing oxidized Met residues, deamidation of Gln to Glu, which occasionally happens during saponification of PAM ester, sodium adducts, and a few less prominent side-reactions are identified, and their corresponding sequences are discarded.

### Statistics

Statistical analysis and graphing was performed using Prism (Graphpad) or Excel (Microsoft). Concentration-response curves were fitted using Prism using nonlinear regression. The listed replicates for each experiment indicates the number of distinct samples measured for a given assay, all experiments were performed in triplicate unless otherwise indicated. Significance for activities between constructs was determined using a student’s two-sided, unpaired t-test.

## Acknowledgements

This research was funded by Sarepta Therapeutics. C.K.S. (4000057398) and C.E.F. (4000057441) acknowledge the National Science Foundation Graduate Research Fellowship (NSF Grant No. 1122374) for research support. We acknowledge support from the Swanson Biotechnology Center Flow Cytometry Facility at the Koch Institute for Integrative Cancer Research at MIT through the use of their flow cytometers (NCI Cancer Center Support Grant P30-CA14051). We thank Cassandra Rogers and Brandyn Braswell at the Whitehead Institute’s Keck Imaging facility for use of the confocal microscope.

## Conflict of Interest

B.L.P. is a co-founder and/or member of the scientific advisory board of several companies focusing on the development of protein and peptide therapeutics. A provisional patent disclosure was filed regarding the methodology and compounds described in this study.

## Abbreviations

AS-MS: affinity selection-mass spectrometry
CPPs: cell-penetrating peptides
DBCO: dibenzocyclooctyne
DEAC: 7-Diethylaminocoumarin-3-carboxylic acid
EGFP: enhanced green fluorescent protein
ETD: electron-transfer dissociation
HCD: higher-energy collisional dissociation
LDH: lactate dehydrogenase
MALDI-ToF: Matrix-assisted laser esorption/ionization – time of flight
nLC-MS/MS: nano-liquid chromatography-tandem mass spectrometry
OBOC: one-bead one-compound
PMO: phosphorodiamidate morpholino oligomer
PS-MS: Penetration-selection mass spectrometry
SulfoCy5: Sulfo-Cyanine 5 fluorophore

