## Supporting Information for "In-cell penetration selection—mass spectrometry produces noncanonical peptides for antisense delivery"

#### Supplemental Figures

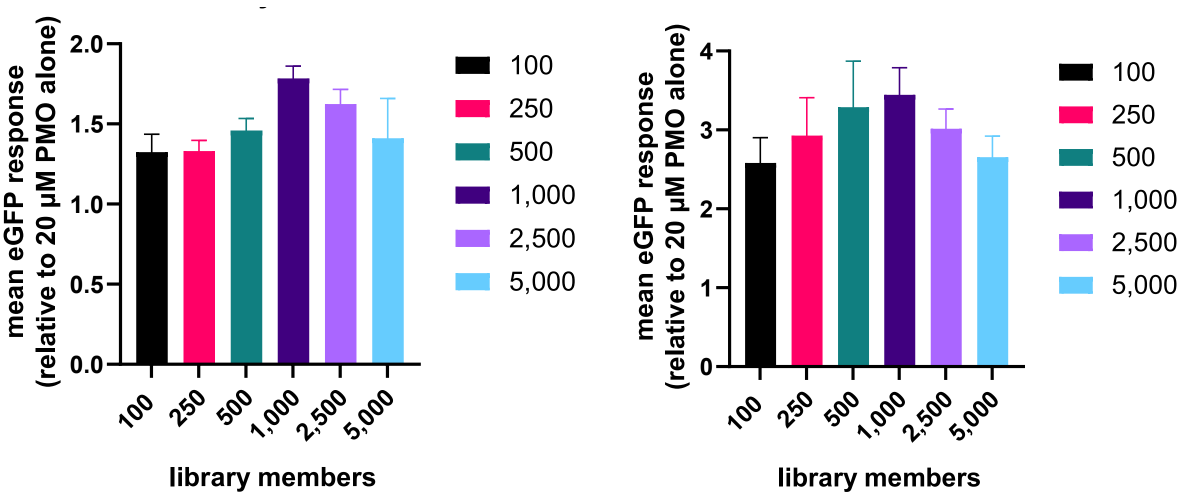

**Figure S1 Biological replicates 1 and 2 of EGFP assay with peptide libraries 100-5,000 members (replicates of Figure 3C)**. HeLa 654 cells were treated with 20 µM PMO-Library of varying member sizes or 20 µM PMO alone for 22 h prior to flow cytometry. Results are given relative to the fluorescence of PMO-treated cells. Bars represent mean ± SD, N = 3. There is no significant difference ($p < 0.01$) in EGFP fluorescence between the libraries of different sizes (N=3, student's two-sided, unpaired t-test). The two graphs represent independent biological replicates, performed and measured on different days.

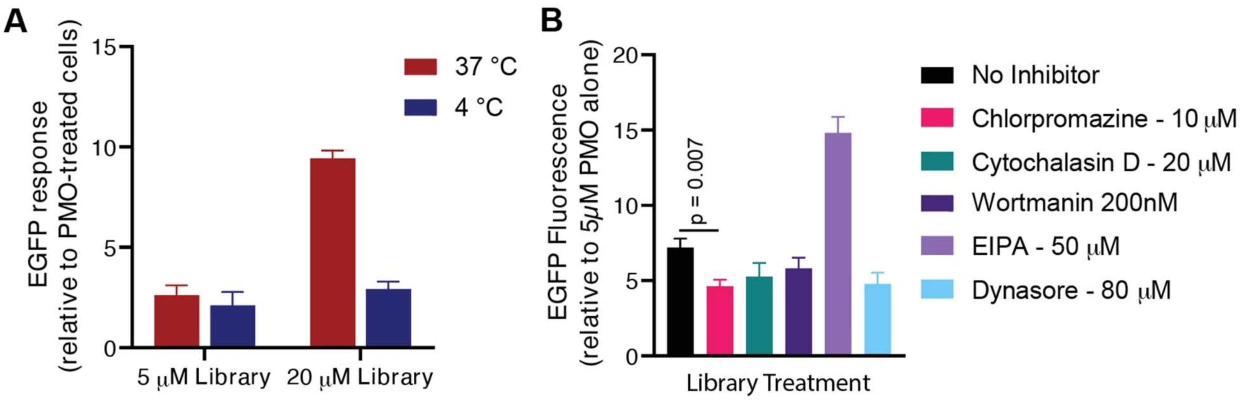

**Figure S2 Biological replicate of EGFP assay with peptide library incubated at 4˚ or 37˚ C (replicate of Figure 3D and 3E).**(A) HeLa 654 cells were pre-incubated at 4 ˚C or 37 ˚C for 30 min prior to treatment with 20 µM PMO-Library or 20 µM PMO alone for 2 h at the indicated temperature. After treatment, cells were washed with 0.1 mg/mL heparin and incubated in media for 22 h prior to flow cytometry. Results are given relative to the fluorescence of PMO-treated cells. Bars represent mean ± SD, N = 3. There was a significant difference ($p<0.0001$) between the 4 ˚C and 37 ˚C treatment conditions at 20 µM of PMO-library (N=3, student's two-sided, unpaired t-test). (B) HeLa 654 cells were pre-incubated for 30 min with the indicated compound and then 10 µM PMO-library (1,000 members) was added. After treatment with the construct for 3 h, the cells were washed with PBS and 0.1 mg/mL heparin and the media was exchanged for fresh, untreated media for 22 h prior to flow cytometry. Results are given normalized to the fluorescence of cells treated with 10μM PMO. Bars represent mean ± SD, N = 3. At 10 μM chlorpromazine, EGFP fluorescence significantly decreased (p=0.007, N=3, student's two-sided, unpaired t-test).

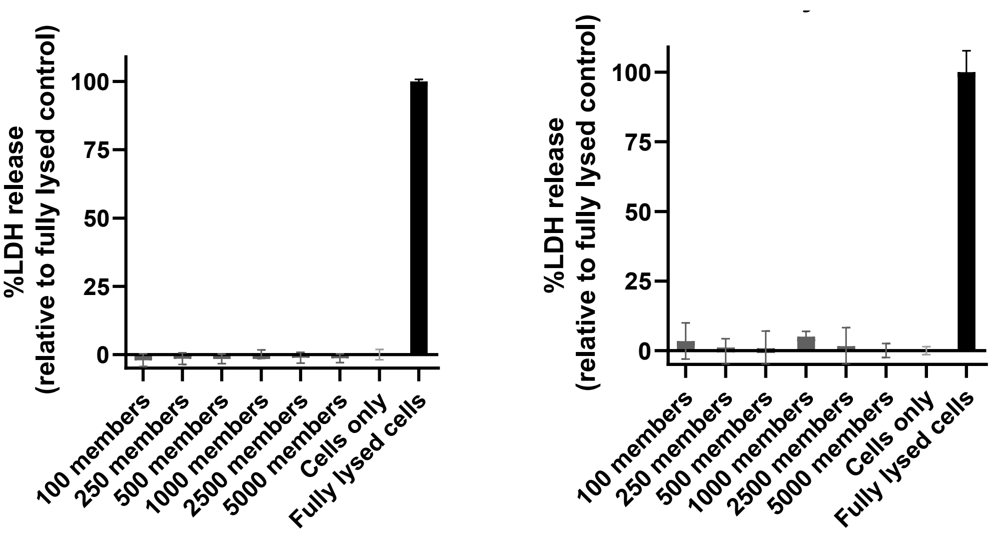

**Figure S3 LDH release toxicity assay of cells treated with peptide libraries of 100-5,000 members (Figures 3C and Figure S1)**. LDH assay conducted on supernatant of cells treated for the EGFP assay shown in Fig 3C and Fig S1. Concentration of PMO conjugates was 20 μM. Bars represent mean ± SD, N = 3. Results are given as LDH release above vehicle relative to fully lysed cells. No compounds showed LDH release significantly above vehicle-treated cells.

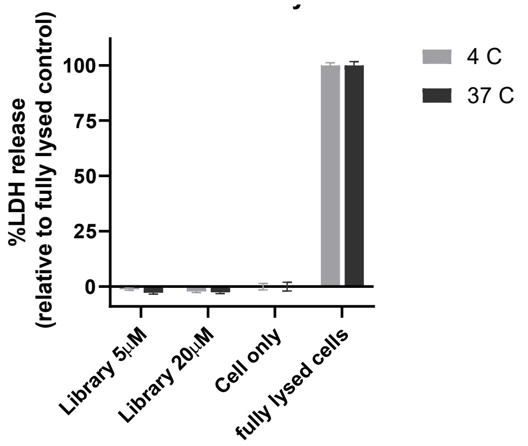

**Figure S4 LDH release toxicity assay of peptide library incubated at 4° or 37° C (Figure 3D and Figure S2)**. LDH assay conducted on supernatant of cells treated for the EGFP assay shown in Fig 3D and Fig S2. HeLa 654 cells were pre-incubated at 4 °C or 37 °C for 30 min prior to treatment with 20 μM PMO-Library or 20 μM PMO alone for 2 h at the indicated temperature. After treatment, cells were washed with 0.1 mg/mL heparin and incubated in media for 22 h prior to flow cytometry. Bars represent mean ± SD, N = 3. Results are given as LDH release above vehicle relative to fully lysed cells. No compounds showed LDH release significantly above vehicle-treated cells.

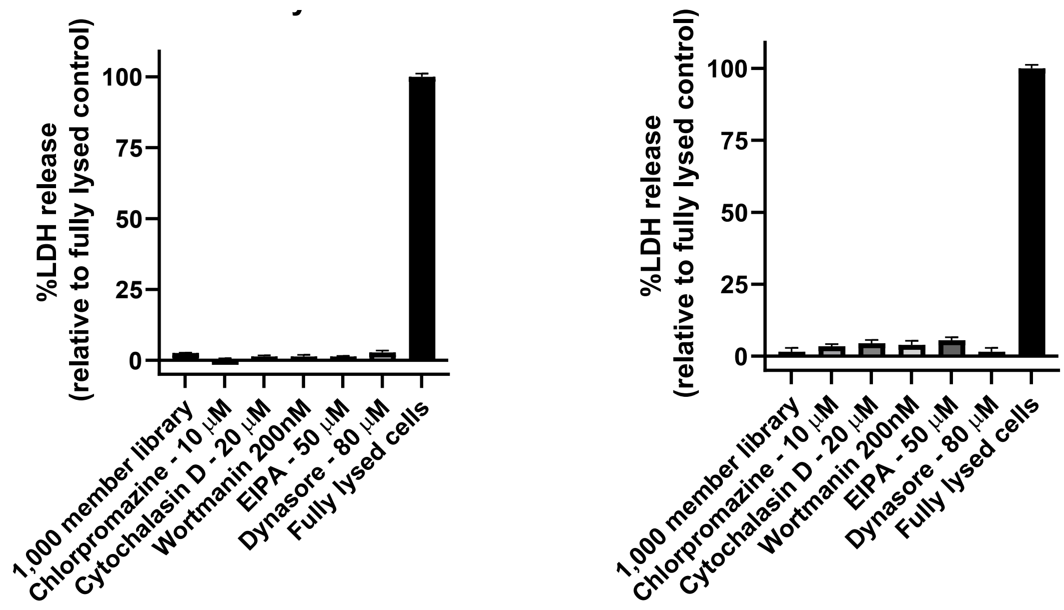

**Figure S5 LDH release toxicity assay of peptide library co-incubated with endocytosis inhibitors (Figure 3E and Figure S2)** LDH assay conducted on supernatant of cells treated for the EGFP assay shown in Fig 3E and Fig S2. HeLa 654 cells were pre-incubated for 30 min with the indicated compound and then 10 μM PMO-library (1,000 members) was added. After treatment with the construct for 3 h, the cells were washed with PBS and 0.1 mg/mL heparin and the media was exchanged for fresh, untreated media for 22 h prior to flow cytometry. Bars represent mean ± SD, N = 3. Results are given as LDH release above vehicle relative to fully lysed cells. No LDH release significantly above vehicle-treated cells was demonstrated for the incubation with chlorpromazine.

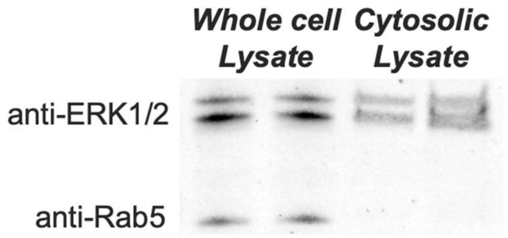

**Figure S6 Extraction of the cytosol was verified via Western blot**. The protein in the no treatment control lysates were analyzed via sodium dodecyl sulfate–polyacrylamide gel electrophoresis (SDS-PAGE) and Western blot to visualize presence of ERK1/2 (cytosolic marker) and Rab5 (an endosomal marker). Rab5 is observed in the whole cell lysate but not in the cytosolic extract.

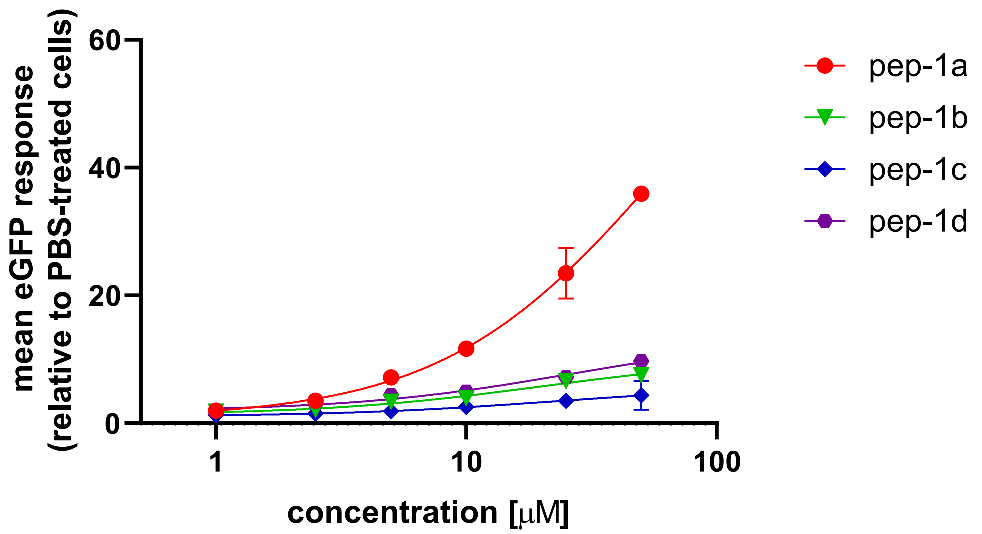

**Figure S7 Biological replicate of EGFP assay with candidate peptides at increasing concentrations (replicate of Figure 5C)** HeLa 654 cells were treated with 1, 2.5, 5, 10, 25, or 50 µM PMO-CPP for 22 h prior to flow cytometry. Results are given as the mean EGFP fluorescence of cells treated with PMO-peptide relative to the fluorescence of cells treated with vehicle only. Bars represent mean ± SD, N = 3.

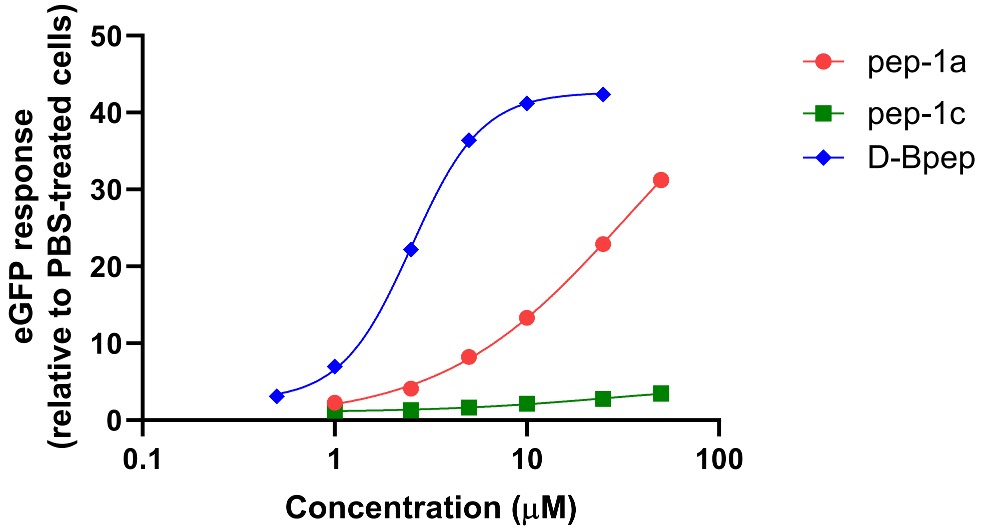

**Figure S8 Biological replicate of EGFP assay with pep1a, pep1c, and Bpep (replicate of Figure 5E)** HeLa 654 cells were treated with 0.5, 1, 2.5, 5, 10, 25, or 50 µM PMO-CPP for 22 h. Results are given as the mean EGFP fluorescence of cells treated with PMO-peptide relative to the fluorescence of cells treated with vehicle only. Bas represent mean ± SD, N = 3.

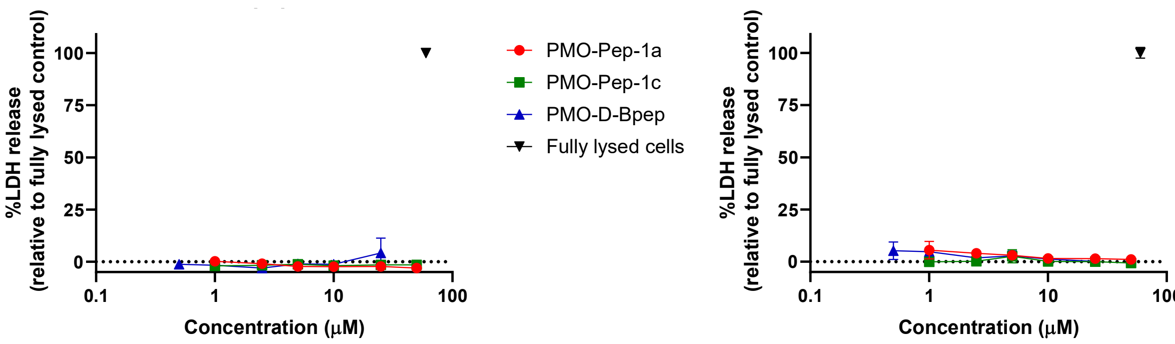

**Figure S9 LDH release toxicity assay with pep1a, pep1c, and Bpep (Figure 7C and Figure S8)** LDH assay conducted on supernatant of cells treated for the EGFP assay shown in Fig 7C and Fig S8. HeLa 654 cells were treated with 0.5, 1, 2.5, 5, 10, 25, or 50 µM PMO-CPP for 22 h. Bars represent mean ± SD, N = 3. Results are given as LDH release above vehicle relative to fully lysed cells.

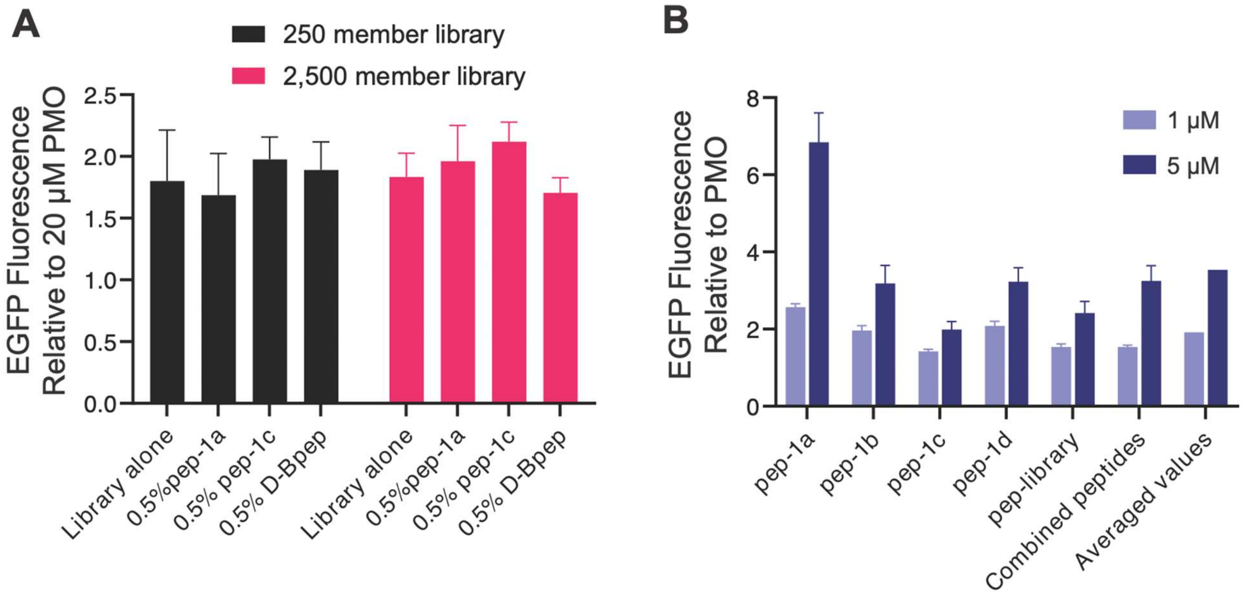

**Figure S10 Activity of one peptide does not influence library activity**. (A) HeLa 654 cells were treated with 20 µM PMO-Library or 19.9 µM PMO-Library and 0.1 µM PMO-CPP for 22 h prior to flow cytometry. Results are given as the mean EGFP fluorescence of cells treated with PMO-peptide relative to the fluorescence of cells treated with vehicle only. Bars represent mean ± SD, N = 3. All 3 PMO-CPP treatment conditions are not significantly different from the library alone. (N=3, student's two-sided, unpaired t-test). (B) HeLa 654 cells were treated with 1 or 5 µM PMO-CPP or a combined solution of 5 PMO-CPPs for 22 h prior to flow cytometry. Results are given as the mean EGFP fluorescence of cells treated with PMO-peptide relative to the fluorescence of cells treated with vehicle only. Bars represent mean ± SD, N = 3. Indicated concentration represents total PMO-CPP present in the sample. All PMO-CPPs at 1 µM showed significantly less PMO delivery activity than 5 µM of the combined peptides ($p < 0.05$).

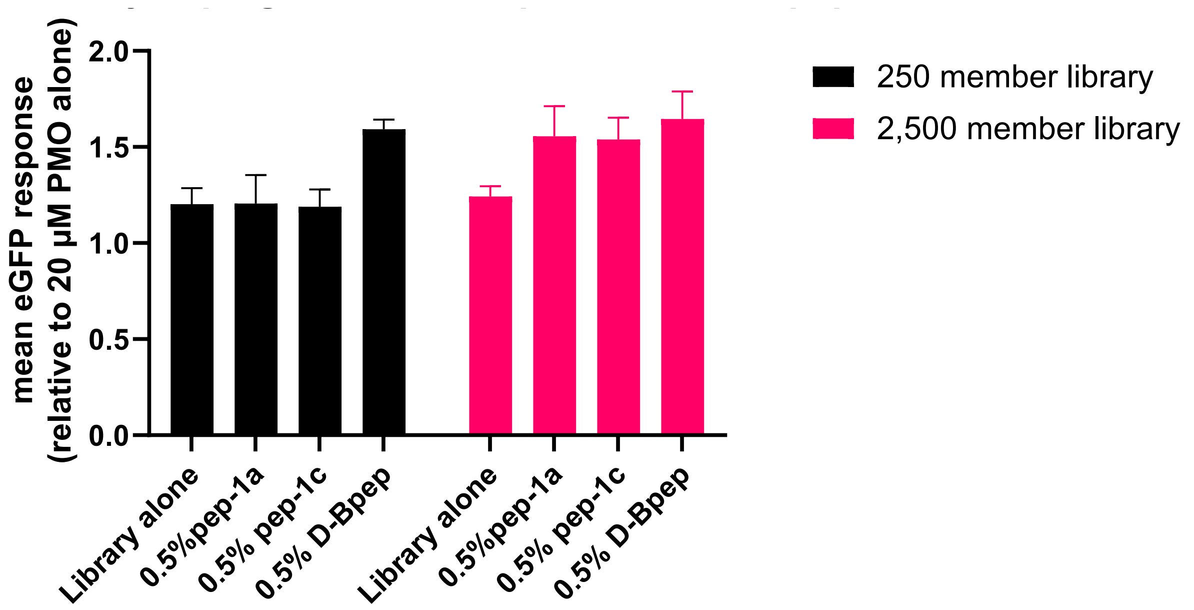

**Figure S11 Biological replicate of EGFP assay of peptide library doped with penetrant peptides (replicate of Figure S10)** HeLa 654 cells were treated with 20 µM PMO-Library or 19.9 µM PMO-Library and 0.1 µM PMO-CPP for 22 h prior to flow cytometry. Results are given as the mean EGFP fluorescence of cells treated with PMO-peptide relative to the fluorescence of cells treated with vehicle only. Bars represent mean ± SD, N = 3. All 3 PMO-CPP treatment conditions are not significantly different ($p < 0.01$) from the library alone (N=3, student's two-sided, unpaired t-test).

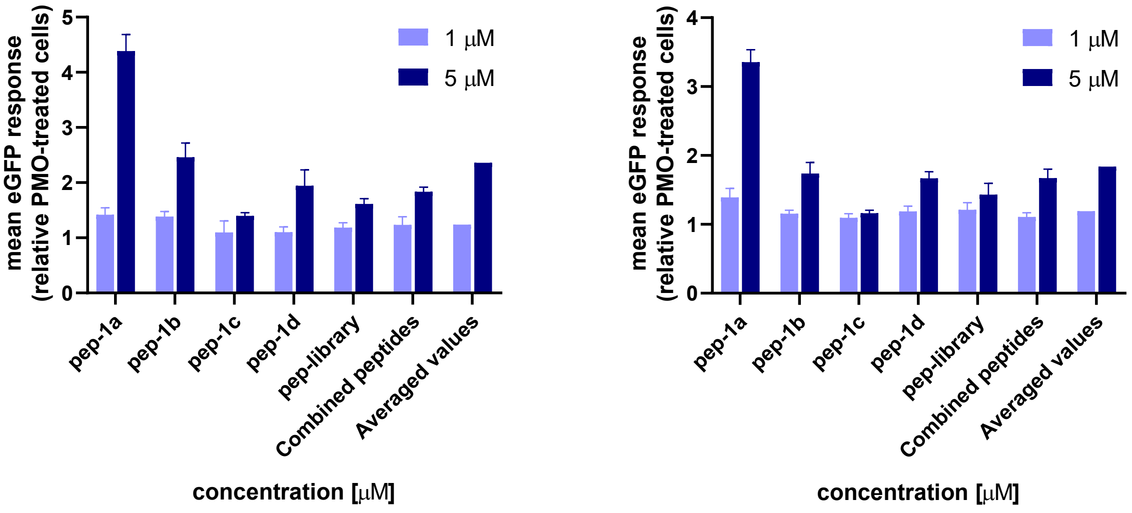

**Figure S12 Biological replicates 1 and 2 of EGFP assay with candidate peptides at 1 or 5µM (replicates of Figure S10)** HeLa 654 cells were treated with 1 or 5 µM PMO-CPP or a combined solution of 5 PMO-CPPs for 22 h prior to flow cytometry. Results are given as the mean EGFP fluorescence of cells treated with PMO-peptide relative to the fluorescence of cells treated with vehicle only. Bars represent mean ± SD, N = 3. Indicated concentration represents total PMO-CPP present in the sample. All PMO-CPPs at 1 µM showed significantly less PMO delivery activity than 5 μM of the combined peptides ($p < 0.05$). The two graphs represent independent biological replicates, performed and measured on different days.

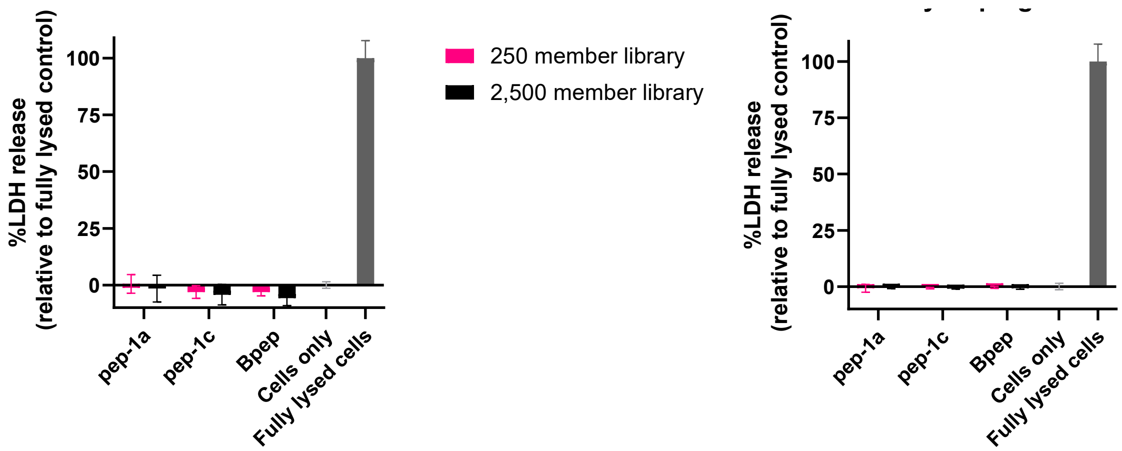

**Figure S13 LDH release toxicity assay of peptide library doped with penetrant peptides (Figures S10A and S11)** LDH assay conducted on supernatant of cells treated for the EGFP assay shown in Fig S10A and Fig S11. HeLa 654 cells were treated with 20 μM PMO-Library or 19.9 μM PMO-Library and 0.1 μM PMO-CPP for 22 h prior to flow cytometry. Bars represent mean ± SD, N = 3. Results are given as LDH release above vehicle relative to fully lysed cells.

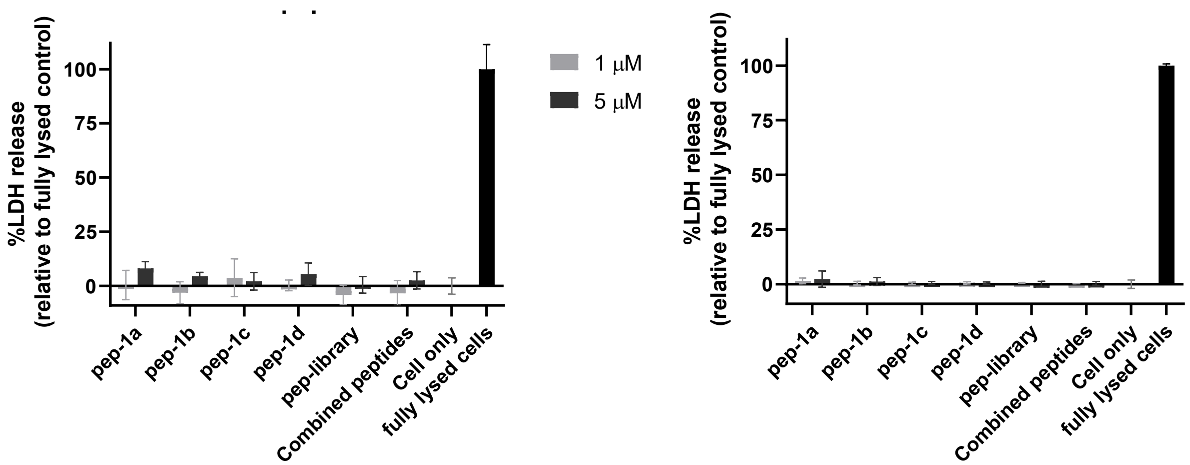

**Figure S14 LDH release toxicity assay with candidate peptides at 1 or 5μM (Figure S10B and S12)** LDH assay conducted on supernatant of cells treated for the EGFP assay shown in Fig S10B and Fig S12. HeLa 654 cells were treated with 1 or 5 μM PMO-CPP or a combined solution of 5 PMO-CPPs for 22 h prior to flow cytometry. Results are given as the mean EGFP fluorescence of cells treated with PMO-peptide relative to the fluorescence of cells treated with vehicle only. Indicated concentration represents total PMO-CPP present in the sample. Bars represent mean ± SD, N = 3. Results are given as LDH release above vehicle relative to fully lysed cells.

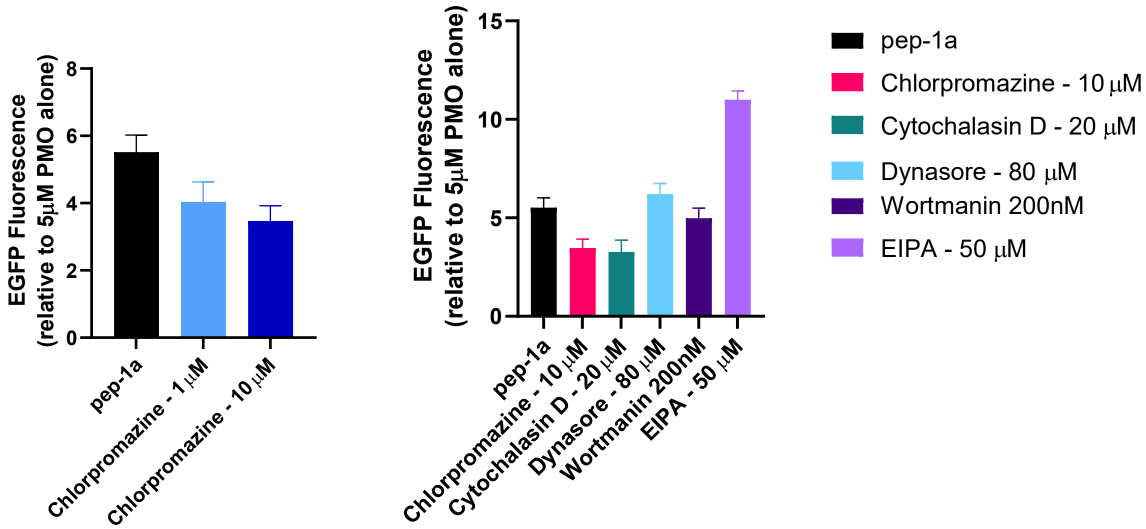

**Figure S15 Biological replicate of EGFP assay of pep1a co-incubated with endocytosis inhibitors (replicate of Figure 6A)** HeLa 654 cells were pre-incubated for 30 min with the indicated compound and then 5 µM PMO-Pep1a was added. After treatment with the construct for 3 h, the cells were washed with 0.1 mg/mL heparin and the media was exchanged for fresh, untreated media for 22 h prior to flow cytometry. Data are shown as EGFP mean fluorescence intensity relative to 5 µM PMO for cells treated with different endocytosis inhibitors. Bars represent mean ± SD, N = 3. At 10 µM chlorpromazine, EGFP fluorescence significantly decreased (p=0.0066, N=3, student's two-sided, unpaired t-test), and this decrease is demonstrated to be dose-dependent.

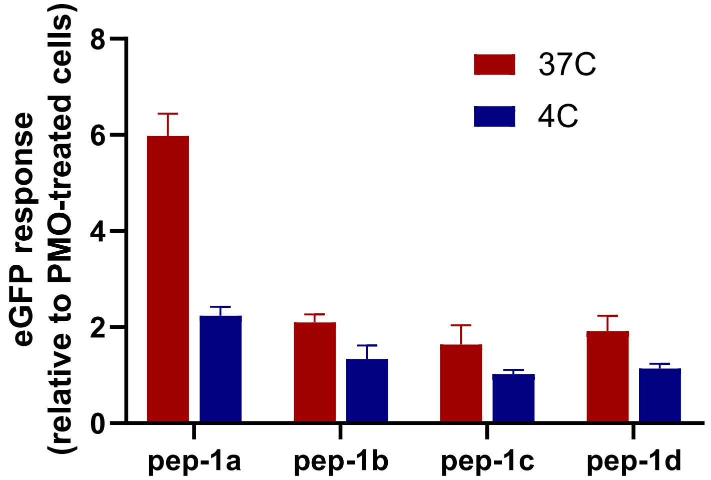

**Figure S16 Biological replicate of EGFP assay of candidate peptides treated at 4° or 37°C (replicate of Figure 6B)** HeLa 654 cells were pre-incubated for 30 min at 4 °C or 37 °C, followed by the addition of PMO-peptide conjugate to each well at a concentration of 5 µM. After incubation at 4°C or 37 °C for 2 h, the cells were washed with 0.1 mg/mL heparin and the media was exchanged for fresh, untreated media for 22 h prior to flow cytometry. Data are shown as EGFP mean fluorescence intensity relative to 5 µM PMO at the indicated temperature. Bars represent mean ± SD, N = 3.

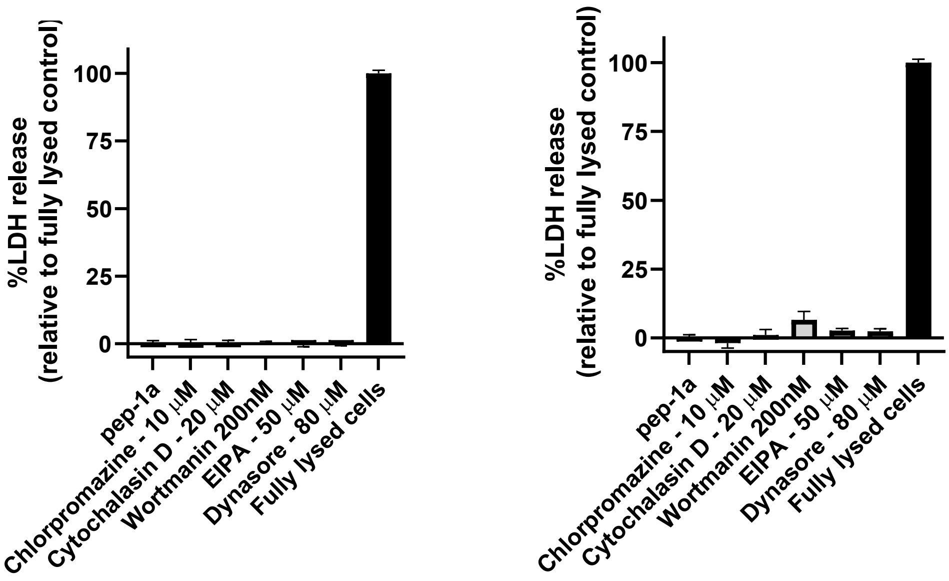

**Figure S17 LDH release toxicity assay of pep1a co-incubated with endocytosis inhibitors (Figure 6A and Figure S15)** LDH assay conducted on supernatant of cells treated for the EGFP assay shown in Fig 6A and Fig S15. HeLa 654 cells were pre-incubated for 30 min with the indicated compound and then 5 μM PMO-Pep1a was added. After treatment with the construct for 3 h, the cells were washed with 0.1 mg/mL heparin and the media was exchanged for fresh, untreated media for 22 h prior to flow cytometry. Bars represent mean ± SD, N = 3. Results are given as LDH release above vehicle relative to fully lysed cells. There is no significant LDH release from chlorpromazine-treated cells.

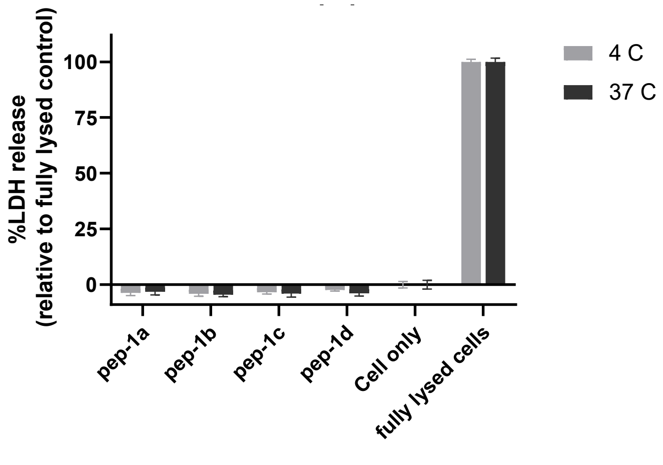

**Figure S18 LDH release toxicity assay with candidate peptides treated at 4° or 37°C (Figure 6)** LDH assay conducted on supernatant of cells treated for the EGFP assay shown in Fig 9B. HeLa 654 cells were pre-incubated for 30 min at 4 ˚C or 37 ˚C, followed by the addition of PMO-peptide conjugate to each well at a concentration of 5 μM. After incubation at 4˚C or 37 ˚C for 2 h, the cells were washed with 0.1 mg/mL heparin and the media was exchanged for fresh, untreated media for 22 h prior to flow cytometry. Bars represent mean ± SD, N = 3. Results are given as LDH release above vehicle relative to fully lysed cells.

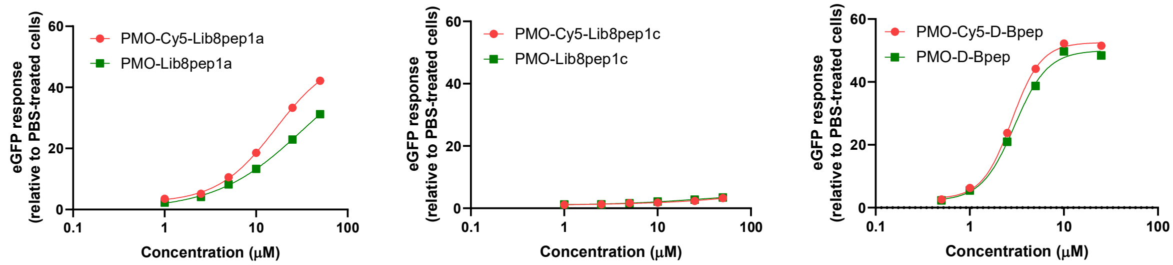

**Figure S19 Biological replicate of EGFP assay with candidate peptides with and without the sulfo-Cy5 fluorophore appended (replicate of Figure 7B-D)** HeLa 654 cells were treated with 1, 2.5, 5, 10, 25, or 50 μM PMO-CPP or PMO-SulfoCy5-CPP for 22 h prior to flow-cytometry. Results are given as the mean EGFP fluorescence of cells treated with PMO-peptide relative to the fluorescence of cells treated with vehicle only. Bars represent mean ± SD, N = 3. No peptides had significant differences in EC50 between SulfoCy5 and unlabeled constructs.

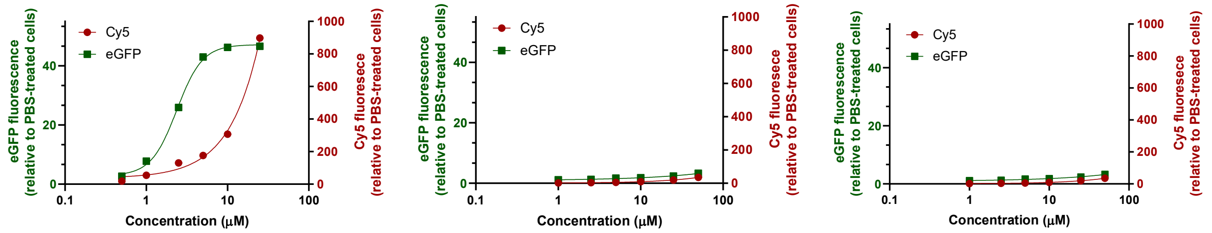

**Figure S20 Biological replicate of EGFP assay with Cy5-labelled candidate peptides comparing total Cy5 fluorescence to EGFP fluorescence (replicate of Figure 7E-G)** HeLa 654 cells were treated with 1, 2.5, 5, 10, 25, or 50 μM PMO-SulfoCy5-CPP for 22 h prior to flow cytometry. Results are given as the mean fluorescence of cells treated with PMO-SulfoCy5-peptide relative to the fluorescence of cells treated with vehicle only for each channel. Bars represent mean ± SD, N = 3.

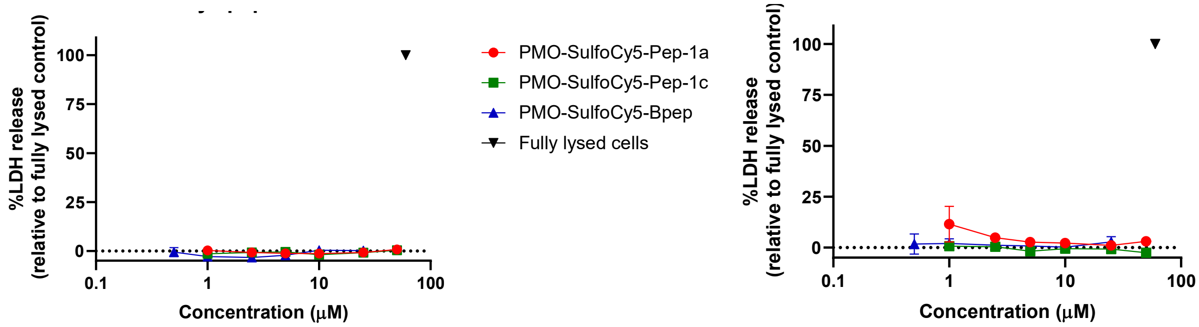

**Figure S21 LDH release toxicity assay with candidate peptides with the sulfo-Cy5 fluorophore appended (Figure 7B-D, Figure 7E-G, Figure S19, Figure S20)** LDH assay conducted on supernatant of cells treated for the EGFP assay shown in Fig 7, Fig S19, and Fig S20. HeLa 654 cells were treated with 1, 2.5, 5, 10, 25, or 50 μM PMO-CPP or PMO-SulfoCy5-CPP for 22 h prior to flow-cytometry. Bars represent mean ± SD, N = 3. Results are given as LDH release above vehicle relative to fully lysed cells.

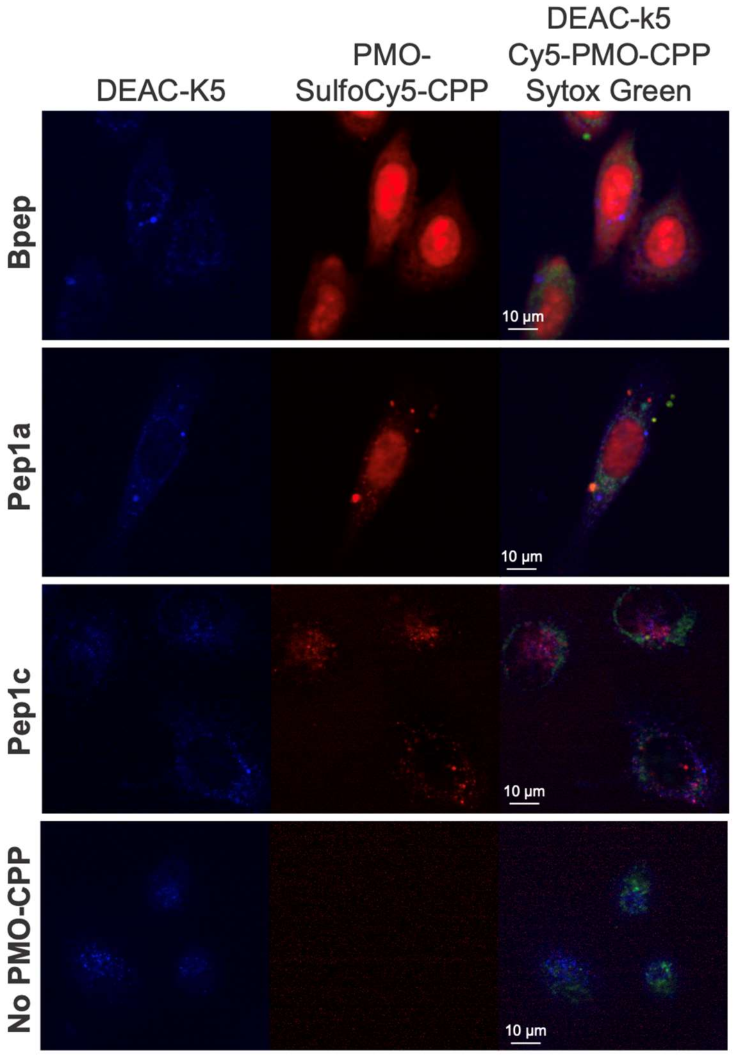

**Figure S22 PMO-CPPs do not appear to permeabilize endosomes for general cargo release.** Confocal micrographs of HeLa cells treated with 50 μM DEAC-k5 (endosome localizing peptide) followed by 25 μM of PMO-SulfoCy5-CPPs. All conjugates demonstrate fluorescent puncta likely due to accumulation in endosomes, but Bpep and Pep1a show intense nuclear staining, indicating endosomal escape. However, the DEAC-k5 appears to remain as puncta and does not show diffuse fluorescence in cytosol or nucleus. Sytox Green was added to exclude observation of dead cells.

### Appendix I: Gel Images

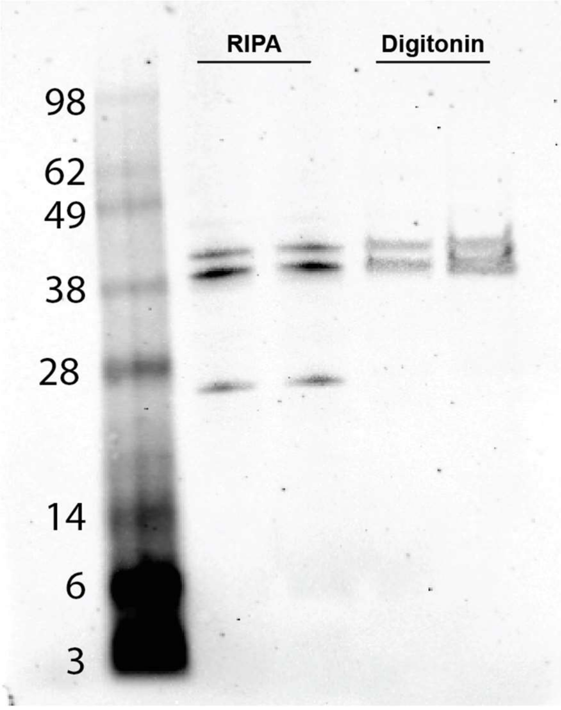

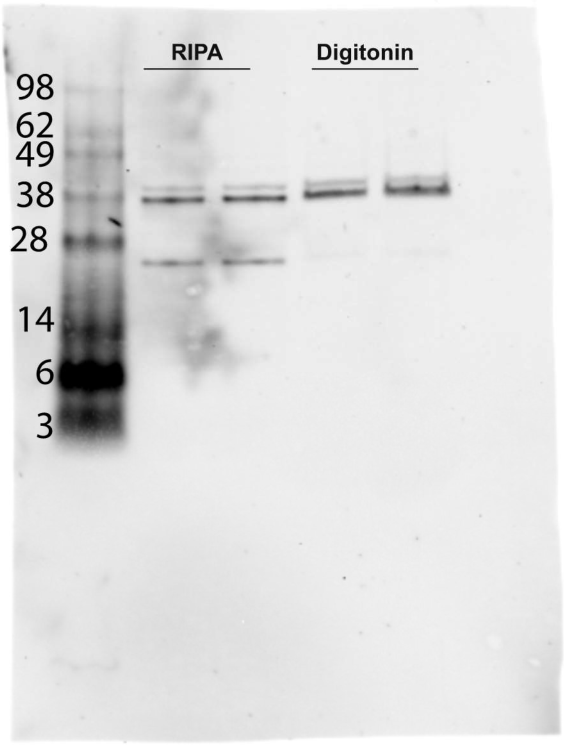
RIPA
Digitonin
98
62
49
38
28
14
6
3

### Appendix II: Peptide Library sequencing

All unique peptides found through sequencing a 500-member portion of Library 8 prior to appending the isoseramox and linker. D-residues are given their one-letter abbreviation, while non-canonical residues are given the three-letter abbreviation listed in Figure 2.

Legend:

| Monomer code | Monomer name |
|---|---|
| A | Beta-alanine |
| D | D-Asp |
| E | D-Glu |
| F | D-Phe |
| G | Gly |
| H | D-His |
| K | D-Lys |
| L | Norleucine |
| N | D-Asn |
| P | D-Pro |
| Q | D-Gln |
| R | D-Arg |
| S | D-Ser |
| T | D-Thr |
| V | D-Val |
| a | Dab |
| b | Pip |
| c | Gba |
| d | Amf |
| e | Nap |
| f | Dpa |
| g | Hle |
| h | Abu |
| i | Oxidized His |
| j | Oxidize Trp |
| k | Oxidized Nap |
| l | Oxidized Dpa |
| m | Oxidized Amf |
| X or Z | C-terminal amide |

| Scan | Peptide | ALC (%) | m/z | z | RT | Mass | ppm |
|---|---|---|---|---|---|---|---|
| 7553 | gbDeDfKWKKZ | 99 | 373.7082 | 4 | 35.06 | 1490.807 | -2.4 |
| 7566 | gbDeDfKWKKZ | 99 | 373.7083 | 4 | 35.11 | 1490.807 | -2.2 |
| 7582 | gbDeDfKWKKZ | 99 | 373.7081 | 4 | 35.18 | 1490.807 | -2.7 |
| 7598 | gbDeDfKWKKZ | 99 | 373.7082 | 4 | 35.24 | 1490.807 | -2.4 |

| | | | | | | | |
|---|---|---|---|---|---|---|---|
| 5323 | fFKFNTKWKKZ | 98 | 362.9596 | 4 | 26.04 | 1447.813 | -2.3 |
| 5420 | eLhPLdKWKKZ | 98 | 343.2176 | 4 | 26.42 | 1368.843 | -1.6 |
| 5482 | fNTRfNKWKKZ | 98 | 380.7128 | 4 | 26.68 | 1518.825 | -1.8 |
| 6373 | eTFLGEKWKKZ | 98 | 444.9195 | 3 | 30.31 | 1331.739 | -1.7 |
| 6516 | fFKeSDKWKKZ | 98 | 495.9386 | 3 | 30.88 | 1484.797 | -2 |
| 7595 | GfAfFQKWKKZ | 98 | 479.9317 | 3 | 35.23 | 1436.776 | -1.6 |
| 8322 | DTALffKWKKZ | 98 | 478.9348 | 3 | 38.22 | 1433.786 | -2.3 |
| 9306 | ePLEegKWKKZ | 98 | 362.9661 | 4 | 42.47 | 1447.838 | -1.9 |
| 7543 | gbDeDfKWKKZ | 98 | 373.7083 | 4 | 35.02 | 1490.807 | -2.2 |
| 5251 | gFgNTQKWKKZ | 97 | 444.942 | 3 | 25.75 | 1331.808 | -2.9 |
| 5515 | NLhLgQKWKKZ | 97 | 423.6107 | 3 | 26.82 | 1267.813 | -2.2 |
| 5991 | DfdDfKKWKKZ | 97 | 523.6174 | 3 | 28.76 | 1567.834 | -2.2 |
| 6100 | SaeDfQKWKKZ | 97 | 360.4453 | 4 | 29.21 | 1437.756 | -2.4 |
| 6159 | FSfGgRKWKKZ | 97 | 347.2096 | 4 | 29.45 | 1384.813 | -2.9 |
| 6587 | GfLHfGKWKKZ | 97 | 466.9314 | 3 | 31.16 | 1397.776 | -2.6 |
| 6771 | SEfTeGKWKKZ | 97 | 461.579 | 3 | 31.9 | 1381.718 | -2.1 |
| 7083 | gTEgfPKWKKZ | 97 | 348.9647 | 4 | 33.15 | 1391.833 | -2.4 |
| 7216 | GFPLEfKWKKZ | 97 | 452.2594 | 3 | 33.67 | 1353.76 | -2.4 |
| 7805 | egTQeEKWKKZ | 97 | 489.9422 | 3 | 36.08 | 1466.807 | -1.7 |
| 8065 | egTQeEKWKKZ | 97 | 367.7082 | 4 | 37.12 | 1466.807 | -2.5 |
| 8706 | DTALffKWKKZ | 97 | 478.935 | 3 | 39.88 | 1433.786 | -1.9 |
| 8757 | fegQbFKWKKZ | 97 | 512.9662 | 3 | 40.11 | 1535.881 | -2.4 |
| 5336 | fFKFNTKWKKZ | 97 | 483.6109 | 3 | 26.09 | 1447.813 | -1.4 |
| 5349 | fFKFNTKWKKZ | 97 | 483.6106 | 3 | 26.14 | 1447.813 | -2 |
| 5365 | fFKFNTKWKKZ | 97 | 483.6103 | 3 | 26.2 | 1447.813 | -2.6 |
| 5394 | fFKFNTKWKKZ | 97 | 483.6103 | 3 | 26.32 | 1447.813 | -2.6 |
| 5404 | fFKFNTKWKKZ | 97 | 483.6104 | 3 | 26.36 | 1447.813 | -2.4 |
| 5417 | fFKFNTKWKKZ | 97 | 483.6109 | 3 | 26.41 | 1447.813 | -1.4 |
| 5528 | fNTRfNKWKKZ | 97 | 380.7125 | 4 | 26.87 | 1518.825 | -2.4 |
| 5570 | fNTRfNKWKKZ | 97 | 380.7126 | 4 | 27.04 | 1518.825 | -2.1 |
| 5998 | DfdDfKKWKKZ | 97 | 523.6172 | 3 | 28.79 | 1567.834 | -2.6 |
| 6014 | DfdDfKKWKKZ | 97 | 523.6177 | 3 | 28.85 | 1567.834 | -1.7 |
| 6027 | DfdDfKKWKKZ | 97 | 523.6177 | 3 | 28.91 | 1567.834 | -1.7 |
| 6040 | DfdDfKKWKKZ | 97 | 523.6174 | 3 | 28.96 | 1567.834 | -2.4 |
| 6053 | DfdDfKKWKKZ | 97 | 523.6176 | 3 | 29.01 | 1567.834 | -2 |
| 6066 | DfdDfKKWKKZ | 97 | 523.6173 | 3 | 29.07 | 1567.834 | -2.5 |
| 6079 | DfdDfKKWKKZ | 97 | 523.6174 | 3 | 29.12 | 1567.834 | -2.2 |
| 6088 | DfdDfKKWKKZ | 97 | 523.6179 | 3 | 29.16 | 1567.834 | -1.4 |
| 6103 | SaeDfQKWKKZ | 97 | 480.258 | 3 | 29.22 | 1437.756 | -2.4 |
| 6115 | SaeDfQKWKKZ | 97 | 480.258 | 3 | 29.28 | 1437.756 | -2.5 |
| 6143 | SaeDfQKWKKZ | 97 | 480.2581 | 3 | 29.39 | 1437.756 | -2.2 |
| 6172 | FSfGgRKWKKZ | 97 | 347.2097 | 4 | 29.5 | 1384.813 | -2.5 |
| 6185 | FSfGgRKWKKZ | 97 | 347.2097 | 4 | 29.56 | 1384.813 | -2.5 |

| | | | | | | | |
|---|---|---|---|---|---|---|---|
| 6198 | FSfGgRKWKKZ | 97 | 347.2097 | 4 | 29.61 | 1384.813 | -2.7 |
| 6532 | fFKeSDKWKKZ | 97 | 495.9384 | 3 | 30.94 | 1484.797 | -2.4 |
| 6558 | fFKeSDKWKKZ | 97 | 495.9389 | 3 | 31.05 | 1484.797 | -1.4 |
| 6593 | GfLHfGKWKKZ | 97 | 350.4503 | 4 | 31.19 | 1397.776 | -2.8 |
| 6781 | SEfTeGKWKKZ | 97 | 461.579 | 3 | 31.94 | 1381.718 | -2.1 |
| 6794 | SEfTeGKWKKZ | 97 | 461.579 | 3 | 31.99 | 1381.718 | -2.2 |
| 6807 | SEfTeGKWKKZ | 97 | 461.579 | 3 | 32.04 | 1381.718 | -2.1 |
| 6820 | SEfTeGKWKKZ | 97 | 461.5791 | 3 | 32.09 | 1381.718 | -1.9 |
| 6833 | SEfTeGKWKKZ | 97 | 461.5791 | 3 | 32.14 | 1381.718 | -2 |
| 6846 | SEfTeGKWKKZ | 97 | 461.579 | 3 | 32.19 | 1381.718 | -2.3 |
| 6859 | SEfTeGKWKKZ | 97 | 461.579 | 3 | 32.24 | 1381.718 | -2.2 |
| 6878 | SEfTeGKWKKZ | 97 | 461.579 | 3 | 32.32 | 1381.718 | -2.3 |
| 6894 | SEfTeGKWKKZ | 97 | 461.5793 | 3 | 32.39 | 1381.718 | -1.6 |
| 7585 | gbDeDfKWKKZ | 97 | 497.9421 | 3 | 35.19 | 1490.807 | -2 |
| 7815 | egTQeEKWKKZ | 97 | 489.9422 | 3 | 36.12 | 1466.807 | -1.8 |
| 7828 | egTQeEKWKKZ | 97 | 489.9422 | 3 | 36.17 | 1466.807 | -1.8 |
| 7841 | egTQeEKWKKZ | 97 | 489.9419 | 3 | 36.23 | 1466.807 | -2.5 |
| 7851 | egTQeEKWKKZ | 97 | 367.7082 | 4 | 36.27 | 1466.807 | -2.4 |
| 7864 | egTQeEKWKKZ | 97 | 489.9418 | 3 | 36.32 | 1466.807 | -2.6 |
| 7867 | egTQeEKWKKZ | 97 | 367.7081 | 4 | 36.33 | 1466.807 | -2.9 |
| 7880 | egTQeEKWKKZ | 97 | 489.942 | 3 | 36.38 | 1466.807 | -2.3 |
| 7883 | egTQeEKWKKZ | 97 | 367.7082 | 4 | 36.39 | 1466.807 | -2.6 |
| 7893 | egTQeEKWKKZ | 97 | 489.942 | 3 | 36.43 | 1466.807 | -2.2 |
| 7903 | egTQeEKWKKZ | 97 | 489.9421 | 3 | 36.47 | 1466.807 | -2 |
| 7909 | egTQeEKWKKZ | 97 | 367.7083 | 4 | 36.5 | 1466.807 | -2.4 |
| 7916 | egTQeEKWKKZ | 97 | 489.942 | 3 | 36.52 | 1466.807 | -2.2 |
| 7922 | egTQeEKWKKZ | 97 | 367.7083 | 4 | 36.55 | 1466.807 | -2.4 |
| 7926 | egTQeEKWKKZ | 97 | 489.9422 | 3 | 36.56 | 1466.807 | -1.8 |
| 7935 | egTQeEKWKKZ | 97 | 367.7083 | 4 | 36.6 | 1466.807 | -2.4 |
| 7942 | egTQeEKWKKZ | 97 | 489.942 | 3 | 36.63 | 1466.807 | -2.1 |
| 7948 | egTQeEKWKKZ | 97 | 367.7082 | 4 | 36.65 | 1466.807 | -2.6 |
| 7952 | egTQeEKWKKZ | 97 | 489.9421 | 3 | 36.66 | 1466.807 | -1.9 |
| 7961 | egTQeEKWKKZ | 97 | 367.7083 | 4 | 36.7 | 1466.807 | -2.3 |
| 7965 | egTQeEKWKKZ | 97 | 489.9421 | 3 | 36.71 | 1466.807 | -2 |
| 7974 | egTQeEKWKKZ | 97 | 367.7083 | 4 | 36.75 | 1466.807 | -2.4 |
| 7978 | egTQeEKWKKZ | 97 | 489.9424 | 3 | 36.77 | 1466.807 | -1.3 |
| 7984 | egTQeEKWKKZ | 97 | 367.7085 | 4 | 36.79 | 1466.807 | -1.8 |
| 7991 | egTQeEKWKKZ | 97 | 489.9423 | 3 | 36.82 | 1466.807 | -1.5 |
| 7997 | egTQeEKWKKZ | 97 | 367.7084 | 4 | 36.85 | 1466.807 | -2 |
| 8004 | egTQeEKWKKZ | 97 | 489.942 | 3 | 36.87 | 1466.807 | -2.1 |
| 8020 | egTQeEKWKKZ | 97 | 489.9423 | 3 | 36.94 | 1466.807 | -1.5 |
| 8033 | egTQeEKWKKZ | 97 | 489.942 | 3 | 36.99 | 1466.807 | -2.3 |
| 8046 | egTQeEKWKKZ | 97 | 489.9418 | 3 | 37.04 | 1466.807 | -2.6 |

| | | | | | | | |
|---|---|---|---|---|---|---|---|
| 8049 | egTQeEKWKKZ | 97 | 367.7082 | 4 | 37.05 | 1466.807 | -2.6 |
| 8087 | egTQeEKWKKZ | 97 | 489.9422 | 3 | 37.21 | 1466.807 | -1.8 |
| 8334 | DTALffKWKKZ | 97 | 478.9348 | 3 | 38.28 | 1433.786 | -2.2 |
| 8346 | DTALffKWKKZ | 97 | 478.9348 | 3 | 38.33 | 1433.786 | -2.3 |
| 8409 | DTALffKWKKZ | 97 | 359.4525 | 4 | 38.6 | 1433.786 | -3.4 |
| 8469 | DTALffKWKKZ | 97 | 478.9349 | 3 | 38.87 | 1433.786 | -2.1 |
| 8769 | fegQbFKWKKZ | 97 | 512.9662 | 3 | 40.17 | 1535.881 | -2.4 |
| 8802 | fegQbFKWKKZ | 97 | 512.9664 | 3 | 40.31 | 1535.881 | -2.1 |
| 8817 | fegQbFKWKKZ | 97 | 512.9662 | 3 | 40.38 | 1535.881 | -2.4 |
| 8841 | fegQbFKWKKZ | 97 | 512.9663 | 3 | 40.48 | 1535.881 | -2.3 |
| 8853 | fegQbFKWKKZ | 97 | 512.9662 | 3 | 40.54 | 1535.881 | -2.5 |
| 8865 | fegQbFKWKKZ | 97 | 512.9659 | 3 | 40.59 | 1535.881 | -3 |
| 8886 | fegQbFKWKKZ | 97 | 512.9662 | 3 | 40.68 | 1535.881 | -2.5 |
| 8898 | fegQbFKWKKZ | 97 | 512.9667 | 3 | 40.74 | 1535.881 | -1.5 |
| 8913 | fegQbFKWKKZ | 97 | 512.9667 | 3 | 40.8 | 1535.881 | -1.6 |
| 9333 | ePLEegKWKKZ | 97 | 362.9661 | 4 | 42.58 | 1447.838 | -1.8 |
| 5014 | PKLDFLKWKKZ | 96 | 434.6068 | 3 | 24.8 | 1300.802 | -2.5 |
| 5186 | gaFQRfKWKKZ | 96 | 368.2266 | 4 | 25.49 | 1468.882 | -3.1 |
| 5914 | ffKHgTKWKKZ | 96 | 382.7294 | 4 | 28.44 | 1526.891 | -1.9 |
| 6438 | SNfDfbKWKKZ | 96 | 492.9297 | 3 | 30.57 | 1475.772 | -2.9 |
| 6628 | TNFLGfKWKKZ | 96 | 336.6952 | 4 | 31.34 | 1342.755 | -2.4 |
| 5261 | gFgNTQKWKKZ | 96 | 333.9584 | 4 | 25.79 | 1331.808 | -2.5 |
| 5268 | gFgNTQKWKKZ | 96 | 444.942 | 3 | 25.81 | 1331.808 | -2.7 |
| 5281 | FggNTQKWKKZ | 96 | 444.9425 | 3 | 25.86 | 1331.808 | -1.6 |
| 5284 | gFgNTQKWKKZ | 96 | 333.9586 | 4 | 25.88 | 1331.808 | -1.9 |
| 5333 | fFKFNTKWKKZ | 96 | 362.9598 | 4 | 26.07 | 1447.813 | -1.7 |
| 5375 | fFKFNTKWKKZ | 96 | 362.9595 | 4 | 26.24 | 1447.813 | -2.6 |
| 5388 | fFKFNTKWKKZ | 96 | 362.9596 | 4 | 26.29 | 1447.813 | -2.5 |
| 5401 | fFKFNTKWKKZ | 96 | 362.9596 | 4 | 26.35 | 1447.813 | -2.4 |
| 5440 | fFKFNTKWKKZ | 96 | 362.9598 | 4 | 26.51 | 1447.813 | -1.9 |
| 5499 | fNTRfNKWKKZ | 96 | 380.7127 | 4 | 26.75 | 1518.825 | -1.9 |
| 5541 | fNTRfNKWKKZ | 96 | 380.7127 | 4 | 26.92 | 1518.825 | -2 |
| 5554 | fNTRfNKWKKZ | 96 | 380.7127 | 4 | 26.98 | 1518.825 | -2 |
| 5993 | DfdDfKKWKKZ | 96 | 523.6174 | 3 | 28.77 | 1567.834 | -2.2 |
| 6029 | DfdDfKKWKKZ | 96 | 523.6177 | 3 | 28.91 | 1567.834 | -1.7 |
| 6055 | DfdDfKKWKKZ | 96 | 523.6176 | 3 | 29.02 | 1567.834 | -2 |
| 6081 | DfdDfKKWKKZ | 96 | 523.6174 | 3 | 29.13 | 1567.834 | -2.2 |
| 6188 | FSfGgRKWKKZ | 96 | 462.6107 | 3 | 29.57 | 1384.813 | -2.2 |
| 6211 | FSfGgRKWKKZ | 96 | 347.2097 | 4 | 29.66 | 1384.813 | -2.4 |
| 6529 | fFKeSDKWKKZ | 96 | 372.2056 | 4 | 30.93 | 1484.797 | -2.5 |
| 6542 | fFKeSDKWKKZ | 96 | 372.2056 | 4 | 30.98 | 1484.797 | -2.3 |
| 6638 | TNFLGfKWKKZ | 96 | 448.5913 | 3 | 31.37 | 1342.755 | -2.1 |
| 6641 | TNFLGfKWKKZ | 96 | 336.6952 | 4 | 31.39 | 1342.755 | -2.4 |

| | | | | | | | |
|---|---|---|---|---|---|---|---|
| 6654 | TNFLGfKWKKZ | 96 | 336.6952 | 4 | 31.44 | 1342.755 | -2.5 |
| 6686 | TNFLGfKWKKZ | 96 | 336.6952 | 4 | 31.57 | 1342.755 | -2.3 |
| 7106 | gTEgfPKWKKZ | 96 | 464.9504 | 3 | 33.24 | 1391.833 | -2.6 |
| 7154 | GFPLEfKWKKZ | 96 | 452.2594 | 3 | 33.43 | 1353.76 | -2.6 |
| 7158 | gTEgfPKWKKZ | 96 | 464.9506 | 3 | 33.44 | 1391.833 | -2.1 |
| 7164 | GFPLEfKWKKZ | 96 | 452.2596 | 3 | 33.47 | 1353.76 | -2.2 |
| 7203 | GFPLEfKWKKZ | 96 | 452.2595 | 3 | 33.62 | 1353.76 | -2.2 |
| 7226 | gTEgfPKWKKZ | 96 | 464.9505 | 3 | 33.71 | 1391.833 | -2.5 |
| 7304 | GFPLEfKWKKZ | 96 | 452.2596 | 3 | 34.02 | 1353.76 | -2.2 |
| 7854 | egTQeEKWKKZ | 96 | 489.942 | 3 | 36.28 | 1466.807 | -2.2 |
| 7896 | egTQeEKWKKZ | 96 | 367.7082 | 4 | 36.45 | 1466.807 | -2.5 |
| 8010 | egTQeEKWKKZ | 96 | 367.7082 | 4 | 36.9 | 1466.807 | -2.4 |
| 8036 | egTQeEKWKKZ | 96 | 367.7082 | 4 | 37 | 1466.807 | -2.5 |
| 8059 | egTQeEKWKKZ | 96 | 489.942 | 3 | 37.1 | 1466.807 | -2.1 |
| 8072 | egTQeEKWKKZ | 96 | 489.942 | 3 | 37.15 | 1466.807 | -2.1 |
| 8424 | DTALffKWKKZ | 96 | 359.4531 | 4 | 38.67 | 1433.786 | -1.8 |
| 8589 | DTALffKWKKZ | 96 | 478.9347 | 3 | 39.38 | 1433.786 | -2.5 |
| 8694 | DTALffKWKKZ | 96 | 478.9349 | 3 | 39.82 | 1433.786 | -2.2 |
| 8874 | fegQbFKWKKZ | 96 | 512.9663 | 3 | 40.63 | 1535.881 | -2.3 |
| 9189 | ePLEegKWKKZ | 96 | 362.9659 | 4 | 41.97 | 1447.838 | -2.2 |
| 9213 | ePLEegKWKKZ | 96 | 362.966 | 4 | 42.07 | 1447.838 | -2.1 |
| 9225 | ePLEegKWKKZ | 96 | 362.9659 | 4 | 42.12 | 1447.838 | -2.2 |
| 9258 | ePLEegKWKKZ | 96 | 362.9659 | 4 | 42.26 | 1447.838 | -2.2 |
| 3470 | KFSLEaKWKKZ | 95 | 323.9509 | 4 | 18.33 | 1291.776 | -1.6 |
| 4126 | AKQfFNKWKKZ | 95 | 467.2703 | 3 | 21.15 | 1398.793 | -2.4 |
| 4386 | bgRdGfKWKKZ | 95 | 364.2281 | 4 | 22.25 | 1452.887 | -2.6 |
| 5644 | ffKHgTKWKKZ | 95 | 382.7293 | 4 | 27.35 | 1526.891 | -2.1 |
| 5941 | fGENDLKWKKZ | 95 | 447.2422 | 3 | 28.55 | 1338.709 | -2.7 |
| 6127 | FfQEhLKWKKZ | 95 | 471.9383 | 3 | 29.32 | 1412.797 | -2.6 |
| 6539 | eAFfdEKWKKZ | 95 | 511.2785 | 3 | 30.97 | 1530.817 | -2.4 |
| 8601 | DTALefKWKKZ | 95 | 470.2629 | 3 | 39.43 | 1407.77 | -2.4 |
| 4150 | AKQfFNKWKKZ | 95 | 467.2706 | 3 | 21.26 | 1398.793 | -1.7 |
| 4396 | bgRdGfKWKKZ | 95 | 364.2281 | 4 | 22.29 | 1452.887 | -2.6 |
| 4406 | bgRdGfKWKKZ | 95 | 364.2282 | 4 | 22.33 | 1452.887 | -2.3 |
| 4416 | bgRdGfKWKKZ | 95 | 364.2282 | 4 | 22.36 | 1452.887 | -2.1 |
| 4429 | bgRdGfKWKKZ | 95 | 364.2284 | 4 | 22.41 | 1452.887 | -1.6 |
| 4442 | bgRdGfKWKKZ | 95 | 364.2282 | 4 | 22.47 | 1452.887 | -2.3 |
| 4455 | bgRdGfKWKKZ | 95 | 364.2283 | 4 | 22.52 | 1452.887 | -2.1 |
| 4468 | bgRdGfKWKKZ | 95 | 364.2281 | 4 | 22.57 | 1452.887 | -2.4 |
| 4481 | bgRdGfKWKKZ | 95 | 364.2281 | 4 | 22.62 | 1452.887 | -2.6 |
| 4494 | bgRdGfKWKKZ | 95 | 364.2281 | 4 | 22.67 | 1452.887 | -2.5 |
| 4520 | bgRdGfKWKKZ | 95 | 364.2281 | 4 | 22.78 | 1452.887 | -2.5 |
| 4533 | bgRdGfKWKKZ | 95 | 364.2282 | 4 | 22.83 | 1452.887 | -2.1 |

| | | | | | | | |
|---|---|---|---|---|---|---|---|
| 4546 | bgRdGfKWKKZ | 95 | 364.2283 | 4 | 22.88 | 1452.887 | -2.1 |
| 4565 | bgRdGfKWKKZ | 95 | 364.2283 | 4 | 22.96 | 1452.887 | -2.1 |
| 4575 | bgRdGfKWKKZ | 95 | 364.2282 | 4 | 23.01 | 1452.887 | -2.2 |
| 4594 | bgRdGfKWKKZ | 95 | 364.2282 | 4 | 23.08 | 1452.887 | -2.3 |
| 5258 | gFgNTQKWKKZ | 95 | 444.9422 | 3 | 25.78 | 1331.808 | -2.3 |
| 5294 | gFgNTQKWKKZ | 95 | 444.9422 | 3 | 25.92 | 1331.808 | -2.4 |
| 5307 | gFgNTQKWKKZ | 95 | 444.9422 | 3 | 25.97 | 1331.808 | -2.4 |
| 5346 | fFKFNTKWKKZ | 95 | 362.9598 | 4 | 26.12 | 1447.813 | -1.9 |
| 5359 | fFKFNTKWKKZ | 95 | 362.9596 | 4 | 26.18 | 1447.813 | -2.3 |
| 5414 | fFKFNTKWKKZ | 95 | 362.9598 | 4 | 26.4 | 1447.813 | -1.8 |
| 5427 | fFKFNTKWKKZ | 95 | 362.96 | 4 | 26.45 | 1447.813 | -1.4 |
| 5486 | fNTRfNKWKKZ | 95 | 380.7127 | 4 | 26.7 | 1518.825 | -2 |
| 5502 | fNTRfNKWKKZ | 95 | 507.2813 | 3 | 26.76 | 1518.825 | -1.7 |
| 5512 | fNTRfNKWKKZ | 95 | 380.7126 | 4 | 26.8 | 1518.825 | -2.1 |
| 5978 | DfdDfKKWKKZ | 95 | 392.9649 | 4 | 28.7 | 1567.834 | -2.2 |
| 6000 | DfdDfKKWKKZ | 95 | 523.6172 | 3 | 28.8 | 1567.834 | -2.6 |
| 6001 | DfdDfKKWKKZ | 95 | 392.9647 | 4 | 28.8 | 1567.834 | -2.8 |
| 6011 | DfdDfKKWKKZ | 95 | 392.9652 | 4 | 28.84 | 1567.834 | -1.5 |
| 6016 | DfdDfKKWKKZ | 95 | 523.6177 | 3 | 28.86 | 1567.834 | -1.7 |
| 6037 | DfdDfKKWKKZ | 95 | 392.9648 | 4 | 28.95 | 1567.834 | -2.6 |
| 6042 | DfdDfKKWKKZ | 95 | 523.6174 | 3 | 28.97 | 1567.834 | -2.4 |
| 6063 | DfdDfKKWKKZ | 95 | 392.9647 | 4 | 29.06 | 1567.834 | -2.7 |
| 6068 | DfdDfKKWKKZ | 95 | 523.6173 | 3 | 29.07 | 1567.834 | -2.5 |
| 6090 | DfdDfKKWKKZ | 95 | 523.6179 | 3 | 29.17 | 1567.834 | -1.4 |
| 6124 | SaeDfQKWKKZ | 95 | 360.445 | 4 | 29.31 | 1437.756 | -3.3 |
| 6130 | SaeDfQKWKKZ | 95 | 480.258 | 3 | 29.34 | 1437.756 | -2.3 |
| 6140 | SaeDfQKWKKZ | 95 | 360.4452 | 4 | 29.38 | 1437.756 | -2.7 |
| 6153 | SaeDfQKWKKZ | 95 | 480.2581 | 3 | 29.43 | 1437.756 | -2.2 |
| 6375 | eTFLGEKWKKZ | 95 | 444.9195 | 3 | 30.32 | 1331.739 | -1.7 |
| 6518 | fFKeSDKWKKZ | 95 | 495.9386 | 3 | 30.89 | 1484.797 | -2 |
| 7096 | gTEgfPKWKKZ | 95 | 348.9647 | 4 | 33.2 | 1391.833 | -2.3 |
| 7145 | gTEgfPKWKKZ | 95 | 464.9505 | 3 | 33.39 | 1391.833 | -2.5 |
| 7187 | FGPLEfKWKKZ | 95 | 452.2595 | 3 | 33.56 | 1353.76 | -2.2 |
| 7229 | FGPLEfKWKKZ | 95 | 452.2595 | 3 | 33.72 | 1353.76 | -2.4 |
| 7252 | gTEgfPKWKKZ | 95 | 464.9501 | 3 | 33.81 | 1391.833 | -3.2 |
| 7281 | GFPLEfKWKKZ | 95 | 452.2596 | 3 | 33.93 | 1353.76 | -2.2 |
| 7346 | FGPLEfKWKKZ | 95 | 452.2596 | 3 | 34.19 | 1353.76 | -2.2 |
| 7572 | fGAfFQKWKKZ | 95 | 479.9315 | 3 | 35.14 | 1436.776 | -2.2 |
| 7587 | gbDeDfKWKKZ | 95 | 497.9421 | 3 | 35.2 | 1490.807 | -2 |
| 8023 | egTQeEKWKKZ | 95 | 367.7084 | 4 | 36.95 | 1466.807 | -2 |
| 8160 | egTQeEKWKKZ | 95 | 489.942 | 3 | 37.52 | 1466.807 | -2.1 |
| 8184 | egTQeEKWKKZ | 95 | 489.9419 | 3 | 37.62 | 1466.807 | -2.3 |
| 8244 | egTQeEKWKKZ | 95 | 489.9423 | 3 | 37.88 | 1466.807 | -1.6 |

| | | | | | | | |
|---|---|---|---|---|---|---|---|
| 8376 | DTALffKWKKZ | 95 | 359.4526 | 4 | 38.46 | 1433.786 | -3.1 |
| 8394 | DTALffKWKKZ | 95 | 478.9345 | 3 | 38.53 | 1433.786 | -2.9 |
| 8434 | DTALffKWKKZ | 95 | 359.4531 | 4 | 38.71 | 1433.786 | -1.7 |
| 8613 | DTALefKWKKZ | 95 | 470.2629 | 3 | 39.48 | 1407.77 | -2.5 |
| 8685 | DTALffKWKKZ | 95 | 478.9349 | 3 | 39.79 | 1433.786 | -2.2 |
| 8745 | DTALffKWKKZ | 95 | 478.9352 | 3 | 40.06 | 1433.786 | -1.6 |
| 8804 | fegQbFKWKKZ | 95 | 512.9664 | 3 | 40.32 | 1535.881 | -2.1 |
| 9270 | ePLEegKWKKZ | 95 | 362.9661 | 4 | 42.31 | 1447.838 | -1.7 |
| 9282 | ePLEegKWKKZ | 95 | 362.9661 | 4 | 42.36 | 1447.838 | -1.9 |
| 4659 | bgRdGfKWKKZ | 94 | 364.2283 | 4 | 23.35 | 1452.887 | -2 |
| 4965 | FKeNFKKWKKZ | 94 | 363.2178 | 4 | 24.6 | 1448.845 | -1.6 |
| 5027 | NFDSfdKWKKZ | 94 | 484.2582 | 3 | 24.85 | 1449.756 | -2 |
| 5595 | PSFfEAKWKKZ | 94 | 336.4374 | 4 | 27.15 | 1341.723 | -2.2 |
| 8292 | PLLEegKWKKZ | 94 | 455.6191 | 3 | 38.09 | 1363.838 | -1.8 |
| 8483 | hfFgSgKWKKZ | 94 | 346.9674 | 4 | 38.93 | 1383.843 | -2.1 |
| 9801 | GfLfGKWWKKZ | 94 | 483.2719 | 3 | 44.64 | 1446.796 | -1.8 |
| 3446 | KFSLEaKWKKZ | 94 | 323.9506 | 4 | 18.22 | 1291.776 | -2.3 |
| 3452 | KFSLEaKWKKZ | 94 | 323.9507 | 4 | 18.25 | 1291.776 | -1.9 |
| 3482 | KFSLEaKWKKZ | 94 | 323.9506 | 4 | 18.38 | 1291.776 | -2.5 |
| 4507 | bgRdGfKWKKZ | 94 | 364.2282 | 4 | 22.72 | 1452.887 | -2.2 |
| 4607 | bgRdGfKWKKZ | 94 | 364.2281 | 4 | 23.14 | 1452.887 | -2.6 |
| 4975 | PKLDFLKWKKZ | 94 | 326.207 | 4 | 24.64 | 1300.802 | -2.5 |
| 4982 | PKLDFLKWKKZ | 94 | 326.2068 | 4 | 24.67 | 1300.802 | -2.9 |
| 5008 | PKLDFLKWKKZ | 94 | 326.207 | 4 | 24.77 | 1300.802 | -2.4 |
| 5016 | PKLDFLKWKKZ | 94 | 434.6068 | 3 | 24.8 | 1300.802 | -2.5 |
| 5021 | PKLDFLKWKKZ | 94 | 326.2069 | 4 | 24.82 | 1300.802 | -2.6 |
| 5037 | PKLDFLKWKKZ | 94 | 326.2072 | 4 | 24.89 | 1300.802 | -1.9 |
| 5271 | gFgNTQKWKKZ | 94 | 333.9584 | 4 | 25.83 | 1331.808 | -2.4 |
| 5300 | gFgNTQKWKKZ | 94 | 333.9586 | 4 | 25.94 | 1331.808 | -2.1 |
| 5338 | fFKFNTKWKKZ | 94 | 483.6109 | 3 | 26.09 | 1447.813 | -1.4 |
| 5351 | fFKFNTKWKKZ | 94 | 483.6106 | 3 | 26.15 | 1447.813 | -2 |
| 5367 | fFKFNTKWKKZ | 94 | 483.6103 | 3 | 26.21 | 1447.813 | -2.6 |
| 5396 | fFKFNTKWKKZ | 94 | 483.6103 | 3 | 26.32 | 1447.813 | -2.6 |
| 5406 | fFKFNTKWKKZ | 94 | 483.6104 | 3 | 26.36 | 1447.813 | -2.4 |
| 5419 | fFKFNTKWKKZ | 94 | 483.6109 | 3 | 26.42 | 1447.813 | -1.4 |
| 5618 | PSFfEAKWKKZ | 94 | 336.4376 | 4 | 27.24 | 1341.723 | -1.6 |
| 5660 | ffKHgTKWKKZ | 94 | 382.7293 | 4 | 27.41 | 1526.891 | -2.1 |
| 5673 | ffKHgTKWKKZ | 94 | 382.7292 | 4 | 27.46 | 1526.891 | -2.4 |
| 5689 | ffKHgTKWKKZ | 94 | 382.7295 | 4 | 27.53 | 1526.891 | -1.7 |
| 5696 | ffKHgTKWKKZ | 94 | 382.7293 | 4 | 27.56 | 1526.891 | -2 |
| 5988 | DfdDfKKWKKZ | 94 | 392.9648 | 4 | 28.75 | 1567.834 | -2.5 |
| 6021 | DfdDfKKWKKZ | 94 | 392.9651 | 4 | 28.88 | 1567.834 | -1.6 |
| 6076 | DfdDfKKWKKZ | 94 | 392.9649 | 4 | 29.11 | 1567.834 | -2.2 |

| | | | | | | | |
|---|---|---|---|---|---|---|---|
| 6091 | DfdDfKKWKKZ | 94 | 392.9651 | 4 | 29.17 | 1567.834 | -1.8 |
| 6117 | SaeDfQKWKKZ | 94 | 480.258 | 3 | 29.28 | 1437.756 | -2.5 |
| 6145 | SaeDfQKWKKZ | 94 | 480.2581 | 3 | 29.4 | 1437.756 | -2.2 |
| 6513 | fFKeSDKWKKZ | 94 | 372.2057 | 4 | 30.87 | 1484.797 | -2 |
| 6534 | fFKeSDKWKKZ | 94 | 495.9384 | 3 | 30.95 | 1484.797 | -2.4 |
| 6560 | fFKeSDKWKKZ | 94 | 495.9389 | 3 | 31.05 | 1484.797 | -1.4 |
| 6565 | eAFfdEKWKKZ | 94 | 511.2789 | 3 | 31.08 | 1530.817 | -1.7 |
| 6581 | eAFfdEKWKKZ | 94 | 511.2785 | 3 | 31.14 | 1530.817 | -2.5 |
| 6589 | GfLHfGKWKKZ | 94 | 466.9314 | 3 | 31.18 | 1397.776 | -2.6 |
| 6677 | TNFLGfKWKKZ | 94 | 448.5912 | 3 | 31.53 | 1342.755 | -2.2 |
| 6722 | TNFLGfKWKKZ | 94 | 448.5912 | 3 | 31.71 | 1342.755 | -2.3 |
| 7050 | gTEgfPKWKKZ | 94 | 348.9648 | 4 | 33.02 | 1391.833 | -2.2 |
| 7057 | gTEgfPKWKKZ | 94 | 464.9505 | 3 | 33.05 | 1391.833 | -2.4 |
| 7076 | gTEgfPKWKKZ | 94 | 348.9647 | 4 | 33.12 | 1391.833 | -2.5 |
| 7167 | gTEgfPKWKKZ | 94 | 348.9647 | 4 | 33.48 | 1391.833 | -2.4 |
| 7197 | gTEgfPKWKKZ | 94 | 464.9505 | 3 | 33.6 | 1391.833 | -2.5 |
| 7255 | FGPLEfKWKKZ | 94 | 452.2591 | 3 | 33.83 | 1353.76 | -3.3 |
| 7271 | FGPLEfKWKKZ | 94 | 452.2593 | 3 | 33.89 | 1353.76 | -2.6 |
| 7330 | FGPLEfKWKKZ | 94 | 452.2595 | 3 | 34.13 | 1353.76 | -2.2 |
| 7359 | FGPLEfKWKKZ | 94 | 452.2594 | 3 | 34.25 | 1353.76 | -2.6 |
| 7636 | fGAfFQKWKKZ | 94 | 479.9315 | 3 | 35.4 | 1436.776 | -2 |
| 7779 | GfAfFQKWKKZ | 94 | 479.9315 | 3 | 35.97 | 1436.776 | -2.2 |
| 8304 | PLLEegKWKKZ | 94 | 455.619 | 3 | 38.15 | 1363.838 | -2 |
| 8310 | DTALffKWKKZ | 94 | 478.9347 | 3 | 38.17 | 1433.786 | -2.5 |
| 8511 | DTALffKWKKZ | 94 | 478.9348 | 3 | 39.04 | 1433.786 | -2.3 |
| 8523 | DTALffKWKKZ | 94 | 478.9349 | 3 | 39.09 | 1433.786 | -2.1 |
| 8571 | DTALffKWKKZ | 94 | 478.9347 | 3 | 39.3 | 1433.786 | -2.5 |
| 8625 | DTALefKWKKZ | 94 | 470.2631 | 3 | 39.53 | 1407.77 | -2.1 |
| 8771 | fegQbFKWKKZ | 94 | 512.9662 | 3 | 40.17 | 1535.881 | -2.4 |
| 8819 | fegQbFKWKKZ | 94 | 512.9662 | 3 | 40.39 | 1535.881 | -2.4 |
| 8889 | fegQbFKWKKZ | 94 | 384.9765 | 4 | 40.7 | 1535.881 | -2.4 |
| 8900 | fegQbFKWKKZ | 94 | 512.9667 | 3 | 40.74 | 1535.881 | -1.5 |
| 9174 | ePLEegKWKKZ | 94 | 362.9661 | 4 | 41.9 | 1447.838 | -1.9 |
| 9471 | ePLEegKWKKZ | 94 | 483.6192 | 3 | 43.18 | 1447.838 | -1.5 |
| 9807 | GfLfGKWWKKZ | 94 | 483.2719 | 3 | 44.66 | 1446.796 | -1.8 |
| 9819 | GfLfGKWWKKZ | 94 | 483.2717 | 3 | 44.72 | 1446.796 | -2.2 |
| 9843 | GfLfGKWWKKZ | 94 | 483.2719 | 3 | 44.82 | 1446.796 | -1.7 |
| 4147 | AgTLGNKWKKZ | 93 | 293.6877 | 4 | 21.24 | 1170.724 | -1.8 |
| 4246 | KfGQdLKWKKZ | 93 | 354.2176 | 4 | 21.67 | 1412.845 | -2.2 |
| 4445 | fhNRLTKWKKZ | 93 | 345.9611 | 4 | 22.48 | 1379.819 | -2.8 |
| 4766 | EAfaLGKWKKZ | 93 | 427.9195 | 3 | 23.8 | 1280.739 | -2 |
| 4793 | TRfQAFKWKKZ | 93 | 354.4572 | 4 | 23.91 | 1413.803 | -2.4 |
| 5134 | TKfhKfKWKKZ | 93 | 492.9661 | 3 | 25.28 | 1475.881 | -2.8 |

| | | | | | | | |
|---|---|---|---|---|---|---|---|
| 6735 | afNfgAKWKKZ | 93 | 362.4647 | 4 | 31.76 | 1445.834 | -2.6 |
| 8514 | EgTeQeKWKKZ | 93 | 489.9418 | 3 | 39.06 | 1466.807 | -2.5 |
| 9549 | ePLEegKWKKZ | 93 | 483.6192 | 3 | 43.52 | 1447.838 | -1.5 |
| 4135 | AKQfFNKWKKZ | 93 | 467.2702 | 3 | 21.19 | 1398.793 | -2.7 |
| 4156 | AgTLGNKWKKZ | 93 | 293.6875 | 4 | 21.28 | 1170.724 | -2.2 |
| 4192 | AgTLGNKWKKZ | 93 | 293.6874 | 4 | 21.43 | 1170.724 | -2.5 |
| 4195 | AgTLGNKWKKZ | 93 | 391.2474 | 3 | 21.45 | 1170.724 | -2.8 |
| 4497 | fhNRLTKWKKZ | 93 | 345.9611 | 4 | 22.69 | 1379.819 | -2.8 |
| 4781 | hDfaLGKWKKZ | 93 | 427.9194 | 3 | 23.86 | 1280.739 | -2.2 |
| 4978 | FKeNFKKWKKZ | 93 | 363.2176 | 4 | 24.66 | 1448.845 | -2.3 |
| 4995 | PKLDFLKWKKZ | 93 | 326.207 | 4 | 24.72 | 1300.802 | -2.4 |
| 5001 | NFDSfdKWKKZ | 93 | 363.4454 | 4 | 24.75 | 1449.756 | -2.2 |
| 5011 | NFDSfdKWKKZ | 93 | 484.2582 | 3 | 24.79 | 1449.756 | -2 |
| 5531 | fNTRfNKWKKZ | 93 | 507.2809 | 3 | 26.88 | 1518.825 | -2.5 |
| 5608 | PSFfEAKWKKZ | 93 | 336.4376 | 4 | 27.2 | 1341.723 | -1.7 |
| 6109 | SaeDfQKWKKZ | 93 | 360.4451 | 4 | 29.25 | 1437.756 | -2.9 |
| 6137 | FfQEhLKWKKZ | 93 | 471.9384 | 3 | 29.37 | 1412.797 | -2.5 |
| 6150 | FfQEhLKWKKZ | 93 | 471.9384 | 3 | 29.42 | 1412.797 | -2.6 |
| 6165 | FSfGgRKWKKZ | 93 | 462.6103 | 3 | 29.48 | 1384.813 | -3 |
| 6168 | FfQEhLKWKKZ | 93 | 471.938 | 3 | 29.49 | 1412.797 | -3.4 |
| 6175 | FSfGgRKWKKZ | 93 | 462.6104 | 3 | 29.52 | 1384.813 | -2.9 |
| 6227 | FSfGgRKWKKZ | 93 | 347.2098 | 4 | 29.73 | 1384.813 | -2.2 |
| 6440 | SNfDfbKWKKZ | 93 | 492.9297 | 3 | 30.58 | 1475.772 | -2.9 |
| 6536 | eAFfdEKWKKZ | 93 | 383.7108 | 4 | 30.96 | 1530.817 | -2.3 |
| 6607 | GfLHfGKWKKZ | 93 | 466.9313 | 3 | 31.25 | 1397.776 | -2.8 |
| 6625 | TNFLGfKWKKZ | 93 | 448.5913 | 3 | 31.32 | 1342.755 | -2.1 |
| 6651 | TNFLGfKWKKZ | 93 | 448.5912 | 3 | 31.43 | 1342.755 | -2.3 |
| 6664 | TNFLGfKWKKZ | 93 | 448.5914 | 3 | 31.48 | 1342.755 | -1.9 |
| 6690 | TNFLGfKWKKZ | 93 | 448.5912 | 3 | 31.58 | 1342.755 | -2.3 |
| 6706 | TNFLGfKWKKZ | 93 | 448.5911 | 3 | 31.65 | 1342.755 | -2.6 |
| 6773 | SEfTeGKWKKZ | 93 | 461.579 | 3 | 31.91 | 1381.718 | -2.1 |
| 6783 | SEfTeGKWKKZ | 93 | 461.579 | 3 | 31.95 | 1381.718 | -2.1 |
| 6796 | SEfTeGKWKKZ | 93 | 461.579 | 3 | 32 | 1381.718 | -2.2 |
| 6809 | SEfTeGKWKKZ | 93 | 461.579 | 3 | 32.05 | 1381.718 | -2.1 |
| 6822 | SEfTeGKWKKZ | 93 | 461.5791 | 3 | 32.1 | 1381.718 | -1.9 |
| 6835 | SEfTeGKWKKZ | 93 | 461.5791 | 3 | 32.15 | 1381.718 | -2 |
| 6848 | SEfTeGKWKKZ | 93 | 461.579 | 3 | 32.2 | 1381.718 | -2.3 |
| 6861 | SEfTeGKWKKZ | 93 | 461.579 | 3 | 32.26 | 1381.718 | -2.2 |
| 6880 | SEfTeGKWKKZ | 93 | 461.579 | 3 | 32.33 | 1381.718 | -2.3 |
| 6896 | SEfTeGKWKKZ | 93 | 461.5793 | 3 | 32.39 | 1381.718 | -1.6 |
| 7108 | gTEgfPKWKKZ | 93 | 464.9504 | 3 | 33.25 | 1391.833 | -2.6 |
| 7109 | gTEgfPKWKKZ | 93 | 348.9646 | 4 | 33.26 | 1391.833 | -2.8 |
| 7138 | gTEgfPKWKKZ | 93 | 348.9646 | 4 | 33.37 | 1391.833 | -2.6 |

| | | | | | | | |
|---|---|---|---|---|---|---|---|
| 7151 | gTEgfPKWKKZ | 93 | 348.9646 | 4 | 33.42 | 1391.833 | -2.8 |
| 7171 | gTEgfPKWKKZ | 93 | 464.9505 | 3 | 33.49 | 1391.833 | -2.4 |
| 7241 | gTEgfPKWKKZ | 93 | 464.9507 | 3 | 33.77 | 1391.833 | -2.1 |
| 7265 | gTEgfPKWKKZ | 93 | 464.9505 | 3 | 33.87 | 1391.833 | -2.5 |
| 7320 | FGPLEfKWKKZ | 93 | 452.2596 | 3 | 34.09 | 1353.76 | -2.1 |
| 7649 | fGAfFQKWKKZ | 93 | 479.9317 | 3 | 35.45 | 1436.776 | -1.7 |
| 7766 | fGAfFQKWKKZ | 93 | 479.9316 | 3 | 35.92 | 1436.776 | -1.8 |
| 7807 | egTQeEKWKKZ | 93 | 489.9422 | 3 | 36.09 | 1466.807 | -1.7 |
| 7980 | egTQeEKWKKZ | 93 | 489.9424 | 3 | 36.78 | 1466.807 | -1.3 |
| 8022 | egTQeEKWKKZ | 93 | 489.9423 | 3 | 36.94 | 1466.807 | -1.5 |
| 8301 | DTALffKWKKZ | 93 | 478.9351 | 3 | 38.13 | 1433.786 | -1.8 |
| 8358 | DTALffKWKKZ | 93 | 478.9349 | 3 | 38.38 | 1433.786 | -2.1 |
| 8370 | DTALffKWKKZ | 93 | 478.9346 | 3 | 38.43 | 1433.786 | -2.7 |
| 8382 | DTALffKWKKZ | 93 | 478.9346 | 3 | 38.48 | 1433.786 | -2.7 |
| 8428 | DTALffKWKKZ | 93 | 478.9348 | 3 | 38.68 | 1433.786 | -2.3 |
| 8450 | DTALffKWKKZ | 93 | 478.9348 | 3 | 38.78 | 1433.786 | -2.2 |
| 8457 | DTALffKWKKZ | 93 | 478.9348 | 3 | 38.81 | 1433.786 | -2.3 |
| 8480 | DTALffKWKKZ | 93 | 478.9348 | 3 | 38.91 | 1433.786 | -2.3 |
| 8499 | DTALffKWKKZ | 93 | 478.9349 | 3 | 38.99 | 1433.786 | -2.2 |
| 8535 | DTALffKWKKZ | 93 | 478.9348 | 3 | 39.14 | 1433.786 | -2.3 |
| 8547 | DTALffKWKKZ | 93 | 478.935 | 3 | 39.19 | 1433.786 | -1.9 |
| 8559 | DTALffKWKKZ | 93 | 478.9347 | 3 | 39.25 | 1433.786 | -2.5 |
| 8759 | fegQbFKWKKZ | 93 | 512.9662 | 3 | 40.12 | 1535.881 | -2.4 |
| 8793 | fegQbFKWKKZ | 93 | 384.9766 | 4 | 40.27 | 1535.881 | -2.3 |
| 8843 | fegQbFKWKKZ | 93 | 512.9663 | 3 | 40.49 | 1535.881 | -2.3 |
| 8855 | fegQbFKWKKZ | 93 | 512.9662 | 3 | 40.54 | 1535.881 | -2.5 |
| 8867 | fegQbFKWKKZ | 93 | 512.9659 | 3 | 40.6 | 1535.881 | -3 |
| 8888 | fegQbFKWKKZ | 93 | 512.9662 | 3 | 40.69 | 1535.881 | -2.5 |
| 8901 | fegQbFKWKKZ | 93 | 384.9769 | 4 | 40.75 | 1535.881 | -1.5 |
| 8915 | fegQbFKWKKZ | 93 | 512.9667 | 3 | 40.81 | 1535.881 | -1.6 |
| 9150 | ePLEegKWKKZ | 93 | 483.6188 | 3 | 41.8 | 1447.838 | -2.3 |
| 9159 | ePLEegKWKKZ | 93 | 483.6191 | 3 | 41.83 | 1447.838 | -1.6 |
| 9171 | ePLEegKWKKZ | 93 | 483.6188 | 3 | 41.88 | 1447.838 | -2.2 |
| 9183 | ePLEegKWKKZ | 93 | 483.6188 | 3 | 41.94 | 1447.838 | -2.3 |
| 9207 | ePLEegKWKKZ | 93 | 483.6188 | 3 | 42.04 | 1447.838 | -2.3 |
| 9219 | ePLEegKWKKZ | 93 | 483.6188 | 3 | 42.09 | 1447.838 | -2.3 |
| 9231 | ePLEegKWKKZ | 93 | 483.619 | 3 | 42.14 | 1447.838 | -1.8 |
| 9234 | ePLEegKWKKZ | 93 | 362.966 | 4 | 42.16 | 1447.838 | -2 |
| 9243 | ePLEegKWKKZ | 93 | 483.6187 | 3 | 42.19 | 1447.838 | -2.5 |
| 9255 | ePLEegKWKKZ | 93 | 483.6188 | 3 | 42.24 | 1447.838 | -2.3 |
| 9279 | ePLEegKWKKZ | 93 | 483.6191 | 3 | 42.35 | 1447.838 | -1.6 |
| 9330 | ePLEegKWKKZ | 93 | 483.6191 | 3 | 42.57 | 1447.838 | -1.6 |
| 9363 | ePLEegKWKKZ | 93 | 483.6191 | 3 | 42.71 | 1447.838 | -1.7 |

| | | | | | | | |
|---|---|---|---|---|---|---|---|
| 9375 | ePLEegKWKKZ | 93 | 483.619 | 3 | 42.76 | 1447.838 | -1.8 |
| 9387 | ePLEegKWKKZ | 93 | 483.6191 | 3 | 42.81 | 1447.838 | -1.7 |
| 9402 | ePLEegKWKKZ | 93 | 483.6191 | 3 | 42.88 | 1447.838 | -1.6 |
| 9411 | ePLEegKWKKZ | 93 | 483.619 | 3 | 42.92 | 1447.838 | -1.8 |
| 9423 | ePLEegKWKKZ | 93 | 483.6187 | 3 | 42.97 | 1447.838 | -2.5 |
| 9435 | ePLEegKWKKZ | 93 | 483.6191 | 3 | 43.02 | 1447.838 | -1.7 |
| 9450 | ePLEegKWKKZ | 93 | 483.619 | 3 | 43.09 | 1447.838 | -1.8 |
| 9459 | ePLEegKWKKZ | 93 | 483.6194 | 3 | 43.12 | 1447.838 | -1 |
| 9483 | ePLEegKWKKZ | 93 | 483.6192 | 3 | 43.23 | 1447.838 | -1.4 |
| 9501 | ePLEegKWKKZ | 93 | 483.6192 | 3 | 43.31 | 1447.838 | -1.5 |
| 9513 | ePLEegKWKKZ | 93 | 483.6193 | 3 | 43.36 | 1447.838 | -1.3 |
| 9522 | ePLEegKWKKZ | 93 | 483.6192 | 3 | 43.4 | 1447.838 | -1.5 |
| 9534 | ePLEegKWKKZ | 93 | 483.6192 | 3 | 43.46 | 1447.838 | -1.5 |
| 9561 | ePLEegKWKKZ | 93 | 483.6192 | 3 | 43.57 | 1447.838 | -1.4 |
| 9573 | ePLEegKWKKZ | 93 | 483.6191 | 3 | 43.63 | 1447.838 | -1.6 |
| 9858 | GfLfGKWWKKZ | 93 | 483.272 | 3 | 44.89 | 1446.796 | -1.6 |
| 5193 | gKfGTPKWKKZ | 92 | 441.2754 | 3 | 25.52 | 1320.807 | -2.2 |
| 5430 | QeLbdgKWKKZ | 92 | 364.7295 | 4 | 26.47 | 1454.891 | -1.7 |
| 8583 | NGfTeGKWKKZ | 92 | 446.5716 | 3 | 39.35 | 1336.708 | -11.3 |
| 4168 | AgTLGNKWKKZ | 92 | 293.6875 | 4 | 21.33 | 1170.724 | -2.3 |
| 4171 | AgTLGNKWKKZ | 92 | 391.2475 | 3 | 21.35 | 1170.724 | -2.5 |
| 4180 | AgTLGNKWKKZ | 92 | 293.6876 | 4 | 21.38 | 1170.724 | -2 |
| 4751 | hDfaLGKWKKZ | 92 | 427.9192 | 3 | 23.73 | 1280.739 | -2.7 |
| 4796 | hDfaLGKWKKZ | 92 | 427.9194 | 3 | 23.92 | 1280.739 | -2.2 |
| 5283 | gFgNTQKWKKZ | 92 | 444.9425 | 3 | 25.87 | 1331.808 | -1.6 |
| 5518 | fNTRfNKWKKZ | 92 | 507.2813 | 3 | 26.83 | 1518.825 | -1.7 |
| 5611 | PSFfEAKWKKZ | 92 | 448.2477 | 3 | 27.21 | 1341.723 | -1.6 |
| 5621 | PSFfEAKWKKZ | 92 | 448.2477 | 3 | 27.25 | 1341.723 | -1.5 |
| 5646 | ffKHgTKWKKZ | 92 | 382.7293 | 4 | 27.35 | 1526.891 | -2.1 |
| 5709 | ffKHgTKWKKZ | 92 | 382.7293 | 4 | 27.61 | 1526.891 | -2.2 |
| 6050 | DfdDfKKWKKZ | 92 | 392.9648 | 4 | 29 | 1567.834 | -2.5 |
| 6105 | SaeDfQKWKKZ | 92 | 480.258 | 3 | 29.23 | 1437.756 | -2.4 |
| 6178 | FfQEhLKWKKZ | 92 | 354.2057 | 4 | 29.53 | 1412.797 | -2.4 |
| 6519 | eAFfdEKWKKZ | 92 | 383.7109 | 4 | 30.89 | 1530.817 | -1.9 |
| 6523 | eAFfdEKWKKZ | 92 | 383.7107 | 4 | 30.91 | 1530.817 | -2.4 |
| 6526 | eAFfdEKWKKZ | 92 | 511.2785 | 3 | 30.92 | 1530.817 | -2.4 |
| 6549 | eAFfdEKWKKZ | 92 | 383.711 | 4 | 31.01 | 1530.817 | -1.5 |
| 6575 | eAFfdEKWKKZ | 92 | 383.7107 | 4 | 31.11 | 1530.817 | -2.4 |
| 6631 | afNfgAKWKKZ | 92 | 362.4647 | 4 | 31.35 | 1445.834 | -2.7 |
| 6640 | TNFLGfKWKKZ | 92 | 448.5913 | 3 | 31.38 | 1342.755 | -2.1 |
| 6679 | TNFLGfKWKKZ | 92 | 448.5912 | 3 | 31.54 | 1342.755 | -2.2 |
| 6724 | TNFLGfKWKKZ | 92 | 448.5912 | 3 | 31.72 | 1342.755 | -2.3 |
| 6891 | afNfgAKWKKZ | 92 | 362.4649 | 4 | 32.37 | 1445.834 | -2.1 |

| | | | | | | | |
|---|---|---|---|---|---|---|---|
| 7069 | gTEgfPKWKKZ | 92 | 464.9505 | 3 | 33.09 | 1391.833 | -2.5 |
| 7095 | gTEgfPKWKKZ | 92 | 464.9506 | 3 | 33.2 | 1391.833 | -2.3 |
| 7121 | gTEgfPKWKKZ | 92 | 464.9504 | 3 | 33.3 | 1391.833 | -2.6 |
| 7147 | gTEgfPKWKKZ | 92 | 464.9505 | 3 | 33.4 | 1391.833 | -2.5 |
| 7160 | gTEgfPKWKKZ | 92 | 464.9506 | 3 | 33.45 | 1391.833 | -2.1 |
| 7177 | GFPLEfKWKKZ | 92 | 452.2594 | 3 | 33.52 | 1353.76 | -2.4 |
| 7212 | gTEgfPKWKKZ | 92 | 464.9505 | 3 | 33.66 | 1391.833 | -2.4 |
| 7218 | GFPLEfKWKKZ | 92 | 452.2594 | 3 | 33.68 | 1353.76 | -2.4 |
| 7228 | gTEgfPKWKKZ | 92 | 464.9505 | 3 | 33.72 | 1391.833 | -2.5 |
| 7555 | gbDeDfKWKKZ | 92 | 373.7082 | 4 | 35.07 | 1490.807 | -2.4 |
| 7568 | gbDeDfKWKKZ | 92 | 373.7083 | 4 | 35.12 | 1490.807 | -2.2 |
| 7817 | egTQeEKWKKZ | 92 | 489.9422 | 3 | 36.13 | 1466.807 | -1.8 |
| 7830 | egTQeEKWKKZ | 92 | 489.9422 | 3 | 36.18 | 1466.807 | -1.8 |
| 7866 | egTQeEKWKKZ | 92 | 489.9418 | 3 | 36.32 | 1466.807 | -2.6 |
| 7905 | egTQeEKWKKZ | 92 | 489.9421 | 3 | 36.48 | 1466.807 | -2 |
| 7918 | egTQeEKWKKZ | 92 | 489.942 | 3 | 36.53 | 1466.807 | -2.2 |
| 7944 | egTQeEKWKKZ | 92 | 489.942 | 3 | 36.63 | 1466.807 | -2.1 |
| 7954 | egTQeEKWKKZ | 92 | 489.9421 | 3 | 36.67 | 1466.807 | -1.9 |
| 7967 | egTQeEKWKKZ | 92 | 489.9421 | 3 | 36.73 | 1466.807 | -2 |
| 7993 | egTQeEKWKKZ | 92 | 489.9423 | 3 | 36.83 | 1466.807 | -1.5 |
| 8006 | egTQeEKWKKZ | 92 | 489.942 | 3 | 36.88 | 1466.807 | -2.1 |
| 8035 | egTQeEKWKKZ | 92 | 489.942 | 3 | 37 | 1466.807 | -2.3 |
| 8089 | egTQeEKWKKZ | 92 | 489.9422 | 3 | 37.22 | 1466.807 | -1.8 |
| 8406 | DTALffKWKKZ | 92 | 478.9348 | 3 | 38.58 | 1433.786 | -2.3 |
| 8418 | DTALffKWKKZ | 92 | 478.9348 | 3 | 38.64 | 1433.786 | -2.2 |
| 8440 | DTALffKWKKZ | 92 | 478.9348 | 3 | 38.73 | 1433.786 | -2.2 |
| 8487 | DTALffKWKKZ | 92 | 478.9348 | 3 | 38.94 | 1433.786 | -2.3 |
| 8619 | GNfTeGKWKKZ | 92 | 446.5721 | 3 | 39.5 | 1336.708 | -10 |
| 8634 | NGfTeGKWKKZ | 92 | 446.5716 | 3 | 39.57 | 1336.708 | -11.3 |
| 8646 | NGfTeGKWKKZ | 92 | 446.5716 | 3 | 39.61 | 1336.708 | -11.3 |
| 8687 | DTALffKWKKZ | 92 | 478.9349 | 3 | 39.79 | 1433.786 | -2.2 |
| 9195 | ePLEegKWKKZ | 92 | 483.619 | 3 | 41.99 | 1447.838 | -1.9 |
| 9201 | ePLEegKWKKZ | 92 | 362.9661 | 4 | 42.02 | 1447.838 | -1.9 |
| 9267 | ePLEegKWKKZ | 92 | 483.6192 | 3 | 42.29 | 1447.838 | -1.5 |
| 9291 | ePLEegKWKKZ | 92 | 483.6191 | 3 | 42.4 | 1447.838 | -1.6 |
| 9303 | ePLEegKWKKZ | 92 | 483.619 | 3 | 42.45 | 1447.838 | -2 |
| 9315 | ePLEegKWKKZ | 92 | 483.6192 | 3 | 42.5 | 1447.838 | -1.5 |
| 9339 | ePLEegKWKKZ | 92 | 483.6191 | 3 | 42.6 | 1447.838 | -1.7 |
| 9351 | ePLEegKWKKZ | 92 | 483.619 | 3 | 42.66 | 1447.838 | -1.8 |
| 9425 | ePLEegKWKKZ | 92 | 483.6187 | 3 | 42.98 | 1447.838 | -2.5 |
| 9524 | ePLEegKWKKZ | 92 | 483.6192 | 3 | 43.41 | 1447.838 | -1.5 |
| 9609 | ePLEegKWKKZ | 92 | 483.6188 | 3 | 43.79 | 1447.838 | -2.2 |
| 9645 | ePLEegKWKKZ | 92 | 483.6187 | 3 | 43.95 | 1447.838 | -2.5 |

| | | | | | | | |
|---|---|---|---|---|---|---|---|
| 9803 | GfLfGKWWKKZ | 92 | 483.2719 | 3 | 44.65 | 1446.796 | -1.8 |
| 9809 | GfLfGKWWKKZ | 92 | 483.2719 | 3 | 44.67 | 1446.796 | -1.8 |
| 9821 | GfLfGKWWKKZ | 92 | 483.2717 | 3 | 44.73 | 1446.796 | -2.2 |
| 9845 | GfLfGKWWKKZ | 92 | 483.2719 | 3 | 44.83 | 1446.796 | -1.7 |
| 3139 | SFEDGGKWKKZ | 91 | 394.2076 | 3 | 16.87 | 1179.604 | -2.2 |
| 5290 | FFTFThKWKKZ | 91 | 439.5879 | 3 | 25.9 | 1315.744 | -1.7 |
| 7128 | PfgghPKWKKZ | 91 | 336.9684 | 4 | 33.33 | 1343.848 | -2.8 |
| 8950 | NhEKLeKWKKZ | 91 | 452.2706 | 3 | 40.96 | 1353.792 | -1.5 |
| 9474 | FffbFfKWKKZ | 91 | 420.2331 | 4 | 43.19 | 1676.906 | -1.6 |
| 3157 | SFEDGGKWKKZ | 91 | 394.2078 | 3 | 16.95 | 1179.604 | -1.7 |
| 4183 | AgTLGNKWKKZ | 91 | 391.2477 | 3 | 21.4 | 1170.724 | -2 |
| 4438 | fhNRLTKWKKZ | 91 | 345.9613 | 4 | 22.45 | 1379.819 | -2.1 |
| 4458 | fhNRLTKWKKZ | 91 | 345.9612 | 4 | 22.53 | 1379.819 | -2.3 |
| 4471 | fhNRLTKWKKZ | 91 | 345.9612 | 4 | 22.58 | 1379.819 | -2.6 |
| 4484 | fhNRLTKWKKZ | 91 | 345.9611 | 4 | 22.63 | 1379.819 | -2.7 |
| 5089 | TKfhKfKWKKZ | 91 | 492.9663 | 3 | 25.1 | 1475.881 | -2.3 |
| 5105 | TKfhKfKWKKZ | 91 | 492.9666 | 3 | 25.17 | 1475.881 | -1.8 |
| 5222 | KgfGTPKWKKZ | 91 | 441.2751 | 3 | 25.63 | 1320.807 | -2.8 |
| 5235 | gKfGTPKWKKZ | 91 | 441.2754 | 3 | 25.68 | 1320.807 | -2 |
| 5253 | gFgNTQKWKKZ | 91 | 444.942 | 3 | 25.75 | 1331.808 | -2.9 |
| 5260 | gFgNTQKWKKZ | 91 | 444.9422 | 3 | 25.78 | 1331.808 | -2.3 |
| 5270 | gFgNTQKWKKZ | 91 | 444.942 | 3 | 25.82 | 1331.808 | -2.7 |
| 5313 | FFTFThKWKKZ | 91 | 439.5876 | 3 | 25.99 | 1315.744 | -2.3 |
| 5504 | fNTRfNKWKKZ | 91 | 507.2813 | 3 | 26.77 | 1518.825 | -1.7 |
| 5538 | NLhLgQKWKKZ | 91 | 423.6108 | 3 | 26.91 | 1267.813 | -1.9 |
| 5547 | fNTRfNKWKKZ | 91 | 507.2814 | 3 | 26.95 | 1518.825 | -1.6 |
| 5551 | NLhLgQKWKKZ | 91 | 423.6108 | 3 | 26.96 | 1267.813 | -2 |
| 5577 | NLhLgQKWKKZ | 91 | 423.6109 | 3 | 27.07 | 1267.813 | -1.6 |
| 6545 | fFKeSDKWKKZ | 91 | 495.9384 | 3 | 31 | 1484.797 | -2.4 |
| 6555 | fFKeSDKWKKZ | 91 | 372.2065 | 4 | 31.04 | 1484.797 | 0 |
| 6562 | eAFfdEKWKKZ | 91 | 383.711 | 4 | 31.06 | 1530.817 | -1.7 |
| 6567 | eAFfdEKWKKZ | 91 | 511.2789 | 3 | 31.08 | 1530.817 | -1.7 |
| 6583 | eAFfdEKWKKZ | 91 | 511.2785 | 3 | 31.15 | 1530.817 | -2.5 |
| 6627 | TNFLGfKWKKZ | 91 | 448.5913 | 3 | 31.33 | 1342.755 | -2.1 |
| 6644 | afNfgAKWKKZ | 91 | 362.4648 | 4 | 31.4 | 1445.834 | -2.4 |
| 6666 | TNFLGfKWKKZ | 91 | 448.5914 | 3 | 31.49 | 1342.755 | -1.9 |
| 6692 | TNFLGfKWKKZ | 91 | 448.5912 | 3 | 31.59 | 1342.755 | -2.3 |
| 6839 | afNfgAKWKKZ | 91 | 362.4648 | 4 | 32.17 | 1445.834 | -2.2 |
| 7059 | gTEgfPKWKKZ | 91 | 464.9505 | 3 | 33.05 | 1391.833 | -2.4 |
| 7082 | gTEgfPKWKKZ | 91 | 464.9506 | 3 | 33.15 | 1391.833 | -2.2 |
| 7134 | gTEgfPKWKKZ | 91 | 464.9506 | 3 | 33.35 | 1391.833 | -2.3 |
| 7156 | GFPLEfKWKKZ | 91 | 452.2594 | 3 | 33.44 | 1353.76 | -2.6 |
| 7189 | GFPLEfKWKKZ | 91 | 452.2595 | 3 | 33.57 | 1353.76 | -2.2 |

| | | | | | | | |
|---|---|---|---|---|---|---|---|
| 7199 | gTEgfPKWKKZ | 91 | 464.9505 | 3 | 33.61 | 1391.833 | -2.5 |
| 7231 | GFPLEfKWKKZ | 91 | 452.2595 | 3 | 33.73 | 1353.76 | -2.4 |
| 7242 | FGPLEfKWKKZ | 91 | 452.2596 | 3 | 33.78 | 1353.76 | -2.2 |
| 7294 | GFPLEfKWKKZ | 91 | 452.2595 | 3 | 33.98 | 1353.76 | -2.2 |
| 7323 | gTEgPfKWKKZ | 91 | 464.9506 | 3 | 34.1 | 1391.833 | -2.1 |
| 7348 | GFPLEfKWKKZ | 91 | 452.2596 | 3 | 34.2 | 1353.76 | -2.2 |
| 7361 | GFPLEfKWKKZ | 91 | 452.2594 | 3 | 34.25 | 1353.76 | -2.6 |
| 7545 | gbDeDfKWKKZ | 91 | 373.7083 | 4 | 35.03 | 1490.807 | -2.2 |
| 7584 | gbDeDfKWKKZ | 91 | 373.7081 | 4 | 35.18 | 1490.807 | -2.7 |
| 7597 | GfAfFQKWKKZ | 91 | 479.9317 | 3 | 35.24 | 1436.776 | -1.6 |
| 7600 | gbDeDfKWKKZ | 91 | 373.7082 | 4 | 35.25 | 1490.807 | -2.4 |
| 7843 | egTQeEKWKKZ | 91 | 489.9419 | 3 | 36.23 | 1466.807 | -2.5 |
| 7856 | egTQeEKWKKZ | 91 | 489.942 | 3 | 36.28 | 1466.807 | -2.2 |
| 7882 | egTQeEKWKKZ | 91 | 489.942 | 3 | 36.39 | 1466.807 | -2.3 |
| 7895 | egTQeEKWKKZ | 91 | 489.942 | 3 | 36.44 | 1466.807 | -2.2 |
| 7928 | egTQeEKWKKZ | 91 | 489.9422 | 3 | 36.57 | 1466.807 | -1.8 |
| 8048 | egTQeEKWKKZ | 91 | 489.9418 | 3 | 37.05 | 1466.807 | -2.6 |
| 8061 | egTQeEKWKKZ | 91 | 489.942 | 3 | 37.1 | 1466.807 | -2.1 |
| 8324 | DTALffKWKKZ | 91 | 478.9348 | 3 | 38.23 | 1433.786 | -2.3 |
| 8336 | DTALffKWKKZ | 91 | 478.9348 | 3 | 38.29 | 1433.786 | -2.2 |
| 8501 | DTALffKWKKZ | 91 | 478.9349 | 3 | 39 | 1433.786 | -2.2 |
| 8513 | DTALffKWKKZ | 91 | 478.9348 | 3 | 39.05 | 1433.786 | -2.3 |
| 8525 | DTALffKWKKZ | 91 | 478.9349 | 3 | 39.1 | 1433.786 | -2.1 |
| 8573 | DTALffKWKKZ | 91 | 478.9347 | 3 | 39.31 | 1433.786 | -2.5 |
| 8963 | NhEKLeKWKKZ | 91 | 452.2701 | 3 | 41.01 | 1353.792 | -2.6 |
| 8989 | NhEKLeKWKKZ | 91 | 452.2705 | 3 | 41.12 | 1353.792 | -1.6 |
| 9002 | NhEKLeKWKKZ | 91 | 452.2705 | 3 | 41.17 | 1353.792 | -1.7 |
| 9015 | NhEKLeKWKKZ | 91 | 452.2701 | 3 | 41.22 | 1353.792 | -2.6 |
| 9028 | NhEKLeKWKKZ | 91 | 452.2699 | 3 | 41.28 | 1353.792 | -3 |
| 9041 | NhEKLeKWKKZ | 91 | 452.2702 | 3 | 41.33 | 1353.792 | -2.4 |
| 9054 | NhEKLeKWKKZ | 91 | 452.2701 | 3 | 41.39 | 1353.792 | -2.6 |
| 9066 | NhEKLeKWKKZ | 91 | 452.2701 | 3 | 41.44 | 1353.792 | -2.6 |
| 9079 | NhEKLeKWKKZ | 91 | 452.2704 | 3 | 41.49 | 1353.792 | -2 |
| 9091 | NhEKLeKWKKZ | 91 | 452.2703 | 3 | 41.54 | 1353.792 | -2.1 |
| 9117 | NhEKLeKWKKZ | 91 | 452.2708 | 3 | 41.65 | 1353.792 | -1.1 |
| 9185 | ePLEegKWKKZ | 91 | 483.6188 | 3 | 41.95 | 1447.838 | -2.3 |
| 9221 | ePLEegKWKKZ | 91 | 483.6188 | 3 | 42.1 | 1447.838 | -2.3 |
| 9233 | ePLEegKWKKZ | 91 | 483.619 | 3 | 42.15 | 1447.838 | -1.8 |
| 9245 | ePLEegKWKKZ | 91 | 483.6187 | 3 | 42.2 | 1447.838 | -2.5 |
| 9257 | ePLEegKWKKZ | 91 | 483.6188 | 3 | 42.25 | 1447.838 | -2.3 |
| 9281 | ePLEegKWKKZ | 91 | 483.6191 | 3 | 42.36 | 1447.838 | -1.6 |
| 9377 | ePLEegKWKKZ | 91 | 483.619 | 3 | 42.77 | 1447.838 | -1.8 |
| 9389 | ePLEegKWKKZ | 91 | 483.6191 | 3 | 42.82 | 1447.838 | -1.7 |

| | | | | | | | |
|---|---|---|---|---|---|---|---|
| 9437 | ePLEegKWKKZ | 91 | 483.6191 | 3 | 43.03 | 1447.838 | -1.7 |
| 9452 | ePLEegKWKKZ | 91 | 483.619 | 3 | 43.1 | 1447.838 | -1.8 |
| 9461 | ePLEegKWKKZ | 91 | 483.6194 | 3 | 43.13 | 1447.838 | -1 |
| 9473 | ePLEegKWKKZ | 91 | 483.6192 | 3 | 43.19 | 1447.838 | -1.5 |
| 9485 | ePLEegKWKKZ | 91 | 483.6192 | 3 | 43.24 | 1447.838 | -1.4 |
| 9503 | ePLEegKWKKZ | 91 | 483.6192 | 3 | 43.32 | 1447.838 | -1.5 |
| 9515 | ePLEegKWKKZ | 91 | 483.6193 | 3 | 43.37 | 1447.838 | -1.3 |
| 9551 | ePLEegKWKKZ | 91 | 483.6192 | 3 | 43.53 | 1447.838 | -1.5 |
| 9563 | ePLEegKWKKZ | 91 | 483.6192 | 3 | 43.58 | 1447.838 | -1.4 |
| 9657 | ePLEegKWKKZ | 91 | 483.6191 | 3 | 44 | 1447.838 | -1.6 |
| 9860 | GfLfGKWWKKZ | 91 | 483.272 | 3 | 44.9 | 1446.796 | -1.6 |
| 3931 | TaDLKeKWKKZ | 90 | 336.4546 | 4 | 20.31 | 1341.792 | -2.1 |
| 6764 | EFeNedKWKKZ | 90 | 516.9419 | 3 | 31.87 | 1547.808 | -2.4 |
| 7024 | afNfgAKWKKZ | 90 | 362.4648 | 4 | 32.91 | 1445.834 | -2.5 |
| 7753 | APPeeNKWKKZ | 90 | 454.5879 | 3 | 35.87 | 1360.744 | -1.9 |
| 3431 | KFSLEaKWKKZ | 90 | 323.9508 | 4 | 18.16 | 1291.776 | -1.6 |
| 4529 | fhNRLTKWKKZ | 90 | 345.9611 | 4 | 22.82 | 1379.819 | -2.8 |
| 5013 | NFDSfdKWKKZ | 90 | 484.2582 | 3 | 24.79 | 1449.756 | -2 |
| 5023 | PKLDFLKWKKZ | 90 | 326.2069 | 4 | 24.83 | 1300.802 | -2.6 |
| 5029 | NFDSfdKWKKZ | 90 | 484.2582 | 3 | 24.86 | 1449.756 | -2 |
| 5039 | PKLDFLKWKKZ | 90 | 326.2072 | 4 | 24.9 | 1300.802 | -1.9 |
| 5121 | KTfhKfKWKKZ | 90 | 492.9664 | 3 | 25.23 | 1475.881 | -2.2 |
| 5274 | FFTFThKWKKZ | 90 | 439.5876 | 3 | 25.84 | 1315.744 | -2.3 |
| 5296 | gFgNTQKWKKZ | 90 | 444.9422 | 3 | 25.93 | 1331.808 | -2.4 |
| 5326 | FFTFThKWKKZ | 90 | 439.5877 | 3 | 26.05 | 1315.744 | -2.2 |
| 5484 | fNTRfNKWKKZ | 90 | 380.7128 | 4 | 26.69 | 1518.825 | -1.8 |
| 5525 | NLhLgQKWKKZ | 90 | 423.6104 | 3 | 26.85 | 1267.813 | -2.8 |
| 5549 | fNTRfNKWKKZ | 90 | 507.2814 | 3 | 26.95 | 1518.825 | -1.6 |
| 5564 | NLhLgQKWKKZ | 90 | 423.6106 | 3 | 27.02 | 1267.813 | -2.3 |
| 5589 | NLhLgQKWKKZ | 90 | 423.6105 | 3 | 27.12 | 1267.813 | -2.5 |
| 5675 | ffKHgTKWKKZ | 90 | 382.7292 | 4 | 27.47 | 1526.891 | -2.4 |
| 5980 | DfdDfKKWKKZ | 90 | 392.9649 | 4 | 28.71 | 1567.834 | -2.2 |
| 6132 | SaeDfQKWKKZ | 90 | 480.258 | 3 | 29.34 | 1437.756 | -2.3 |
| 6155 | SaeDfQKWKKZ | 90 | 480.2581 | 3 | 29.44 | 1437.756 | -2.2 |
| 6541 | eAFfdEKWKKZ | 90 | 511.2785 | 3 | 30.98 | 1530.817 | -2.4 |
| 6653 | TNFLGfKWKKZ | 90 | 448.5912 | 3 | 31.44 | 1342.755 | -2.3 |
| 6680 | afNfgAKWKKZ | 90 | 362.4648 | 4 | 31.54 | 1445.834 | -2.5 |
| 6708 | TNFLGfKWKKZ | 90 | 448.5911 | 3 | 31.65 | 1342.755 | -2.6 |
| 6766 | EFeNedKWKKZ | 90 | 516.9419 | 3 | 31.88 | 1547.808 | -2.4 |
| 7122 | gTEgfPKWKKZ | 90 | 348.9646 | 4 | 33.31 | 1391.833 | -2.8 |
| 7166 | GFPLEfKWKKZ | 90 | 452.2596 | 3 | 33.48 | 1353.76 | -2.2 |
| 7173 | gTEgfPKWKKZ | 90 | 464.9505 | 3 | 33.5 | 1391.833 | -2.4 |
| 7179 | GFPLEfKWKKZ | 90 | 452.2594 | 3 | 33.53 | 1353.76 | -2.4 |

| | | | | | | | |
|---|---|---|---|---|---|---|---|
| 7186 | gTEgfPKWKKZ | 90 | 464.9506 | 3 | 33.55 | 1391.833 | -2.2 |
| 7205 | GFPLEfKWKKZ | 90 | 452.2595 | 3 | 33.63 | 1353.76 | -2.2 |
| 7254 | gTEgfPKWKKZ | 90 | 464.9501 | 3 | 33.82 | 1391.833 | -3.2 |
| 7257 | GFPLEfKWKKZ | 90 | 452.2591 | 3 | 33.83 | 1353.76 | -3.3 |
| 7267 | gTEgfPKWKKZ | 90 | 464.9505 | 3 | 33.87 | 1391.833 | -2.5 |
| 7284 | gTEgfPKWKKZ | 90 | 464.9505 | 3 | 33.94 | 1391.833 | -2.4 |
| 7296 | GFPLEfKWKKZ | 90 | 452.2595 | 3 | 33.99 | 1353.76 | -2.2 |
| 7306 | GFPLEfKWKKZ | 90 | 452.2596 | 3 | 34.03 | 1353.76 | -2.2 |
| 7332 | GFPLEfKWKKZ | 90 | 452.2595 | 3 | 34.13 | 1353.76 | -2.2 |
| 8074 | egTQeEKWKKZ | 90 | 489.942 | 3 | 37.16 | 1466.807 | -2.1 |
| 8294 | PLLEegKWKKZ | 90 | 455.6191 | 3 | 38.1 | 1363.838 | -1.8 |
| 8303 | DTALffKWKKZ | 90 | 478.9351 | 3 | 38.14 | 1433.786 | -1.8 |
| 8306 | PLLEegKWKKZ | 90 | 455.619 | 3 | 38.16 | 1363.838 | -2 |
| 8312 | DTALffKWKKZ | 90 | 478.9347 | 3 | 38.18 | 1433.786 | -2.5 |
| 8360 | DTALffKWKKZ | 90 | 478.9349 | 3 | 38.39 | 1433.786 | -2.1 |
| 8372 | DTALffKWKKZ | 90 | 478.9346 | 3 | 38.44 | 1433.786 | -2.7 |
| 8452 | DTALffKWKKZ | 90 | 478.9348 | 3 | 38.79 | 1433.786 | -2.2 |
| 8459 | DTALffKWKKZ | 90 | 478.9348 | 3 | 38.82 | 1433.786 | -2.3 |
| 8493 | EgTeQeKWKKZ | 90 | 489.9419 | 3 | 38.97 | 1466.807 | -2.5 |
| 8537 | DTALffKWKKZ | 90 | 478.9348 | 3 | 39.15 | 1433.786 | -2.3 |
| 8549 | DTALffKWKKZ | 90 | 478.935 | 3 | 39.2 | 1433.786 | -1.9 |
| 8591 | DTALffKWKKZ | 90 | 478.9347 | 3 | 39.39 | 1433.786 | -2.5 |
| 8595 | NGfTeGKWKKZ | 90 | 446.5718 | 3 | 39.4 | 1336.708 | -10.7 |
| 8607 | GNfTeGKWKKZ | 90 | 446.5715 | 3 | 39.45 | 1336.708 | -11.4 |
| 8615 | DTALefKWKKZ | 90 | 470.2629 | 3 | 39.49 | 1407.77 | -2.5 |
| 8627 | DTALefKWKKZ | 90 | 470.2631 | 3 | 39.54 | 1407.77 | -2.1 |
| 8670 | NGfTeGKWKKZ | 90 | 446.5722 | 3 | 39.72 | 1336.708 | -10 |
| 8708 | DTALffKWKKZ | 90 | 478.935 | 3 | 39.89 | 1433.786 | -1.9 |
| 8876 | fegQbFKWKKZ | 90 | 512.9663 | 3 | 40.64 | 1535.881 | -2.3 |
| 8937 | NhEKLeKWKKZ | 90 | 452.2706 | 3 | 40.9 | 1353.792 | -1.4 |
| 8976 | NhEKLeKWKKZ | 90 | 452.2706 | 3 | 41.06 | 1353.792 | -1.5 |
| 9101 | NhEKLeKWKKZ | 90 | 452.2704 | 3 | 41.58 | 1353.792 | -2 |
| 9152 | ePLEegKWKKZ | 90 | 483.6188 | 3 | 41.8 | 1447.838 | -2.3 |
| 9161 | ePLEegKWKKZ | 90 | 483.6191 | 3 | 41.84 | 1447.838 | -1.6 |
| 9173 | ePLEegKWKKZ | 90 | 483.6188 | 3 | 41.89 | 1447.838 | -2.2 |
| 9197 | ePLEegKWKKZ | 90 | 483.619 | 3 | 42 | 1447.838 | -1.9 |
| 9209 | ePLEegKWKKZ | 90 | 483.6188 | 3 | 42.05 | 1447.838 | -2.3 |
| 9269 | ePLEegKWKKZ | 90 | 483.6192 | 3 | 42.3 | 1447.838 | -1.5 |
| 9293 | ePLEegKWKKZ | 90 | 483.6191 | 3 | 42.41 | 1447.838 | -1.6 |
| 9305 | ePLEegKWKKZ | 90 | 483.619 | 3 | 42.46 | 1447.838 | -2 |
| 9353 | ePLEegKWKKZ | 90 | 483.619 | 3 | 42.67 | 1447.838 | -1.8 |
| 9365 | ePLEegKWKKZ | 90 | 483.6191 | 3 | 42.72 | 1447.838 | -1.7 |
| 9404 | ePLEegKWKKZ | 90 | 483.6191 | 3 | 42.89 | 1447.838 | -1.6 |

| | | | | | | | |
|---|---|---|---|---|---|---|---|
| 9413 | ePLEegKWKKZ | 90 | 483.619 | 3 | 42.93 | 1447.838 | -1.8 |
| 9489 | FffbFfKWKKZ | 90 | 420.2332 | 4 | 43.26 | 1676.906 | -1.5 |
| 9498 | FffbFfKWKKZ | 90 | 420.2332 | 4 | 43.3 | 1676.906 | -1.4 |
| 9536 | ePLEegKWKKZ | 90 | 483.6192 | 3 | 43.46 | 1447.838 | -1.5 |
| 9831 | GfLfGKWWKKZ | 90 | 483.2715 | 3 | 44.77 | 1446.796 | -2.5 |
| 9833 | GfLfGKWWKKZ | 90 | 483.2715 | 3 | 44.78 | 1446.796 | -2.5 |
| 9879 | GfLfGKWWKKZ | 90 | 483.272 | 3 | 44.98 | 1446.796 | -1.6 |
| 3832 | GKEgFdKWKKZ | 89 | 338.9601 | 4 | 19.88 | 1351.813 | -1.2 |
| 4174 | KaTfQLKWKKZ | 89 | 346.2163 | 4 | 21.36 | 1380.839 | -2.3 |
| 4933 | FLAhaeKWKKZ | 89 | 326.2018 | 4 | 24.48 | 1300.781 | -2.1 |
| 5708 | fgAGbFKWKKZ | 89 | 447.2719 | 3 | 27.6 | 1338.797 | -2 |
| 5786 | fQhhbfKWKKZ | 89 | 365.465 | 4 | 27.92 | 1457.834 | -2 |
| 3145 | SFEDGGKWKKZ | 89 | 394.2074 | 3 | 16.89 | 1179.604 | -2.7 |
| 3472 | KFSLEaKWKKZ | 89 | 323.9509 | 4 | 18.34 | 1291.776 | -1.6 |
| 3847 | KGEgFdKWKKZ | 89 | 338.9597 | 4 | 19.95 | 1351.813 | -2.1 |
| 4132 | AKQfFNKWKKZ | 89 | 350.7045 | 4 | 21.18 | 1398.793 | -2.6 |
| 4977 | PKLDFLKWKKZ | 89 | 326.207 | 4 | 24.65 | 1300.802 | -2.5 |
| 4984 | PKLDFLKWKKZ | 89 | 326.2068 | 4 | 24.68 | 1300.802 | -2.9 |
| 4997 | PKLDFLKWKKZ | 89 | 326.207 | 4 | 24.73 | 1300.802 | -2.4 |
| 5160 | gKfGTPKWKKZ | 89 | 331.2084 | 4 | 25.39 | 1320.807 | -2 |
| 5173 | gKfGTPKWKKZ | 89 | 331.2082 | 4 | 25.44 | 1320.807 | -2.7 |
| 5183 | gKfGTPKWKKZ | 89 | 441.2751 | 3 | 25.48 | 1320.807 | -2.9 |
| 5195 | gKfGTPKWKKZ | 89 | 441.2754 | 3 | 25.53 | 1320.807 | -2.2 |
| 5209 | gKfGTPKWKKZ | 89 | 441.2751 | 3 | 25.58 | 1320.807 | -2.8 |
| 5245 | gKfGTPKWKKZ | 89 | 331.2082 | 4 | 25.72 | 1320.807 | -2.5 |
| 5297 | FFTFThKWKKZ | 89 | 439.5878 | 3 | 25.93 | 1315.744 | -1.9 |
| 5309 | gFgNTQKWKKZ | 89 | 444.9422 | 3 | 25.98 | 1331.808 | -2.4 |
| 5325 | fFKFNTKWKKZ | 89 | 362.9596 | 4 | 26.04 | 1447.813 | -2.3 |
| 5377 | fFKFNTKWKKZ | 89 | 362.9595 | 4 | 26.25 | 1447.813 | -2.6 |
| 5533 | fNTRfNKWKKZ | 89 | 507.2809 | 3 | 26.89 | 1518.825 | -2.5 |
| 5613 | PSFfEAKWKKZ | 89 | 448.2477 | 3 | 27.22 | 1341.723 | -1.6 |
| 5662 | ffKHgTKWKKZ | 89 | 382.7293 | 4 | 27.42 | 1526.891 | -2.1 |
| 5796 | fQhhbfKWKKZ | 89 | 365.4647 | 4 | 27.95 | 1457.834 | -2.8 |
| 5847 | fQhhbfKWKKZ | 89 | 365.4651 | 4 | 28.16 | 1457.834 | -1.7 |
| 5943 | fGENDLKWKKZ | 89 | 447.2422 | 3 | 28.56 | 1338.709 | -2.7 |
| 6129 | FfQEhLKWKKZ | 89 | 471.9383 | 3 | 29.33 | 1412.797 | -2.6 |
| 6167 | FSfGgRKWKKZ | 89 | 462.6103 | 3 | 29.49 | 1384.813 | -3 |
| 6528 | eAFfdEKWKKZ | 89 | 511.2785 | 3 | 30.93 | 1530.817 | -2.4 |
| 6538 | eAFfdEKWKKZ | 89 | 383.7108 | 4 | 30.97 | 1530.817 | -2.3 |
| 6571 | fFKeSDKWKKZ | 89 | 372.2076 | 4 | 31.1 | 1484.797 | 2.8 |
| 6657 | afNfgAKWKKZ | 89 | 362.4648 | 4 | 31.45 | 1445.834 | -2.3 |
| 6693 | afNfgAKWKKZ | 89 | 362.4648 | 4 | 31.6 | 1445.834 | -2.5 |
| 6797 | afNfgAKWKKZ | 89 | 362.4647 | 4 | 32 | 1445.834 | -2.7 |

| | | | | | | | |
|---|---|---|---|---|---|---|---|
| 6813 | afNfgAKWKKZ | 89 | 362.4648 | 4 | 32.07 | 1445.834 | -2.5 |
| 6901 | afNfgAKWKKZ | 89 | 362.4647 | 4 | 32.42 | 1445.834 | -2.6 |
| 6998 | afNfgAKWKKZ | 89 | 362.4648 | 4 | 32.81 | 1445.834 | -2.4 |
| 7244 | GFPLEfKWKKZ | 89 | 452.2596 | 3 | 33.78 | 1353.76 | -2.2 |
| 7273 | GFPLEfKWKKZ | 89 | 452.2593 | 3 | 33.9 | 1353.76 | -2.6 |
| 7283 | GFPLEfKWKKZ | 89 | 452.2596 | 3 | 33.94 | 1353.76 | -2.2 |
| 7291 | gTEgfPKWKKZ | 89 | 464.9506 | 3 | 33.97 | 1391.833 | -2.3 |
| 7310 | gTEgfPKWKKZ | 89 | 464.9504 | 3 | 34.05 | 1391.833 | -2.5 |
| 7356 | PfgghPKWKKZ | 89 | 336.9685 | 4 | 34.23 | 1343.848 | -2.6 |
| 7638 | GfAfFQKWKKZ | 89 | 479.9315 | 3 | 35.41 | 1436.776 | -2 |
| 7768 | GfAfFQKWKKZ | 89 | 479.9316 | 3 | 35.93 | 1436.776 | -1.8 |
| 8162 | egTQeEKWKKZ | 89 | 489.942 | 3 | 37.52 | 1466.807 | -2.1 |
| 8186 | egTQeEKWKKZ | 89 | 489.9419 | 3 | 37.63 | 1466.807 | -2.3 |
| 8196 | DTALffKWKKZ | 89 | 478.9348 | 3 | 37.67 | 1433.786 | -2.4 |
| 8246 | egTQeEKWKKZ | 89 | 489.9423 | 3 | 37.89 | 1466.807 | -1.6 |
| 8348 | DTALffKWKKZ | 89 | 478.9348 | 3 | 38.34 | 1433.786 | -2.3 |
| 8384 | DTALffKWKKZ | 89 | 478.9346 | 3 | 38.49 | 1433.786 | -2.7 |
| 8396 | DTALffKWKKZ | 89 | 478.9345 | 3 | 38.54 | 1433.786 | -2.9 |
| 8408 | DTALffKWKKZ | 89 | 478.9348 | 3 | 38.59 | 1433.786 | -2.3 |
| 8430 | DTALffKWKKZ | 89 | 478.9348 | 3 | 38.69 | 1433.786 | -2.3 |
| 8442 | DTALffKWKKZ | 89 | 478.9348 | 3 | 38.74 | 1433.786 | -2.2 |
| 8471 | DTALffKWKKZ | 89 | 478.9349 | 3 | 38.87 | 1433.786 | -2.1 |
| 8482 | DTALffKWKKZ | 89 | 478.9348 | 3 | 38.92 | 1433.786 | -2.3 |
| 8489 | DTALffKWKKZ | 89 | 478.9348 | 3 | 38.95 | 1433.786 | -2.3 |
| 8561 | DTALffKWKKZ | 89 | 478.9347 | 3 | 39.26 | 1433.786 | -2.5 |
| 8585 | NGfTeGKWKKZ | 89 | 446.5716 | 3 | 39.36 | 1336.708 | -11.3 |
| 8603 | DTALefKWKKZ | 89 | 470.2629 | 3 | 39.44 | 1407.77 | -2.4 |
| 8658 | GNfTeGKWKKZ | 89 | 446.5717 | 3 | 39.66 | 1336.708 | -11.1 |
| 8747 | DTALffKWKKZ | 89 | 478.9352 | 3 | 40.07 | 1433.786 | -1.6 |
| 9246 | FFLEegKWKKZ | 89 | 362.9659 | 4 | 42.21 | 1447.838 | -2.5 |
| 9317 | ePLEegKWKKZ | 89 | 483.6192 | 3 | 42.51 | 1447.838 | -1.5 |
| 9332 | ePLEegKWKKZ | 89 | 483.6191 | 3 | 42.58 | 1447.838 | -1.6 |
| 9341 | ePLEegKWKKZ | 89 | 483.6191 | 3 | 42.62 | 1447.838 | -1.7 |
| 9575 | ePLEegKWKKZ | 89 | 483.6191 | 3 | 43.63 | 1447.838 | -1.6 |
| 9611 | ePLEegKWKKZ | 89 | 483.6188 | 3 | 43.79 | 1447.838 | -2.2 |
| 9647 | ePLEegKWKKZ | 89 | 483.6187 | 3 | 43.96 | 1447.838 | -2.5 |
| 6590 | SNFLGfKWKKZ | 88 | 443.9192 | 3 | 31.18 | 1328.739 | -2.6 |
| 8778 | FFNeAeKWKKZ | 88 | 366.2002 | 4 | 40.21 | 1460.776 | -2.7 |
| 3862 | GKEgFdKWKKZ | 88 | 338.9598 | 4 | 20.01 | 1351.813 | -1.9 |
| 4102 | AKQfFNKWKKZ | 88 | 350.7045 | 4 | 21.05 | 1398.793 | -2.6 |
| 4138 | KaTfQLKWKKZ | 88 | 346.2164 | 4 | 21.21 | 1380.839 | -2 |
| 4159 | AKQfFNKWKKZ | 88 | 350.7046 | 4 | 21.3 | 1398.793 | -2.3 |
| 4753 | EAfaLGKWKKZ | 88 | 427.9192 | 3 | 23.74 | 1280.739 | -2.7 |

| | | | | | | | |
|---|---|---|---|---|---|---|---|
| 4783 | DhfaLGKWKKZ | 88 | 427.9194 | 3 | 23.87 | 1280.739 | -2.2 |
| 4798 | EAfaLGKWKKZ | 88 | 427.9194 | 3 | 23.93 | 1280.739 | -2.2 |
| 5010 | PKLDFLKWKKZ | 88 | 326.207 | 4 | 24.78 | 1300.802 | -2.4 |
| 5190 | gKfGTPKWKKZ | 88 | 331.2083 | 4 | 25.5 | 1320.807 | -2.3 |
| 5232 | gKfGTPKWKKZ | 88 | 331.2084 | 4 | 25.67 | 1320.807 | -2 |
| 5237 | gKfGTPKWKKZ | 88 | 441.2754 | 3 | 25.69 | 1320.807 | -2 |
| 5292 | FFTFThKWKKZ | 88 | 439.5879 | 3 | 25.91 | 1315.744 | -1.7 |
| 5335 | fFKFNTKWKKZ | 88 | 362.9598 | 4 | 26.08 | 1447.813 | -1.7 |
| 5361 | fFKFNTKWKKZ | 88 | 362.9596 | 4 | 26.19 | 1447.813 | -2.3 |
| 5442 | fFKFNTKWKKZ | 88 | 362.9598 | 4 | 26.51 | 1447.813 | -1.9 |
| 5488 | fNTRfNKWKKZ | 88 | 380.7127 | 4 | 26.71 | 1518.825 | -2 |
| 5514 | fNTRfNKWKKZ | 88 | 380.7126 | 4 | 26.81 | 1518.825 | -2.1 |
| 5517 | NLhLgQKWKKZ | 88 | 423.6107 | 3 | 26.82 | 1267.813 | -2.2 |
| 5605 | NLhLgQKWKKZ | 88 | 423.6109 | 3 | 27.19 | 1267.813 | -1.6 |
| 5623 | PSFfEAKWKKZ | 88 | 448.2477 | 3 | 27.26 | 1341.723 | -1.5 |
| 5631 | PSFfEAKWKKZ | 88 | 336.4375 | 4 | 27.29 | 1341.723 | -1.9 |
| 5691 | ffKHgTKWKKZ | 88 | 382.7295 | 4 | 27.54 | 1526.891 | -1.7 |
| 5698 | ffKHgTKWKKZ | 88 | 382.7293 | 4 | 27.56 | 1526.891 | -2 |
| 6003 | DfdDfKKWKKZ | 88 | 392.9647 | 4 | 28.81 | 1567.834 | -2.8 |
| 6065 | DfdDfKKWKKZ | 88 | 392.9647 | 4 | 29.06 | 1567.834 | -2.7 |
| 6093 | DfdDfKKWKKZ | 88 | 392.9651 | 4 | 29.18 | 1567.834 | -1.8 |
| 6139 | FfQEhLKWKKZ | 88 | 471.9384 | 3 | 29.37 | 1412.797 | -2.5 |
| 6142 | SaeDfQKWKKZ | 88 | 360.4452 | 4 | 29.38 | 1437.756 | -2.7 |
| 6162 | FfQEhLKWKKZ | 88 | 354.2052 | 4 | 29.47 | 1412.797 | -3.7 |
| 6177 | FSfGgRKWKKZ | 88 | 462.6104 | 3 | 29.53 | 1384.813 | -2.9 |
| 6190 | FSfGgRKWKKZ | 88 | 462.6107 | 3 | 29.58 | 1384.813 | -2.2 |
| 6203 | FSfGgRKWKKZ | 88 | 462.6104 | 3 | 29.63 | 1384.813 | -2.9 |
| 6521 | eAFfdEKWKKZ | 88 | 383.7109 | 4 | 30.9 | 1530.817 | -1.9 |
| 6551 | eAFfdEKWKKZ | 88 | 383.711 | 4 | 31.02 | 1530.817 | -1.5 |
| 6554 | eAFfdEKWKKZ | 88 | 511.279 | 3 | 31.03 | 1530.817 | -1.5 |
| 6667 | afNfgAKWKKZ | 88 | 362.4649 | 4 | 31.49 | 1445.834 | -2.1 |
| 6774 | afNfgAKWKKZ | 88 | 362.4648 | 4 | 31.91 | 1445.834 | -2.4 |
| 6855 | afNfgAKWKKZ | 88 | 362.4647 | 4 | 32.23 | 1445.834 | -2.6 |
| 6881 | afNfgAKWKKZ | 88 | 362.4648 | 4 | 32.33 | 1445.834 | -2.3 |
| 6927 | afNfgAKWKKZ | 88 | 362.4647 | 4 | 32.52 | 1445.834 | -2.7 |
| 6953 | afNfgAKWKKZ | 88 | 362.4648 | 4 | 32.63 | 1445.834 | -2.5 |
| 7200 | PfgghPKWKKZ | 88 | 336.9686 | 4 | 33.61 | 1343.848 | -2.4 |
| 7322 | GFPLEfKWKKZ | 88 | 452.2596 | 3 | 34.09 | 1353.76 | -2.1 |
| 7574 | GfAfFQKWKKZ | 88 | 479.9315 | 3 | 35.14 | 1436.776 | -2.2 |
| 7724 | APPeeNKWKKZ | 88 | 454.5879 | 3 | 35.75 | 1360.744 | -1.8 |
| 7755 | APPeeNKWKKZ | 88 | 454.5879 | 3 | 35.88 | 1360.744 | -1.9 |
| 8420 | DTALffKWKKZ | 88 | 478.9348 | 3 | 38.65 | 1433.786 | -2.2 |
| 8516 | EgTeQeKWKKZ | 88 | 489.9418 | 3 | 39.07 | 1466.807 | -2.5 |

| | | | | | | | |
|---|---|---|---|---|---|---|---|
| 8696 | DTALffKWKKZ | 88 | 478.9349 | 3 | 39.83 | 1433.786 | -2.2 |
| 8903 | fegQbFKWKKZ | 88 | 384.9769 | 4 | 40.76 | 1535.881 | -1.5 |
| 8952 | NhEKLeKWKKZ | 88 | 452.2706 | 3 | 40.96 | 1353.792 | -1.5 |
| 8965 | NhEKLeKWKKZ | 88 | 452.2701 | 3 | 41.02 | 1353.792 | -2.6 |
| 8978 | NhEKLeKWKKZ | 88 | 452.2706 | 3 | 41.07 | 1353.792 | -1.5 |
| 8991 | NhEKLeKWKKZ | 88 | 452.2705 | 3 | 41.12 | 1353.792 | -1.6 |
| 9004 | NhEKLeKWKKZ | 88 | 452.2705 | 3 | 41.18 | 1353.792 | -1.7 |
| 9017 | NhEKLeKWKKZ | 88 | 452.2701 | 3 | 41.23 | 1353.792 | -2.6 |
| 9030 | NhEKLeKWKKZ | 88 | 452.2699 | 3 | 41.29 | 1353.792 | -3 |
| 9068 | NhEKLeKWKKZ | 88 | 452.2701 | 3 | 41.44 | 1353.792 | -2.6 |
| 9081 | NhEKLeKWKKZ | 88 | 452.2704 | 3 | 41.5 | 1353.792 | -2 |
| 9294 | PeLEegKWKKZ | 88 | 362.9661 | 4 | 42.41 | 1447.838 | -1.9 |
| 9465 | FffbFfKWKKZ | 88 | 420.2333 | 4 | 43.15 | 1676.906 | -1.2 |
| 9659 | ePLEegKWKKZ | 88 | 483.6191 | 3 | 44.01 | 1447.838 | -1.6 |
| 3883 | AAKDLFKWKKZ | 87 | 309.1916 | 4 | 20.1 | 1232.739 | -1.7 |
| 6389 | FdDDfFKWKKZ | 87 | 378.7004 | 4 | 30.38 | 1510.776 | -2.5 |
| 8703 | LNTQeEKWKKZ | 87 | 457.5862 | 3 | 39.86 | 1369.751 | -10 |
| 11244 | eLhPLdKWKKZ | 87 | 457.2876 | 3 | 50.98 | 1368.843 | -1.7 |
| 3172 | FSEDGGKWKKZ | 87 | 394.2078 | 3 | 17.01 | 1179.604 | -1.6 |
| 3448 | KFSLEaKWKKZ | 87 | 323.9506 | 4 | 18.23 | 1291.776 | -2.3 |
| 3484 | KFSLEaKWKKZ | 87 | 323.9506 | 4 | 18.39 | 1291.776 | -2.5 |
| 4111 | AKQfFNKWKKZ | 87 | 350.7044 | 4 | 21.09 | 1398.793 | -2.7 |
| 4120 | AKQfFNKWKKZ | 87 | 350.7045 | 4 | 21.12 | 1398.793 | -2.5 |
| 4144 | AKQfFNKWKKZ | 87 | 350.7047 | 4 | 21.23 | 1398.793 | -2.1 |
| 4173 | AgTLGNKWKKZ | 87 | 391.2475 | 3 | 21.35 | 1170.724 | -2.5 |
| 5224 | gKfGTPKWKKZ | 87 | 441.2751 | 3 | 25.64 | 1320.807 | -2.8 |
| 5315 | FFTFThKWKKZ | 87 | 439.5876 | 3 | 26 | 1315.744 | -2.3 |
| 5328 | FFTFThKWKKZ | 87 | 439.5877 | 3 | 26.05 | 1315.744 | -2.2 |
| 5403 | fFKFNTKWKKZ | 87 | 362.9596 | 4 | 26.35 | 1447.813 | -2.4 |
| 5416 | fFKFNTKWKKZ | 87 | 362.9598 | 4 | 26.41 | 1447.813 | -1.8 |
| 5501 | fNTRfNKWKKZ | 87 | 380.7127 | 4 | 26.76 | 1518.825 | -1.9 |
| 5540 | NLhLgQKWKKZ | 87 | 423.6108 | 3 | 26.92 | 1267.813 | -1.9 |
| 5809 | fQhhbfKWKKZ | 87 | 365.4648 | 4 | 28.01 | 1457.834 | -2.6 |
| 5860 | fQhhbfKWKKZ | 87 | 365.4646 | 4 | 28.21 | 1457.834 | -3 |
| 5902 | fQhhbfKWKKZ | 87 | 365.4648 | 4 | 28.39 | 1457.834 | -2.5 |
| 5990 | DfdDfKKWKKZ | 87 | 392.9648 | 4 | 28.75 | 1567.834 | -2.5 |
| 6023 | DfdDfKKWKKZ | 87 | 392.9651 | 4 | 28.89 | 1567.834 | -1.6 |
| 6078 | DfdDfKKWKKZ | 87 | 392.9649 | 4 | 29.12 | 1567.834 | -2.2 |
| 6102 | SaeDfQKWKKZ | 87 | 360.4453 | 4 | 29.22 | 1437.756 | -2.4 |
| 6111 | SaeDfQKWKKZ | 87 | 360.4451 | 4 | 29.26 | 1437.756 | -2.9 |
| 6121 | FfQEhLKWKKZ | 87 | 354.2056 | 4 | 29.3 | 1412.797 | -2.6 |
| 6126 | SaeDfQKWKKZ | 87 | 360.445 | 4 | 29.32 | 1437.756 | -3.3 |
| 6134 | FfQEhLKWKKZ | 87 | 354.2056 | 4 | 29.35 | 1412.797 | -2.5 |

| | | | | | | | |
|---|---|---|---|---|---|---|---|
| 6147 | FfQEhLKWKKZ | 87 | 354.2056 | 4 | 29.4 | 1412.797 | -2.5 |
| 6152 | FfQEhLKWKKZ | 87 | 471.9384 | 3 | 29.43 | 1412.797 | -2.6 |
| 6170 | FfQEhLKWKKZ | 87 | 471.938 | 3 | 29.5 | 1412.797 | -3.4 |
| 6515 | fFKeSDKWKKZ | 87 | 372.2057 | 4 | 30.88 | 1484.797 | -2 |
| 6525 | eAFfdEKWKKZ | 87 | 383.7107 | 4 | 30.92 | 1530.817 | -2.4 |
| 6577 | eAFfdEKWKKZ | 87 | 383.7107 | 4 | 31.13 | 1530.817 | -2.4 |
| 6673 | afNfgAKWKKZ | 87 | 482.9508 | 3 | 31.52 | 1445.834 | -1.9 |
| 6683 | afNfgAKWKKZ | 87 | 482.9507 | 3 | 31.56 | 1445.834 | -2.2 |
| 6748 | afNfgAKWKKZ | 87 | 362.4648 | 4 | 31.81 | 1445.834 | -2.4 |
| 6761 | afNfgAKWKKZ | 87 | 362.4647 | 4 | 31.86 | 1445.834 | -2.6 |
| 6787 | afNfgAKWKKZ | 87 | 362.4648 | 4 | 31.96 | 1445.834 | -2.4 |
| 7651 | GfAfFQKWKKZ | 87 | 479.9317 | 3 | 35.46 | 1436.776 | -1.7 |
| 8198 | DTALffKWKKZ | 87 | 478.9348 | 3 | 37.68 | 1433.786 | -2.4 |
| 8787 | FFNeAeKWKKZ | 87 | 366.2003 | 4 | 40.24 | 1460.776 | -2.4 |
| 9043 | NhEKLeKWKKZ | 87 | 452.2702 | 3 | 41.34 | 1353.792 | -2.4 |
| 9056 | NhEKLeKWKKZ | 87 | 452.2701 | 3 | 41.4 | 1353.792 | -2.6 |
| 9093 | NhEKLeKWKKZ | 87 | 452.2703 | 3 | 41.55 | 1353.792 | -2.1 |
| 9119 | NhEKLeKWKKZ | 87 | 452.2708 | 3 | 41.66 | 1353.792 | -1.1 |
| 9284 | ePLEegKWKKZ | 87 | 362.9661 | 4 | 42.37 | 1447.838 | -1.9 |
| 9881 | GfLfGKWWKKZ | 87 | 483.272 | 3 | 44.99 | 1446.796 | -1.6 |
| 11246 | eLhPLdKWKKZ | 87 | 457.2876 | 3 | 50.99 | 1368.843 | -1.7 |
| 11292 | eLhPLdKWKKZ | 87 | 457.2873 | 3 | 51.19 | 1368.843 | -2.4 |
| 4913 | dAQgfGKWKKZ | 86 | 343.4569 | 4 | 24.4 | 1369.802 | -2.6 |
| 4964 | GffGGaKWKKZ | 86 | 435.9126 | 3 | 24.6 | 1304.718 | -1.8 |
| 5650 | hfFgSbKWKKZ | 86 | 346.7124 | 4 | 27.37 | 1382.823 | -1.7 |
| 3454 | KFSLEaKWKKZ | 86 | 323.9507 | 4 | 18.26 | 1291.776 | -1.9 |
| 3802 | KGEgFdKWKKZ | 86 | 338.9598 | 4 | 19.75 | 1351.813 | -1.9 |
| 3874 | AAKDLFKWKKZ | 86 | 309.1915 | 4 | 20.06 | 1232.739 | -1.9 |
| 3895 | AAKDLFKWKKZ | 86 | 309.1914 | 4 | 20.15 | 1232.739 | -2.1 |
| 4128 | AKQfFNKWKKZ | 86 | 467.2703 | 3 | 21.16 | 1398.793 | -2.4 |
| 4185 | AgTLGNKWKKZ | 86 | 391.2477 | 3 | 21.41 | 1170.724 | -2 |
| 4768 | hDfaLGKWKKZ | 86 | 427.9195 | 3 | 23.8 | 1280.739 | -2 |
| 4967 | FKeNFKKWKKZ | 86 | 363.2178 | 4 | 24.61 | 1448.845 | -1.6 |
| 5073 | KTfhKfKWKKZ | 86 | 369.9769 | 4 | 25.04 | 1475.881 | -1.5 |
| 5112 | KTfhKfKWKKZ | 86 | 369.9766 | 4 | 25.19 | 1475.881 | -2.3 |
| 5125 | KTfhKfKWKKZ | 86 | 369.9763 | 4 | 25.25 | 1475.881 | -2.9 |
| 5136 | TKfhKfKWKKZ | 86 | 492.9661 | 3 | 25.29 | 1475.881 | -2.8 |
| 5177 | gKfGTPKWKKZ | 86 | 331.2082 | 4 | 25.45 | 1320.807 | -2.6 |
| 5185 | gKfGTPKWKKZ | 86 | 441.2751 | 3 | 25.49 | 1320.807 | -2.9 |
| 5203 | KgfGTPKWKKZ | 86 | 331.2082 | 4 | 25.56 | 1320.807 | -2.7 |
| 5216 | KgfGTPKWKKZ | 86 | 331.2082 | 4 | 25.61 | 1320.807 | -2.5 |
| 5276 | FFTFThKWKKZ | 86 | 439.5876 | 3 | 25.85 | 1315.744 | -2.3 |
| 5299 | FFTFThKWKKZ | 86 | 439.5878 | 3 | 25.94 | 1315.744 | -1.9 |

| | | | | | | | |
|---|---|---|---|---|---|---|---|
| 5348 | fFKFNTKWKKZ | 86 | 362.9598 | 4 | 26.13 | 1447.813 | -1.9 |
| 5390 | fFKFNTKWKKZ | 86 | 362.9596 | 4 | 26.3 | 1447.813 | -2.5 |
| 5520 | fNTRfNKWKKZ | 86 | 507.2813 | 3 | 26.83 | 1518.825 | -1.7 |
| 5527 | NLhLgQKWKKZ | 86 | 423.6104 | 3 | 26.86 | 1267.813 | -2.8 |
| 5543 | fNTRfNKWKKZ | 86 | 380.7127 | 4 | 26.93 | 1518.825 | -2 |
| 5553 | NLhLgQKWKKZ | 86 | 423.6108 | 3 | 26.97 | 1267.813 | -2 |
| 5579 | NLhLgQKWKKZ | 86 | 423.6109 | 3 | 27.08 | 1267.813 | -1.6 |
| 5586 | PSFfEAKKWKZ | 86 | 336.4375 | 4 | 27.11 | 1341.723 | -1.8 |
| 5634 | PSFfEAKWKKZ | 86 | 448.2477 | 3 | 27.31 | 1341.723 | -1.5 |
| 5695 | fgAGbFKWKKZ | 86 | 447.272 | 3 | 27.55 | 1338.797 | -1.8 |
| 5835 | fQhhbfKWKKZ | 86 | 365.4647 | 4 | 28.11 | 1457.834 | -2.7 |
| 5872 | fQhhbfKWKKZ | 86 | 365.4648 | 4 | 28.26 | 1457.834 | -2.6 |
| 6094 | SaeDfQKWKKZ | 86 | 360.4456 | 4 | 29.19 | 1437.756 | -1.7 |
| 6647 | TNFLGfKWKKZ | 86 | 672.3831 | 2 | 31.41 | 1342.755 | -2.4 |
| 6738 | afNfgAKWKKZ | 86 | 482.9507 | 3 | 31.77 | 1445.834 | -2.3 |
| 6868 | afNfgAKWKKZ | 86 | 362.4648 | 4 | 32.28 | 1445.834 | -2.4 |
| 7037 | afNfgAKWKKZ | 86 | 362.4647 | 4 | 32.96 | 1445.834 | -2.7 |
| 7135 | PfgghPKWKKZ | 86 | 336.9685 | 4 | 33.36 | 1343.848 | -2.6 |
| 7726 | APPeeNKWKKZ | 86 | 454.5879 | 3 | 35.76 | 1360.744 | -1.8 |
| 7781 | GfAfFQKWKKZ | 86 | 479.9315 | 3 | 35.98 | 1436.776 | -2.2 |
| 8597 | GNfTeGKWKKZ | 86 | 446.5718 | 3 | 39.41 | 1336.708 | -10.7 |
| 8621 | NGfTeGKWKKZ | 86 | 446.5721 | 3 | 39.51 | 1336.708 | -10 |
| 8643 | TbTQeEKWKKZ | 86 | 457.5858 | 3 | 39.6 | 1369.751 | -11.2 |
| 8891 | fegQbFKWKKZ | 86 | 384.9765 | 4 | 40.7 | 1535.881 | -2.4 |
| 8939 | NhEKLeKWKKZ | 86 | 452.2706 | 3 | 40.91 | 1353.792 | -1.4 |
| 9103 | NhEKLeKWKKZ | 86 | 452.2704 | 3 | 41.59 | 1353.792 | -2 |
| 9215 | ePLEegKWKKZ | 86 | 362.966 | 4 | 42.07 | 1447.838 | -2.1 |
| 9308 | ePLEegKWKKZ | 86 | 362.9661 | 4 | 42.47 | 1447.838 | -1.9 |
| 9476 | FffbFfKWKKZ | 86 | 420.2331 | 4 | 43.2 | 1676.906 | -1.6 |
| 11184 | eLhPLdKWKKZ | 86 | 457.2875 | 3 | 50.72 | 1368.843 | -1.9 |
| 11294 | eLhPLdKWKKZ | 86 | 457.2873 | 3 | 51.2 | 1368.843 | -2.4 |
| 3958 | RKLgKLKWKKZ | 85 | 339.2438 | 4 | 20.42 | 1352.95 | -2.7 |
| 4050 | GGaLgLKWKKZ | 85 | 385.928 | 3 | 20.82 | 1154.765 | -2.5 |
| 4234 | NhEKLeKWKKZ | 85 | 339.4546 | 4 | 21.62 | 1353.792 | -1.9 |
| 10275 | QHNfgAKWKKZ | 85 | 463.5999 | 3 | 46.72 | 1387.788 | -7.1 |
| 3117 | SFEDGGKWKKZ | 85 | 394.2078 | 3 | 16.77 | 1179.604 | -1.7 |
| 3141 | SFEDGGKWKKZ | 85 | 394.2076 | 3 | 16.88 | 1179.604 | -2.2 |
| 3159 | SFEDGGKWKKZ | 85 | 394.2078 | 3 | 16.96 | 1179.604 | -1.7 |
| 3811 | GKEgFdKWKKZ | 85 | 338.9597 | 4 | 19.79 | 1351.813 | -2.2 |
| 3871 | GKEgFdKWKKZ | 85 | 338.9599 | 4 | 20.05 | 1351.813 | -1.7 |
| 3922 | AAKDLFKWKKZ | 85 | 309.191 | 4 | 20.27 | 1232.739 | -3.4 |
| 4027 | RKLgKLKWKKZ | 85 | 339.2438 | 4 | 20.72 | 1352.95 | -2.7 |
| 5151 | TKfhKfKWKKZ | 85 | 369.9766 | 4 | 25.35 | 1475.881 | -2.2 |

| | | | | | | | |
|---|---|---|---|---|---|---|---|
| 5167 | TKfhKfKWKKZ | 85 | 369.9765 | 4 | 25.42 | 1475.881 | -2.6 |
| 5211 | gKfGTPKWKKZ | 85 | 441.2751 | 3 | 25.59 | 1320.807 | -2.8 |
| 5302 | gFgNTQKWKKZ | 85 | 333.9586 | 4 | 25.95 | 1331.808 | -2.1 |
| 5429 | fFKFNTKWKKZ | 85 | 362.96 | 4 | 26.46 | 1447.813 | -1.4 |
| 5432 | QeLbdgKWKKZ | 85 | 364.7295 | 4 | 26.47 | 1454.891 | -1.7 |
| 5572 | fNTRfNKWKKZ | 85 | 380.7126 | 4 | 27.05 | 1518.825 | -2.1 |
| 5591 | NLhLgQKWKKZ | 85 | 423.6105 | 3 | 27.13 | 1267.813 | -2.5 |
| 5669 | fgAGbFKWKKZ | 85 | 447.2719 | 3 | 27.45 | 1338.797 | -2 |
| 5822 | fQhhbfKWKKZ | 85 | 365.4651 | 4 | 28.06 | 1457.834 | -1.8 |
| 6013 | DfdDfKKWKKZ | 85 | 392.9652 | 4 | 28.85 | 1567.834 | -1.5 |
| 6039 | DfdDfKKWKKZ | 85 | 392.9648 | 4 | 28.95 | 1567.834 | -2.6 |
| 6161 | FSfGgRKWKKZ | 85 | 347.2096 | 4 | 29.46 | 1384.813 | -2.9 |
| 6174 | FSfGgRKWKKZ | 85 | 347.2097 | 4 | 29.51 | 1384.813 | -2.5 |
| 6187 | FSfGgRKWKKZ | 85 | 347.2097 | 4 | 29.57 | 1384.813 | -2.5 |
| 6200 | FSfGgRKWKKZ | 85 | 347.2097 | 4 | 29.62 | 1384.813 | -2.7 |
| 6531 | fFKeSDKWKKZ | 85 | 372.2056 | 4 | 30.94 | 1484.797 | -2.5 |
| 6660 | TNFLGfKWKKZ | 85 | 672.3832 | 2 | 31.46 | 1342.755 | -2.2 |
| 6803 | afNfgAKWKKZ | 85 | 482.9505 | 3 | 32.03 | 1445.834 | -2.6 |
| 6940 | afNfgAKWKKZ | 85 | 362.4648 | 4 | 32.57 | 1445.834 | -2.4 |
| 7008 | afNfgAKWKKZ | 85 | 362.4645 | 4 | 32.85 | 1445.834 | -3.2 |
| 8364 | DTALffKWKKZ | 85 | 717.8981 | 2 | 38.41 | 1433.786 | -2.9 |
| 8431 | DTALffKWKKZ | 85 | 717.8983 | 2 | 38.7 | 1433.786 | -2.7 |
| 8609 | GNfTeGKWKKZ | 85 | 446.5715 | 3 | 39.46 | 1336.708 | -11.4 |
| 8631 | TbTQeEKWKKZ | 85 | 457.5858 | 3 | 39.55 | 1369.751 | -11 |
| 8648 | NGfTeGKWKKZ | 85 | 446.5716 | 3 | 39.62 | 1336.708 | -11.3 |
| 8660 | NGfTeGKWKKZ | 85 | 446.5717 | 3 | 39.67 | 1336.708 | -11.1 |
| 8667 | TbTQeEKWKKZ | 85 | 457.586 | 3 | 39.7 | 1369.751 | -10.7 |
| 8691 | TbTQeEKWKKZ | 85 | 457.586 | 3 | 39.81 | 1369.751 | -10.7 |
| 8715 | LNTQeEKWKKZ | 85 | 457.5858 | 3 | 39.92 | 1369.751 | -10.8 |
| 9176 | ePLEegKWKKZ | 85 | 362.9661 | 4 | 41.91 | 1447.838 | -1.9 |
| 9191 | fALEegKWKKZ | 85 | 362.9659 | 4 | 41.97 | 1447.838 | -2.2 |
| 9360 | ePLEegKWKKZ | 85 | 362.966 | 4 | 42.7 | 1447.838 | -2 |
| 9525 | FffbFfKWKKZ | 85 | 420.233 | 4 | 43.42 | 1676.906 | -1.8 |
| 11172 | eLhPLdKWKKZ | 85 | 457.2879 | 3 | 50.67 | 1368.843 | -1.1 |
| 11186 | eLhPLdKWKKZ | 85 | 457.2875 | 3 | 50.73 | 1368.843 | -1.9 |
| 11268 | eLhPLdKWKKZ | 85 | 457.2875 | 3 | 51.09 | 1368.843 | -1.9 |
| 11280 | eLhPLdKWKKZ | 85 | 457.288 | 3 | 51.14 | 1368.843 | -0.7 |
| 11304 | eLhPLdKWKKZ | 85 | 457.2878 | 3 | 51.24 | 1368.843 | -1.1 |
| 4744 | EFGGedKWKKZ | 84 | 451.2473 | 3 | 23.7 | 1350.724 | -2.7 |
| 5495 | FFNRhfKWKKZ | 84 | 365.9623 | 4 | 26.73 | 1459.824 | -2.5 |
| 8538 | NEHTeGKWKKZ | 84 | 441.9 | 3 | 39.16 | 1322.688 | -7.7 |
| 4074 | GGaLgLKWKKZ | 84 | 385.9279 | 3 | 20.93 | 1154.765 | -2.7 |
| 4137 | AKQfFNKWKKZ | 84 | 467.2702 | 3 | 21.2 | 1398.793 | -2.7 |

| | | | | | | | |
|---|---|---|---|---|---|---|---|
| 4149 | AgTLGNKWKKZ | 84 | 293.6877 | 4 | 21.25 | 1170.724 | -1.8 |
| 4152 | AKQfFNKWKKZ | 84 | 467.2706 | 3 | 21.26 | 1398.793 | -1.7 |
| 4194 | AgTLGNKWKKZ | 84 | 293.6874 | 4 | 21.44 | 1170.724 | -2.5 |
| 4197 | AgTLGNKWKKZ | 84 | 391.2474 | 3 | 21.46 | 1170.724 | -2.8 |
| 4248 | KfGQdLKWKKZ | 84 | 354.2176 | 4 | 21.67 | 1412.845 | -2.2 |
| 4398 | bgRdGfKWKKZ | 84 | 364.2281 | 4 | 22.29 | 1452.887 | -2.6 |
| 4418 | bgRdGfKWKKZ | 84 | 364.2282 | 4 | 22.37 | 1452.887 | -2.1 |
| 4522 | bgRdGfKWKKZ | 84 | 364.2281 | 4 | 22.79 | 1452.887 | -2.5 |
| 4548 | bgRdGfKWKKZ | 84 | 364.2283 | 4 | 22.89 | 1452.887 | -2.1 |
| 4855 | RTfQAFKWKKZ | 84 | 354.4575 | 4 | 24.16 | 1413.803 | -1.8 |
| 5086 | TKfhKfKWKKZ | 84 | 369.9765 | 4 | 25.09 | 1475.881 | -2.4 |
| 5091 | TKfhKfKWKKZ | 84 | 492.9663 | 3 | 25.11 | 1475.881 | -2.3 |
| 5099 | KTfhKfKWKKZ | 84 | 369.9767 | 4 | 25.14 | 1475.881 | -2.1 |
| 5138 | TKfhKfKWKKZ | 84 | 369.9765 | 4 | 25.3 | 1475.881 | -2.5 |
| 5263 | gFgNTQKWKKZ | 84 | 333.9584 | 4 | 25.79 | 1331.808 | -2.5 |
| 5286 | gFgNTQKWKKZ | 84 | 333.9586 | 4 | 25.89 | 1331.808 | -1.9 |
| 5433 | eLhPLdKWKKZ | 84 | 457.2878 | 3 | 26.48 | 1368.843 | -1.3 |
| 5530 | fNTRfNKWKKZ | 84 | 380.7125 | 4 | 26.88 | 1518.825 | -2.4 |
| 5566 | NLhLgQKWKKZ | 84 | 423.6106 | 3 | 27.03 | 1267.813 | -2.3 |
| 5659 | fgAGbFKWKKZ | 84 | 447.272 | 3 | 27.41 | 1338.797 | -1.9 |
| 5720 | fgAGbFKWKKZ | 84 | 447.2714 | 3 | 27.65 | 1338.797 | -3.1 |
| 6052 | DfdDfKKWKKZ | 84 | 392.9648 | 4 | 29.01 | 1567.834 | -2.5 |
| 6630 | TNFLGfKWKKZ | 84 | 336.6952 | 4 | 31.34 | 1342.755 | -2.4 |
| 6688 | TNFLGfKWKKZ | 84 | 336.6952 | 4 | 31.57 | 1342.755 | -2.3 |
| 6703 | afNfgAKWKKZ | 84 | 362.4647 | 4 | 31.63 | 1445.834 | -2.6 |
| 6972 | afNfgAKWKKZ | 84 | 362.465 | 4 | 32.7 | 1445.834 | -1.8 |
| 7278 | PfgghPKWKKZ | 84 | 448.9557 | 3 | 33.92 | 1343.848 | -2.4 |
| 7286 | gTEgfPKWKKZ | 84 | 464.9505 | 3 | 33.95 | 1391.833 | -2.4 |
| 7325 | gTEgPfKWKKZ | 84 | 464.9506 | 3 | 34.1 | 1391.833 | -2.1 |
| 7853 | egTQeEKWKKZ | 84 | 367.7082 | 4 | 36.27 | 1466.807 | -2.4 |
| 8378 | DTALffKWKKZ | 84 | 359.4526 | 4 | 38.47 | 1433.786 | -3.1 |
| 8426 | DTALffKWKKZ | 84 | 359.4531 | 4 | 38.67 | 1433.786 | -1.8 |
| 8436 | DTALffKWKKZ | 84 | 359.4531 | 4 | 38.72 | 1433.786 | -1.7 |
| 8443 | DTALffKWKKZ | 84 | 717.8982 | 2 | 38.75 | 1433.786 | -2.8 |
| 8460 | DTALffKWKKZ | 84 | 717.8975 | 2 | 38.83 | 1433.786 | -3.8 |
| 8495 | EgTeQeKWKKZ | 84 | 489.9419 | 3 | 38.98 | 1466.807 | -2.5 |
| 8636 | GNfTeGKWKKZ | 84 | 446.5716 | 3 | 39.57 | 1336.708 | -11.3 |
| 8795 | fegQbFKWKKZ | 84 | 384.9766 | 4 | 40.28 | 1535.881 | -2.3 |
| 9227 | ePLEegKWKKZ | 84 | 362.9659 | 4 | 42.13 | 1447.838 | -2.2 |
| 9236 | ePLEegKWKKZ | 84 | 362.966 | 4 | 42.16 | 1447.838 | -2 |
| 9248 | FFLEegKWKKZ | 84 | 362.9659 | 4 | 42.21 | 1447.838 | -2.5 |
| 9260 | ePLEegKWKKZ | 84 | 362.9659 | 4 | 42.27 | 1447.838 | -2.2 |
| 9272 | ePLEegKWKKZ | 84 | 362.9661 | 4 | 42.32 | 1447.838 | -1.7 |

| | | | | | | | |
|---|---|---|---|---|---|---|---|
| 9467 | FffbFfKWKKZ | 84 | 420.2333 | 4 | 43.16 | 1676.906 | -1.2 |
| 10287 | HQNfgAKWKKZ | 84 | 463.5997 | 3 | 46.77 | 1387.788 | -7.5 |
| 10299 | QHNfgAKWKKZ | 84 | 463.6002 | 3 | 46.82 | 1387.788 | -6.6 |
| 11157 | eLhPLdKWKKZ | 84 | 457.2874 | 3 | 50.61 | 1368.843 | -2.1 |
| 11174 | eLhPLdKWKKZ | 84 | 457.2879 | 3 | 50.68 | 1368.843 | -1.1 |
| 11232 | eLhPLdKWKKZ | 84 | 457.2878 | 3 | 50.93 | 1368.843 | -1.3 |
| 11256 | eLhPLdKWKKZ | 84 | 457.2877 | 3 | 51.03 | 1368.843 | -1.5 |
| 11270 | eLhPLdKWKKZ | 84 | 457.2875 | 3 | 51.09 | 1368.843 | -1.9 |
| 11282 | eLhPLdKWKKZ | 84 | 457.288 | 3 | 51.15 | 1368.843 | -0.7 |
| 11306 | eLhPLdKWKKZ | 84 | 457.2878 | 3 | 51.25 | 1368.843 | -1.1 |
| 3407 | hFEEADKWKKZ | 83 | 316.9222 | 4 | 18.05 | 1263.661 | -1.1 |
| 4359 | AhAfgGKWKKZ | 83 | 408.2563 | 3 | 22.14 | 1221.739 | 6.9 |
| 10120 | NHNfgAKWKKZ | 83 | 458.928 | 3 | 46.06 | 1373.772 | -7 |
| 3823 | GKEgFdKWKKZ | 83 | 338.9598 | 4 | 19.84 | 1351.813 | -1.9 |
| 3849 | KGEgFdKWKKZ | 83 | 338.9597 | 4 | 19.95 | 1351.813 | -2.1 |
| 3933 | TaDLKeKWKKZ | 83 | 336.4546 | 4 | 20.31 | 1341.792 | -2.1 |
| 3943 | RKLgKLKWKKZ | 83 | 339.2439 | 4 | 20.36 | 1352.95 | -2.5 |
| 3964 | RKLgKLKWKKZ | 83 | 339.2439 | 4 | 20.44 | 1352.95 | -2.3 |
| 4032 | GGaLgLKWKKZ | 83 | 385.9281 | 3 | 20.74 | 1154.765 | -2.3 |
| 4098 | GGaLgLKWKKZ | 83 | 385.928 | 3 | 21.03 | 1154.765 | -2.5 |
| 4158 | AgTLGNKWKKZ | 83 | 293.6875 | 4 | 21.29 | 1170.724 | -2.2 |
| 4444 | bgRdGfKWKKZ | 83 | 364.2282 | 4 | 22.48 | 1452.887 | -2.3 |
| 4447 | fhNRLTKWKKZ | 83 | 345.9611 | 4 | 22.49 | 1379.819 | -2.8 |
| 4470 | bgRdGfKWKKZ | 83 | 364.2281 | 4 | 22.58 | 1452.887 | -2.4 |
| 4483 | bgRdGfKWKKZ | 83 | 364.2281 | 4 | 22.63 | 1452.887 | -2.6 |
| 4499 | fhNRLTKWKKZ | 83 | 345.9611 | 4 | 22.69 | 1379.819 | -2.8 |
| 4577 | bgRdGfKWKKZ | 83 | 364.2282 | 4 | 23.01 | 1452.887 | -2.2 |
| 4596 | bgRdGfKWKKZ | 83 | 364.2282 | 4 | 23.09 | 1452.887 | -2.3 |
| 4990 | GffGGaKWKKZ | 83 | 435.9122 | 3 | 24.7 | 1304.718 | -2.7 |
| 5000 | GffGGaKWKKZ | 83 | 435.9124 | 3 | 24.74 | 1304.718 | -2.3 |
| 5107 | TKfhKfKWKKZ | 83 | 492.9666 | 3 | 25.18 | 1475.881 | -1.8 |
| 5123 | TKfhKfKWKKZ | 83 | 492.9664 | 3 | 25.24 | 1475.881 | -2.2 |
| 5337 | fFKFNTKWKKZ | 83 | 483.6109 | 3 | 26.09 | 1447.813 | -1.4 |
| 5418 | fFKFNTKWKKZ | 83 | 483.6109 | 3 | 26.41 | 1447.813 | -1.4 |
| 5435 | eLhPLdKWKKZ | 83 | 457.2878 | 3 | 26.48 | 1368.843 | -1.3 |
| 5556 | fNTRfNKWKKZ | 83 | 380.7127 | 4 | 26.98 | 1518.825 | -2 |
| 5607 | NLhLgQKWKKZ | 83 | 423.6109 | 3 | 27.2 | 1267.813 | -1.6 |
| 5682 | fgAGbFKWKKZ | 83 | 447.272 | 3 | 27.5 | 1338.797 | -1.9 |
| 5711 | ffKHgTKWKKZ | 83 | 382.7293 | 4 | 27.62 | 1526.891 | -2.2 |
| 5887 | fQhhbfKWKKZ | 83 | 365.4648 | 4 | 28.33 | 1457.834 | -2.5 |
| 6649 | TNFLGfKWKKZ | 83 | 672.3831 | 2 | 31.42 | 1342.755 | -2.4 |
| 6696 | afNfgAKWKKZ | 83 | 482.9506 | 3 | 31.61 | 1445.834 | -2.5 |
| 6719 | afNfgAKWKKZ | 83 | 362.4648 | 4 | 31.7 | 1445.834 | -2.5 |

| | | | | | | | |
|---|---|---|---|---|---|---|---|
| 7148 | PfgghPKWKKZ | 83 | 336.9685 | 4 | 33.41 | 1343.848 | -2.6 |
| 7174 | PfgghPKWKKZ | 83 | 336.9686 | 4 | 33.51 | 1343.848 | -2.4 |
| 7268 | PfgghPKWKKZ | 83 | 448.9556 | 3 | 33.88 | 1343.848 | -2.5 |
| 7293 | gTEgfPKWKKZ | 83 | 464.9506 | 3 | 33.98 | 1391.833 | -2.3 |
| 7297 | PfgghPKWKKZ | 83 | 448.9557 | 3 | 33.99 | 1343.848 | -2.2 |
| 7307 | PfgghPKWKKZ | 83 | 448.9557 | 3 | 34.03 | 1343.848 | -2.4 |
| 7986 | egTQeEKWKKZ | 83 | 367.7085 | 4 | 36.8 | 1466.807 | -1.8 |
| 8051 | egTQeEKWKKZ | 83 | 367.7082 | 4 | 37.06 | 1466.807 | -2.6 |
| 8366 | DTALffKWKKZ | 83 | 717.8981 | 2 | 38.41 | 1433.786 | -2.9 |
| 8385 | DTALffKWKKZ | 83 | 717.8979 | 2 | 38.5 | 1433.786 | -3.2 |
| 8411 | DTALffKWKKZ | 83 | 359.4525 | 4 | 38.61 | 1433.786 | -3.4 |
| 8433 | DTALffKWKKZ | 83 | 717.8983 | 2 | 38.7 | 1433.786 | -2.7 |
| 8453 | DTALffKWKKZ | 83 | 717.8981 | 2 | 38.8 | 1433.786 | -2.9 |
| 8622 | TbTQeEKWKKZ | 83 | 457.5863 | 3 | 39.52 | 1369.751 | -10 |
| 8705 | TbTQeEKWKKZ | 83 | 457.5862 | 3 | 39.87 | 1369.751 | -10.2 |
| 9296 | PeLEegKWKKZ | 83 | 362.9661 | 4 | 42.42 | 1447.838 | -1.9 |
| 9335 | ePLEegKWKKZ | 83 | 362.9661 | 4 | 42.59 | 1447.838 | -1.8 |
| 9441 | FffbFfKWKKZ | 83 | 420.233 | 4 | 43.05 | 1676.906 | -1.8 |
| 9491 | FffbFfKWKKZ | 83 | 420.2332 | 4 | 43.27 | 1676.906 | -1.5 |
| 11234 | eLhPLdKWKKZ | 83 | 457.2878 | 3 | 50.94 | 1368.843 | -1.3 |
| 11258 | eLhPLdKWKKZ | 83 | 457.2877 | 3 | 51.04 | 1368.843 | -1.5 |
| 5266 | hPPASEKWKKZ | 82 | 385.5613 | 3 | 25.81 | 1153.661 | 1.1 |
| 3174 | FSEDGGKWKKZ | 82 | 394.2078 | 3 | 17.02 | 1179.604 | -1.6 |
| 3398 | hFEEADKWKKZ | 82 | 316.9218 | 4 | 18.02 | 1263.661 | -2.5 |
| 4038 | GGaLgLKWKKZ | 82 | 385.928 | 3 | 20.77 | 1154.765 | -2.6 |
| 4062 | GGaLgLKWKKZ | 82 | 385.9279 | 3 | 20.87 | 1154.765 | -2.7 |
| 4086 | GGaLgLKWKKZ | 82 | 385.9281 | 3 | 20.98 | 1154.765 | -2.3 |
| 4170 | AgTLGNKWKKZ | 82 | 293.6875 | 4 | 21.34 | 1170.724 | -2.3 |
| 4182 | AgTLGNKWKKZ | 82 | 293.6876 | 4 | 21.39 | 1170.724 | -2 |
| 4375 | AhAfgGKWKKZ | 82 | 408.2563 | 3 | 22.2 | 1221.739 | 6.8 |
| 4388 | bgRdGfKWKKZ | 82 | 364.2281 | 4 | 22.25 | 1452.887 | -2.6 |
| 4440 | fhNRLTKWKKZ | 82 | 345.9613 | 4 | 22.46 | 1379.819 | -2.1 |
| 4457 | bgRdGfKWKKZ | 82 | 364.2283 | 4 | 22.53 | 1452.887 | -2.1 |
| 4460 | fhNRLTKWKKZ | 82 | 345.9612 | 4 | 22.54 | 1379.819 | -2.3 |
| 4473 | fhNRLTKWKKZ | 82 | 345.9612 | 4 | 22.59 | 1379.819 | -2.6 |
| 4486 | fhNRLTKWKKZ | 82 | 345.9611 | 4 | 22.64 | 1379.819 | -2.7 |
| 4496 | bgRdGfKWKKZ | 82 | 364.2281 | 4 | 22.68 | 1452.887 | -2.5 |
| 4535 | bgRdGfKWKKZ | 82 | 364.2282 | 4 | 22.84 | 1452.887 | -2.1 |
| 4756 | EFGGedKWKKZ | 82 | 451.2474 | 3 | 23.75 | 1350.724 | -2.4 |
| 4835 | TRfhNFKWKKZ | 82 | 472.2737 | 3 | 24.08 | 1413.804 | -2.9 |
| 4974 | GffGGaKWKKZ | 82 | 435.9124 | 3 | 24.64 | 1304.718 | -2.3 |
| 4980 | FKeNFKKWKKZ | 82 | 363.2176 | 4 | 24.66 | 1448.845 | -2.3 |
| 5279 | hPPASEKWKKZ | 82 | 385.5613 | 3 | 25.86 | 1153.661 | 1 |

| | | | | | | | |
|---|---|---|---|---|---|---|---|
| 5289 | hPPASEKWKKZ | 82 | 385.5615 | 3 | 25.9 | 1153.661 | 1.6 |
| 5350 | fFKFNTKWKKZ | 82 | 483.6106 | 3 | 26.14 | 1447.813 | -2 |
| 5366 | fFKFNTKWKKZ | 82 | 483.6103 | 3 | 26.2 | 1447.813 | -2.6 |
| 5395 | fFKFNTKWKKZ | 82 | 483.6103 | 3 | 26.32 | 1447.813 | -2.6 |
| 5405 | fFKFNTKWKKZ | 82 | 483.6104 | 3 | 26.36 | 1447.813 | -2.4 |
| 5446 | fFKFNTKWKKZ | 82 | 483.6105 | 3 | 26.53 | 1447.813 | -2 |
| 5447 | fFKFNTKWKKZ | 82 | 483.6105 | 3 | 26.53 | 1447.813 | -2 |
| 5798 | fQhhbfKWKKZ | 82 | 365.4647 | 4 | 27.96 | 1457.834 | -2.8 |
| 5849 | fQhhbfKWKKZ | 82 | 365.4651 | 4 | 28.17 | 1457.834 | -1.7 |
| 6213 | FSfGgRKWKKZ | 82 | 347.2097 | 4 | 29.67 | 1384.813 | -2.4 |
| 6217 | FSfGgRKWKKZ | 82 | 462.6107 | 3 | 29.69 | 1384.813 | -2.2 |
| 6391 | FdDDfFKWKKZ | 82 | 378.7004 | 4 | 30.38 | 1510.776 | -2.5 |
| 6514 | fFKeSDKWKKZ | 82 | 372.2057 | 4 | 30.87 | 1484.797 | -2 |
| 6643 | TNFLGfKWKKZ | 82 | 336.6952 | 4 | 31.39 | 1342.755 | -2.4 |
| 6656 | TNFLGfKWKKZ | 82 | 336.6952 | 4 | 31.45 | 1342.755 | -2.5 |
| 7161 | PfgghPKWKKZ | 82 | 336.9686 | 4 | 33.46 | 1343.848 | -2.2 |
| 7190 | PfgghPKWKKZ | 82 | 336.9686 | 4 | 33.57 | 1343.848 | -2.4 |
| 7317 | PfgghPKWKKZ | 82 | 448.9557 | 3 | 34.07 | 1343.848 | -2.2 |
| 7583 | gbDeDfKWKKZ | 82 | 373.7081 | 4 | 35.18 | 1490.807 | -2.7 |
| 7885 | egTQeEKWKKZ | 82 | 367.7082 | 4 | 36.4 | 1466.807 | -2.6 |
| 7911 | egTQeEKWKKZ | 82 | 367.7083 | 4 | 36.5 | 1466.807 | -2.4 |
| 7950 | egTQeEKWKKZ | 82 | 367.7082 | 4 | 36.66 | 1466.807 | -2.6 |
| 8387 | DTALffKWKKZ | 82 | 717.8979 | 2 | 38.5 | 1433.786 | -3.2 |
| 8462 | DTALffKWKKZ | 82 | 717.8975 | 2 | 38.84 | 1433.786 | -3.8 |
| 8473 | EgTeQeKWKKZ | 82 | 367.7085 | 4 | 38.88 | 1466.807 | -1.8 |
| 8529 | NEHTeGKWKKZ | 82 | 441.8999 | 3 | 39.12 | 1322.688 | -7.8 |
| 8550 | NEHTeGKWKKZ | 82 | 441.9 | 3 | 39.21 | 1322.688 | -7.7 |
| 8586 | NEHTeGKWKKZ | 82 | 441.8998 | 3 | 39.37 | 1322.688 | -8.1 |
| 8672 | GNfTeGKWKKZ | 82 | 446.5722 | 3 | 39.72 | 1336.708 | -10 |
| 8681 | TbTQeEKWKKZ | 82 | 457.5862 | 3 | 39.77 | 1369.751 | -10.2 |
| 8693 | TbTQeEKWKKZ | 82 | 457.586 | 3 | 39.82 | 1369.751 | -10.7 |
| 8733 | TbTQeEKWKKZ | 82 | 457.5859 | 3 | 40 | 1369.751 | -10.9 |
| 9456 | ePLEegKWKKZ | 82 | 362.966 | 4 | 43.12 | 1447.838 | -2 |
| 9500 | FffbFfKWKKZ | 82 | 420.2332 | 4 | 43.3 | 1676.906 | -1.4 |
| 11159 | eLhPLdKWKKZ | 82 | 457.2874 | 3 | 50.61 | 1368.843 | -2.1 |
| 4206 | PhPGGHKWKKZ | 81 | 373.558 | 3 | 21.5 | 1117.651 | 1.2 |
| 4991 | begDTbKWKKZ | 81 | 345.9583 | 4 | 24.71 | 1379.808 | -2.9 |
| 6466 | AGPhfeKWKKZ | 81 | 440.2523 | 3 | 30.68 | 1317.739 | -2.7 |
| 3129 | FSEDGGKWKKZ | 81 | 394.2075 | 3 | 16.83 | 1179.604 | -2.4 |
| 3497 | KFSLEaKWKKZ | 81 | 323.9505 | 4 | 18.45 | 1291.776 | -2.8 |
| 3498 | KFSLEaKWKKZ | 81 | 323.9505 | 4 | 18.45 | 1291.776 | -2.8 |
| 3834 | GKEgFdKWKKZ | 81 | 338.9601 | 4 | 19.89 | 1351.813 | -1.2 |
| 3864 | GKEgFdKWKKZ | 81 | 338.9598 | 4 | 20.02 | 1351.813 | -1.9 |

| | | | | | | | |
|---|---|---|---|---|---|---|---|
| 4110 | GGaLgLKWKKZ | 81 | 385.928 | 3 | 21.08 | 1154.765 | -2.6 |
| 4408 | bgRdGfKWKKZ | 81 | 364.2282 | 4 | 22.33 | 1452.887 | -2.3 |
| 4431 | bgRdGfKWKKZ | 81 | 364.2284 | 4 | 22.42 | 1452.887 | -1.6 |
| 4735 | EFGGedKWKKZ | 81 | 451.2475 | 3 | 23.67 | 1350.724 | -2.2 |
| 4800 | TRfQAFKWKKZ | 81 | 354.4572 | 4 | 23.93 | 1413.803 | -2.4 |
| 4813 | TRfQAFKWKKZ | 81 | 354.4574 | 4 | 23.99 | 1413.803 | -1.8 |
| 4826 | TRfQAFKWKKZ | 81 | 354.4572 | 4 | 24.04 | 1413.803 | -2.5 |
| 4943 | FLAhaeKWKKZ | 81 | 326.202 | 4 | 24.52 | 1300.781 | -1.4 |
| 5004 | begDTbKWKKZ | 81 | 345.9584 | 4 | 24.76 | 1379.808 | -2.4 |
| 5449 | FFNRhfKWKKZ | 81 | 365.9624 | 4 | 26.54 | 1459.824 | -2.3 |
| 5476 | FFNRhfKWKKZ | 81 | 365.9625 | 4 | 26.66 | 1459.824 | -2.2 |
| 5483 | fNTRfNKWKKZ | 81 | 380.7128 | 4 | 26.68 | 1518.825 | -1.8 |
| 5597 | PSFfEAKWKKZ | 81 | 336.4374 | 4 | 27.16 | 1341.723 | -2.2 |
| 5610 | PSFfEAKWKKZ | 81 | 336.4376 | 4 | 27.21 | 1341.723 | -1.7 |
| 5620 | PSFfEAKWKKZ | 81 | 336.4376 | 4 | 27.25 | 1341.723 | -1.6 |
| 5904 | fQhhbfKWKKZ | 81 | 365.4648 | 4 | 28.4 | 1457.834 | -2.5 |
| 6118 | FfQEhLKWKKZ | 81 | 354.2055 | 4 | 29.29 | 1412.797 | -2.7 |
| 6517 | fFKeSDKWKKZ | 81 | 495.9386 | 3 | 30.88 | 1484.797 | -2 |
| 6530 | fFKeSDKWKKZ | 81 | 372.2056 | 4 | 30.93 | 1484.797 | -2.5 |
| 6533 | fFKeSDKWKKZ | 81 | 495.9384 | 3 | 30.95 | 1484.797 | -2.4 |
| 6543 | fFKeSDKWKKZ | 81 | 372.2056 | 4 | 30.99 | 1484.797 | -2.3 |
| 6546 | fFKeSDKWKKZ | 81 | 495.9384 | 3 | 31 | 1484.797 | -2.4 |
| 6662 | TNFLGfKWKKZ | 81 | 672.3832 | 2 | 31.47 | 1342.755 | -2.2 |
| 6672 | TNFLGfKWKKZ | 81 | 336.6954 | 4 | 31.51 | 1342.755 | -1.9 |
| 7078 | gTEgfPKWKKZ | 81 | 348.9647 | 4 | 33.13 | 1391.833 | -2.5 |
| 7098 | gTEgfPKWKKZ | 81 | 348.9647 | 4 | 33.21 | 1391.833 | -2.3 |
| 7111 | gTEgfPKWKKZ | 81 | 348.9646 | 4 | 33.26 | 1391.833 | -2.8 |
| 7124 | gTEgfPKWKKZ | 81 | 348.9646 | 4 | 33.31 | 1391.833 | -2.8 |
| 7312 | gTEgfPKWKKZ | 81 | 464.9504 | 3 | 34.05 | 1391.833 | -2.5 |
| 7333 | PfgghPKWKKZ | 81 | 448.9557 | 3 | 34.14 | 1343.848 | -2.4 |
| 7554 | gbDeDfKWKKZ | 81 | 373.7082 | 4 | 35.06 | 1490.807 | -2.4 |
| 7567 | gbDeDfKWKKZ | 81 | 373.7083 | 4 | 35.11 | 1490.807 | -2.2 |
| 7586 | gbDeDfKWKKZ | 81 | 497.9421 | 3 | 35.19 | 1490.807 | -2 |
| 7599 | gbDeDfKWKKZ | 81 | 373.7082 | 4 | 35.24 | 1490.807 | -2.4 |
| 7924 | egTQeEKWKKZ | 81 | 367.7083 | 4 | 36.55 | 1466.807 | -2.4 |
| 7927 | egTQeEKWKKZ | 81 | 489.9422 | 3 | 36.57 | 1466.807 | -1.8 |
| 7937 | egTQeEKWKKZ | 81 | 367.7083 | 4 | 36.61 | 1466.807 | -2.4 |
| 7943 | egTQeEKWKKZ | 81 | 489.942 | 3 | 36.63 | 1466.807 | -2.1 |
| 7963 | egTQeEKWKKZ | 81 | 367.7083 | 4 | 36.71 | 1466.807 | -2.3 |
| 7999 | egTQeEKWKKZ | 81 | 367.7084 | 4 | 36.85 | 1466.807 | -2 |
| 8067 | egTQeEKWKKZ | 81 | 367.7082 | 4 | 37.13 | 1466.807 | -2.5 |
| 8421 | DTALffKWKKZ | 81 | 717.8982 | 2 | 38.65 | 1433.786 | -2.8 |
| 8445 | DTALffKWKKZ | 81 | 717.8982 | 2 | 38.76 | 1433.786 | -2.8 |

| | | | | | | | |
|---|---|---|---|---|---|---|---|
| 8562 | NHETeGKWKKZ | 81 | 441.8998 | 3 | 39.26 | 1322.688 | -8 |
| 8624 | TbTQeEKWKKZ | 81 | 457.5863 | 3 | 39.52 | 1369.751 | -10 |
| 8645 | TbTQeEKWKKZ | 81 | 457.5858 | 3 | 39.61 | 1369.751 | -11.2 |
| 8811 | FFNeAeKWKKZ | 81 | 366.2002 | 4 | 40.35 | 1460.776 | -2.6 |
| 8829 | FFNeAeKWKKZ | 81 | 366.2001 | 4 | 40.43 | 1460.776 | -2.9 |
| 10311 | HQNfgAKWKKZ | 81 | 463.5998 | 3 | 46.87 | 1387.788 | -7.5 |
| 11145 | eLhPLdKWKKZ | 81 | 457.2876 | 3 | 50.55 | 1368.843 | -1.7 |
| 2893 | LAhSNDKWKKZ | 80 | 294.1734 | 4 | 15.79 | 1172.667 | -1.8 |
| 3781 | gKLESEKWKKZ | 80 | 434.6019 | 3 | 19.66 | 1300.787 | -2 |
| 4261 | dTKLQfKWKKZ | 80 | 365.2239 | 4 | 21.73 | 1456.871 | -2.8 |
| 7491 | fAfFAGKWKKZ | 80 | 460.9241 | 3 | 34.8 | 1379.754 | -2.7 |
| 8490 | ASSSggKWKKZ | 80 | 392.2487 | 3 | 38.96 | 1173.723 | 0.6 |
| 9198 | ESiSfGKKKKZ | 80 | 422.9051 | 3 | 42 | 1265.688 | 4.2 |
| 9914 | aGiEegKWKKZ | 80 | 451.2632 | 3 | 45.13 | 1350.756 | 8.5 |
| 3419 | hFEEADKWKKZ | 80 | 316.9219 | 4 | 18.11 | 1263.661 | -2 |
| 3471 | KFSLEaKWKKZ | 80 | 323.9509 | 4 | 18.33 | 1291.776 | -1.6 |
| 3782 | gKLESEKWKKZ | 80 | 434.6019 | 3 | 19.66 | 1300.787 | -2 |
| 4140 | KaTfQLKWKKZ | 80 | 346.2164 | 4 | 21.21 | 1380.839 | -2 |
| 4305 | AhAfgGKWKKZ | 80 | 408.2562 | 3 | 21.91 | 1221.739 | 6.5 |
| 4330 | AhAfgGKWKKZ | 80 | 408.2564 | 3 | 22.02 | 1221.739 | 7 |
| 4346 | AhAfgGKWKKZ | 80 | 408.2564 | 3 | 22.08 | 1221.739 | 7 |
| 4509 | bgRdGfKWKKZ | 80 | 364.2282 | 4 | 22.73 | 1452.887 | -2.2 |
| 4531 | fhNRLTKWKKZ | 80 | 345.9611 | 4 | 22.82 | 1379.819 | -2.8 |
| 4609 | bgRdGfKWKKZ | 80 | 364.2281 | 4 | 23.14 | 1452.887 | -2.6 |
| 4822 | TRfQAFKWKKZ | 80 | 472.2743 | 3 | 24.03 | 1413.803 | -1.5 |
| 4839 | TRfQAFKWKKZ | 80 | 354.4574 | 4 | 24.09 | 1413.803 | -1.9 |
| 4848 | TRfQAFKWKKZ | 80 | 472.274 | 3 | 24.13 | 1413.803 | -2.1 |
| 4861 | TRfQAFKWKKZ | 80 | 472.2744 | 3 | 24.19 | 1413.803 | -1.3 |
| 4871 | TRfQAFKWKKZ | 80 | 354.4573 | 4 | 24.23 | 1413.803 | -2.1 |
| 5162 | KgfGTPKWKKZ | 80 | 331.2084 | 4 | 25.4 | 1320.807 | -2 |
| 5389 | fFKFNTKWKKZ | 80 | 362.9596 | 4 | 26.3 | 1447.813 | -2.5 |
| 5428 | fFKFNTKWKKZ | 80 | 362.96 | 4 | 26.45 | 1447.813 | -1.4 |
| 5636 | PSFfEAKWKKZ | 80 | 448.2477 | 3 | 27.31 | 1341.723 | -1.5 |
| 5862 | fQhhbfKWKKZ | 80 | 365.4646 | 4 | 28.22 | 1457.834 | -3 |
| 5915 | ffKHgTKWKKZ | 80 | 382.7294 | 4 | 28.44 | 1526.891 | -1.9 |
| 6116 | SaeDfQKWKKZ | 80 | 480.258 | 3 | 29.28 | 1437.756 | -2.5 |
| 6489 | AGPhfeKWKKZ | 80 | 440.2528 | 3 | 30.77 | 1317.739 | -1.5 |
| 6544 | fFKeSDKWKKZ | 80 | 372.2056 | 4 | 30.99 | 1484.797 | -2.3 |
| 6559 | fFKeSDKWKKZ | 80 | 495.9389 | 3 | 31.05 | 1484.797 | -1.4 |
| 7169 | gTEgfPKWKKZ | 80 | 348.9647 | 4 | 33.49 | 1391.833 | -2.4 |
| 7280 | PfgghPKWKKZ | 80 | 448.9557 | 3 | 33.92 | 1343.848 | -2.4 |
| 7309 | PfgghPKWKKZ | 80 | 448.9557 | 3 | 34.04 | 1343.848 | -2.4 |
| 7544 | gbDeDfKWKKZ | 80 | 373.7083 | 4 | 35.02 | 1490.807 | -2.2 |

| | | | | | | | |
|---|---|---|---|---|---|---|---|
| 7816 | egTQeEKWKKZ | 80 | 489.9422 | 3 | 36.12 | 1466.807 | -1.8 |
| 7842 | egTQeEKWKKZ | 80 | 489.9419 | 3 | 36.23 | 1466.807 | -2.5 |
| 7855 | egTQeEKWKKZ | 80 | 489.942 | 3 | 36.28 | 1466.807 | -2.2 |
| 7869 | egTQeEKWKKZ | 80 | 367.7081 | 4 | 36.34 | 1466.807 | -2.9 |
| 7881 | egTQeEKWKKZ | 80 | 489.942 | 3 | 36.38 | 1466.807 | -2.3 |
| 7894 | egTQeEKWKKZ | 80 | 489.942 | 3 | 36.44 | 1466.807 | -2.2 |
| 7898 | egTQeEKWKKZ | 80 | 367.7082 | 4 | 36.45 | 1466.807 | -2.5 |
| 7904 | egTQeEKWKKZ | 80 | 489.9421 | 3 | 36.47 | 1466.807 | -2 |
| 7917 | egTQeEKWKKZ | 80 | 489.942 | 3 | 36.53 | 1466.807 | -2.2 |
| 7953 | egTQeEKWKKZ | 80 | 489.9421 | 3 | 36.67 | 1466.807 | -1.9 |
| 7966 | egTQeEKWKKZ | 80 | 489.9421 | 3 | 36.72 | 1466.807 | -2 |
| 7975 | egTQeEKWKKZ | 80 | 367.7083 | 4 | 36.75 | 1466.807 | -2.4 |
| 7976 | egTQeEKWKKZ | 80 | 367.7083 | 4 | 36.76 | 1466.807 | -2.4 |
| 8005 | egTQeEKWKKZ | 80 | 489.942 | 3 | 36.87 | 1466.807 | -2.1 |
| 8038 | egTQeEKWKKZ | 80 | 367.7082 | 4 | 37.01 | 1466.807 | -2.5 |
| 8455 | DTALffKWKKZ | 80 | 717.8981 | 2 | 38.8 | 1433.786 | -2.9 |
| 8485 | hfFgSgKWKKZ | 80 | 346.9674 | 4 | 38.94 | 1383.843 | -2.1 |
| 8540 | NEHTeGKWKKZ | 80 | 441.9 | 3 | 39.17 | 1322.688 | -7.7 |
| 8633 | TbTQeEKWKKZ | 80 | 457.5858 | 3 | 39.56 | 1369.751 | -11 |
| 8655 | TbTQeEKWKKZ | 80 | 457.586 | 3 | 39.65 | 1369.751 | -10.7 |
| 8657 | TbTQeEKWKKZ | 80 | 457.586 | 3 | 39.66 | 1369.751 | -10.7 |
| 8669 | TbTQeEKWKKZ | 80 | 457.586 | 3 | 39.71 | 1369.751 | -10.7 |
| 8717 | LNTQeEKWKKZ | 80 | 457.5858 | 3 | 39.93 | 1369.751 | -10.8 |
| 8799 | FFNeAeKWKKZ | 80 | 366.2003 | 4 | 40.3 | 1460.776 | -2.5 |
| 9451 | fALEegKWKKZ | 80 | 483.619 | 3 | 43.09 | 1447.838 | -1.8 |
| 2851 | AhLSQQKWKKZ | 79 | 400.9111 | 3 | 15.62 | 1199.714 | -2 |
| 2929 | hLASNDKWKKZ | 79 | 294.1733 | 4 | 15.95 | 1172.667 | -1.9 |
| 3886 | fEPAHGKWKKZ | 79 | 326.4325 | 4 | 20.11 | 1301.703 | -1.8 |
| 4538 | GGgRLFKWKKZ | 79 | 415.9354 | 3 | 22.85 | 1244.787 | -2.3 |
| 4736 | fFKFDaKWKKZ | 79 | 362.9596 | 4 | 23.67 | 1447.813 | -2.4 |
| 2869 | AhLSQQKWKKZ | 79 | 300.9353 | 4 | 15.69 | 1199.714 | -1.4 |
| 3147 | SFEDGGKWKKZ | 79 | 394.2074 | 3 | 16.9 | 1179.604 | -2.7 |
| 3453 | KFSLEaKWKKZ | 79 | 323.9507 | 4 | 18.25 | 1291.776 | -1.9 |
| 3483 | KFSLEaKWKKZ | 79 | 323.9506 | 4 | 18.38 | 1291.776 | -2.5 |
| 4262 | dTKLQfKWKKZ | 79 | 365.2239 | 4 | 21.73 | 1456.871 | -2.8 |
| 4317 | AhAfgGKWKKZ | 79 | 408.256 | 3 | 21.96 | 1221.739 | 6.2 |
| 4567 | bgRdGfKWKKZ | 79 | 364.2283 | 4 | 22.97 | 1452.887 | -2.1 |
| 4661 | bgRdGfKWKKZ | 79 | 364.2283 | 4 | 23.36 | 1452.887 | -2 |
| 4737 | fFKFDaKWKKZ | 79 | 362.9596 | 4 | 23.67 | 1447.813 | -2.4 |
| 4920 | FLAhaeKWKKZ | 79 | 326.2018 | 4 | 24.43 | 1300.781 | -2.1 |
| 5015 | PKLDFLKWKKZ | 79 | 434.6068 | 3 | 24.8 | 1300.802 | -2.5 |
| 5188 | gaFQRfKWKKZ | 79 | 368.2266 | 4 | 25.5 | 1468.882 | -3.1 |
| 5205 | KgfGTPKWKKZ | 79 | 331.2082 | 4 | 25.57 | 1320.807 | -2.7 |

| | | | | | | | |
|---|---|---|---|---|---|---|---|
| 5218 | KgfGTPKWKKZ | 79 | 331.2082 | 4 | 25.62 | 1320.807 | -2.5 |
| 5273 | gFgNTQKWKKZ | 79 | 333.9584 | 4 | 25.83 | 1331.808 | -2.4 |
| 5324 | fFKFNTKWKKZ | 79 | 362.9596 | 4 | 26.04 | 1447.813 | -2.3 |
| 5334 | fFKFNTKWKKZ | 79 | 362.9598 | 4 | 26.08 | 1447.813 | -1.7 |
| 5347 | fFKFNTKWKKZ | 79 | 362.9598 | 4 | 26.13 | 1447.813 | -1.9 |
| 5360 | fFKFNTKWKKZ | 79 | 362.9596 | 4 | 26.18 | 1447.813 | -2.3 |
| 5376 | fFKFNTKWKKZ | 79 | 362.9595 | 4 | 26.24 | 1447.813 | -2.6 |
| 5402 | fFKFNTKWKKZ | 79 | 362.9596 | 4 | 26.35 | 1447.813 | -2.4 |
| 5415 | fFKFNTKWKKZ | 79 | 362.9598 | 4 | 26.4 | 1447.813 | -1.8 |
| 5461 | FFNRhfKWKKZ | 79 | 365.9625 | 4 | 26.59 | 1459.824 | -2.2 |
| 5487 | fNTRfNKWKKZ | 79 | 380.7127 | 4 | 26.7 | 1518.825 | -2 |
| 5500 | fNTRfNKWKKZ | 79 | 380.7127 | 4 | 26.75 | 1518.825 | -1.9 |
| 5503 | fNTRfNKWKKZ | 79 | 507.2813 | 3 | 26.76 | 1518.825 | -1.7 |
| 5513 | fNTRfNKWKKZ | 79 | 380.7126 | 4 | 26.81 | 1518.825 | -2.1 |
| 5529 | fNTRfNKWKKZ | 79 | 380.7125 | 4 | 26.87 | 1518.825 | -2.4 |
| 5633 | PSFfEAKWKKZ | 79 | 336.4375 | 4 | 27.3 | 1341.723 | -1.9 |
| 5811 | fQhhbfKWKKZ | 79 | 365.4648 | 4 | 28.02 | 1457.834 | -2.6 |
| 5837 | fQhhbfKWKKZ | 79 | 365.4647 | 4 | 28.12 | 1457.834 | -2.7 |
| 6106 | DfdDfKKWKKZ | 79 | 392.9647 | 4 | 29.24 | 1567.834 | -2.7 |
| 6144 | SaeDfQKWKKZ | 79 | 480.2581 | 3 | 29.39 | 1437.756 | -2.2 |
| 6556 | fFKeSDKWKKZ | 79 | 372.2065 | 4 | 31.04 | 1484.797 | 0 |
| 6592 | SNFLGfKWKKZ | 79 | 443.9192 | 3 | 31.19 | 1328.739 | -2.6 |
| 7085 | gTEgfPKWKKZ | 79 | 348.9647 | 4 | 33.16 | 1391.833 | -2.4 |
| 7270 | PfgghPKWKKZ | 79 | 448.9556 | 3 | 33.88 | 1343.848 | -2.5 |
| 7299 | PfgghPKWKKZ | 79 | 448.9557 | 3 | 34 | 1343.848 | -2.2 |
| 7806 | egTQeEKWKKZ | 79 | 489.9422 | 3 | 36.08 | 1466.807 | -1.7 |
| 7829 | egTQeEKWKKZ | 79 | 489.9422 | 3 | 36.17 | 1466.807 | -1.8 |
| 7852 | egTQeEKWKKZ | 79 | 367.7082 | 4 | 36.27 | 1466.807 | -2.4 |
| 7868 | egTQeEKWKKZ | 79 | 367.7081 | 4 | 36.33 | 1466.807 | -2.9 |
| 7897 | egTQeEKWKKZ | 79 | 367.7082 | 4 | 36.45 | 1466.807 | -2.5 |
| 7936 | egTQeEKWKKZ | 79 | 367.7083 | 4 | 36.6 | 1466.807 | -2.4 |
| 7949 | egTQeEKWKKZ | 79 | 367.7082 | 4 | 36.65 | 1466.807 | -2.6 |
| 7962 | egTQeEKWKKZ | 79 | 367.7083 | 4 | 36.7 | 1466.807 | -2.3 |
| 7979 | egTQeEKWKKZ | 79 | 489.9424 | 3 | 36.77 | 1466.807 | -1.3 |
| 7985 | egTQeEKWKKZ | 79 | 367.7085 | 4 | 36.8 | 1466.807 | -1.8 |
| 7992 | egTQeEKWKKZ | 79 | 489.9423 | 3 | 36.82 | 1466.807 | -1.5 |
| 8034 | egTQeEKWKKZ | 79 | 489.942 | 3 | 36.99 | 1466.807 | -2.3 |
| 8047 | egTQeEKWKKZ | 79 | 489.9418 | 3 | 37.04 | 1466.807 | -2.6 |
| 8050 | egTQeEKWKKZ | 79 | 367.7082 | 4 | 37.06 | 1466.807 | -2.6 |
| 8103 | egTQeEKWKKZ | 79 | 489.9421 | 3 | 37.28 | 1466.807 | -2 |
| 8104 | egTQeEKWKKZ | 79 | 489.9421 | 3 | 37.28 | 1466.807 | -2 |
| 8574 | ENHTeGKWKKZ | 79 | 441.8998 | 3 | 39.31 | 1322.688 | -8.1 |
| 8735 | TbTQeEKWKKZ | 79 | 457.5859 | 3 | 40.01 | 1369.751 | -10.9 |

| | | | | | | | |
|---|---|---|---|---|---|---|---|
| 9249 | ESiSfGKKKKZ | 79 | 422.9048 | 3 | 42.22 | 1265.688 | 3.7 |
| 9527 | FffbFfKWKKZ | 79 | 420.233 | 4 | 43.42 | 1676.906 | -1.8 |
| 10033 | aGiEegKWKKZ | 79 | 451.263 | 3 | 45.67 | 1350.756 | 8.2 |
| 10035 | aGiEegKWKKZ | 79 | 451.263 | 3 | 45.68 | 1350.756 | 8.2 |
| 11147 | eLhPLdKWKKZ | 79 | 457.2876 | 3 | 50.56 | 1368.843 | -1.7 |
| 4069 | LQREQeKWKKZ | 78 | 360.7112 | 4 | 20.91 | 1438.82 | -2.8 |
| 4225 | LEFTDTKWKKZ | 78 | 432.2424 | 3 | 21.58 | 1293.708 | -2 |
| 5626 | EgTGAeKWKKZ | 78 | 424.2477 | 3 | 27.27 | 1269.723 | -1.6 |
| 9928 | bSPEegKWKKZ | 78 | 451.2633 | 3 | 45.21 | 1350.781 | -9.8 |
| 9940 | QPhEegKWKKZ | 78 | 451.2632 | 3 | 45.26 | 1350.781 | -10 |
| 3432 | KFSLEaKWKKZ | 78 | 323.9508 | 4 | 18.16 | 1291.776 | -1.6 |
| 3447 | KFSLEaKWKKZ | 78 | 323.9506 | 4 | 18.23 | 1291.776 | -2.3 |
| 3804 | KGEgFdKWKKZ | 78 | 338.9598 | 4 | 19.76 | 1351.813 | -1.9 |
| 3897 | AAKDLFKWKKZ | 78 | 309.1914 | 4 | 20.16 | 1232.739 | -2.1 |
| 4176 | KaTfQLKWKKZ | 78 | 346.2163 | 4 | 21.37 | 1380.839 | -2.3 |
| 4226 | LEFTDTKWKKZ | 78 | 432.2424 | 3 | 21.58 | 1293.708 | -2 |
| 4936 | dAQgfGKWKKZ | 78 | 343.4571 | 4 | 24.49 | 1369.802 | -2.2 |
| 4985 | dAQgfGKWKKZ | 78 | 343.4569 | 4 | 24.68 | 1369.802 | -2.7 |
| 5175 | KgfGTPKWKKZ | 78 | 331.2082 | 4 | 25.45 | 1320.807 | -2.7 |
| 5179 | gKfGTPKWKKZ | 78 | 331.2082 | 4 | 25.46 | 1320.807 | -2.6 |
| 5247 | KgfGTPKWKKZ | 78 | 331.2082 | 4 | 25.73 | 1320.807 | -2.5 |
| 5441 | fFKFNTKWKKZ | 78 | 362.9598 | 4 | 26.51 | 1447.813 | -1.9 |
| 5532 | fNTRfNKWKKZ | 78 | 507.2809 | 3 | 26.88 | 1518.825 | -2.5 |
| 5542 | fNTRfNKWKKZ | 78 | 380.7127 | 4 | 26.92 | 1518.825 | -2 |
| 5571 | fNTRfNKWKKZ | 78 | 380.7126 | 4 | 27.04 | 1518.825 | -2.1 |
| 5647 | PSFfEAKWKKZ | 78 | 448.2476 | 3 | 27.36 | 1341.723 | -1.8 |
| 5788 | fQhhbfKWKKZ | 78 | 365.465 | 4 | 27.92 | 1457.834 | -2 |
| 5824 | fQhhbfKWKKZ | 78 | 365.4651 | 4 | 28.07 | 1457.834 | -1.8 |
| 5874 | fQhhbfKWKKZ | 78 | 365.4648 | 4 | 28.27 | 1457.834 | -2.6 |
| 6104 | SaeDfQKWKKZ | 78 | 480.258 | 3 | 29.23 | 1437.756 | -2.4 |
| 6154 | SaeDfQKWKKZ | 78 | 480.2581 | 3 | 29.43 | 1437.756 | -2.2 |
| 6219 | FSfGgRKWKKZ | 78 | 462.6107 | 3 | 29.69 | 1384.813 | -2.2 |
| 6572 | fFKeSDKWKKZ | 78 | 372.2076 | 4 | 31.1 | 1484.797 | 2.8 |
| 6685 | afNfgAKWKKZ | 78 | 482.9507 | 3 | 31.56 | 1445.834 | -2.2 |
| 6805 | afNfgAKWKKZ | 78 | 482.9505 | 3 | 32.03 | 1445.834 | -2.6 |
| 7052 | gTEgfPKWKKZ | 78 | 348.9648 | 4 | 33.02 | 1391.833 | -2.2 |
| 7140 | gTEgfPKWKKZ | 78 | 348.9646 | 4 | 33.37 | 1391.833 | -2.6 |
| 7153 | gTEgfPKWKKZ | 78 | 348.9646 | 4 | 33.42 | 1391.833 | -2.8 |
| 7319 | PfgghPKWKKZ | 78 | 448.9557 | 3 | 34.08 | 1343.848 | -2.2 |
| 7371 | PfgghPKWKKZ | 78 | 336.9685 | 4 | 34.29 | 1343.848 | -2.6 |
| 7506 | fAfFAGKWKKZ | 78 | 460.9243 | 3 | 34.86 | 1379.754 | -2.2 |
| 7884 | egTQeEKWKKZ | 78 | 367.7082 | 4 | 36.4 | 1466.807 | -2.6 |
| 7910 | egTQeEKWKKZ | 78 | 367.7083 | 4 | 36.5 | 1466.807 | -2.4 |

| | | | | | | | |
|---|---|---|---|---|---|---|---|
| 7923 | egTQeEKWKKZ | 78 | 367.7083 | 4 | 36.55 | 1466.807 | -2.4 |
| 7998 | egTQeEKWKKZ | 78 | 367.7084 | 4 | 36.85 | 1466.807 | -2 |
| 8011 | egTQeEKWKKZ | 78 | 367.7082 | 4 | 36.9 | 1466.807 | -2.4 |
| 8012 | egTQeEKWKKZ | 78 | 367.7082 | 4 | 36.9 | 1466.807 | -2.4 |
| 8037 | egTQeEKWKKZ | 78 | 367.7082 | 4 | 37 | 1466.807 | -2.5 |
| 8066 | egTQeEKWKKZ | 78 | 367.7082 | 4 | 37.12 | 1466.807 | -2.5 |
| 8073 | egTQeEKWKKZ | 78 | 489.942 | 3 | 37.15 | 1466.807 | -2.1 |
| 8552 | NEHTeGKWKKZ | 78 | 441.9 | 3 | 39.22 | 1322.688 | -7.7 |
| 8890 | fegQbFKWKKZ | 78 | 384.9765 | 4 | 40.7 | 1535.881 | -2.4 |
| 9203 | ePLEegKWKKZ | 78 | 362.9661 | 4 | 42.02 | 1447.838 | -1.9 |
| 9324 | ePLEegKWKKZ | 78 | 362.9661 | 4 | 42.54 | 1447.838 | -1.9 |
| 9916 | aGiEegKWKKZ | 78 | 451.2632 | 3 | 45.15 | 1350.756 | 8.5 |
| 9975 | QPhEegKWKKZ | 78 | 451.263 | 3 | 45.42 | 1350.781 | -10.5 |
| 3790 | SRNfhGKWKKZ | 77 | 328.4417 | 4 | 19.7 | 1309.741 | -2.2 |
| 4033 | fTNETSKWKKZ | 77 | 336.6824 | 4 | 20.75 | 1342.703 | -2.1 |
| 5469 | NgPeASKWKKZ | 77 | 427.9195 | 3 | 26.63 | 1280.739 | -2.1 |
| 6349 | bGGFgfKWKKZ | 77 | 442.6001 | 3 | 30.21 | 1324.781 | -1.9 |
| 6476 | hPAGfeKWKKZ | 77 | 440.2523 | 3 | 30.72 | 1317.739 | -2.5 |
| 7046 | GaSLEfKWKKZ | 77 | 433.2525 | 3 | 33 | 1296.734 | 1.2 |
| 8258 | hQhLgQKWKKZ | 77 | 418.935 | 3 | 37.94 | 1253.797 | -11.4 |
| 3813 | GKEgFdKWKKZ | 77 | 338.9597 | 4 | 19.8 | 1351.813 | -2.2 |
| 3873 | GKEgFdKWKKZ | 77 | 338.9599 | 4 | 20.06 | 1351.813 | -1.7 |
| 3924 | AAKDLFKWKKZ | 77 | 309.191 | 4 | 20.28 | 1232.739 | -3.4 |
| 4127 | AKQfFNKWKKZ | 77 | 467.2703 | 3 | 21.16 | 1398.793 | -2.4 |
| 4314 | AhAfgGKWKKZ | 77 | 306.444 | 4 | 21.95 | 1221.739 | 6.8 |
| 4411 | AhAfgGKWKKZ | 77 | 306.444 | 4 | 22.34 | 1221.739 | 6.7 |
| 4525 | GGgRLFKWKKZ | 77 | 415.9353 | 3 | 22.8 | 1244.787 | -2.5 |
| 4561 | GGgRLFKWKKZ | 77 | 415.9354 | 3 | 22.95 | 1244.787 | -2.3 |
| 4857 | TRfQAFKWKKZ | 77 | 354.4575 | 4 | 24.17 | 1413.803 | -1.8 |
| 4915 | dNhgfGKWKKZ | 77 | 343.4569 | 4 | 24.41 | 1369.802 | -2.6 |
| 4923 | dAQgfGKWKKZ | 77 | 343.4571 | 4 | 24.44 | 1369.802 | -2.3 |
| 4946 | dAQgfGKWKKZ | 77 | 343.4573 | 4 | 24.53 | 1369.802 | -1.5 |
| 4956 | dAQgfGKWKKZ | 77 | 343.4573 | 4 | 24.57 | 1369.802 | -1.6 |
| 5003 | NFDSfdKWKKZ | 77 | 363.4454 | 4 | 24.75 | 1449.756 | -2.2 |
| 5192 | gKfGTPKWKKZ | 77 | 331.2083 | 4 | 25.51 | 1320.807 | -2.3 |
| 5234 | gKfGTPKWKKZ | 77 | 331.2084 | 4 | 25.68 | 1320.807 | -2 |
| 5521 | NgPeASKWKKZ | 77 | 427.9197 | 3 | 26.84 | 1280.739 | -1.5 |
| 5889 | fQhhbfKWKKZ | 77 | 365.4648 | 4 | 28.34 | 1457.834 | -2.5 |
| 5979 | DfdDfKKWKKZ | 77 | 392.9649 | 4 | 28.71 | 1567.834 | -2.2 |
| 6002 | DfdDfKKWKKZ | 77 | 392.9647 | 4 | 28.8 | 1567.834 | -2.8 |
| 6022 | DfdDfKKWKKZ | 77 | 392.9651 | 4 | 28.88 | 1567.834 | -1.6 |
| 6077 | DfdDfKKWKKZ | 77 | 392.9649 | 4 | 29.11 | 1567.834 | -2.2 |
| 6095 | SaeDfQKWKKZ | 77 | 360.4456 | 4 | 29.19 | 1437.756 | -1.7 |

| | | | | | | | |
|---|---|---|---|---|---|---|---|
| 6101 | SaeDfQKWKKZ | 77 | 360.4453 | 4 | 29.21 | 1437.756 | -2.4 |
| 6123 | FfQEhLKWKKZ | 77 | 354.2056 | 4 | 29.31 | 1412.797 | -2.6 |
| 6160 | FSfGgRKWKKZ | 77 | 347.2096 | 4 | 29.46 | 1384.813 | -2.9 |
| 6164 | FfQEhLKWKKZ | 77 | 354.2052 | 4 | 29.47 | 1412.797 | -3.7 |
| 6675 | afNfgAKWKKZ | 77 | 482.9508 | 3 | 31.52 | 1445.834 | -1.9 |
| 6815 | afNfgAKWKKZ | 77 | 362.4648 | 4 | 32.07 | 1445.834 | -2.5 |
| 7865 | egTQeEKWKKZ | 77 | 489.9418 | 3 | 36.32 | 1466.807 | -2.6 |
| 8024 | egTQeEKWKKZ | 77 | 367.7084 | 4 | 36.95 | 1466.807 | -2 |
| 8025 | egTQeEKWKKZ | 77 | 367.7084 | 4 | 36.96 | 1466.807 | -2 |
| 8060 | egTQeEKWKKZ | 77 | 489.942 | 3 | 37.1 | 1466.807 | -2.1 |
| 8088 | egTQeEKWKKZ | 77 | 489.9422 | 3 | 37.21 | 1466.807 | -1.8 |
| 8423 | DTALffKWKKZ | 77 | 717.8982 | 2 | 38.66 | 1433.786 | -2.8 |
| 8492 | ASSSggKWKKZ | 77 | 392.2487 | 3 | 38.97 | 1173.723 | 0.6 |
| 8526 | ASSSggKWKKZ | 77 | 392.2489 | 3 | 39.11 | 1173.723 | 1.3 |
| 8564 | NHETeGKWKKZ | 77 | 441.8998 | 3 | 39.27 | 1322.688 | -8 |
| 8866 | fegQbFKWKKZ | 77 | 512.9659 | 3 | 40.59 | 1535.881 | -3 |
| 9186 | ESiSfGKKKKZ | 77 | 422.905 | 3 | 41.95 | 1265.688 | 4.1 |
| 9261 | EiSSfGKKKKZ | 77 | 422.905 | 3 | 42.27 | 1265.688 | 4 |
| 10099 | bSPEegKWKKZ | 77 | 451.263 | 3 | 45.96 | 1350.781 | -10.5 |
| 10313 | QHNfgAKWKKZ | 77 | 463.5998 | 3 | 46.88 | 1387.788 | -7.5 |
| 2968 | RNESHFKWKKZ | 76 | 340.4408 | 4 | 16.12 | 1357.737 | -2 |
| 3292 | QQSPLGKWKKZ | 76 | 400.2391 | 3 | 17.55 | 1197.698 | -2.4 |
| 8361 | NGHNAfKWKKZ | 76 | 435.5683 | 3 | 38.39 | 1303.694 | -8.2 |
| 2956 | hLASNDKWKKZ | 76 | 294.1734 | 4 | 16.07 | 1172.667 | -1.7 |
| 3876 | AAKDLFKWKKZ | 76 | 309.1915 | 4 | 20.07 | 1232.739 | -1.9 |
| 3960 | RKLgKLKWKKZ | 76 | 339.2438 | 4 | 20.43 | 1352.95 | -2.7 |
| 4065 | GGaLgLKWKKZ | 76 | 289.6978 | 4 | 20.89 | 1154.765 | -2.7 |
| 4134 | AKQfFNKWKKZ | 76 | 350.7045 | 4 | 21.19 | 1398.793 | -2.6 |
| 4161 | AKQfFNKWKKZ | 76 | 350.7046 | 4 | 21.3 | 1398.793 | -2.3 |
| 4302 | AhAfgGKWKKZ | 76 | 306.444 | 4 | 21.9 | 1221.739 | 6.8 |
| 4340 | AhAfgGKWKKZ | 76 | 306.444 | 4 | 22.06 | 1221.739 | 6.8 |
| 4366 | AhAfgGKWKKZ | 76 | 306.4441 | 4 | 22.16 | 1221.739 | 7 |
| 4379 | AhAfgGKWKKZ | 76 | 306.4438 | 4 | 22.22 | 1221.739 | 6.2 |
| 4387 | bgRdGfKWKKZ | 76 | 364.2281 | 4 | 22.25 | 1452.887 | -2.6 |
| 4395 | AhAfgGKWKKZ | 76 | 306.4439 | 4 | 22.28 | 1221.739 | 6.4 |
| 4397 | bgRdGfKWKKZ | 76 | 364.2281 | 4 | 22.29 | 1452.887 | -2.6 |
| 4407 | bgRdGfKWKKZ | 76 | 364.2282 | 4 | 22.33 | 1452.887 | -2.3 |
| 4417 | bgRdGfKWKKZ | 76 | 364.2282 | 4 | 22.37 | 1452.887 | -2.1 |
| 4430 | bgRdGfKWKKZ | 76 | 364.2284 | 4 | 22.42 | 1452.887 | -1.6 |
| 4443 | bgRdGfKWKKZ | 76 | 364.2282 | 4 | 22.47 | 1452.887 | -2.3 |
| 4456 | bgRdGfKWKKZ | 76 | 364.2283 | 4 | 22.52 | 1452.887 | -2.1 |
| 4482 | bgRdGfKWKKZ | 76 | 364.2281 | 4 | 22.62 | 1452.887 | -2.6 |
| 4508 | bgRdGfKWKKZ | 76 | 364.2282 | 4 | 22.73 | 1452.887 | -2.2 |

| | | | | | | | |
|---|---|---|---|---|---|---|---|
| 4521 | bgRdGfKWKKZ | 76 | 364.2281 | 4 | 22.78 | 1452.887 | -2.5 |
| 4534 | bgRdGfKWKKZ | 76 | 364.2282 | 4 | 22.83 | 1452.887 | -2.1 |
| 4566 | bgRdGfKWKKZ | 76 | 364.2283 | 4 | 22.97 | 1452.887 | -2.1 |
| 4576 | bgRdGfKWKKZ | 76 | 364.2282 | 4 | 23.01 | 1452.887 | -2.2 |
| 4595 | bgRdGfKWKKZ | 76 | 364.2282 | 4 | 23.09 | 1452.887 | -2.3 |
| 4660 | bgRdGfKWKKZ | 76 | 364.2283 | 4 | 23.36 | 1452.887 | -2 |
| 4749 | EAfaLGKWKKZ | 76 | 427.9193 | 3 | 23.72 | 1280.739 | -2.5 |
| 4752 | EAfaLGKWKKZ | 76 | 427.9192 | 3 | 23.74 | 1280.739 | -2.7 |
| 4767 | EAfaLGKWKKZ | 76 | 427.9195 | 3 | 23.8 | 1280.739 | -2 |
| 4797 | EAfaLGKWKKZ | 76 | 427.9194 | 3 | 23.92 | 1280.739 | -2.2 |
| 4966 | FKeNFKKWKKZ | 76 | 363.2178 | 4 | 24.61 | 1448.845 | -1.6 |
| 4969 | dAQgfGKWKKZ | 76 | 343.4571 | 4 | 24.62 | 1369.802 | -2.3 |
| 5422 | eLhPLdKWKKZ | 76 | 343.2176 | 4 | 26.43 | 1368.843 | -1.6 |
| 5481 | NgPeASKWKKZ | 76 | 427.9196 | 3 | 26.68 | 1280.739 | -1.9 |
| 5494 | NgPeASKWKKZ | 76 | 427.9196 | 3 | 26.73 | 1280.739 | -1.9 |
| 5555 | fNTRfNKWKKZ | 76 | 380.7127 | 4 | 26.98 | 1518.825 | -2 |
| 5838 | fQhhbfKWKKZ | 76 | 486.9506 | 3 | 28.12 | 1457.834 | -2.5 |
| 5866 | fQhhbfKWKKZ | 76 | 486.9503 | 3 | 28.24 | 1457.834 | -3.1 |
| 6012 | DfdDfKKWKKZ | 76 | 392.9652 | 4 | 28.84 | 1567.834 | -1.5 |
| 6051 | DfdDfKKWKKZ | 76 | 392.9648 | 4 | 29 | 1567.834 | -2.5 |
| 6064 | DfdDfKKWKKZ | 76 | 392.9647 | 4 | 29.06 | 1567.834 | -2.7 |
| 6092 | DfdDfKKWKKZ | 76 | 392.9651 | 4 | 29.18 | 1567.834 | -1.8 |
| 6125 | SaeDfQKWKKZ | 76 | 360.445 | 4 | 29.31 | 1437.756 | -3.3 |
| 6173 | FSfGgRKWKKZ | 76 | 347.2097 | 4 | 29.51 | 1384.813 | -2.5 |
| 6186 | FSfGgRKWKKZ | 76 | 347.2097 | 4 | 29.56 | 1384.813 | -2.5 |
| 6212 | FSfGgRKWKKZ | 76 | 347.2097 | 4 | 29.67 | 1384.813 | -2.4 |
| 6359 | bGGFgfKWKKZ | 76 | 332.2018 | 4 | 30.26 | 1324.781 | -2.3 |
| 6512 | hPAGfeKWKKZ | 76 | 440.2526 | 3 | 30.87 | 1317.739 | -2 |
| 6659 | afNfgAKWKKZ | 76 | 362.4648 | 4 | 31.46 | 1445.834 | -2.3 |
| 6740 | afNfgAKWKKZ | 76 | 482.9507 | 3 | 31.78 | 1445.834 | -2.3 |
| 6776 | afNfgAKWKKZ | 76 | 362.4648 | 4 | 31.92 | 1445.834 | -2.4 |
| 7213 | PfgghPKWKKZ | 76 | 336.9685 | 4 | 33.66 | 1343.848 | -2.5 |
| 7335 | PfgghPKWKKZ | 76 | 448.9557 | 3 | 34.14 | 1343.848 | -2.4 |
| 8021 | egTQeEKWKKZ | 76 | 489.9423 | 3 | 36.94 | 1466.807 | -1.5 |
| 8475 | DTALffKWKKZ | 76 | 717.8991 | 2 | 38.9 | 1433.786 | -1.5 |
| 8477 | DTALffKWKKZ | 76 | 717.8991 | 2 | 38.9 | 1433.786 | -1.5 |
| 8784 | fegQbFKWKKZ | 76 | 512.9662 | 3 | 40.23 | 1535.881 | -2.4 |
| 8785 | fegQbFKWKKZ | 76 | 512.9662 | 3 | 40.24 | 1535.881 | -2.4 |
| 8842 | fegQbFKWKKZ | 76 | 512.9663 | 3 | 40.48 | 1535.881 | -2.3 |
| 8854 | fegQbFKWKKZ | 76 | 512.9662 | 3 | 40.54 | 1535.881 | -2.5 |
| 8875 | fegQbFKWKKZ | 76 | 512.9663 | 3 | 40.63 | 1535.881 | -2.3 |
| 8899 | fegQbFKWKKZ | 76 | 512.9667 | 3 | 40.74 | 1535.881 | -1.5 |
| 9443 | FffbFfKWKKZ | 76 | 420.233 | 4 | 43.06 | 1676.906 | -1.8 |

| | | | | | | | |
|---|---|---|---|---|---|---|---|
| 9600 | AfLEegKWKKZ | 76 | 483.6192 | 3 | 43.75 | 1447.838 | -1.5 |
| 9601 | AfLEegKWKKZ | 76 | 483.6192 | 3 | 43.75 | 1447.838 | -1.5 |
| 9646 | AfLEegKWKKZ | 76 | 483.6187 | 3 | 43.95 | 1447.838 | -2.5 |
| 9963 | bSPEegKWKKZ | 76 | 451.2633 | 3 | 45.36 | 1350.781 | -9.7 |
| 9987 | bSPEegKWKKZ | 76 | 451.2633 | 3 | 45.47 | 1350.781 | -9.8 |
| 10011 | bSPEegKWKKZ | 76 | 451.2632 | 3 | 45.57 | 1350.781 | -9.9 |
| 10047 | bSPEegKWKKZ | 76 | 451.2632 | 3 | 45.73 | 1350.781 | -10 |
| 10086 | bSPEegKWKKZ | 76 | 451.263 | 3 | 45.9 | 1350.781 | -10.4 |
| 10158 | NHNfgAKWKKZ | 76 | 458.928 | 3 | 46.23 | 1373.772 | -7 |
| 5643 | AGAgfGKWKKZ | 75 | 398.9089 | 3 | 27.34 | 1193.707 | -2.1 |
| 6071 | bHDFRDKWKKZ | 75 | 462.2582 | 3 | 29.09 | 1383.752 | 0.3 |
| 2848 | AhLSQQKWKKZ | 75 | 300.9352 | 4 | 15.6 | 1199.714 | -1.9 |
| 3146 | SFEDGGKWKKZ | 75 | 394.2074 | 3 | 16.9 | 1179.604 | -2.7 |
| 4034 | fTNETSKWKKZ | 75 | 336.6824 | 4 | 20.75 | 1342.703 | -2.1 |
| 4104 | AKQfFNKWKKZ | 75 | 350.7045 | 4 | 21.06 | 1398.793 | -2.6 |
| 4136 | AKQfFNKWKKZ | 75 | 467.2702 | 3 | 21.19 | 1398.793 | -2.7 |
| 4151 | AKQfFNKWKKZ | 75 | 467.2706 | 3 | 21.26 | 1398.793 | -1.7 |
| 4172 | AgTLGNKWKKZ | 75 | 391.2475 | 3 | 21.35 | 1170.724 | -2.5 |
| 4184 | AgTLGNKWKKZ | 75 | 391.2477 | 3 | 21.4 | 1170.724 | -2 |
| 4196 | AgTLGNKWKKZ | 75 | 391.2474 | 3 | 21.45 | 1170.724 | -2.8 |
| 4247 | KfGQdLKWKKZ | 75 | 354.2176 | 4 | 21.67 | 1412.845 | -2.2 |
| 4353 | AhAfgGKWKKZ | 75 | 306.4441 | 4 | 22.11 | 1221.739 | 7 |
| 4469 | bgRdGfKWKKZ | 75 | 364.2281 | 4 | 22.57 | 1452.887 | -2.4 |
| 4495 | bgRdGfKWKKZ | 75 | 364.2281 | 4 | 22.68 | 1452.887 | -2.5 |
| 4547 | bgRdGfKWKKZ | 75 | 364.2283 | 4 | 22.89 | 1452.887 | -2.1 |
| 4593 | GGgRLFKWKKZ | 75 | 415.9353 | 3 | 23.08 | 1244.787 | -2.5 |
| 4782 | EAfaLGKWKKZ | 75 | 427.9194 | 3 | 23.86 | 1280.739 | -2.2 |
| 4993 | begDTbKWKKZ | 75 | 345.9583 | 4 | 24.71 | 1379.808 | -2.9 |
| 5127 | KTfhKfKWKKZ | 75 | 369.9763 | 4 | 25.26 | 1475.881 | -2.9 |
| 5690 | ffKHgTKWKKZ | 75 | 382.7295 | 4 | 27.53 | 1526.891 | -1.7 |
| 5697 | ffKHgTKWKKZ | 75 | 382.7293 | 4 | 27.56 | 1526.891 | -2 |
| 5831 | fQLGbfKWKKZ | 75 | 486.9511 | 3 | 28.1 | 1457.834 | -1.3 |
| 5868 | fQhhbfKWKKZ | 75 | 486.9503 | 3 | 28.25 | 1457.834 | -3.1 |
| 6038 | DfdDfKKWKKZ | 75 | 392.9648 | 4 | 28.95 | 1567.834 | -2.6 |
| 6110 | SaeDfQKWKKZ | 75 | 360.4451 | 4 | 29.25 | 1437.756 | -2.9 |
| 6131 | SaeDfQKWKKZ | 75 | 480.258 | 3 | 29.34 | 1437.756 | -2.3 |
| 6136 | FfQEhLKWKKZ | 75 | 354.2056 | 4 | 29.36 | 1412.797 | -2.5 |
| 6141 | SaeDfQKWKKZ | 75 | 360.4452 | 4 | 29.38 | 1437.756 | -2.7 |
| 6149 | FfQEhLKWKKZ | 75 | 354.2056 | 4 | 29.41 | 1412.797 | -2.5 |
| 6374 | eTFLGEKWKKZ | 75 | 444.9195 | 3 | 30.31 | 1331.739 | -1.7 |
| 6499 | hPAGfeKWKKZ | 75 | 440.2524 | 3 | 30.81 | 1317.739 | -2.4 |
| 6799 | afNfgAKWKKZ | 75 | 362.4647 | 4 | 32.01 | 1445.834 | -2.7 |
| 6893 | afNfLhKWKKZ | 75 | 362.4649 | 4 | 32.38 | 1445.834 | -2.1 |

| | | | | | | | |
|---|---|---|---|---|---|---|---|
| 7036 | GaSLEfKWKKZ | 75 | 433.2524 | 3 | 32.96 | 1296.734 | 1 |
| 7062 | GaSLEfKWKKZ | 75 | 433.2523 | 3 | 33.06 | 1296.734 | 0.6 |
| 7075 | GaSLEfKWKKZ | 75 | 433.2522 | 3 | 33.12 | 1296.734 | 0.5 |
| 7091 | GaSLEfKWKKZ | 75 | 433.2524 | 3 | 33.18 | 1296.734 | 1 |
| 8531 | NEHTeGKWKKZ | 75 | 441.8999 | 3 | 39.13 | 1322.688 | -7.8 |
| 8588 | NEHTeGKWKKZ | 75 | 441.8998 | 3 | 39.37 | 1322.688 | -8.1 |
| 8758 | fegQbFKWKKZ | 75 | 512.9662 | 3 | 40.11 | 1535.881 | -2.4 |
| 8794 | fegQbFKWKKZ | 75 | 384.9766 | 4 | 40.28 | 1535.881 | -2.3 |
| 8796 | fegQbFKWKKZ | 75 | 512.9664 | 3 | 40.29 | 1535.881 | -2.1 |
| 8797 | fegQbFKWKKZ | 75 | 512.9664 | 3 | 40.29 | 1535.881 | -2.1 |
| 8803 | fegQbFKWKKZ | 75 | 512.9664 | 3 | 40.31 | 1535.881 | -2.1 |
| 8818 | fegQbFKWKKZ | 75 | 512.9662 | 3 | 40.38 | 1535.881 | -2.4 |
| 8887 | fegQbFKWKKZ | 75 | 512.9662 | 3 | 40.68 | 1535.881 | -2.5 |
| 8914 | fegQbFKWKKZ | 75 | 512.9667 | 3 | 40.8 | 1535.881 | -1.6 |
| 9460 | fALEegKWKKZ | 75 | 483.6194 | 3 | 43.13 | 1447.838 | -1 |
| 9574 | AfLEegKWKKZ | 75 | 483.6191 | 3 | 43.63 | 1447.838 | -1.6 |
| 9894 | GfLfGKWWKKZ | 75 | 483.2721 | 3 | 45.05 | 1446.796 | -1.4 |
| 9951 | bSPEegKWKKZ | 75 | 451.2633 | 3 | 45.31 | 1350.781 | -9.7 |
| 9999 | bSPEegKWKKZ | 75 | 451.2633 | 3 | 45.52 | 1350.781 | -9.7 |
| 10023 | bSPEegKWKKZ | 75 | 451.2632 | 3 | 45.62 | 1350.781 | -9.9 |
| 10101 | bSPEegKWKKZ | 75 | 451.263 | 3 | 45.97 | 1350.781 | -10.5 |
| 10122 | NHNfgAKWKKZ | 75 | 458.928 | 3 | 46.07 | 1373.772 | -7 |
| 10277 | QHNfgAKWKKZ | 75 | 463.5999 | 3 | 46.73 | 1387.788 | -7.1 |
| 10301 | QHNfgAKWKKZ | 75 | 463.6002 | 3 | 46.83 | 1387.788 | -6.6 |
| 3395 | PGNgGNKWKKZ | 74 | 289.4247 | 4 | 18 | 1153.672 | -2.3 |
| 5052 | bGHedaKWKKZ | 74 | 461.2683 | 3 | 24.95 | 1380.793 | -7.2 |
| 7370 | GGieDfKWKKZ | 74 | 348.4282 | 4 | 34.29 | 1389.698 | -10.6 |
| 7387 | PfgghPKWKKZ | 74 | 336.9686 | 4 | 34.36 | 1343.848 | -2.2 |
| 8276 | TAbLgQKWKKZ | 74 | 418.9352 | 3 | 38.02 | 1253.797 | -10.7 |
| 8388 | NHGANfKWKKZ | 74 | 435.5684 | 3 | 38.51 | 1303.694 | -8 |
| 10809 | TFDfgAKWKKZ | 74 | 458.2685 | 3 | 49.07 | 1371.77 | 9.7 |
| 10830 | EHPfgAKWKKZ | 74 | 458.2681 | 3 | 49.16 | 1371.782 | 0.8 |
| 2898 | hLASNDKWKKZ | 74 | 391.8955 | 3 | 15.82 | 1172.667 | -1.6 |
| 2943 | LAhSNDKWKKZ | 74 | 294.1733 | 4 | 16.01 | 1172.667 | -2 |
| 4236 | NhEKLeKWKKZ | 74 | 339.4546 | 4 | 21.62 | 1353.792 | -1.9 |
| 4237 | KfGQdLKWKKZ | 74 | 354.2176 | 4 | 21.63 | 1412.845 | -2.3 |
| 4238 | KfGQdLKWKKZ | 74 | 354.2176 | 4 | 21.63 | 1412.845 | -2.3 |
| 4290 | hAAfgGKWKKZ | 74 | 306.4441 | 4 | 21.85 | 1221.739 | 7 |
| 4424 | AhAfgGKWKKZ | 74 | 306.4441 | 4 | 22.4 | 1221.739 | 6.9 |
| 4828 | TRfQAFKWKKZ | 74 | 354.4572 | 4 | 24.05 | 1413.803 | -2.5 |
| 4976 | PKLDFLKWKKZ | 74 | 326.207 | 4 | 24.65 | 1300.802 | -2.5 |
| 4983 | PKLDFLKWKKZ | 74 | 326.2068 | 4 | 24.67 | 1300.802 | -2.9 |
| 4996 | KPLDFLKWKKZ | 74 | 326.207 | 4 | 24.72 | 1300.802 | -2.4 |

| | | | | | | | |
|---|---|---|---|---|---|---|---|
| 5006 | begDTbKWKKZ | 74 | 345.9584 | 4 | 24.76 | 1379.808 | -2.4 |
| 5038 | PKLDFLKWKKZ | 74 | 326.2072 | 4 | 24.89 | 1300.802 | -1.9 |
| 5075 | KTfhKfKWKKZ | 74 | 369.9769 | 4 | 25.05 | 1475.881 | -1.5 |
| 5114 | KTfhKfKWKKZ | 74 | 369.9766 | 4 | 25.2 | 1475.881 | -2.3 |
| 5169 | KTfhKfKWKKZ | 74 | 369.9765 | 4 | 25.42 | 1475.881 | -2.6 |
| 5519 | fNTRfNKWKKZ | 74 | 507.2813 | 3 | 26.83 | 1518.825 | -1.7 |
| 5600 | AGAgfGKWKKZ | 74 | 398.9088 | 3 | 27.17 | 1193.707 | -2.4 |
| 5833 | fQLGbfKWKKZ | 74 | 486.9511 | 3 | 28.1 | 1457.834 | -1.3 |
| 5840 | fQhhbfKWKKZ | 74 | 486.9506 | 3 | 28.13 | 1457.834 | -2.5 |
| 5853 | fQhhbfKWKKZ | 74 | 486.9512 | 3 | 28.19 | 1457.834 | -1.3 |
| 6229 | FSfGgRKWKKZ | 74 | 347.2098 | 4 | 29.73 | 1384.813 | -2.2 |
| 6390 | FdDDfFKWKKZ | 74 | 378.7004 | 4 | 30.38 | 1510.776 | -2.5 |
| 6669 | afNfgAKWKKZ | 74 | 362.4649 | 4 | 31.5 | 1445.834 | -2.1 |
| 6763 | afNfgAKWKKZ | 74 | 362.4647 | 4 | 31.87 | 1445.834 | -2.6 |
| 6789 | afNfgAKWKKZ | 74 | 362.4648 | 4 | 31.97 | 1445.834 | -2.4 |
| 6841 | afNfLhKWKKZ | 74 | 362.4648 | 4 | 32.18 | 1445.834 | -2.2 |
| 6903 | afNfgAKWKKZ | 74 | 362.4647 | 4 | 32.42 | 1445.834 | -2.6 |
| 6974 | afNfLhKWKKZ | 74 | 362.465 | 4 | 32.71 | 1445.834 | -1.8 |
| 7301 | PfgghPKWKKZ | 74 | 336.9685 | 4 | 34.01 | 1343.848 | -2.6 |
| 8127 | egTQeEKWKKZ | 74 | 489.9424 | 3 | 37.38 | 1466.807 | -1.4 |
| 8128 | egTQeEKWKKZ | 74 | 489.9424 | 3 | 37.38 | 1466.807 | -1.4 |
| 8187 | egTQeEKWKKZ | 74 | 367.7083 | 4 | 37.63 | 1466.807 | -2.3 |
| 8293 | PLLEegKWKKZ | 74 | 455.6191 | 3 | 38.1 | 1363.838 | -1.8 |
| 8337 | NGHNAfKWKKZ | 74 | 435.5683 | 3 | 38.29 | 1303.694 | -8.3 |
| 8363 | NGHNAfKWKKZ | 74 | 435.5683 | 3 | 38.4 | 1303.694 | -8.2 |
| 8467 | EgTeQeKWKKZ | 74 | 367.7081 | 4 | 38.86 | 1466.807 | -2.8 |
| 8528 | ASSSggKWKKZ | 74 | 392.2489 | 3 | 39.12 | 1173.723 | 1.3 |
| 8770 | fegQbFKWKKZ | 74 | 512.9662 | 3 | 40.17 | 1535.881 | -2.4 |
| 9220 | fALEegKWKKZ | 74 | 483.6188 | 3 | 42.09 | 1447.838 | -2.3 |
| 9403 | AfLEegKWKKZ | 74 | 483.6191 | 3 | 42.88 | 1447.838 | -1.6 |
| 9588 | AfLEegKWKKZ | 74 | 483.6191 | 3 | 43.69 | 1447.838 | -1.6 |
| 9589 | AfLEegKWKKZ | 74 | 483.6191 | 3 | 43.7 | 1447.838 | -1.6 |
| 9624 | fALEegKWKKZ | 74 | 483.6191 | 3 | 43.85 | 1447.838 | -1.6 |
| 9625 | fALEegKWKKZ | 74 | 483.6191 | 3 | 43.85 | 1447.838 | -1.6 |
| 10059 | bSPEegKWKKZ | 74 | 451.2629 | 3 | 45.78 | 1350.781 | -10.6 |
| 10134 | NHNfgAKWKKZ | 74 | 458.9277 | 3 | 46.12 | 1373.772 | -7.7 |
| 10143 | NHNfgAKWKKZ | 74 | 458.9277 | 3 | 46.16 | 1373.772 | -7.8 |
| 10289 | QHNfgAKWKKZ | 74 | 463.5997 | 3 | 46.78 | 1387.788 | -7.5 |
| 3780 | gPLShAKWKKZ | 73 | 390.2599 | 3 | 19.65 | 1167.749 | 7.3 |
| 4941 | fTAhfNKWKKZ | 73 | 469.2668 | 3 | 24.51 | 1404.771 | 5.6 |
| 9768 | hfFgSfKWKKZ | 73 | 494.2876 | 3 | 44.5 | 1479.843 | -1.6 |
| 10818 | HPEfgAKWKKZ | 73 | 458.2684 | 3 | 49.11 | 1371.782 | 1.4 |
| 3409 | hFEEADKWKKZ | 73 | 316.9222 | 4 | 18.06 | 1263.661 | -1.1 |

| | | | | | | | |
|---|---|---|---|---|---|---|---|
| 4122 | AKQfFNKWKKZ | 73 | 350.7045 | 4 | 21.14 | 1398.793 | -2.5 |
| 4146 | AKQfFNKWKKZ | 73 | 350.7047 | 4 | 21.24 | 1398.793 | -2.1 |
| 4148 | AgTLGNKWKKZ | 73 | 293.6877 | 4 | 21.25 | 1170.724 | -1.8 |
| 4181 | AgTLGNKWKKZ | 73 | 293.6876 | 4 | 21.39 | 1170.724 | -2 |
| 4213 | LhTLGNKWKKZ | 73 | 293.6875 | 4 | 21.53 | 1170.724 | -2.3 |
| 4214 | LhTLGNKWKKZ | 73 | 293.6875 | 4 | 21.53 | 1170.724 | -2.3 |
| 4608 | bgRdGfKWKKZ | 73 | 364.2281 | 4 | 23.14 | 1452.887 | -2.6 |
| 4795 | TRfQAFKWKKZ | 73 | 354.4572 | 4 | 23.91 | 1413.803 | -2.4 |
| 4802 | TRfQAFKWKKZ | 73 | 354.4572 | 4 | 23.94 | 1413.803 | -2.4 |
| 4815 | TRfQAFKWKKZ | 73 | 354.4574 | 4 | 24 | 1413.803 | -1.8 |
| 4824 | TRfQAFKWKKZ | 73 | 472.2743 | 3 | 24.03 | 1413.803 | -1.5 |
| 4837 | TRfhNFKWKKZ | 73 | 472.2737 | 3 | 24.09 | 1413.804 | -2.9 |
| 4841 | TRfQAFKWKKZ | 73 | 354.4574 | 4 | 24.1 | 1413.803 | -1.9 |
| 4863 | TRfQAFKWKKZ | 73 | 472.2744 | 3 | 24.19 | 1413.803 | -1.3 |
| 4979 | FKeNFKKWKKZ | 73 | 363.2176 | 4 | 24.66 | 1448.845 | -2.3 |
| 5009 | PKLDFLKWKKZ | 73 | 326.207 | 4 | 24.78 | 1300.802 | -2.4 |
| 5022 | PKLDFLKWKKZ | 73 | 326.2069 | 4 | 24.83 | 1300.802 | -2.6 |
| 5042 | bGHedaKWKKZ | 73 | 461.2685 | 3 | 24.91 | 1380.793 | -6.8 |
| 5088 | KTfhKfKWKKZ | 73 | 369.9765 | 4 | 25.1 | 1475.881 | -2.4 |
| 5187 | gaFQRfKWKKZ | 73 | 368.2266 | 4 | 25.49 | 1468.882 | -3.1 |
| 5652 | hfFgSbKWKKZ | 73 | 346.7124 | 4 | 27.38 | 1382.823 | -1.7 |
| 5688 | fgAGbFKWKKZ | 73 | 335.7057 | 4 | 27.52 | 1338.797 | -2 |
| 5701 | fgAGbFKWKKZ | 73 | 335.7059 | 4 | 27.58 | 1338.797 | -1.7 |
| 5855 | fQhhbfKWKKZ | 73 | 486.9512 | 3 | 28.19 | 1457.834 | -1.3 |
| 5989 | DfdDfKKWKKZ | 73 | 392.9648 | 4 | 28.75 | 1567.834 | -2.5 |
| 6107 | DfdDfKKWKKZ | 73 | 392.9647 | 4 | 29.24 | 1567.834 | -2.7 |
| 6199 | FSfGgRKWKKZ | 73 | 347.2097 | 4 | 29.62 | 1384.813 | -2.7 |
| 6527 | eAFfdEKWKKZ | 73 | 511.2785 | 3 | 30.92 | 1530.817 | -2.4 |
| 6582 | AeFfdEKWKKZ | 73 | 511.2785 | 3 | 31.14 | 1530.817 | -2.5 |
| 6594 | fGLHfGKWKKZ | 73 | 350.4503 | 4 | 31.2 | 1397.776 | -2.8 |
| 6633 | afNfgAKWKKZ | 73 | 362.4647 | 4 | 31.35 | 1445.834 | -2.7 |
| 6646 | afNfgAKWKKZ | 73 | 362.4648 | 4 | 31.41 | 1445.834 | -2.4 |
| 6698 | afNfgAKWKKZ | 73 | 482.9506 | 3 | 31.61 | 1445.834 | -2.5 |
| 6705 | afNfLhKWKKZ | 73 | 362.4647 | 4 | 31.64 | 1445.834 | -2.6 |
| 6737 | afNfLhKWKKZ | 73 | 362.4647 | 4 | 31.77 | 1445.834 | -2.6 |
| 6750 | afNfgAKWKKZ | 73 | 362.4648 | 4 | 31.82 | 1445.834 | -2.4 |
| 6870 | afNfgAKWKKZ | 73 | 362.4648 | 4 | 32.29 | 1445.834 | -2.4 |
| 6883 | afNfgAKWKKZ | 73 | 362.4648 | 4 | 32.34 | 1445.834 | -2.3 |
| 6929 | afNfgAKWKKZ | 73 | 362.4647 | 4 | 32.53 | 1445.834 | -2.7 |
| 6942 | afNfLhKWKKZ | 73 | 362.4648 | 4 | 32.58 | 1445.834 | -2.4 |
| 6955 | afNfgAKWKKZ | 73 | 362.4648 | 4 | 32.63 | 1445.834 | -2.5 |
| 6984 | GaSLEfKWKKZ | 73 | 433.2524 | 3 | 32.75 | 1296.734 | 0.9 |
| 7020 | GaSLEfKWKKZ | 73 | 433.2522 | 3 | 32.9 | 1296.734 | 0.5 |

| | | | | | | | |
|---|---|---|---|---|---|---|---|
| 7223 | PfgghPKWKKZ | 73 | 336.9685 | 4 | 33.7 | 1343.848 | -2.6 |
| 7327 | PfgghPKWKKZ | 73 | 336.9686 | 4 | 34.11 | 1343.848 | -2.2 |
| 7376 | GGieDfKWKKZ | 73 | 464.2352 | 3 | 34.31 | 1389.698 | -10.5 |
| 8188 | egTQeEKWKKZ | 73 | 367.7083 | 4 | 37.63 | 1466.807 | -2.3 |
| 8259 | egTQeEKWKKZ | 73 | 489.9419 | 3 | 37.95 | 1466.807 | -2.5 |
| 8260 | egTQeEKWKKZ | 73 | 489.9419 | 3 | 37.95 | 1466.807 | -2.5 |
| 8270 | bShLgQKWKKZ | 73 | 418.9351 | 3 | 38 | 1253.797 | -11.1 |
| 8339 | NGHNAfKWKKZ | 73 | 435.5683 | 3 | 38.3 | 1303.694 | -8.3 |
| 8349 | NGHNAfKWKKZ | 73 | 435.5683 | 3 | 38.34 | 1303.694 | -8.3 |
| 8373 | NGHNAfKWKKZ | 73 | 435.5683 | 3 | 38.45 | 1303.694 | -8.2 |
| 8400 | NHGANfKWKKZ | 73 | 435.5687 | 3 | 38.56 | 1303.694 | -7.4 |
| 8519 | ASSSggKWKKZ | 73 | 392.2486 | 3 | 39.08 | 1173.723 | 0.4 |
| 8576 | NEHTeGKWKKZ | 73 | 441.8998 | 3 | 39.32 | 1322.688 | -8.1 |
| 8604 | DhSLffKWKKZ | 73 | 478.9348 | 3 | 39.44 | 1433.786 | -2.4 |
| 8724 | DhSLffKWKKZ | 73 | 478.9347 | 3 | 39.96 | 1433.786 | -2.5 |
| 8780 | FFNeAeKWKKZ | 73 | 366.2002 | 4 | 40.21 | 1460.776 | -2.7 |
| 9151 | FFLEegKWKKZ | 73 | 483.6188 | 3 | 41.8 | 1447.838 | -2.3 |
| 9160 | AfLEegKWKKZ | 73 | 483.6191 | 3 | 41.83 | 1447.838 | -1.6 |
| 9172 | AfLEegKWKKZ | 73 | 483.6188 | 3 | 41.89 | 1447.838 | -2.2 |
| 9184 | AfLEegKWKKZ | 73 | 483.6188 | 3 | 41.94 | 1447.838 | -2.3 |
| 9196 | AfLEegKWKKZ | 73 | 483.619 | 3 | 41.99 | 1447.838 | -1.9 |
| 9208 | AfLEegKWKKZ | 73 | 483.6188 | 3 | 42.04 | 1447.838 | -2.3 |
| 9232 | AfLEegKWKKZ | 73 | 483.619 | 3 | 42.15 | 1447.838 | -1.8 |
| 9244 | AfLEegKWKKZ | 73 | 483.6187 | 3 | 42.2 | 1447.838 | -2.5 |
| 9256 | AfLEegKWKKZ | 73 | 483.6188 | 3 | 42.25 | 1447.838 | -2.3 |
| 9268 | AfLEegKWKKZ | 73 | 483.6192 | 3 | 42.3 | 1447.838 | -1.5 |
| 9280 | AfLEegKWKKZ | 73 | 483.6191 | 3 | 42.35 | 1447.838 | -1.6 |
| 9292 | AfLEegKWKKZ | 73 | 483.6191 | 3 | 42.4 | 1447.838 | -1.6 |
| 9304 | AfLEegKWKKZ | 73 | 483.619 | 3 | 42.45 | 1447.838 | -2 |
| 9316 | AfLEegKWKKZ | 73 | 483.6192 | 3 | 42.51 | 1447.838 | -1.5 |
| 9331 | fALEegKWKKZ | 73 | 483.6191 | 3 | 42.57 | 1447.838 | -1.6 |
| 9340 | AfLEegKWKKZ | 73 | 483.6191 | 3 | 42.61 | 1447.838 | -1.7 |
| 9352 | AfLEegKWKKZ | 73 | 483.619 | 3 | 42.66 | 1447.838 | -1.8 |
| 9364 | AfLEegKWKKZ | 73 | 483.6191 | 3 | 42.71 | 1447.838 | -1.7 |
| 9376 | fALEegKWKKZ | 73 | 483.619 | 3 | 42.76 | 1447.838 | -1.8 |
| 9388 | AfLEegKWKKZ | 73 | 483.6191 | 3 | 42.82 | 1447.838 | -1.7 |
| 9412 | AfLEegKWKKZ | 73 | 483.619 | 3 | 42.92 | 1447.838 | -1.8 |
| 9436 | AfLEegKWKKZ | 73 | 483.6191 | 3 | 43.02 | 1447.838 | -1.7 |
| 9472 | AfLEegKWKKZ | 73 | 483.6192 | 3 | 43.18 | 1447.838 | -1.5 |
| 9484 | AfLEegKWKKZ | 73 | 483.6192 | 3 | 43.23 | 1447.838 | -1.4 |
| 9502 | AfLEegKWKKZ | 73 | 483.6192 | 3 | 43.31 | 1447.838 | -1.5 |
| 9514 | FFLEegKWKKZ | 73 | 483.6193 | 3 | 43.36 | 1447.838 | -1.3 |
| 9523 | AfLEegKWKKZ | 73 | 483.6192 | 3 | 43.4 | 1447.838 | -1.5 |

| | | | | | | | |
|---|---|---|---|---|---|---|---|
| 9535 | FFLEegKWKKZ | 73 | 483.6192 | 3 | 43.46 | 1447.838 | -1.5 |
| 9550 | AfLEegKWKKZ | 73 | 483.6192 | 3 | 43.52 | 1447.838 | -1.5 |
| 9562 | AfLEegKWKKZ | 73 | 483.6192 | 3 | 43.58 | 1447.838 | -1.4 |
| 9610 | AfLEegKWKKZ | 73 | 483.6188 | 3 | 43.79 | 1447.838 | -2.2 |
| 10071 | QhPEegKWKKZ | 73 | 451.2632 | 3 | 45.83 | 1350.781 | -9.9 |
| 10110 | NHNfgAKWKKZ | 73 | 458.9279 | 3 | 46.01 | 1373.772 | -7.2 |
| 10811 | TFDfgAKWKKZ | 73 | 458.2685 | 3 | 49.08 | 1371.77 | 9.7 |
| 4684 | dhAhgfKWKKZ | 72 | 452.619 | 3 | 23.46 | 1354.828 | 5.3 |
| 5149 | LhLgAGKWKKZ | 72 | 385.5963 | 3 | 25.34 | 1153.77 | -2.6 |
| 10005 | HGALfGKWKKZ | 72 | 416.2419 | 3 | 45.55 | 1245.713 | -7.7 |
| 10038 | GAHLfGKWKKZ | 72 | 416.2417 | 3 | 45.69 | 1245.713 | -8 |
| 3116 | SFEDGGKWKKZ | 72 | 394.2078 | 3 | 16.77 | 1179.604 | -1.7 |
| 3128 | SFEDGGKWKKZ | 72 | 394.2075 | 3 | 16.82 | 1179.604 | -2.4 |
| 3140 | SFEDGGKWKKZ | 72 | 394.2076 | 3 | 16.87 | 1179.604 | -2.2 |
| 3400 | hFEEADKWKKZ | 72 | 316.9218 | 4 | 18.02 | 1263.661 | -2.5 |
| 3433 | KFSLEaKWKKZ | 72 | 323.9508 | 4 | 18.17 | 1291.776 | -1.6 |
| 4157 | AgTLGNKWKKZ | 72 | 293.6875 | 4 | 21.28 | 1170.724 | -2.2 |
| 4193 | AgTLGNKWKKZ | 72 | 293.6874 | 4 | 21.44 | 1170.724 | -2.5 |
| 5055 | bHGedaKWKKZ | 72 | 346.203 | 4 | 24.96 | 1380.793 | -7.2 |
| 5182 | LhLgAGKWKKZ | 72 | 385.5963 | 3 | 25.47 | 1153.77 | -2.6 |
| 5252 | gFgNTQKWKKZ | 72 | 444.942 | 3 | 25.75 | 1331.808 | -2.9 |
| 5259 | gFgNTQKWKKZ | 72 | 444.9422 | 3 | 25.78 | 1331.808 | -2.3 |
| 5269 | gFgNTQKWKKZ | 72 | 444.942 | 3 | 25.82 | 1331.808 | -2.7 |
| 5295 | gFgNTQKWKKZ | 72 | 444.9422 | 3 | 25.92 | 1331.808 | -2.4 |
| 5678 | fgGAbFKWKKZ | 72 | 335.7057 | 4 | 27.48 | 1338.797 | -2.2 |
| 6228 | FSfGgRKWKKZ | 72 | 347.2098 | 4 | 29.73 | 1384.813 | -2.2 |
| 6642 | TNFLGfKWKKZ | 72 | 336.6952 | 4 | 31.39 | 1342.755 | -2.4 |
| 6919 | afNfgAKWKKZ | 72 | 362.465 | 4 | 32.49 | 1445.834 | -1.9 |
| 7010 | afNfLhKWKKZ | 72 | 362.4645 | 4 | 32.85 | 1445.834 | -3.2 |
| 7026 | afNfgAKWKKZ | 72 | 362.4648 | 4 | 32.92 | 1445.834 | -2.5 |
| 7275 | PfgghPKWKKZ | 72 | 336.9686 | 4 | 33.9 | 1343.848 | -2.4 |
| 7288 | PfgghPKWKKZ | 72 | 336.9686 | 4 | 33.95 | 1343.848 | -2.2 |
| 8235 | egTQeEKWKKZ | 72 | 489.9422 | 3 | 37.84 | 1466.807 | -1.8 |
| 8236 | egTQeEKWKKZ | 72 | 489.9422 | 3 | 37.84 | 1466.807 | -1.8 |
| 8305 | LPLEegKWKKZ | 72 | 455.619 | 3 | 38.15 | 1363.838 | -2 |
| 8351 | NGHNAfKWKKZ | 72 | 435.5683 | 3 | 38.35 | 1303.694 | -8.3 |
| 8390 | NHGANfKWKKZ | 72 | 435.5684 | 3 | 38.52 | 1303.694 | -8 |
| 8707 | DhSLffKWKKZ | 72 | 478.935 | 3 | 39.88 | 1433.786 | -1.9 |
| 9424 | AfLEegKWKKZ | 72 | 483.6187 | 3 | 42.97 | 1447.838 | -2.5 |
| 10842 | TFDfgAKWKKZ | 72 | 458.2684 | 3 | 49.22 | 1371.77 | 9.6 |
| 4020 | bALGgHKWKKZ | 71 | 305.6994 | 4 | 20.69 | 1218.772 | -2.5 |
| 5081 | dSGedaKWKKZ | 71 | 461.2686 | 3 | 25.07 | 1380.782 | 1.7 |
| 5445 | gPAGGeKWKKZ | 71 | 398.9089 | 3 | 26.53 | 1193.707 | -1.9 |

| | | | | | | | |
|---|---|---|---|---|---|---|---|
| 2889 | hLASNDKWKKZ | 71 | 391.8953 | 3 | 15.78 | 1172.667 | -2.2 |
| 2909 | AhLSQQKWKKZ | 71 | 400.9114 | 3 | 15.86 | 1199.714 | -1.3 |
| 3884 | AAKDLFKWKKZ | 71 | 309.1916 | 4 | 20.1 | 1232.739 | -1.7 |
| 3896 | AAKDLFKWKKZ | 71 | 309.1914 | 4 | 20.15 | 1232.739 | -2.1 |
| 4029 | RKLgLKKWKKZ | 71 | 339.2438 | 4 | 20.73 | 1352.95 | -2.7 |
| 4070 | LQREQeKWKKZ | 71 | 360.7112 | 4 | 20.91 | 1438.82 | -2.8 |
| 4113 | AKQfFNKWKKZ | 71 | 350.7044 | 4 | 21.1 | 1398.793 | -2.7 |
| 4169 | AgTLGNKWKKZ | 71 | 293.6875 | 4 | 21.34 | 1170.724 | -2.3 |
| 4327 | AhAfgGKWKKZ | 71 | 306.4441 | 4 | 22 | 1221.739 | 6.9 |
| 4606 | LGARLFKWKKZ | 71 | 415.9349 | 3 | 23.13 | 1244.787 | -3.3 |
| 4850 | TRfQAFKWKKZ | 71 | 472.274 | 3 | 24.14 | 1413.803 | -2.1 |
| 4873 | TRfQAFKWKKZ | 71 | 354.4573 | 4 | 24.24 | 1413.803 | -2.1 |
| 4935 | FLAhaeKWKKZ | 71 | 326.2018 | 4 | 24.49 | 1300.781 | -2.1 |
| 5140 | KTfhKfKWKKZ | 71 | 369.9765 | 4 | 25.31 | 1475.881 | -2.5 |
| 5153 | TKfhKfKWKKZ | 71 | 369.9766 | 4 | 25.36 | 1475.881 | -2.2 |
| 5159 | LhLgAGKWKKZ | 71 | 385.5965 | 3 | 25.38 | 1153.77 | -2.1 |
| 5166 | LhLgAGKWKKZ | 71 | 385.5963 | 3 | 25.41 | 1153.77 | -2.7 |
| 5282 | gFgNTQKWKKZ | 71 | 444.9425 | 3 | 25.87 | 1331.808 | -1.6 |
| 5308 | gFgNTQKWKKZ | 71 | 444.9422 | 3 | 25.97 | 1331.808 | -2.4 |
| 5630 | GGhgfGKWKKZ | 71 | 398.9089 | 3 | 27.29 | 1193.707 | -2 |
| 5645 | ffKHgTKWKKZ | 71 | 382.7293 | 4 | 27.35 | 1526.891 | -2.1 |
| 5661 | ffKHgTKWKKZ | 71 | 382.7293 | 4 | 27.41 | 1526.891 | -2.1 |
| 5674 | ffKHgTKWKKZ | 71 | 382.7292 | 4 | 27.47 | 1526.891 | -2.4 |
| 5942 | fGENDLKWKKZ | 71 | 447.2422 | 3 | 28.55 | 1338.709 | -2.7 |
| 6180 | FfQEhLKWKKZ | 71 | 354.2057 | 4 | 29.54 | 1412.797 | -2.4 |
| 6584 | GfLHfGKWKKZ | 71 | 466.9316 | 3 | 31.16 | 1397.776 | -2.3 |
| 6585 | GfLHfGKWKKZ | 71 | 466.9316 | 3 | 31.16 | 1397.776 | -2.3 |
| 6596 | eAFfdEKWKKZ | 71 | 383.7106 | 4 | 31.21 | 1530.817 | -2.6 |
| 6597 | eAFfdEKWKKZ | 71 | 383.7106 | 4 | 31.21 | 1530.817 | -2.6 |
| 6629 | TNFLGfKWKKZ | 71 | 336.6952 | 4 | 31.34 | 1342.755 | -2.4 |
| 6655 | TNFLGfKWKKZ | 71 | 336.6952 | 4 | 31.44 | 1342.755 | -2.5 |
| 6682 | afNfLhKWKKZ | 71 | 362.4648 | 4 | 31.55 | 1445.834 | -2.5 |
| 6687 | TNFLGfKWKKZ | 71 | 336.6952 | 4 | 31.57 | 1342.755 | -2.3 |
| 6721 | afNfLhKWKKZ | 71 | 362.4648 | 4 | 31.7 | 1445.834 | -2.5 |
| 6857 | afNfgAKWKKZ | 71 | 362.4647 | 4 | 32.24 | 1445.834 | -2.6 |
| 6997 | GaSLEfKWKKZ | 71 | 325.1912 | 4 | 32.8 | 1296.734 | 1.3 |
| 7262 | PfgghPKWKKZ | 71 | 336.9685 | 4 | 33.85 | 1343.848 | -2.7 |
| 8245 | egTQeEKWKKZ | 71 | 489.9423 | 3 | 37.88 | 1466.807 | -1.6 |
| 8302 | DhSLffKWKKZ | 71 | 478.9351 | 3 | 38.14 | 1433.786 | -1.8 |
| 8375 | NGHNAfKWKKZ | 71 | 435.5683 | 3 | 38.45 | 1303.694 | -8.2 |
| 8831 | FFNeAeKWKKZ | 71 | 366.2001 | 4 | 40.44 | 1460.776 | -2.9 |
| 9175 | FFLEegKWKKZ | 71 | 362.9661 | 4 | 41.9 | 1447.838 | -1.9 |
| 9190 | FFLEegKWKKZ | 71 | 362.9659 | 4 | 41.97 | 1447.838 | -2.2 |

| | | | | | | | |
|---|---|---|---|---|---|---|---|
| 9214 | FFLEegKWKKZ | 71 | 362.966 | 4 | 42.07 | 1447.838 | -2.1 |
| 9226 | FFLEegKWKKZ | 71 | 362.9659 | 4 | 42.12 | 1447.838 | -2.2 |
| 9235 | FFLEegKWKKZ | 71 | 362.966 | 4 | 42.16 | 1447.838 | -2 |
| 9259 | FFLEegKWKKZ | 71 | 362.9659 | 4 | 42.26 | 1447.838 | -2.2 |
| 9283 | FFLEegKWKKZ | 71 | 362.9661 | 4 | 42.36 | 1447.838 | -1.9 |
| 9307 | FFLEegKWKKZ | 71 | 362.9661 | 4 | 42.47 | 1447.838 | -1.9 |
| 9334 | FFLEegKWKKZ | 71 | 362.9661 | 4 | 42.58 | 1447.838 | -1.8 |
| 9361 | FFLEegKWKKZ | 71 | 362.966 | 4 | 42.7 | 1447.838 | -2 |
| 9420 | FFLEegKWKKZ | 71 | 362.9661 | 4 | 42.96 | 1447.838 | -1.8 |
| 9421 | FFLEegKWKKZ | 71 | 362.9661 | 4 | 42.96 | 1447.838 | -1.8 |
| 9457 | FFLEegKWKKZ | 71 | 362.966 | 4 | 43.12 | 1447.838 | -2 |
| 9658 | AfLEegKWKKZ | 71 | 483.6191 | 3 | 44 | 1447.838 | -1.6 |
| 9757 | hfFgSfKWKKZ | 71 | 494.2874 | 3 | 44.45 | 1479.843 | -1.9 |
| 10053 | GAHLfGKWKKZ | 71 | 416.2415 | 3 | 45.75 | 1245.713 | -8.6 |
| 10820 | HPEfgAKWKKZ | 71 | 458.2684 | 3 | 49.12 | 1371.782 | 1.4 |
| 10832 | EHPfgAKWKKZ | 71 | 458.2681 | 3 | 49.17 | 1371.782 | 0.8 |
| 10844 | TFDfgAKWKKZ | 71 | 458.2684 | 3 | 49.23 | 1371.77 | 9.6 |
| 8732 | GGHFgSKWKKZ | 70 | 400.9014 | 3 | 40 | 1199.693 | -8.8 |
| 10640 | NFGEPfKWKKZ | 70 | 452.5809 | 3 | 48.31 | 1354.719 | 1.8 |
| 11482 | DeGSfGKWKKZ | 70 | 442.2333 | 3 | 52.03 | 1323.676 | 1.4 |
| 11634 | TiDQPfKWKKZ | 70 | 469.2522 | 3 | 52.73 | 1404.73 | 3.2 |
| 11719 | NeGEPfKWKKZ | 70 | 469.2524 | 3 | 53.12 | 1404.734 | 0.9 |
| 2907 | hLASNDKWKKZ | 70 | 391.8954 | 3 | 15.86 | 1172.667 | -1.8 |
| 2971 | LhASNDKWKKZ | 70 | 294.1734 | 4 | 16.14 | 1172.667 | -1.8 |
| 3158 | SFEDGGKWKKZ | 70 | 394.2078 | 3 | 16.95 | 1179.604 | -1.7 |
| 3792 | TcNfhGKWKKZ | 70 | 328.4417 | 4 | 19.71 | 1309.741 | -2.2 |
| 3978 | RKLgLKKWKKZ | 70 | 339.2437 | 4 | 20.51 | 1352.95 | -2.9 |
| 3990 | RKLgLKKWKKZ | 70 | 339.2438 | 4 | 20.56 | 1352.95 | -2.6 |
| 4005 | RKLgLKKWKKZ | 70 | 339.2439 | 4 | 20.62 | 1352.95 | -2.3 |
| 4512 | LGARLFKWKKZ | 70 | 415.9353 | 3 | 22.75 | 1244.787 | -2.2 |
| 4574 | LGARLFKWKKZ | 70 | 415.9355 | 3 | 23 | 1244.787 | -1.9 |
| 4906 | fTAhfNKWKKZ | 70 | 352.202 | 4 | 24.37 | 1404.771 | 5.7 |
| 4919 | fTAhfNKWKKZ | 70 | 352.2019 | 4 | 24.42 | 1404.771 | 5.5 |
| 4951 | fTAhfNKWKKZ | 70 | 352.2022 | 4 | 24.55 | 1404.771 | 6.5 |
| 5604 | GGhgfGKWKKZ | 70 | 398.909 | 3 | 27.18 | 1193.707 | -1.8 |
| 5617 | GGhgfGKWKKZ | 70 | 398.9091 | 3 | 27.24 | 1193.707 | -1.5 |
| 6588 | GfLHfGKWKKZ | 70 | 466.9314 | 3 | 31.17 | 1397.776 | -2.6 |
| 7000 | afNfLhKWKKZ | 70 | 362.4648 | 4 | 32.81 | 1445.834 | -2.4 |
| 7389 | PfgghPKWKKZ | 70 | 336.9686 | 4 | 34.37 | 1343.848 | -2.2 |
| 8185 | egTQeEKWKKZ | 70 | 489.9419 | 3 | 37.62 | 1466.807 | -2.3 |
| 8300 | TbALgQKWKKZ | 70 | 418.935 | 3 | 38.13 | 1253.797 | -11.3 |
| 8402 | NHGANfKWKKZ | 70 | 435.5687 | 3 | 38.57 | 1303.694 | -7.4 |
| 8481 | DhSLffKWKKZ | 70 | 478.9348 | 3 | 38.92 | 1433.786 | -2.3 |

| | | | | | | | |
|---|---|---|---|---|---|---|---|
| 8640 | DhSLefKWKKZ | 70 | 470.2625 | 3 | 39.59 | 1407.77 | -3.2 |
| 8664 | DhSLffKWKKZ | 70 | 478.9348 | 3 | 39.69 | 1433.786 | -2.2 |
| 8746 | DhSLffKWKKZ | 70 | 478.9352 | 3 | 40.06 | 1433.786 | -1.6 |
| 8789 | FFNeAeKWKKZ | 70 | 366.2003 | 4 | 40.25 | 1460.776 | -2.4 |
| 9202 | FFLEegKWKKZ | 70 | 362.9661 | 4 | 42.02 | 1447.838 | -1.9 |
| 9247 | FFLEegKWKKZ | 70 | 362.9659 | 4 | 42.21 | 1447.838 | -2.5 |
| 9295 | FFLEegKWKKZ | 70 | 362.9661 | 4 | 42.42 | 1447.838 | -1.9 |
| 9325 | FFLEegKWKKZ | 70 | 362.9661 | 4 | 42.55 | 1447.838 | -1.9 |
| 9408 | FFLEegKWKKZ | 70 | 362.966 | 4 | 42.91 | 1447.838 | -2 |
| 9409 | FFLEegKWKKZ | 70 | 362.966 | 4 | 42.91 | 1447.838 | -2 |
| 9672 | AfLEegKWKKZ | 70 | 483.619 | 3 | 44.07 | 1447.838 | -1.8 |
| 9673 | AfLEegKWKKZ | 70 | 483.619 | 3 | 44.07 | 1447.838 | -1.8 |
| 9993 | AGHLfGKWKKZ | 70 | 416.2419 | 3 | 45.5 | 1245.713 | -7.7 |
| 10017 | AGHLfGKWKKZ | 70 | 416.2418 | 3 | 45.6 | 1245.713 | -7.9 |
| 10029 | GAHLfGKWKKZ | 70 | 416.242 | 3 | 45.65 | 1245.713 | -7.3 |
| 11636 | TiDQPfKWKKZ | 70 | 469.2522 | 3 | 52.74 | 1404.73 | 3.2 |
| 11644 | TiDQPfKWKKZ | 70 | 469.2522 | 3 | 52.78 | 1404.73 | 3.2 |
| 11646 | TiDQPfKWKKZ | 70 | 469.2522 | 3 | 52.79 | 1404.73 | 3.2 |
| 11666 | eGDQPfKWKKZ | 70 | 469.2525 | 3 | 52.88 | 1404.734 | 1.1 |

### Appendix III. Peptides identified after cleaving 1,000-member PMO-library from beads

| Scan | Peptide | ALC (%) | m/z | z | RT | Mass | ppm |
|---|---|---|---|---|---|---|---|
| 4733 | gLESTNKWKKX | 99 | 420.5873 | 3 | 23.86 | 1258.74 | 0.2 |
| 5511 | AADGfNKWKKX | 99 | 310.6705 | 4 | 25.92 | 1238.656 | -2.6 |
| 5517 | AADGfNKWKKX | 99 | 413.8917 | 3 | 25.93 | 1238.656 | -2.3 |
| 6118 | GSgFEQKWKKX | 99 | 421.9114 | 3 | 27.64 | 1262.714 | -1 |
| 6120 | GSgFEQKWKKX | 99 | 421.9114 | 3 | 27.65 | 1262.714 | -1 |
| 6131 | PTDGTGKWKKX | 99 | 372.8765 | 3 | 27.68 | 1115.609 | -1 |
| 6134 | GSgFEQKWKKX | 99 | 421.9113 | 3 | 27.69 | 1262.714 | -1.1 |
| 6136 | GSgFEQKWKKX | 99 | 421.9113 | 3 | 27.69 | 1262.714 | -1.1 |
| 6152 | GSgFEQKWKKX | 99 | 421.9116 | 3 | 27.74 | 1262.714 | -0.5 |
| 6154 | GSgFEQKWKKX | 99 | 421.9116 | 3 | 27.74 | 1262.714 | -0.5 |
| 6168 | GSgFEQKWKKX | 99 | 421.9118 | 3 | 27.78 | 1262.714 | 0 |
| 6170 | GSgFEQKWKKX | 99 | 421.9118 | 3 | 27.79 | 1262.714 | 0 |
| 6186 | GSgFEQKWKKX | 99 | 421.9116 | 3 | 27.83 | 1262.714 | -0.5 |
| 6202 | GSgFEQKWKKX | 99 | 421.9117 | 3 | 27.88 | 1262.714 | -0.2 |
| 12680 | PTgSDeKWKKX | 99 | 438.252 | 3 | 49.7 | 1311.734 | 0.1 |
| 4544 | NSSbgLKWKKX | 98 | 414.9274 | 3 | 23.37 | 1241.761 | -0.5 |
| 4735 | gLESTNKWKKX | 98 | 420.5873 | 3 | 23.86 | 1258.74 | 0.2 |

| | | | | | | | |
|---|---|---|---|---|---|---|---|
| 5527 | AADGfNKWKKX | 98 | 413.8925 | 3 | 25.96 | 1238.656 | -0.3 |
| 6161 | GPEEGeKWKKX | 98 | 418.8925 | 3 | 27.76 | 1253.656 | 0.2 |
| 6188 | GSgFEQKWKKX | 98 | 421.9116 | 3 | 27.84 | 1262.714 | -0.5 |
| 6204 | GSgFEQKWKKX | 98 | 421.9117 | 3 | 27.89 | 1262.714 | -0.2 |
| 6226 | GSgFEQKWKKX | 98 | 421.9117 | 3 | 27.95 | 1262.714 | -0.2 |
| 6751 | ANfSgaKWKKX | 98 | 437.5958 | 3 | 29.52 | 1309.766 | -0.3 |
| 6764 | ANfSgaKWKKX | 98 | 437.5955 | 3 | 29.57 | 1309.766 | -0.9 |
| 6786 | ANfSgaKWKKX | 98 | 437.5951 | 3 | 29.64 | 1309.766 | -1.9 |
| 7436 | GDDfNSKWKKX | 98 | 433.8874 | 3 | 31.91 | 1298.641 | -0.2 |
| 7528 | DTgPQgKWKKX | 98 | 428.599 | 3 | 32.28 | 1282.776 | -0.8 |
| 7532 | DTgPQgKWKKX | 98 | 428.5992 | 3 | 32.29 | 1282.776 | -0.2 |
| 8405 | PTDDNgKWKKX | 98 | 419.903 | 3 | 35.17 | 1256.688 | -0.6 |
| 12669 | TPgSDeKWKKX | 98 | 438.2518 | 3 | 49.67 | 1311.734 | -0.2 |
| 3891 | GRETgNKWKKX | 97 | 424.9226 | 3 | 21.73 | 1271.746 | -0.3 |
| 4278 | AgEQEdKWKKX | 97 | 337.9487 | 4 | 22.72 | 1347.766 | -0.5 |
| 4815 | GFPbLGKWKKX | 97 | 395.9133 | 3 | 24.06 | 1184.718 | -0.1 |
| 4855 | ATQgEDKWKKX | 97 | 420.5751 | 3 | 24.17 | 1258.703 | 0 |
| 5065 | FgESATKWKKX | 97 | 417.5798 | 3 | 24.72 | 1249.718 | -0.5 |
| 5519 | AADGfNKWKKX | 97 | 413.8917 | 3 | 25.94 | 1238.656 | -2.3 |
| 5547 | AADGfNKWKKX | 97 | 413.8925 | 3 | 26.01 | 1238.656 | -0.2 |
| 6121 | GPEEGeKWKKX | 97 | 418.8921 | 3 | 27.65 | 1253.656 | -0.9 |
| 6123 | GPEEGeKWKKX | 97 | 418.8921 | 3 | 27.65 | 1253.656 | -0.9 |
| 6163 | GPEEGeKWKKX | 97 | 418.8925 | 3 | 27.77 | 1253.656 | 0.2 |
| 6228 | GSgFEQKWKKX | 97 | 421.9117 | 3 | 27.95 | 1262.714 | -0.2 |
| 6477 | NGgALDKWKKX | 97 | 395.9081 | 3 | 28.69 | 1184.703 | -0.5 |
| 6775 | ANfSgaKWKKX | 97 | 437.5954 | 3 | 29.61 | 1309.766 | -1.2 |
| 7432 | GDDfNSKWKKX | 97 | 433.8874 | 3 | 31.9 | 1298.641 | -0.1 |
| 7438 | GDDfNSKWKKX | 97 | 433.8874 | 3 | 31.92 | 1298.641 | -0.2 |
| 7446 | GDDfNSKWKKX | 97 | 433.8875 | 3 | 31.94 | 1298.641 | 0.1 |
| 7543 | DTgPQgKWKKX | 97 | 428.5991 | 3 | 32.33 | 1282.776 | -0.5 |
| 7557 | DTgPQgKWKKX | 97 | 428.5993 | 3 | 32.37 | 1282.776 | 0 |
| 7579 | DTgPQgKWKKX | 97 | 428.5989 | 3 | 32.43 | 1282.776 | -0.9 |
| 7594 | DTgPQgKWKKX | 97 | 428.599 | 3 | 32.49 | 1282.776 | -0.7 |
| 8389 | PTDDNgKWKKX | 97 | 419.9028 | 3 | 35.12 | 1256.688 | -0.9 |
| 8394 | PTDDNgKWKKX | 97 | 419.9032 | 3 | 35.13 | 1256.688 | 0.1 |
| 8398 | PTDDNgKWKKX | 97 | 419.9033 | 3 | 35.15 | 1256.688 | 0.2 |
| 8407 | PTDDNgKWKKX | 97 | 419.903 | 3 | 35.17 | 1256.688 | -0.6 |
| 12671 | TPgSDeKWKKX | 97 | 438.2518 | 3 | 49.68 | 1311.734 | -0.2 |
| 3540 | ARLREQKWKKX | 96 | 336.211 | 4 | 20.87 | 1340.815 | -0.1 |
| 3577 | ARLREQKWKKX | 96 | 336.2111 | 4 | 20.96 | 1340.815 | 0 |
| 3876 | GRETgNKWKKX | 96 | 318.9434 | 4 | 21.7 | 1271.746 | -1.4 |
| 4049 | GGAFNTKWKKX | 96 | 379.2172 | 3 | 22.14 | 1134.63 | 0 |
| 4208 | LRLQADKWKKX | 96 | 321.9529 | 4 | 22.54 | 1283.783 | 0.1 |

| | | | | | | | |
|---|---|---|---|---|---|---|---|
| 4298 | AgEQEdKWKKX | 96 | 337.9487 | 4 | 22.77 | 1347.766 | -0.6 |
| 4403 | DNQhTFKWKKX | 96 | 426.9028 | 3 | 23.03 | 1277.688 | -1 |
| 4546 | NSSbgLKWKKX | 96 | 414.9274 | 3 | 23.38 | 1241.761 | -0.5 |
| 4742 | LgESTNKWKKX | 96 | 315.6919 | 4 | 23.88 | 1258.74 | -1.1 |
| 4772 | GFPbLGKWKKX | 96 | 395.9133 | 3 | 23.96 | 1184.718 | -0.2 |
| 4774 | GFPbLGKWKKX | 96 | 395.9133 | 3 | 23.96 | 1184.718 | -0.2 |
| 4817 | GFPbLGKWKKX | 96 | 395.9133 | 3 | 24.07 | 1184.718 | -0.1 |
| 4839 | GFPbLGKWKKX | 96 | 395.9131 | 3 | 24.13 | 1184.718 | -0.6 |
| 4846 | ATQgEDKWKKX | 96 | 315.683 | 4 | 24.15 | 1258.703 | -0.2 |
| 5067 | FgESATKWKKX | 96 | 417.5798 | 3 | 24.72 | 1249.718 | -0.5 |
| 5081 | FgESATKWKKX | 96 | 417.58 | 3 | 24.76 | 1249.718 | -0.1 |
| 6133 | PTDGTGKWKKX | 96 | 372.8765 | 3 | 27.69 | 1115.609 | -1 |
| 6149 | PTDGTGKWKKX | 96 | 372.8766 | 3 | 27.73 | 1115.609 | -0.5 |
| 6493 | NGgALDKWKKX | 96 | 395.9081 | 3 | 28.74 | 1184.703 | -0.5 |
| 6753 | ANfSgaKWKKX | 96 | 437.5958 | 3 | 29.53 | 1309.766 | -0.3 |
| 6766 | ANfSgaKWKKX | 96 | 437.5955 | 3 | 29.58 | 1309.766 | -0.9 |
| 6788 | ANfSgaKWKKX | 96 | 437.5951 | 3 | 29.65 | 1309.766 | -1.9 |
| 7434 | GDDfNSKWKKX | 96 | 433.8874 | 3 | 31.91 | 1298.641 | -0.1 |
| 7448 | GDDfNSKWKKX | 96 | 433.8875 | 3 | 31.95 | 1298.641 | 0.1 |
| 7530 | DTgPQgKWKKX | 96 | 428.599 | 3 | 32.29 | 1282.776 | -0.8 |
| 8396 | PTDDNgKWKKX | 96 | 419.9032 | 3 | 35.14 | 1256.688 | 0.1 |
| 8400 | PTDDNgKWKKX | 96 | 419.9033 | 3 | 35.15 | 1256.688 | 0.2 |
| 12682 | PTgSDeKWKKX | 96 | 438.252 | 3 | 49.71 | 1311.734 | 0.1 |
| 12701 | PTgSDeKWKKX | 96 | 438.2519 | 3 | 49.76 | 1311.734 | 0 |
| 3667 | NSdaFGKWKKX | 95 | 423.912 | 3 | 21.18 | 1268.714 | 0 |
| 3799 | QAFRNdKWKKX | 95 | 345.9556 | 4 | 21.51 | 1379.794 | -0.4 |
| 3825 | NKTGghKWKKX | 95 | 300.9449 | 4 | 21.57 | 1199.75 | 0.2 |
| 4841 | GFPbLGKWKKX | 95 | 395.9131 | 3 | 24.14 | 1184.718 | -0.6 |
| 5083 | FgESATKWKKX | 95 | 417.58 | 3 | 24.76 | 1249.718 | -0.1 |
| 5524 | AADGfNKWKKX | 95 | 310.6711 | 4 | 25.95 | 1238.656 | -0.5 |
| 5572 | AAAgHgKWKKX | 95 | 398.2621 | 3 | 26.08 | 1191.761 | 3.3 |
| 6151 | PTDGTGKWKKX | 95 | 372.8766 | 3 | 27.74 | 1115.609 | -0.5 |
| 6479 | NGgALDKWKKX | 95 | 395.9081 | 3 | 28.7 | 1184.703 | -0.5 |
| 6768 | ANfSgaKWKKX | 95 | 437.5955 | 3 | 29.58 | 1309.766 | -0.9 |
| 6803 | ANfSgaKWKKX | 95 | 437.5952 | 3 | 29.7 | 1309.766 | -1.7 |
| 7534 | DTgPQgKWKKX | 95 | 428.5992 | 3 | 32.3 | 1282.776 | -0.2 |
| 7536 | DTgPQgKWKKX | 95 | 428.5992 | 3 | 32.31 | 1282.776 | -0.3 |
| 7545 | DTgPQgKWKKX | 95 | 428.5991 | 3 | 32.34 | 1282.776 | -0.5 |
| 7555 | DTgPQgKWKKX | 95 | 428.5994 | 3 | 32.36 | 1282.776 | 0.3 |
| 7581 | DTgPQgKWKKX | 95 | 428.5989 | 3 | 32.44 | 1282.776 | -0.9 |
| 7583 | DTgPQgKWKKX | 95 | 428.5992 | 3 | 32.45 | 1282.776 | -0.3 |
| 7588 | DTgPQgKWKKX | 95 | 428.599 | 3 | 32.47 | 1282.776 | -0.7 |
| 7603 | DTgPQgKWKKX | 95 | 428.599 | 3 | 32.53 | 1282.776 | -0.8 |

| | | | | | | | |
|---|---|---|---|---|---|---|---|
| 7610 | DTgPQgKWKKX | 95 | 428.5992 | 3 | 32.56 | 1282.776 | -0.4 |
| 7824 | GAAELfKWKKX | 95 | 418.2449 | 3 | 33.32 | 1251.713 | 0.3 |
| 8215 | PTfhAKKWKKX | 95 | 431.9324 | 3 | 34.57 | 1292.776 | -0.2 |
| 8248 | PTfhAKKWKKX | 95 | 431.932 | 3 | 34.67 | 1292.776 | -1.1 |
| 8391 | PTDDNgKWKKX | 95 | 419.9028 | 3 | 35.12 | 1256.688 | -0.9 |
| 12719 | PTgSDeKWKKX | 95 | 438.252 | 3 | 49.81 | 1311.734 | 0.1 |
| 3652 | ALaFGTKWKKX | 94 | 393.2448 | 3 | 21.15 | 1176.713 | -0.4 |
| 3846 | ALaKAFKWKKX | 94 | 305.4512 | 4 | 21.63 | 1217.776 | -0.3 |
| 3864 | ALaKAFKWKKX | 94 | 305.4512 | 4 | 21.67 | 1217.776 | -0.3 |
| 4025 | QSKgFaKWKKX | 94 | 327.2094 | 4 | 22.09 | 1304.808 | 0.2 |
| 4280 | AgEQEdKWKKX | 94 | 337.9487 | 4 | 22.72 | 1347.766 | -0.5 |
| 4609 | RFKDgGKWKKX | 94 | 330.4581 | 4 | 23.53 | 1317.804 | -0.2 |
| 5529 | AADGfNKWKKX | 94 | 413.8925 | 3 | 25.96 | 1238.656 | -0.3 |
| 5544 | hGDGfNKWKKX | 94 | 310.6711 | 4 | 26 | 1238.656 | -0.5 |
| 5549 | AADGfNKWKKX | 94 | 413.8925 | 3 | 26.01 | 1238.656 | -0.2 |
| 5582 | AAAgHgKWKKX | 94 | 298.9472 | 4 | 26.1 | 1191.761 | -0.7 |
| 6192 | GPEEGeKWKKX | 94 | 418.8923 | 3 | 27.85 | 1253.656 | -0.4 |
| 6514 | NGgALDKWKKX | 94 | 395.908 | 3 | 28.8 | 1184.703 | -0.6 |
| 7453 | GDDfNSKWKKX | 94 | 433.8874 | 3 | 31.97 | 1298.641 | -0.3 |
| 7455 | GDDfNSKWKKX | 94 | 433.8874 | 3 | 31.97 | 1298.641 | -0.3 |
| 7559 | DTgPQgKWKKX | 94 | 428.5993 | 3 | 32.38 | 1282.776 | 0 |
| 7568 | DTgPQgKWKKX | 94 | 428.5992 | 3 | 32.4 | 1282.776 | -0.4 |
| 7614 | DTgPQgKWKKX | 94 | 428.5995 | 3 | 32.57 | 1282.776 | 0.4 |
| 8361 | ATfNQeKWKKX | 94 | 474.9272 | 3 | 35 | 1421.761 | -0.6 |
| 3614 | hAEbRLKWKKX | 93 | 317.9541 | 4 | 21.06 | 1267.788 | -0.4 |
| 3816 | QAFRNdKWKKX | 93 | 345.9556 | 4 | 21.55 | 1379.794 | -0.2 |
| 3843 | GDRSaFKWKKX | 93 | 313.4333 | 4 | 21.62 | 1249.704 | -0.2 |
| 3878 | GRETgNKWKKX | 93 | 318.9434 | 4 | 21.7 | 1271.746 | -1.4 |
| 3893 | GRETgNKWKKX | 93 | 424.9226 | 3 | 21.74 | 1271.746 | -0.3 |
| 4051 | GGAFNTKWKKX | 93 | 379.2172 | 3 | 22.15 | 1134.63 | 0 |
| 4300 | AgEQEdKWKKX | 93 | 337.9487 | 4 | 22.77 | 1347.766 | -0.6 |
| 4550 | ARgFGRKWKKX | 93 | 326.4622 | 4 | 23.39 | 1301.82 | 0.1 |
| 5574 | AAAgHgKWKKX | 93 | 398.2621 | 3 | 26.08 | 1191.761 | 3.3 |
| 6194 | GPEEGeKWKKX | 93 | 418.8923 | 3 | 27.86 | 1253.656 | -0.4 |
| 6495 | NGgALDKWKKX | 93 | 395.9081 | 3 | 28.74 | 1184.703 | -0.5 |
| 6777 | ANfSgaKWKKX | 93 | 437.5954 | 3 | 29.61 | 1309.766 | -1.2 |
| 7460 | GDDfNSKWKKX | 93 | 433.8875 | 3 | 31.99 | 1298.641 | 0 |
| 7520 | DTgPQgKWKKX | 93 | 428.5992 | 3 | 32.26 | 1282.776 | -0.2 |
| 7522 | DTgPQgKWKKX | 93 | 428.5992 | 3 | 32.26 | 1282.776 | -0.2 |
| 7538 | DTgPQgKWKKX | 93 | 428.5992 | 3 | 32.31 | 1282.776 | -0.3 |
| 7566 | DTgPQgKWKKX | 93 | 428.5995 | 3 | 32.39 | 1282.776 | 0.4 |
| 7570 | DTgPQgKWKKX | 93 | 428.5992 | 3 | 32.41 | 1282.776 | -0.4 |
| 7585 | DTgPQgKWKKX | 93 | 428.5992 | 3 | 32.45 | 1282.776 | -0.3 |

| | | | | | | | |
|---|---|---|---|---|---|---|---|
| 7596 | DTgPQgKWKKX | 93 | 428.599 | 3 | 32.5 | 1282.776 | -0.7 |
| 7826 | GAAELfKWKKX | 93 | 418.2449 | 3 | 33.33 | 1251.713 | 0.3 |
| 12703 | PTgSDeKWKKX | 93 | 438.2519 | 3 | 49.77 | 1311.734 | 0 |
| 3579 | ARLREQKWKKX | 92 | 336.2111 | 4 | 20.97 | 1340.815 | 0 |
| 3699 | RDgSbQKWKKX | 92 | 332.7039 | 4 | 21.26 | 1326.789 | -1.5 |
| 4210 | LRLQADKWKKX | 92 | 321.9529 | 4 | 22.55 | 1283.783 | 0.1 |
| 4364 | GGSbgLKWKKX | 92 | 385.917 | 3 | 22.93 | 1154.729 | 0.4 |
| 4405 | NDQhTFKWKKX | 92 | 426.9028 | 3 | 23.03 | 1277.688 | -1 |
| 4425 | DNQhTFKWKKX | 92 | 426.9034 | 3 | 23.08 | 1277.688 | 0.3 |
| 4744 | LgESTNKWKKX | 92 | 315.6919 | 4 | 23.89 | 1258.74 | -1.1 |
| 4857 | ATQgEDKWKKX | 92 | 420.5751 | 3 | 24.18 | 1258.703 | 0 |
| 4868 | ATQgEDKWKKX | 92 | 315.683 | 4 | 24.21 | 1258.703 | -0.5 |
| 5513 | AADGfNKWKKX | 92 | 310.6705 | 4 | 25.92 | 1238.656 | -2.6 |
| 5566 | AADGfNKWKKX | 92 | 413.8925 | 3 | 26.06 | 1238.656 | -0.2 |
| 5657 | ALAATgKWKKX | 92 | 286.4404 | 4 | 26.32 | 1141.734 | -0.9 |
| 5806 | GfGDGAKWKKX | 92 | 390.2136 | 3 | 26.72 | 1167.619 | 0 |
| 6821 | ANfSgaKWKKX | 92 | 437.5955 | 3 | 29.76 | 1309.766 | -0.9 |
| 7462 | GDDfNSKWKKX | 92 | 433.8875 | 3 | 32 | 1298.641 | 0 |
| 7590 | DTgPQgKWKKX | 92 | 428.599 | 3 | 32.47 | 1282.776 | -0.7 |
| 7822 | GAAELfKWKKX | 92 | 418.2449 | 3 | 33.31 | 1251.713 | 0.1 |
| 7833 | GAAELfKWKKX | 92 | 418.2448 | 3 | 33.35 | 1251.713 | -0.1 |
| 7840 | GAAELfKWKKX | 92 | 418.2449 | 3 | 33.37 | 1251.713 | 0.1 |
| 12721 | TPgSDeKWKKX | 92 | 438.252 | 3 | 49.82 | 1311.734 | 0.1 |
| 3542 | ARLREQKWKKX | 91 | 336.211 | 4 | 20.87 | 1340.815 | -0.1 |
| 3840 | QAFRNdKWKKX | 91 | 345.9555 | 4 | 21.61 | 1379.794 | -0.5 |
| 3921 | GGNPgaKWKKX | 91 | 380.9047 | 3 | 21.81 | 1139.693 | -0.2 |
| 4230 | FTKNTFKWKKX | 91 | 442.9276 | 3 | 22.6 | 1325.761 | 0.3 |
| 4331 | NDATGFKWKKX | 91 | 398.5523 | 3 | 22.85 | 1192.635 | -0.1 |
| 4848 | ATQgEDKWKKX | 91 | 315.683 | 4 | 24.15 | 1258.703 | -0.2 |
| 5520 | NGQSeDKWKKX | 91 | 429.5584 | 3 | 25.94 | 1285.657 | -2.6 |
| 5526 | GhDGfNKWKKX | 91 | 310.6711 | 4 | 25.96 | 1238.656 | -0.5 |
| 5563 | hGDGfNKWKKX | 91 | 310.6712 | 4 | 26.05 | 1238.656 | -0.1 |
| 5671 | AhDgdLKWKKX | 91 | 425.9356 | 3 | 26.35 | 1274.786 | -1 |
| 5688 | AhDgdLKWKKX | 91 | 425.9355 | 3 | 26.39 | 1274.786 | -1.3 |
| 5695 | AhDgdLKWKKX | 91 | 425.9357 | 3 | 26.41 | 1274.786 | -0.9 |
| 5997 | NGEeNDKWKKX | 91 | 438.8909 | 3 | 27.28 | 1313.652 | -0.6 |
| 6938 | AGGQfFKWKKX | 91 | 424.5728 | 3 | 30.13 | 1270.698 | -0.8 |
| 7605 | DTgPQgKWKKX | 91 | 428.599 | 3 | 32.54 | 1282.776 | -0.8 |
| 3616 | AhEbRLKWKKX | 90 | 317.9541 | 4 | 21.06 | 1267.788 | -0.4 |
| 3669 | NSdaFGKWKKX | 90 | 423.912 | 3 | 21.19 | 1268.714 | 0 |
| 3845 | GDRSaFKWKKX | 90 | 313.4333 | 4 | 21.62 | 1249.704 | -0.2 |
| 4239 | AhLRPPKWKKX | 90 | 302.7001 | 4 | 22.62 | 1206.771 | 0 |
| 5546 | hGDGfNKWKKX | 90 | 310.6711 | 4 | 26.01 | 1238.656 | -0.5 |

| | | | | | | | |
|---|---|---|---|---|---|---|---|
| 5568 | AADGfNKWKKX | 90 | 413.8925 | 3 | 26.07 | 1238.656 | -0.2 |
| 5584 | AAAgHgKWKKX | 90 | 298.9472 | 4 | 26.11 | 1191.761 | -0.7 |
| 5594 | AAAgHgKWKKX | 90 | 398.2604 | 3 | 26.14 | 1191.761 | -0.9 |
| 6304 | GGLTDgKWKKX | 90 | 386.9041 | 3 | 28.18 | 1157.692 | -1.5 |
| 6805 | ANfSgaKWKKX | 90 | 437.5952 | 3 | 29.7 | 1309.766 | -1.7 |
| 6899 | ANGAfFKWKKX | 90 | 424.5732 | 3 | 30.01 | 1270.698 | 0.1 |
| 6925 | ANGAfFKWKKX | 90 | 424.5732 | 3 | 30.09 | 1270.698 | 0.2 |
| 7464 | GDDfNSKWKKX | 90 | 433.8873 | 3 | 32 | 1298.641 | -0.4 |
| 7850 | GGhELfKWKKX | 90 | 418.2448 | 3 | 33.4 | 1251.713 | -0.4 |
| 8217 | PTfhAKKWKKX | 90 | 431.9324 | 3 | 34.58 | 1292.776 | -0.2 |
| 3701 | RDgSbQKWKKX | 89 | 332.7039 | 4 | 21.27 | 1326.789 | -1.5 |
| 3818 | QAFRNdKWKKX | 89 | 345.9556 | 4 | 21.56 | 1379.794 | -0.2 |
| 3923 | GGNPgaKWKKX | 89 | 380.9047 | 3 | 21.81 | 1139.693 | -0.2 |
| 4058 | NQEQSFKWKKX | 89 | 441.2383 | 3 | 22.17 | 1320.694 | -0.6 |
| 4194 | GfGAHAKWKKX | 89 | 402.2295 | 3 | 22.51 | 1203.667 | 0.1 |
| 4333 | NDATGFKWKKX | 89 | 398.5523 | 3 | 22.85 | 1192.635 | -0.1 |
| 4427 | DNQhTFKWKKX | 89 | 426.9034 | 3 | 23.09 | 1277.688 | 0.3 |
| 5216 | GGThQeKWKKX | 89 | 405.2329 | 3 | 25.11 | 1212.677 | 0.1 |
| 5379 | aTDGTGKWKKX | 89 | 373.8811 | 3 | 25.55 | 1118.62 | 1.7 |
| 5649 | AhDgdLKWKKX | 89 | 425.9356 | 3 | 26.29 | 1274.786 | -1.2 |
| 6516 | NGgALDKWKKX | 89 | 395.908 | 3 | 28.8 | 1184.703 | -0.6 |
| 6837 | ANfSgaKWKKX | 89 | 437.596 | 3 | 29.81 | 1309.766 | 0.3 |
| 6909 | AGGQfFKWKKX | 89 | 424.5732 | 3 | 30.04 | 1270.698 | 0.1 |
| 7466 | GDDfNSKWKKX | 89 | 433.8873 | 3 | 32.01 | 1298.641 | -0.4 |
| 8063 | NGDDPeKWKKX | 89 | 428.556 | 3 | 34.13 | 1282.646 | 0.4 |
| 8250 | PTfhAKKWKKX | 89 | 431.932 | 3 | 34.67 | 1292.776 | -1.1 |
| 8306 | PTFEGAKWKKX | 89 | 397.5607 | 3 | 34.84 | 1189.661 | -0.2 |
| 3827 | KNTGghKWKKX | 88 | 300.9449 | 4 | 21.58 | 1199.75 | 0.2 |
| 3866 | ALaKAFKWKKX | 88 | 305.4512 | 4 | 21.67 | 1217.776 | -0.3 |
| 5240 | GGThQeKWKKX | 88 | 405.2328 | 3 | 25.17 | 1212.677 | -0.3 |
| 5596 | AAAgHgKWKKX | 88 | 398.2604 | 3 | 26.14 | 1191.761 | -0.9 |
| 7612 | DTgPQgKWKKX | 88 | 428.5992 | 3 | 32.56 | 1282.776 | -0.4 |
| 7857 | GAAELfKWKKX | 88 | 418.2455 | 3 | 33.42 | 1251.713 | 1.5 |
| 8240 | TPfhAKKWKKX | 88 | 431.9321 | 3 | 34.64 | 1292.776 | -1 |
| 3654 | ALdAGTKWKKX | 87 | 393.2448 | 3 | 21.15 | 1176.713 | -0.4 |
| 3755 | LLREGAKWKKX | 87 | 409.9274 | 3 | 21.4 | 1226.761 | -0.7 |
| 3848 | ALaKAFKWKKX | 87 | 305.4512 | 4 | 21.63 | 1217.776 | -0.3 |
| 3953 | PSPKbQKWKKX | 87 | 313.6975 | 4 | 21.9 | 1250.761 | -0.1 |
| 3987 | NKAFDDKWKKX | 87 | 426.9031 | 3 | 21.99 | 1277.688 | -0.3 |
| 4027 | QSKgFaKWKKX | 87 | 327.2094 | 4 | 22.09 | 1304.808 | 0.2 |
| 4181 | GGfhKSKWKKX | 87 | 409.2448 | 3 | 22.47 | 1224.713 | -0.4 |
| 4269 | AhLPPRKWKKX | 87 | 302.7 | 4 | 22.69 | 1206.771 | -0.3 |
| 4524 | GQcSSNKWKKX | 87 | 401.8979 | 3 | 23.32 | 1202.663 | 7.1 |

| | | | | | | | |
|---|---|---|---|---|---|---|---|
| 4577 | SPPEEaKWKKX | 87 | 409.8994 | 3 | 23.46 | 1226.677 | -0.4 |
| 5234 | GGLNAgKWKKX | 87 | 376.5731 | 3 | 25.15 | 1126.698 | -0.1 |
| 5502 | GGQSeDKWKKX | 87 | 410.5518 | 3 | 25.89 | 1228.635 | -1.3 |
| 5711 | AGAgFSKWKKX | 87 | 383.5691 | 3 | 26.46 | 1147.687 | -0.9 |
| 6823 | ANfSgaKWKKX | 87 | 437.5955 | 3 | 29.76 | 1309.766 | -0.9 |
| 6931 | ANGAfFKWKKX | 87 | 318.6814 | 4 | 30.11 | 1270.698 | -0.9 |
| 7642 | NLPNSeKWKKX | 87 | 437.5837 | 3 | 32.65 | 1309.73 | -0.2 |
| 8038 | NGDDPeKWKKX | 87 | 428.5559 | 3 | 34.06 | 1282.646 | 0.1 |
| 8040 | NGDDPeKWKKX | 87 | 428.5559 | 3 | 34.07 | 1282.646 | 0.1 |
| 8048 | NGDDPeKWKKX | 87 | 428.5559 | 3 | 34.09 | 1282.646 | 0 |
| 8050 | NGDDPeKWKKX | 87 | 428.5559 | 3 | 34.09 | 1282.646 | 0 |
| 3842 | QAFRNdKWKKX | 86 | 345.9555 | 4 | 21.62 | 1379.794 | -0.5 |
| 4151 | GSgDGGKWKKX | 86 | 363.545 | 3 | 22.4 | 1087.614 | -0.4 |
| 4232 | FTKNTFKWKKX | 86 | 442.9276 | 3 | 22.61 | 1325.761 | 0.3 |
| 4248 | AhLPPRKWKKX | 86 | 302.7 | 4 | 22.64 | 1206.771 | -0.1 |
| 4271 | AhLPPRKWKKX | 86 | 302.7 | 4 | 22.7 | 1206.771 | -0.3 |
| 4504 | hPLPPGKWKKX | 86 | 284.434 | 4 | 23.28 | 1133.707 | -0.4 |
| 4777 | hGGGRfKWKKX | 86 | 306.6844 | 4 | 23.97 | 1222.709 | -0.3 |
| 4870 | ATQgEDKWKKX | 86 | 315.683 | 4 | 24.21 | 1258.703 | -0.5 |
| 4908 | QaDbQGKWKKX | 86 | 414.9068 | 3 | 24.3 | 1241.699 | -0.4 |
| 5922 | hAKATSKWKKX | 86 | 377.9059 | 3 | 27.06 | 1130.692 | 3 |
| 6770 | ANfSgaKWKKX | 86 | 437.5955 | 3 | 29.59 | 1309.766 | -0.9 |
| 6776 | ANfSgaKWKKX | 86 | 437.5954 | 3 | 29.61 | 1309.766 | -1.2 |
| 7315 | GLEHAaKWKKX | 86 | 399.2378 | 3 | 31.46 | 1194.699 | -5.9 |
| 8026 | NGDDPeKWKKX | 86 | 428.556 | 3 | 34.03 | 1282.646 | 0.4 |
| 8068 | NGDDPeKWKKX | 86 | 428.5559 | 3 | 34.14 | 1282.646 | 0.2 |
| 8070 | NGDDPeKWKKX | 86 | 428.5559 | 3 | 34.15 | 1282.646 | 0.2 |
| 3905 | hSANdFKWKKX | 85 | 317.9369 | 4 | 21.77 | 1267.719 | -0.3 |
| 4497 | DgAGGEKWKKX | 85 | 382.2205 | 3 | 23.26 | 1143.64 | -0.4 |
| 5194 | NGfGAGKWKKX | 85 | 389.8856 | 3 | 25.05 | 1166.635 | 0.2 |
| 5518 | AADGfNKWKKX | 85 | 413.8917 | 3 | 25.94 | 1238.656 | -2.3 |
| 5522 | NGQSeDKWKKX | 85 | 429.5584 | 3 | 25.95 | 1285.657 | -2.6 |
| 5528 | AADGfNKWKKX | 85 | 413.8925 | 3 | 25.96 | 1238.656 | -0.3 |
| 5565 | hGDGfNKWKKX | 85 | 310.6712 | 4 | 26.06 | 1238.656 | -0.1 |
| 6171 | TPDGTGKWKKX | 85 | 372.8768 | 3 | 27.79 | 1115.609 | -0.1 |
| 6769 | ANfSgaKWKKX | 85 | 437.5955 | 3 | 29.59 | 1309.766 | -0.9 |
| 8017 | NGDDPeKWKKX | 85 | 428.5561 | 3 | 34 | 1282.646 | 0.5 |
| 8019 | NGDDPeKWKKX | 85 | 428.5561 | 3 | 34.01 | 1282.646 | 0.5 |
| 8031 | NDGDPeKWKKX | 85 | 428.5561 | 3 | 34.04 | 1282.646 | 0.5 |
| 8033 | NDGDPeKWKKX | 85 | 428.5561 | 3 | 34.05 | 1282.646 | 0.5 |
| 8323 | PTFEAGKWKKX | 85 | 397.5608 | 3 | 34.89 | 1189.661 | 0 |
| 3981 | KANFDDKWKKX | 84 | 320.4291 | 4 | 21.97 | 1277.688 | -0.6 |
| 3994 | QEKDgGKWKKX | 84 | 318.9409 | 4 | 22.01 | 1271.735 | -0.4 |

| | | | | | | | |
|---|---|---|---|---|---|---|---|
| 4197 | GGfhKSKWKKX | 84 | 409.2451 | 3 | 22.52 | 1224.713 | 0.2 |
| 4216 | GfAGHAKWKKX | 84 | 402.2294 | 3 | 22.57 | 1203.667 | 0 |
| 4503 | ALRgGAKWKKX | 84 | 296.7001 | 4 | 23.27 | 1182.771 | 0 |
| 4760 | hGGGRfKWKKX | 84 | 306.6844 | 4 | 23.93 | 1222.709 | -0.2 |
| 5606 | AhGgHgKWKKX | 84 | 298.9472 | 4 | 26.17 | 1191.761 | -0.6 |
| 5729 | AGAgFSKWKKX | 84 | 383.5692 | 3 | 26.51 | 1147.687 | -0.8 |
| 6787 | ANfSgaKWKKX | 84 | 437.5951 | 3 | 29.64 | 1309.766 | -1.9 |
| 6989 | RASPFEKWKKX | 84 | 425.9154 | 3 | 30.28 | 1274.725 | -0.2 |
| 6991 | RASPFEKWKKX | 84 | 425.9154 | 3 | 30.28 | 1274.725 | -0.2 |
| 7727 | iDDEhSKWKKX | 84 | 424.8837 | 3 | 32.92 | 1271.626 | 2.5 |
| 3983 | KANFDDKWKKX | 83 | 320.4291 | 4 | 21.98 | 1277.688 | -0.6 |
| 3989 | NKAFDDKWKKX | 83 | 426.9031 | 3 | 21.99 | 1277.688 | -0.3 |
| 4865 | hADbEHKWKKX | 83 | 417.9033 | 3 | 24.2 | 1250.689 | -0.4 |
| 4867 | hADbEHKWKKX | 83 | 417.9033 | 3 | 24.2 | 1250.689 | -0.4 |
| 5381 | aTDGTGKWKKX | 83 | 373.8811 | 3 | 25.56 | 1118.62 | 1.7 |
| 5510 | GGQSeDKWKKX | 83 | 410.5515 | 3 | 25.92 | 1228.635 | -2.2 |
| 5548 | AADGfNKWKKX | 83 | 413.8925 | 3 | 26.01 | 1238.656 | -0.2 |
| 5978 | DQgGGLKWKKX | 83 | 395.9082 | 3 | 27.23 | 1184.703 | -0.1 |
| 6609 | PQGeEDKWKKX | 83 | 437.8997 | 3 | 29.08 | 1310.677 | 0.1 |
| 6765 | ANfSgaKWKKX | 83 | 437.5955 | 3 | 29.57 | 1309.766 | -0.9 |
| 6804 | ANfSgaKWKKX | 83 | 437.5952 | 3 | 29.7 | 1309.766 | -1.7 |
| 7329 | GLEHAaKWKKX | 83 | 399.2377 | 3 | 31.5 | 1194.699 | -6 |
| 12734 | PTgSDeKWKKX | 83 | 438.2514 | 3 | 49.85 | 1311.734 | -1.1 |
| 4250 | AhLPPRKWKKX | 82 | 302.7 | 4 | 22.65 | 1206.771 | -0.1 |
| 4526 | GQcSSNKWKKX | 82 | 401.8979 | 3 | 23.33 | 1202.663 | 7.1 |
| 4626 | PSNDEbKWKKX | 82 | 419.5629 | 3 | 23.58 | 1255.667 | -0.4 |
| 4691 | GNFGELKWKKX | 82 | 402.5644 | 3 | 23.74 | 1204.672 | -0.3 |
| 4856 | ATQgEDKWKKX | 82 | 420.5751 | 3 | 24.17 | 1258.703 | 0 |
| 5183 | GNfGAGKWKKX | 82 | 389.8856 | 3 | 25.02 | 1166.635 | 0.2 |
| 5512 | AADGfNKWKKX | 82 | 310.6705 | 4 | 25.92 | 1238.656 | -2.6 |
| 5525 | AADGfNKWKKX | 82 | 310.6711 | 4 | 25.95 | 1238.656 | -0.5 |
| 5590 | AADGfNKWKKX | 82 | 413.8925 | 3 | 26.13 | 1238.656 | -0.2 |
| 5793 | AhDgdLKWKKX | 82 | 425.936 | 3 | 26.69 | 1274.786 | -0.2 |
| 6119 | GSgFEQKWKKX | 82 | 421.9114 | 3 | 27.64 | 1262.714 | -1 |
| 6951 | AGGQfFKWKKX | 82 | 424.5732 | 3 | 30.17 | 1270.698 | 0.2 |
| 7011 | NaaPFEKWKKX | 82 | 425.9156 | 3 | 30.34 | 1274.725 | 0.2 |
| 3892 | GRETgNKWKKX | 81 | 424.9226 | 3 | 21.74 | 1271.746 | -0.3 |
| 3958 | NAGNLRKWKKX | 81 | 304.1873 | 4 | 21.91 | 1212.72 | 0 |
| 4241 | AhLRPPKWKKX | 81 | 302.7001 | 4 | 22.63 | 1206.771 | 0 |
| 4387 | GAQdeQKWKKX | 81 | 337.1936 | 4 | 22.99 | 1344.745 | 0 |
| 4734 | gLESTNKWKKX | 81 | 420.5873 | 3 | 23.86 | 1258.74 | 0.2 |
| 4910 | QaDbQGKWKKX | 81 | 414.9068 | 3 | 24.31 | 1241.699 | -0.4 |
| 5714 | AGAgFSKWKKX | 81 | 287.9286 | 4 | 26.47 | 1147.687 | -1 |

| | | | | | | | |
|---|---|---|---|---|---|---|---|
| 5760 | AhDgdLKWKKX | 81 | 425.9354 | 3 | 26.59 | 1274.786 | -1.6 |
| 6377 | FDSAgbKWKKX | 81 | 421.252 | 3 | 28.4 | 1260.734 | -0.2 |
| 6812 | ANfSgaKWKKX | 81 | 437.5958 | 3 | 29.72 | 1309.766 | -0.2 |
| 3861 | cEGSaFKWKKX | 80 | 313.433 | 4 | 21.66 | 1249.704 | -1 |
| 3877 | GRETgNKWKKX | 80 | 318.9434 | 4 | 21.7 | 1271.746 | -1.4 |
| 3941 | GGNPgaKWKKX | 80 | 285.9302 | 4 | 21.87 | 1139.693 | -0.9 |
| 4800 | hGGGRfKWKKX | 80 | 306.6844 | 4 | 24.02 | 1222.709 | -0.2 |
| 4958 | GGGLPDKWKKX | 80 | 362.2136 | 3 | 24.43 | 1083.619 | 0.1 |
| 5213 | GGGfQGKWKKX | 80 | 389.8856 | 3 | 25.1 | 1166.635 | 0.2 |
| 5732 | AGAgFSKWKKX | 80 | 287.9287 | 4 | 26.52 | 1147.687 | -0.7 |
| 6135 | GSgFEQKWKKX | 80 | 421.9113 | 3 | 27.69 | 1262.714 | -1.1 |
| 7653 | bTPNSeKWKKX | 80 | 437.5838 | 3 | 32.69 | 1309.73 | 0 |
| 3615 | AhEbRLKWKKX | 79 | 317.9541 | 4 | 21.06 | 1267.788 | -0.4 |
| 4279 | LhEQEdKWKKX | 79 | 337.9487 | 4 | 22.72 | 1347.766 | -0.5 |
| 4987 | GGGLPDKWKKX | 79 | 362.2134 | 3 | 24.51 | 1083.619 | -0.5 |
| 5102 | FgEASTKWKKX | 79 | 417.5759 | 3 | 24.81 | 1249.718 | -9.9 |
| 5237 | GGGfQGKWKKX | 79 | 389.8855 | 3 | 25.16 | 1166.635 | -0.1 |
| 5717 | DfSAGAKWKKX | 79 | 404.8885 | 3 | 26.48 | 1211.645 | -1 |
| 5846 | DSFNKFKWKKX | 79 | 442.9151 | 3 | 26.84 | 1325.724 | -0.7 |
| 6379 | FDSAgbKWKKX | 79 | 421.252 | 3 | 28.41 | 1260.734 | -0.2 |
| 6393 | FDSAgbKWKKX | 79 | 421.252 | 3 | 28.45 | 1260.734 | -0.1 |
| 3807 | AGAgARKWKKX | 78 | 381.2484 | 3 | 21.53 | 1140.724 | -0.8 |
| 4040 | hbNRSeKWKKX | 78 | 339.203 | 4 | 22.12 | 1352.783 | 0 |
| 4066 | GGNhFQKWKKX | 78 | 392.8929 | 3 | 22.19 | 1175.657 | 0.3 |
| 4299 | LhEQEdKWKKX | 78 | 337.9487 | 4 | 22.77 | 1347.766 | -0.6 |
| 4971 | GGGLPDKWKKX | 78 | 362.2136 | 3 | 24.47 | 1083.619 | 0 |
| 5545 | AADGfNKWKKX | 78 | 310.6711 | 4 | 26 | 1238.656 | -0.5 |
| 6081 | GNNGfNKWKKX | 78 | 423.2276 | 3 | 27.53 | 1266.662 | -0.9 |
| 7300 | GbDLANKWKKX | 78 | 395.569 | 3 | 31.4 | 1183.683 | 2.2 |
| 8081 | NDGDPeKWKKX | 78 | 428.5557 | 3 | 34.18 | 1282.646 | -0.5 |
| 3975 | KPPSbQKWKKX | 77 | 313.6975 | 4 | 21.96 | 1250.761 | -0.2 |
| 4012 | GAFGGTKWKKX | 77 | 360.21 | 3 | 22.05 | 1077.608 | -0.2 |
| 4026 | QSKgFaKWKKX | 77 | 327.2094 | 4 | 22.09 | 1304.808 | 0.2 |
| 4399 | DAGhTFKWKKX | 77 | 388.8889 | 3 | 23.02 | 1163.645 | -0.4 |
| 4572 | GGGeDaKWKKX | 77 | 391.217 | 3 | 23.44 | 1170.63 | -0.4 |
| 4907 | QaDbQGKWKKX | 77 | 311.432 | 4 | 24.3 | 1241.699 | -0.1 |
| 5255 | GGLNAgKWKKX | 77 | 376.5731 | 3 | 25.21 | 1126.698 | -0.1 |
| 5567 | AADGfNKWKKX | 77 | 413.8925 | 3 | 26.06 | 1238.656 | -0.2 |
| 5993 | DQgGGLKWKKX | 77 | 395.9079 | 3 | 27.27 | 1184.703 | -0.8 |
| 5999 | GNEeNDKWKKX | 77 | 438.8909 | 3 | 27.28 | 1313.652 | -0.6 |
| 6097 | GNNGfNKWKKX | 77 | 423.2277 | 3 | 27.58 | 1266.662 | -0.6 |
| 6153 | GSgFEQKWKKX | 77 | 421.9116 | 3 | 27.74 | 1262.714 | -0.5 |
| 6187 | GSgFEQKWKKX | 77 | 421.9116 | 3 | 27.84 | 1262.714 | -0.5 |

| | | | | | | | |
|---|---|---|---|---|---|---|---|
| 6320 | GGLTDgKWKKX | 77 | 386.9044 | 3 | 28.23 | 1157.692 | -0.7 |
| 8083 | NDGDPeKWKKX | 77 | 428.5557 | 3 | 34.18 | 1282.646 | -0.5 |
| 3674 | ARLPGGKWKKX | 76 | 285.6844 | 4 | 21.2 | 1138.709 | -0.3 |
| 3863 | GcESaFKWKKX | 76 | 313.433 | 4 | 21.67 | 1249.704 | -1 |
| 4078 | GAGFNTKWKKX | 76 | 379.2173 | 3 | 22.22 | 1134.63 | 0 |
| 4084 | ESSTGFKWKKX | 76 | 399.5522 | 3 | 22.23 | 1195.635 | -0.2 |
| 4506 | hPLPPGKWKKX | 76 | 284.434 | 4 | 23.28 | 1133.707 | -0.4 |
| 4551 | ARgFGRKWKKX | 76 | 326.4622 | 4 | 23.39 | 1301.82 | 0.1 |
| 4563 | AFLDhAKWKKX | 76 | 298.4343 | 4 | 23.42 | 1189.697 | 9 |
| 5564 | AADGfNKWKKX | 76 | 310.6712 | 4 | 26.06 | 1238.656 | -0.1 |
| 6203 | GSgFEQKWKKX | 76 | 421.9117 | 3 | 27.88 | 1262.714 | -0.2 |
| 6371 | FDSAgbKWKKX | 76 | 421.2516 | 3 | 28.38 | 1260.734 | -1.1 |
| 7281 | LATeGGKWKKX | 76 | 395.5694 | 3 | 31.33 | 1183.687 | -0.1 |
| 9780 | AaeiDeKWKKX | 76 | 474.5873 | 3 | 40.03 | 1420.74 | -0.1 |
| 3819 | ANPLbQKWKKX | 75 | 310.1935 | 4 | 21.56 | 1236.746 | -0.7 |
| 3955 | hARPbQKWKKX | 75 | 313.6975 | 4 | 21.91 | 1250.773 | -9.1 |
| 4421 | DAGhTFKWKKX | 75 | 388.889 | 3 | 23.07 | 1163.645 | -0.1 |
| 5439 | aTGDTGKWKKX | 75 | 373.8771 | 3 | 25.72 | 1118.62 | -9.1 |
| 5589 | AADGfNKWKKX | 75 | 413.8925 | 3 | 26.12 | 1238.656 | -0.2 |
| 5751 | AGAgFSKWKKX | 75 | 383.5689 | 3 | 26.57 | 1147.687 | -1.4 |
| 5815 | LDHDLFKWKKX | 75 | 443.5868 | 3 | 26.75 | 1327.74 | -1.1 |
| 5848 | DSFNKFKWKKX | 75 | 442.9151 | 3 | 26.84 | 1325.724 | -0.7 |
| 8107 | GNDDPeKWKKX | 75 | 428.5554 | 3 | 34.25 | 1282.646 | -1.2 |
| 3977 | KPPSbQKWKKX | 74 | 313.6975 | 4 | 21.96 | 1250.761 | -0.2 |
| 4803 | GGLGFNKWKKX | 74 | 378.5574 | 3 | 24.03 | 1132.651 | -0.2 |
| 5670 | AhDgdLKWKKX | 74 | 425.9356 | 3 | 26.35 | 1274.786 | -1 |
| 5687 | AhDgdLKWKKX | 74 | 425.9355 | 3 | 26.39 | 1274.786 | -1.3 |
| 8109 | GNDDPeKWKKX | 74 | 428.5554 | 3 | 34.26 | 1282.646 | -1.2 |
| 8307 | KAmDAKKWKKX | 74 | 431.9322 | 3 | 34.84 | 1292.78 | -4.3 |
| 8309 | KAmDAKKWKKX | 74 | 431.9322 | 3 | 34.85 | 1292.78 | -4.3 |
| 12736 | PTgSDeKWKKX | 74 | 438.2514 | 3 | 49.86 | 1311.734 | -1.1 |
| 3653 | LAaFGTKWKKX | 73 | 393.2448 | 3 | 21.15 | 1176.713 | -0.4 |
| 3904 | hASNdFKWKKX | 73 | 317.9369 | 4 | 21.77 | 1267.719 | -0.3 |
| 4753 | gLESTNKWKKX | 73 | 420.5871 | 3 | 23.91 | 1258.74 | -0.2 |
| 4773 | aSNbLGKWKKX | 73 | 395.9133 | 3 | 23.96 | 1184.714 | 3.3 |
| 5441 | aTGDTGKWKKX | 73 | 373.8771 | 3 | 25.72 | 1118.62 | -9.1 |
| 5648 | AhDgdLKWKKX | 73 | 425.9356 | 3 | 26.29 | 1274.786 | -1.2 |
| 5694 | AhDgdLKWKKX | 73 | 425.9357 | 3 | 26.41 | 1274.786 | -0.9 |
| 6169 | GSgFEQKWKKX | 73 | 421.9118 | 3 | 27.79 | 1262.714 | 0 |
| 6565 | FaDhhfKWKKX | 73 | 448.5919 | 3 | 28.95 | 1342.755 | -1 |
| 4332 | NDATGFKWKKX | 72 | 398.5523 | 3 | 22.85 | 1192.635 | -0.1 |

| | | | | | | | |
|---|---|---|---|---|---|---|---|
| 4743 | LgESTNKWKKX | 72 | 315.6919 | 4 | 23.88 | 1258.74 | -1.1 |
| 4816 | aSNbLGKWKKX | 72 | 395.9133 | 3 | 24.07 | 1184.714 | 3.4 |
| 6822 | ANfSgaKWKKX | 72 | 437.5955 | 3 | 29.76 | 1309.766 | -0.9 |
| 6924 | AGGQfFKWKKX | 72 | 424.5732 | 3 | 30.09 | 1270.698 | 0.2 |
| 8702 | kaGhFGKWKKX | 72 | 416.5742 | 3 | 36.25 | 1246.707 | -5.1 |
| 3551 | ARLREQKWKKX | 71 | 336.211 | 4 | 20.9 | 1340.815 | -0.2 |
| 3714 | GbcSTQKWKKX | 71 | 308.1833 | 4 | 21.3 | 1228.715 | -9.2 |
| 3920 | KDGGdFKWKKX | 71 | 317.9369 | 4 | 21.8 | 1267.719 | -0.3 |
| 4050 | GGAFNTKWKKX | 71 | 379.2172 | 3 | 22.15 | 1134.63 | 0 |
| 4517 | DgAGGEKWKKX | 71 | 382.2206 | 3 | 23.31 | 1143.64 | 0 |
| 4714 | AhNRNGKWKKX | 71 | 395.9001 | 3 | 23.81 | 1184.689 | -9 |
| 5066 | FgESATKWKKX | 71 | 417.5798 | 3 | 24.72 | 1249.718 | -0.5 |
| 5274 | NLGGAgKWKKX | 71 | 376.5729 | 3 | 25.26 | 1126.698 | -0.5 |
| 6122 | PGEEGeKWKKX | 71 | 418.8921 | 3 | 27.65 | 1253.656 | -0.9 |
| 6162 | PGEEGeKWKKX | 71 | 418.8925 | 3 | 27.76 | 1253.656 | 0.2 |
| 6908 | AGGQfFKWKKX | 71 | 424.5732 | 3 | 30.04 | 1270.698 | 0.1 |
| 6937 | AGGQfFKWKKX | 71 | 424.5728 | 3 | 30.13 | 1270.698 | -0.8 |
| 6950 | AGGQfFKWKKX | 71 | 424.5732 | 3 | 30.17 | 1270.698 | 0.2 |
| 6963 | AGGQfFKWKKX | 71 | 424.573 | 3 | 30.2 | 1270.698 | -0.3 |
| 8633 | hdHGFGKWKKX | 71 | 416.5744 | 3 | 36.03 | 1246.709 | -5.8 |
| 3541 | ARLREQKWKKX | 70 | 336.211 | 4 | 20.87 | 1340.815 | -0.1 |
| 3578 | ARLREQKWKKX | 70 | 336.2111 | 4 | 20.96 | 1340.815 | 0 |
| 3735 | hagTGhKWKKX | 70 | 286.6893 | 4 | 21.35 | 1142.729 | -0.8 |
| 4209 | LRLhNDKWKKX | 70 | 321.9529 | 4 | 22.55 | 1283.783 | -0.1 |
| 4642 | PGAGhgKWKKX | 70 | 361.5659 | 3 | 23.62 | 1081.676 | -0.1 |
| 4840 | aSNbLGKWKKX | 70 | 395.9131 | 3 | 24.13 | 1184.714 | 2.9 |
| 4847 | ATQgEDKWKKX | 70 | 315.683 | 4 | 24.15 | 1258.703 | -0.2 |
| 5082 | FgESATKWKKX | 70 | 417.58 | 3 | 24.76 | 1249.718 | -0.1 |
| 5509 | GGQSeDKWKKX | 70 | 410.5515 | 3 | 25.91 | 1228.635 | -2.2 |
| 5583 | AAAgHgKWKKX | 70 | 298.9472 | 4 | 26.11 | 1191.761 | -0.7 |
| 5659 | ALAATgKWKKX | 70 | 286.4404 | 4 | 26.32 | 1141.734 | -0.9 |
| 5735 | DfSGGhKWKKX | 70 | 404.8886 | 3 | 26.53 | 1211.645 | -0.9 |
| 6984 | hcSPFEKWKKX | 70 | 425.9154 | 3 | 30.26 | 1274.725 | -0.4 |
| 7024 | hcSPFEKWKKX | 70 | 425.9155 | 3 | 30.38 | 1274.725 | -0.1 |
| 7821 | AGAELfKWKKX | 70 | 418.2449 | 3 | 33.31 | 1251.713 | 0.1 |
| 7832 | AGAELfKWKKX | 70 | 418.2448 | 3 | 33.34 | 1251.713 | -0.1 |
| 7839 | AGAELfKWKKX | 70 | 418.2449 | 3 | 33.37 | 1251.713 | 0.1 |
| 8372 | ATfNQeKWKKX | 70 | 474.9277 | 3 | 35.03 | 1421.761 | 0.4 |
| 9831 | KiiSDeKWKKX | 70 | 474.588 | 3 | 40.19 | 1420.736 | 4.1 |

### Appendix IV. Peptides identified from experimental samples

#### Whole cell extract, PMO-Library

| Scan | Peptide | ALC (%) | m/z | z | RT | Mass | ppm |
|---|---|---|---|---|---|---|---|
| 3025 | hPPKWKKX | 95 | 289.8567 | 3 | 17.64 | 866.5491 | -1.1 |
| 3055 | hPPKWKKX | 95 | 289.8566 | 3 | 17.79 | 866.5491 | -1.2 |
| 3038 | PPhKWKKX | 94 | 289.8566 | 3 | 17.71 | 866.5491 | -1.2 |
| 3027 | hPPKWKKX | 93 | 289.8567 | 3 | 17.65 | 866.5491 | -1.1 |
| 3040 | PPhKWKKX | 93 | 289.8566 | 3 | 17.72 | 866.5491 | -1.2 |
| 3057 | hPPKWKKX | 92 | 289.8566 | 3 | 17.8 | 866.5491 | -1.2 |
| 2634 | LTERKWKKX | 92 | 363.2253 | 3 | 15.78 | 1086.666 | -11.3 |
| 3045 | hPPKWKKX | 91 | 289.8565 | 3 | 17.74 | 866.5491 | -1.6 |
| 3047 | hPPKWKKX | 90 | 289.8565 | 3 | 17.75 | 866.5491 | -1.6 |
| 3077 | hPPKWKKX | 89 | 289.8564 | 3 | 17.89 | 866.5491 | -2.1 |
| 2630 | LTERKWKKX | 89 | 363.2252 | 3 | 15.76 | 1086.666 | -11.5 |
| 2641 | LTERKWKKX | 88 | 363.2254 | 3 | 15.81 | 1086.666 | -11 |
| 2781 | LPDETKWKKX | 87 | 381.8884 | 3 | 16.49 | 1142.645 | -1.3 |
| 2805 | LPDRjWKKX | 82 | 381.8882 | 3 | 16.59 | 1142.635 | 7.1 |
| 2812 | LPDETKWKKX | 81 | 381.8882 | 3 | 16.62 | 1142.645 | -1.8 |
| 2782 | LPDETKWKKX | 78 | 381.8884 | 3 | 16.49 | 1142.645 | -1.3 |
| 3085 | AhShgKWKKX | 78 | 348.5618 | 3 | 17.93 | 1042.665 | -1.7 |
| 2783 | KeETKWKKX | 76 | 381.8884 | 3 | 16.5 | 1142.66 | -14.6 |
| 2648 | LTERKKWKX | 75 | 363.2253 | 3 | 15.85 | 1086.666 | -11.1 |
| 2793 | LPDRWjKKX | 75 | 381.8884 | 3 | 16.54 | 1142.635 | 7.5 |
| 2759 | LPDETKWKKX | 75 | 381.8884 | 3 | 16.39 | 1142.645 | -1.2 |
| 2792 | LPDETKWKKX | 75 | 381.8884 | 3 | 16.54 | 1142.645 | -1.2 |
| 2760 | LPDETKWKKX | 74 | 381.8884 | 3 | 16.39 | 1142.645 | -1.2 |
| 2804 | LPDETKWKKX | 74 | 381.8882 | 3 | 16.59 | 1142.645 | -1.6 |
| 2631 | LTERKWKKX | 73 | 363.2252 | 3 | 15.77 | 1086.666 | -11.5 |
| 2621 | LTERKWKKX | 72 | 363.2255 | 3 | 15.72 | 1086.666 | -10.8 |
| 2635 | LTERKWKKX | 72 | 363.2253 | 3 | 15.78 | 1086.666 | -11.3 |
| 3078 | LDagKWKKX | 72 | 348.5616 | 3 | 17.9 | 1042.665 | -1.9 |
| 2392 | KDLQQKWKKX | 71 | 300.9354 | 4 | 14.6 | 1199.714 | -1.3 |
| 2394 | KDLQQKWKKX | 70 | 300.9354 | 4 | 14.61 | 1199.714 | -1.3 |
| 2813 | LPDETKWKKX | 69 | 381.8882 | 3 | 16.63 | 1142.645 | -1.8 |
| 2621 | LTEAhKWKKX | 68 | 363.2255 | 3 | 15.72 | 1086.655 | -0.4 |
| 2753 | LPDRjWKKX | 67 | 381.8884 | 3 | 16.36 | 1142.635 | 7.6 |
| 2615 | SbSGgKWKKX | 67 | 358.2254 | 3 | 15.69 | 1071.655 | -0.8 |
| 2635 | LTPTSKWKKX | 67 | 363.2253 | 3 | 15.78 | 1086.655 | -0.9 |
| 2631 | LTPTSKWKKX | 66 | 363.2252 | 3 | 15.77 | 1086.655 | -1.2 |
| 3087 | AhATgKWKKX | 65 | 348.5618 | 3 | 17.93 | 1042.665 | -1.7 |
| 3080 | LDagKWKKX | 64 | 348.5616 | 3 | 17.9 | 1042.665 | -1.9 |
| 2805 | LPDETKWKKX | 61 | 381.8882 | 3 | 16.59 | 1142.645 | -1.6 |

**Cytosolic extract, PMO-Library**

| Scan | Peptide | ALC (%) | m/z | z | RT | Mass | ppm |
|---|---|---|---|---|---|---|---|
| 3566 | ARRhaHKWKKX | 90 | 324.2084 | 4 | 21.38 | 1292.805 | -0.8 |
| 3584 | ARRhaHKWKKX | 89 | 324.2085 | 4 | 21.46 | 1292.805 | -0.3 |
| 3504 | ARhRaHKWKKX | 88 | 324.2084 | 4 | 21.11 | 1292.805 | -0.7 |
| 3568 | ARRhaHKWKKX | 88 | 324.2084 | 4 | 21.39 | 1292.805 | -0.8 |
| 3494 | ARhRaHKWKKX | 87 | 324.2085 | 4 | 21.07 | 1292.805 | -0.5 |
| 3586 | ARRhaHKWKKX | 87 | 324.2085 | 4 | 21.47 | 1292.805 | -0.3 |
| 3514 | LNgLTHKWKKX | 86 | 324.2084 | 4 | 21.15 | 1292.808 | -2.9 |
| 3496 | ARhRaHKWKKX | 85 | 324.2085 | 4 | 21.08 | 1292.805 | -0.5 |
| 3516 | LNgLTHKWKKX | 85 | 324.2084 | 4 | 21.16 | 1292.808 | -2.9 |
| 3506 | ARhRaHKWKKX | 84 | 324.2084 | 4 | 21.12 | 1292.805 | -0.7 |
| 3698 | ARhRaHKWKKX | 80 | 324.2086 | 4 | 21.97 | 1292.805 | -0.1 |
| 3579 | cNLNHKWKKX | 79 | 302.9305 | 4 | 21.44 | 1207.705 | -9.9 |
| 3611 | iHgKWKKX | 78 | 335.8711 | 3 | 21.58 | 1004.603 | -11.8 |
| 3612 | iHgKWKKX | 78 | 335.8711 | 3 | 21.58 | 1004.603 | -11.8 |
| 3700 | ARhRaHKWKKX | 76 | 324.2086 | 4 | 21.98 | 1292.805 | -0.1 |
| 3579 | RkNHKWKKX | 71 | 302.9305 | 4 | 21.44 | 1207.682 | 9.1 |
| 3574 | KAhRHKWKKX | 68 | 292.1911 | 4 | 21.42 | 1164.736 | -0.3 |
| 3579 | hGNLNHKWKKX | 67 | 302.9305 | 4 | 21.44 | 1207.694 | -0.6 |
| 3574 | KRRHKWKKX | 66 | 292.1911 | 4 | 21.42 | 1164.747 | -9.9 |

### Appendix V. LC-MS Characterization

Note: Chromatograms were obtained using Method A

**PMO-DBCO**
Mass Expected: 6500.0
Mass Observed: 6499.9

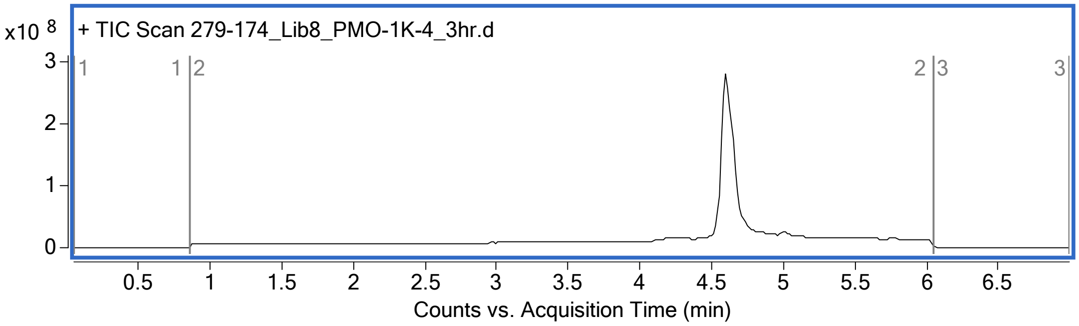

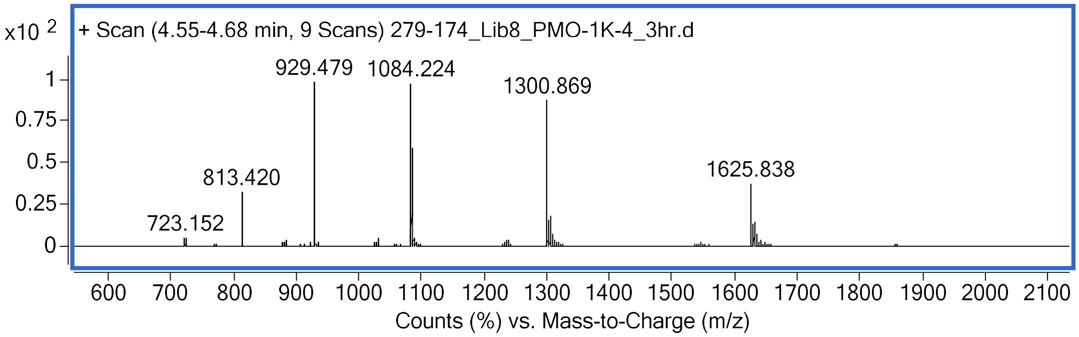

**PMO-Pep1a**
Mass Expected: 8617.3
Mass Observed: 8617.5
Peptide sequence: Biotin-Azidolysine-G-G-K-G-G-s-βal-r-r-abu-dab-h-k-w-k-k

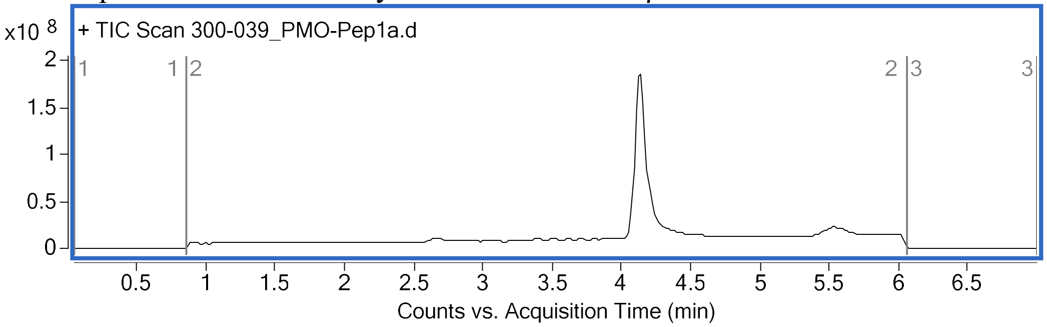

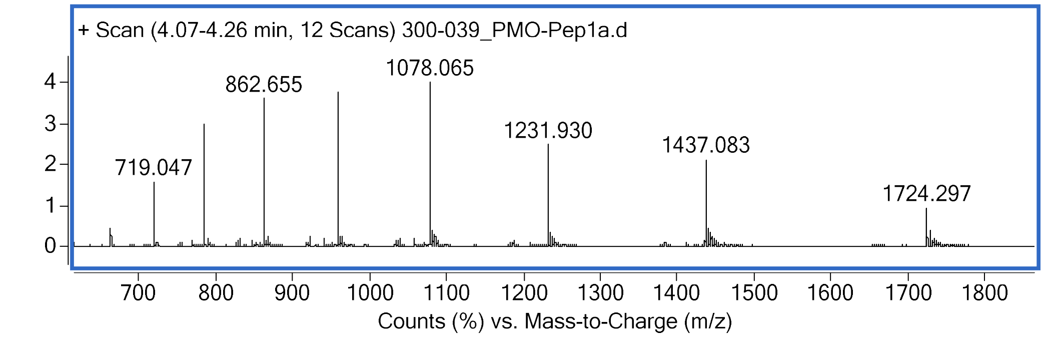

**PMO-Pep1b**
Mass Expected: 8532.4
Mass Observed: 8532.4
Peptide sequence: Biotin-Azidolysine-G-G-K-G-G-s-abu-G-n-nle-n-h-k-w-k-k

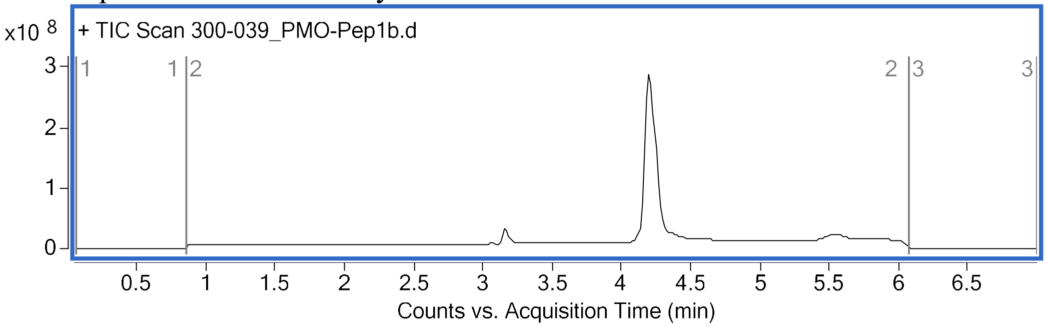

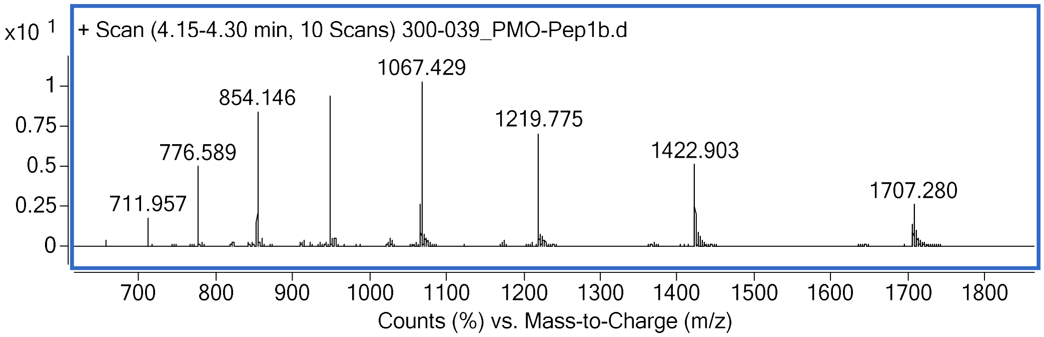

**PMO-Pep1c**
Mass Expected: 8467.3
Mass Observed: 8467.3
Peptide sequence: Biotin-Azidolysine-G-G-K-G-G-s-nle-p-d-e-t-k-w-k-k

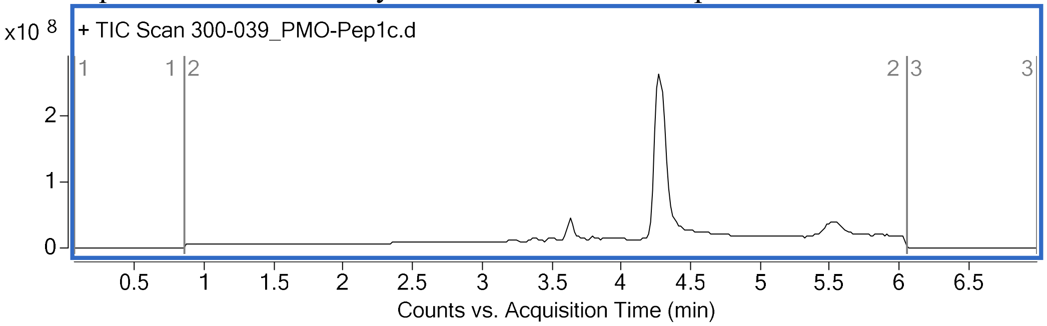

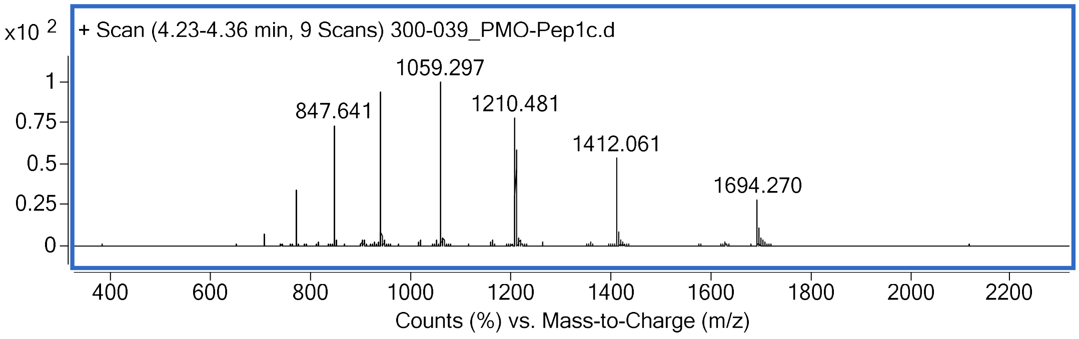

**PMO-Pep1d**
Mass Expected: 8367.2
Mass Observed: 8367.2
Peptide sequence: Biotin-Azidolysine- G-G-K-G-G-s-βal-abu-s-abu-hle-k-w-k-k

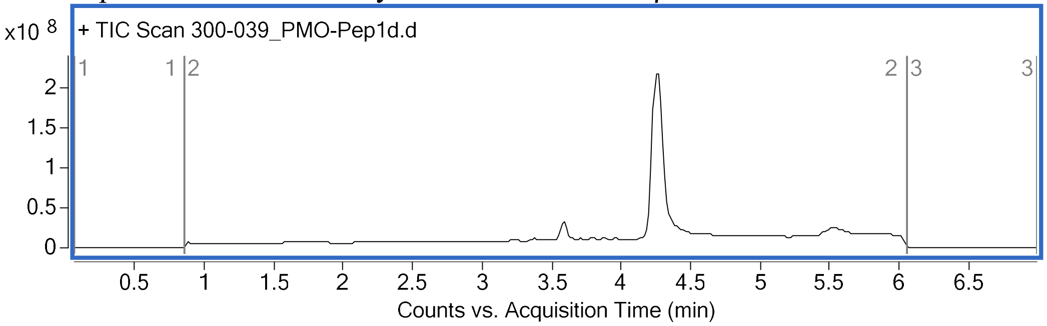

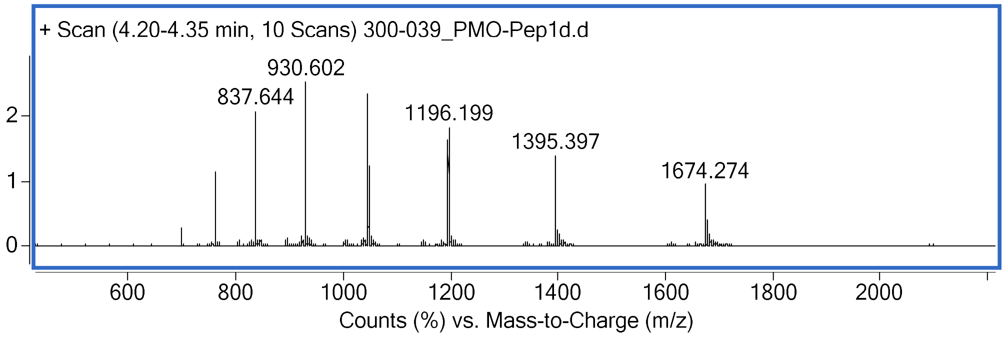

**PMO-library-peptide**
Mass Expected: 8437.1
Mass Observed: 8437.1
Peptide sequence: Biotin-Azidolysine-G-G-K-G-G-s-G-βal-n-d-p-βal-k-w-k-k

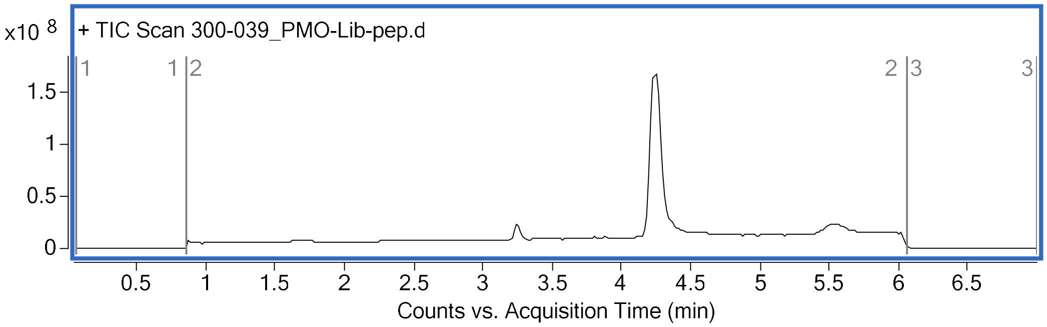

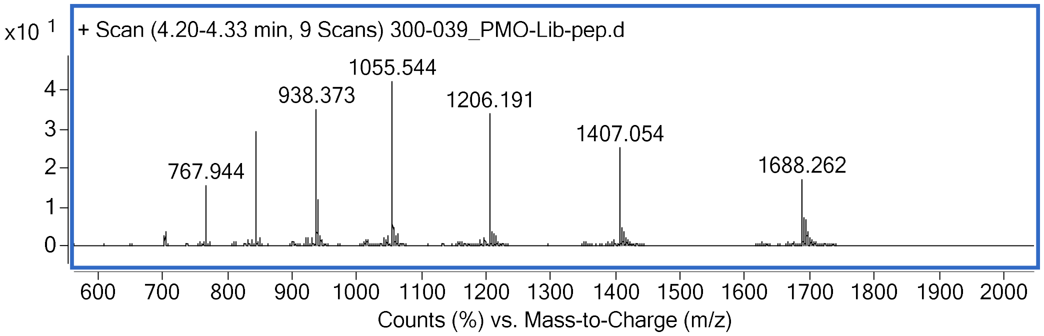

**PMO-D-Bpep**
Mass Expected: 8871.9
Mass Observed: 8871.8
Peptide sequence: Biotin-Azidolysine-G-G-K-G-G-r-ahx-r-r-βal-r-r-ahx-r-r-βal-r

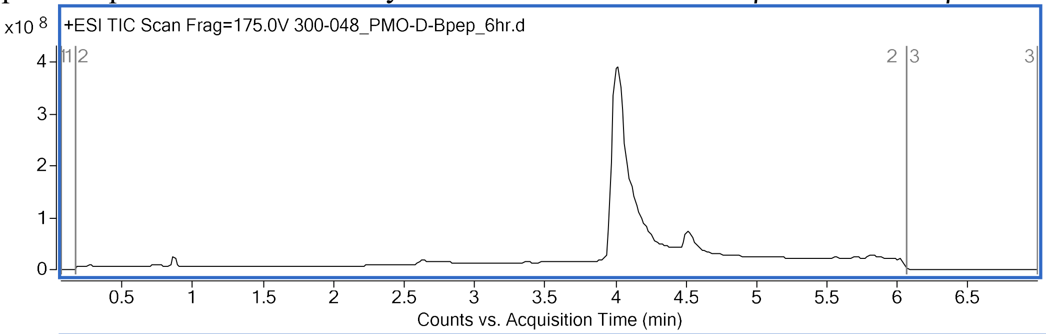

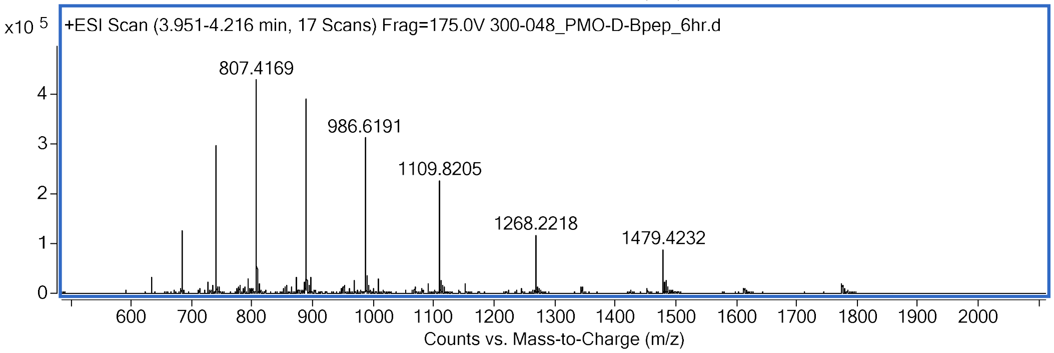

**PMO-Sulfo-Cy5-Pep1a**
Mass Expected: 9485.6
Mass Observed: 9485.7

Peptide sequence: Biotin-Azidolysine-(Sulfo-Cy5)c-G-G-K-G-G-s-βal-r-r-abu-dab-h-k-w-k-k

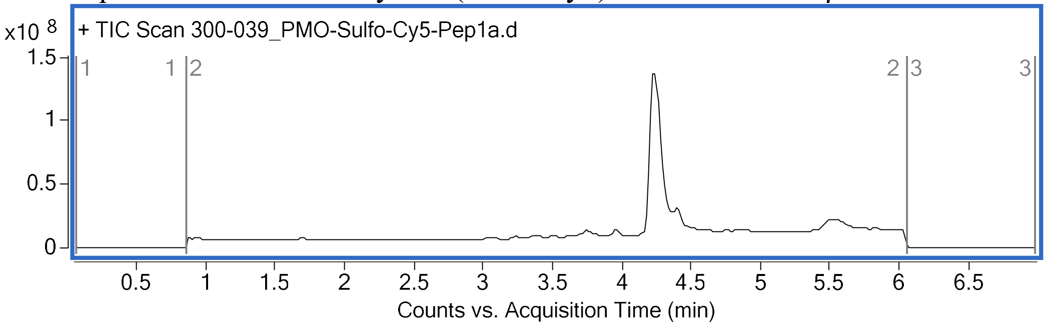

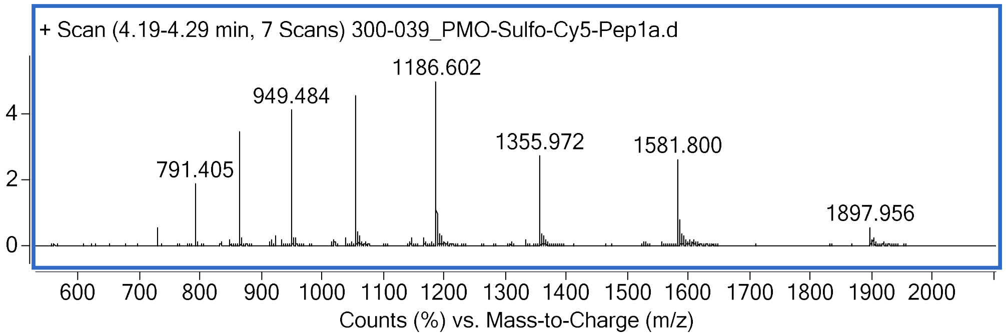

**PMO-Sulfo-Cy5-Pep1c**

Mass Expected:9335.3

Mass Observed: 9335.5

Peptide sequence: Biotin-Azidolysine-(Sulfo-Cy5)c- G-G-K-G-G-s-nle-p-d-e-t-k-w-k-k

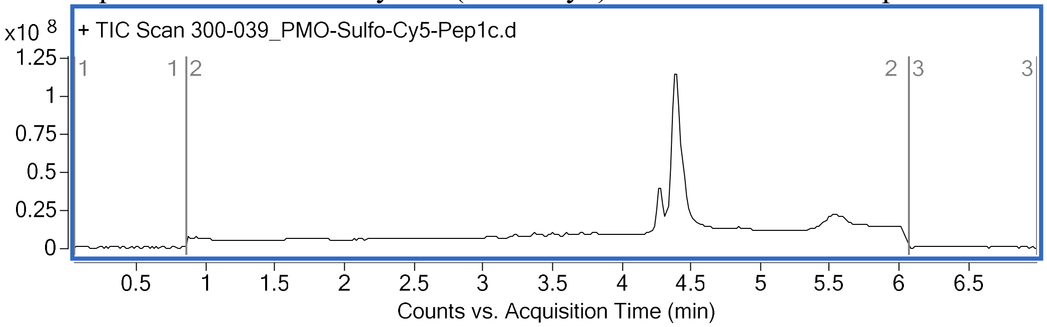

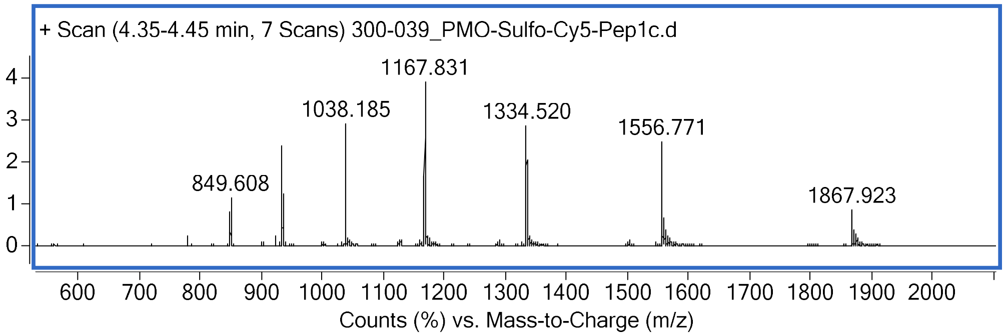

**PMO-Sulfo-Cy5-D-Bpep**
Mass Expected: 9739.9
Mass Observed: 9739.8
Peptide sequence: Biotin-Azidolysine-(Sulfo-Cy5)c-G-G-K-G-G-r-ahx-r-r-βal-r-r-ahx-r-r-βal-r

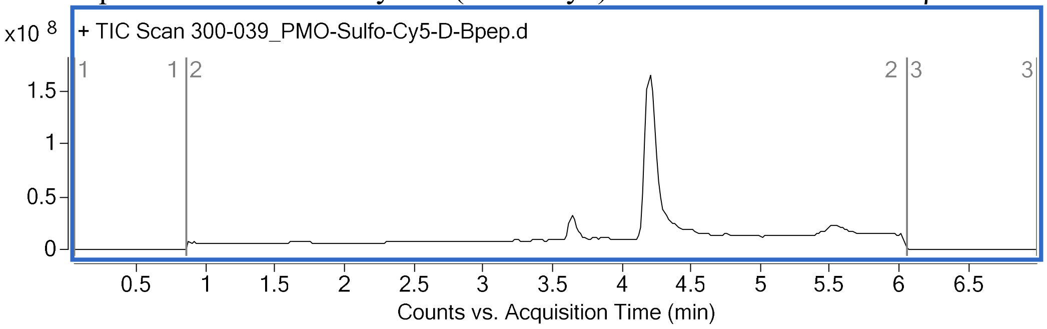

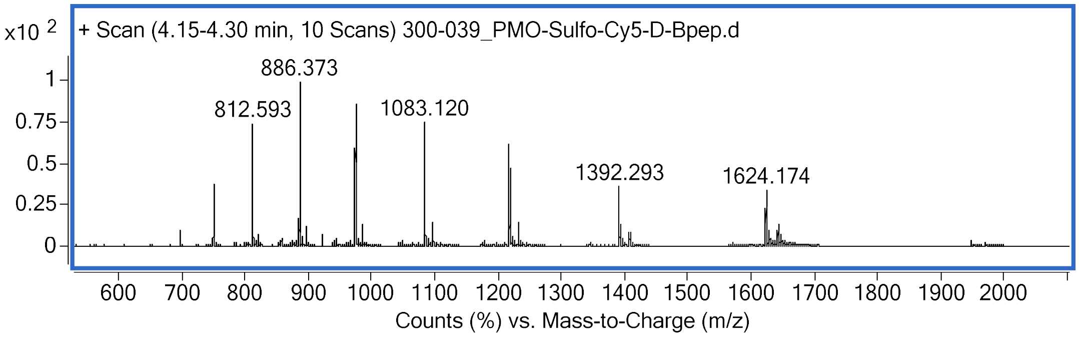

**DEAC-k5**
Mass Expected: 900.591
Mass Observed: 900.593
Peptide sequence: DEAC-k-k-k-k-k

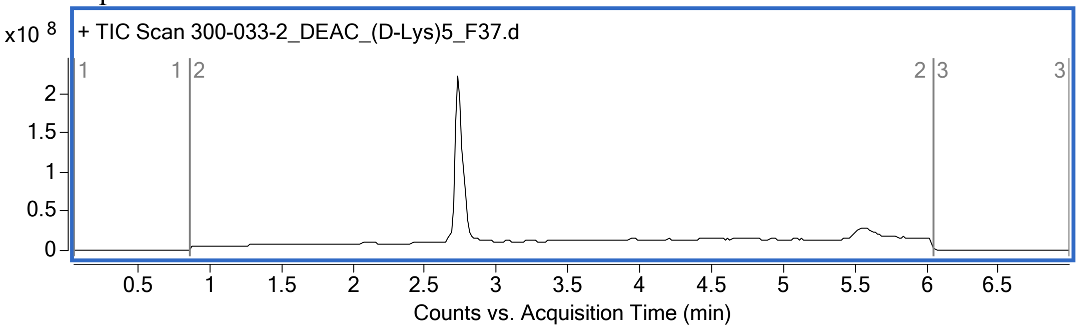

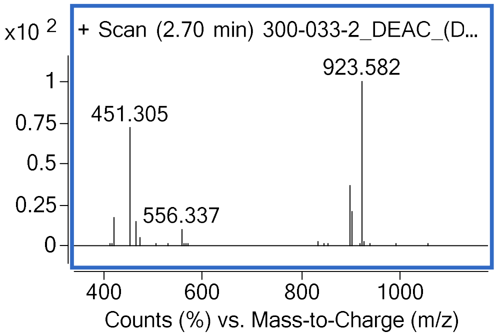

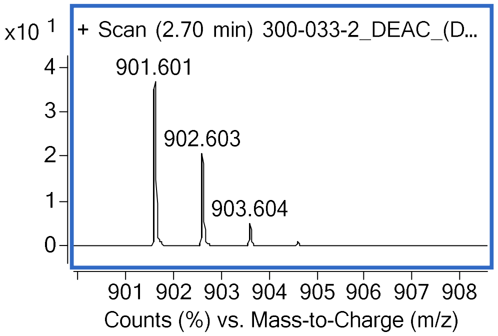

**PMO-Library (1,000 member)**

The libraries shown with greater than 1,000 members were produced by combining 1,000 member libraries. Two of the 1,000-member PMO-libraries are shown here.

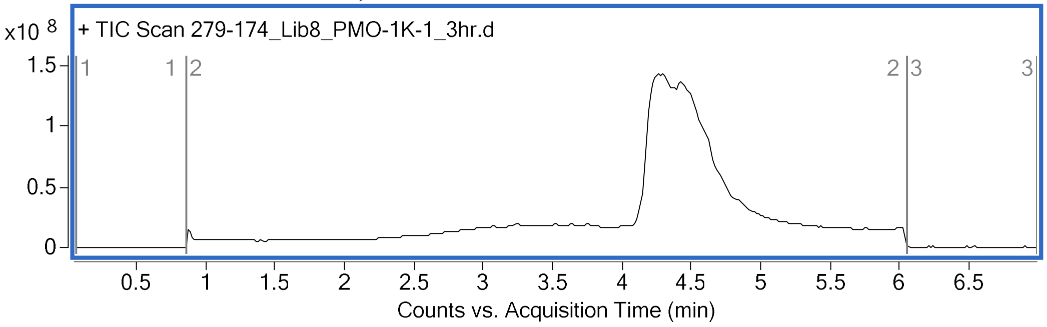

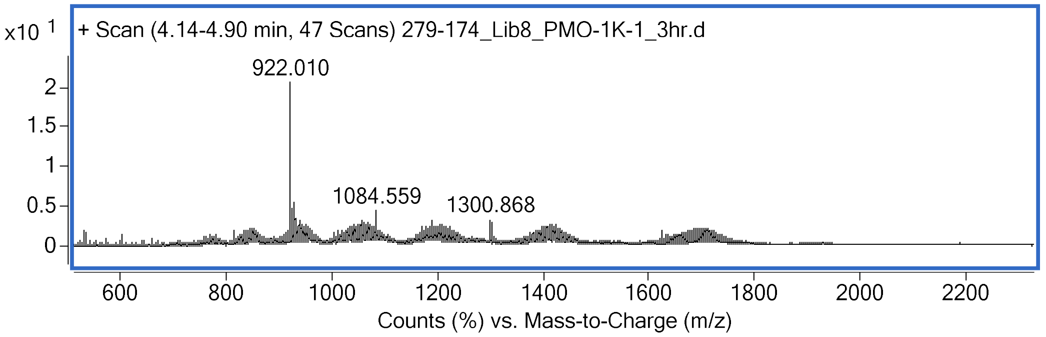

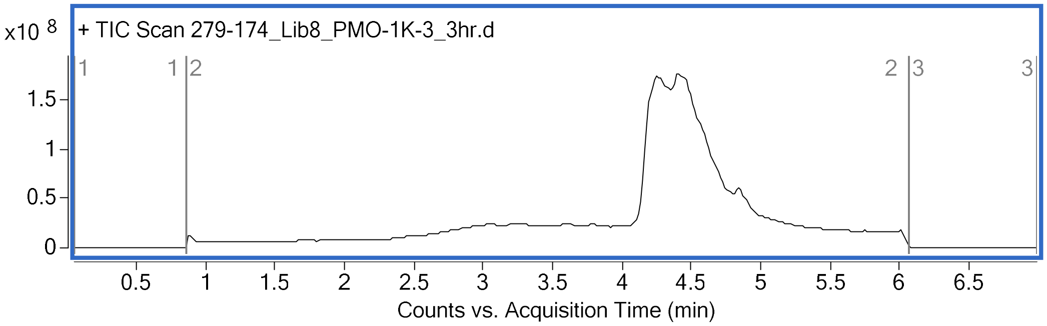

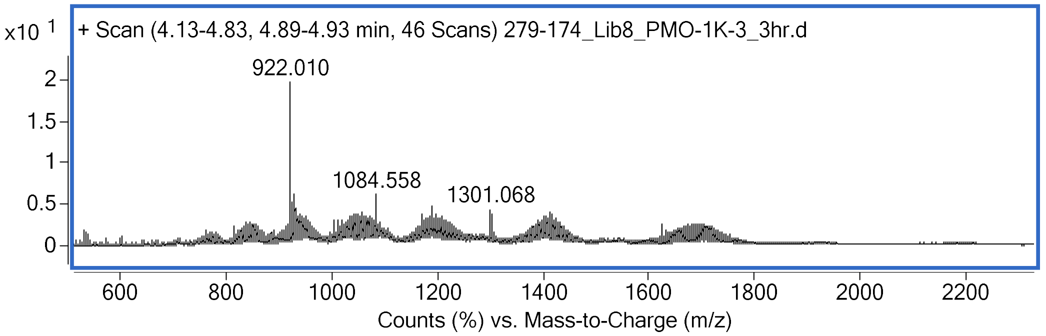
